# Time-resolved bottlenecks reveal concurrent pathogen entry and growth during bacterial infection dynamics

**DOI:** 10.1101/2025.11.21.689558

**Authors:** Sylvain Vicente, Nathanaël Boutillon, Raphael Forien, Isabelle Mila, Lionel Roques, Nemo Peeters

## Abstract

Within-host pathogen dynamics are shaped by successive host-imposed barriers that generate bottlenecks and ultimately determine infection outcome, yet these processes remain difficult to study *in vivo*. Here, we use neutral genetic barcoding to quantify the root-entry bottleneck during infection of tomato by the bacterial soil-borne phytopathogen *Ralstonia pseudosolanacearum*. We establish an updated theoretical framework showing that the temporal structure of pathogen entry, often overlooked, is a key determinant of root colonisation. We introduce the concept of time-resolved bottleneck, capturing infection scenarios in which entry occurs continuously while the pathogen population simultaneously expands within the host. We develop quantitative tools that exploit barcode dynamics to disentangle the respective roles of motility, growth, and spatial competition, enabling direct inference of *in vivo* pathogen fluxes and growth rates and linking host susceptibility to mechanistic features of colonisation. Extending beyond plants, reanalysis of two independent animal infection datasets reveals previously unrecognised within-host development patterns. Together, these results show that time-resolved bottleneck is a general principle of bacterial infection dynamics and propose a new standard for barcoding-based studies: a host- and pathogen-agnostic framework that is directly applicable to a majority of existing STAMP(-R)-style datasets, distributed as an open, annotated R/Python notebook to facilitate community reuse.

---

Understanding the spatiotemporal dynamics of pathogens within their hosts is essential to explaining infection success. Host–pathogen molecular interactions and the mechanisms governing tissue colonisation are ultimately manifested in the growth, spread, and structuring of pathogen populations *in vivo*. Despite major advances in elucidating the molecular determinants of bacterial virulence, quantitatively and descriptively capturing infection dynamics within the host remains challenging, limiting our ability to link molecular mechanisms to population-level outcomes. The STAMP (Sequence TagBased Analysis of Microbial Population dynamics) framework ^1^ and its recent extensions ^2,3^ have been effective in monitoring the dynamics of the infecting population and identifying bottlenecks in different host-bacterial pathosystems ^4^. The overarching goals of such approaches are to identify the locations of host-imposed barriers, quantify their restrictive strength against invading pathogens, and ultimately predict host survival or mortality under defined conditions.

STAMP experiments introduce neutral genetic diversity into pathogen populations by inserting short sequence tags into the genome. These tagged populations behave indistinguishably from wild-type strains, while the engineered neutral diversity acts as a heritable record of the population’s genetic history during infection. A primary application of this approach is the identification and quantification of infectious bottlenecks, during which pathogen populations undergo severe reductions in size that are revealed by characteristic losses of neutral genetic diversity. The bottleneck was originally defined as *number of founding pathogen cells* that cross the barrier and expand ^1^. Although many recent studies have now been published using this type of indicators (see review ^4^ and references within), the link between pathogen population patterns measured through STAMP and the nature of the underlying biological mechanisms so far remains unclear.

Here, we address this theoretical gap by applying STAMP to infections of tomato by the phytopathogen *Ralstonia pseudosolanacearum. R. pseudosolanacearum* is a soil-borne bacterium that causes bacterial wilt, a devastating disease of solanaceous crops. Infection initiates at the root, where the pathogen breaches host defences and rapidly colonises the xylem, leading to vascular occlusion, wilting, and ultimately plant death. Previous work has revealed the presence of localised host barriers along the infection route, including physical defences in the root, which impose strong constraints on pathogen entry and spread. ^5,6,7,8^.

We demonstrate that infection is associated with a stringent root-entry bottleneck, such that only a very small fraction of the invading pathogen population successfully establishes in the xylem. Although this bottleneck is detected by both the original STAMP framework ^1^ and its algorithmically refined implementation STAMPR ^2^, neither approach captures the fine-scale spatio-temporal patterns of root colonisation revealed by our data. By revisiting the theoretical basis of the classical definition of a bottleneck as a fixed “number of founding pathogen cells”, we show that this formulation is valid only under restrictive conditions. Critically, it fails to account for infection mechanisms such as motility, *in situ* growth, and spatial competition, which are central to host colonisation. Instead, our results support a scenario in which pathogen entry, proliferation, and dispersal occur concurrently. We formalise this as a *time-resolved bottleneck model*, in which bottlenecks emerge from explicitly time-dependent population dynamics.

To formalise this conceptual shift, we introduce a new analytical framework for neutral barcoding data. In addition to an *instantaneous bottleneck* model corresponding to the original STAMP framework ^1^, we develop three *time-resolved* models grounded in population genetics. Experimental datasets were systematically evaluated against each model to identify the infection regime that best explains the observed barcode dynamics, providing qualitative insight into the underlying mode of infection. Quantitative metrics can then be extracted to directly characterise the processes governing infection, including pathogen entry, growth, and competition.

Consistently, *time-resolved* models provided a markedly better fit to the plant infection data, indicating that pathogen populations begin to expand immediately upon host entry and that root barriers likely relax over the course of infection. The metrics derived from this framework explicitly capture the temporal dimension of infection, yielding quantitative estimates of pathogen influx into the host and *in planta* growth rates at the level of individual plants. Both parameters strongly correlate with host susceptibility, demonstrating that pathogen motility and growth dynamics are integral determinants of susceptibility rather than secondary consequences of infection.

To assess the generality of our framework, we reanalyse published animal infection datasets ^9,10^ and uncover analogous time-resolved dynamics in *Pseudomonas aeruginosa* and *Klebsiella pneumoniae* infections in mice. These reanalyses serve as a stringent benchmark for the performance of our approach and demonstrate its ability to infer pathogen influxes and *in situ* growth rates at the level of individual organs within individual hosts. Strikingly, this quantitative dissection reveals a previously overlooked trait in that *P. aeruginosa* colonisation is highly heterogeneous across the host, being driven by extensive dispersal among systemically connected organs (lung, spleen and liver), while colonisation of the gastrointestinal tract (gallbladder, stomach, small intestine, caecum and colon) is dominated by local population growth.

To maximise accessibility and reproducibility, models analysis and outputs are provided along with a fully annotated R computational document, enabling straightforward reuse, adaptation, and extension of the methodology for the analysis of neutral barcoding experiments across diverse host–pathogen systems.

## 1 Results

### 1.1 Physical barriers in plant roots impose a stringent pathogen-limiting bottleneck

To investigate the infection dynamics of *Ralstonia pseudosolanacearum*, we constructed a library of 119 neutral genetic tags in the reference strain GMI1000. Co-cultures of this tagged collection were used to implement the STAMP approach ^1^, enabling a quantitative analysis of the earliest stages of pathogen colonisation within the plant host (Fig. 1a). We first asked whether root defences impose a population-limiting barrier on *R. pseudosolanacearum*. We therefore formulated the following prediction: if a bottleneck occurs at root entry, tag diversity should decrease between inoculum and roots then remain stable once the pathogen has successfully colonised host tissues.

**Figure 1.**
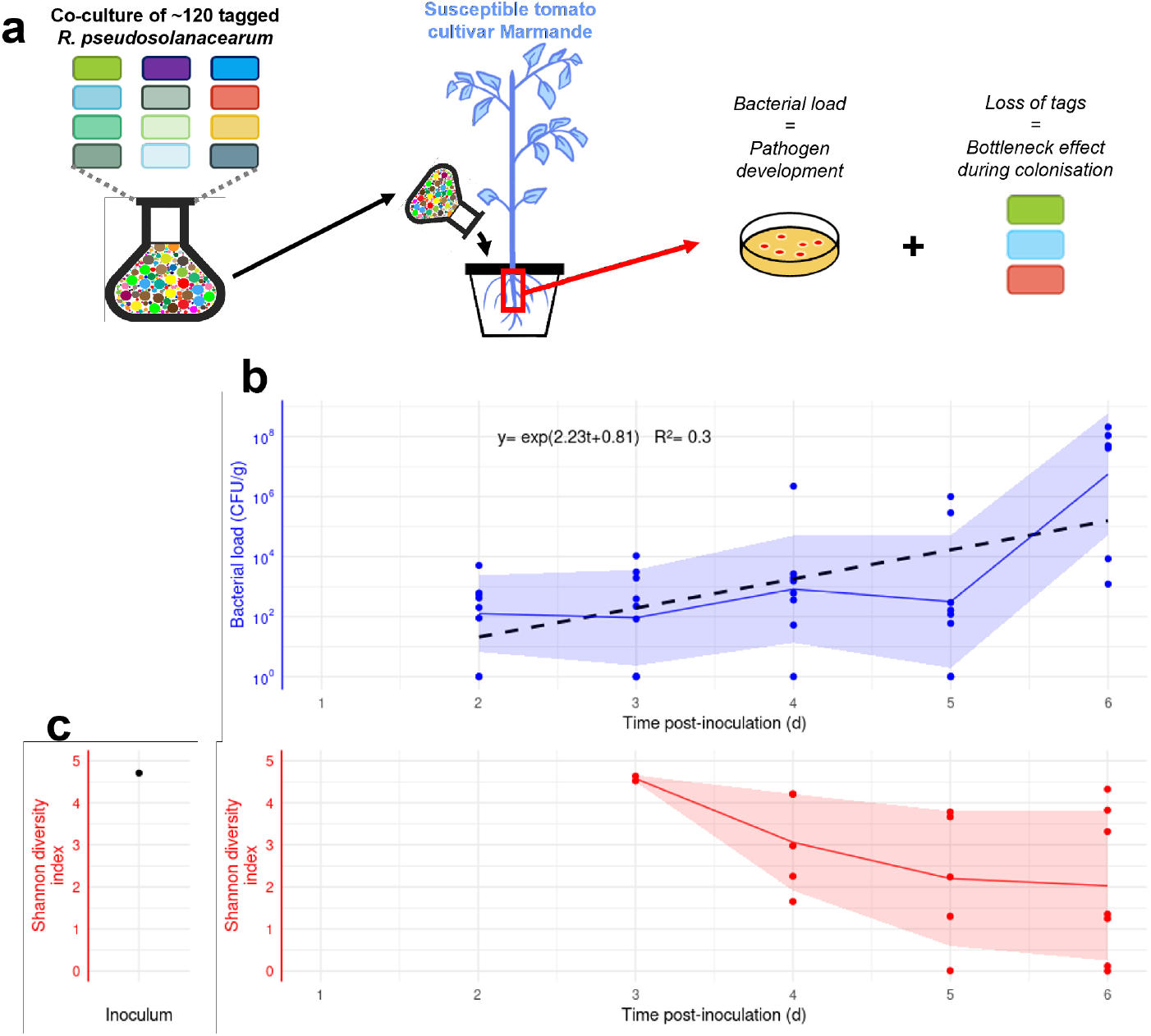
Roots of susceptible tomato (*cv*. Marmande) are infected by limited numbers of individual *R. pseudosolanacearum* cells. **a.** STAMP overview: 119 isogenic GMI1000 strains carrying a neutral 20 bp chromosomal barcode are generated; plants are soil-drenched with a co-culture of the tagged strains. Roots are sampled to measure bacterial load and tag diversity. Because tags are fitness-neutral, losses of diversity report bottlenecks. **b**. Root bacterial load from 2-6 DPI (log scale, *n* = 8-9 plants/day). Fit: *y* = *e*^*at*+*b*^ (black dotted line, equation). Blue line: local mean ± SD. **c**. Root Shannon diversity from 3-6 DPI; sequencing was feasible from 3 DPI onward when loads typically exceeded ∼ 10^4^ CFU/g. Red line: local mean ± SD. Diversity in inoculum: 4.71.

To test this hypothesis, susceptible tomato plants (*cv*. Marmande) were inoculated with the 119tags co-culture and sampled between 2-6 days post-inoculation (DPI). At each day, entire taproots were harvested for colony-forming unit (CFU) quantification (Fig. 1b) and tag sequencing to compute Shannon diversity (Fig. 1c). Bacterial loads increased steadily over time, consistent with active growth and progressive host colonisation. In line with our prediction, the mean Shannon diversity converged to ∼ 2 by 4-6 DPI, indicating establishment and stability of pathogen diversity over time once the population was well-established. Notably, *in planta* diversity was consistently lower than that of the inoculum, demonstrating that a 119-tag collection reliably captured a reduction in pathogen diversity at roots level and providing strong evidence for the existence of a root-entry bottleneck.

To assess whether physical root barriers underlie this bottleneck, we performed a second experiment comparing plants with intact roots systems to plants whose roots were mechanically scarified immediately prior to inoculation. Control Marmande tomato plants were infected as described above with the 119-tag co-culture. For each condition, three independent replicates of *n* = 8 plants were harvested at the onset of first disease symptoms within each experimental batch in order to collect samples at a non-saturation phase of the infection (Supplementary Fig. 1). Root samples were processed for CFU and diversity (Figs. 2a,2b). Control intact Marmande tomato plants with intact roots displayed behaviour similar to that observed in the first experiment, both in terms of CFU accumulation and overall infection dynamics. Intact root plants exhibit a pronounced reduction in diversity relative to the inoculum, reflected by losses in both richness (tag extinction) and evenness (frequency imbalance, Fig. 2c). By contrast, plants with cut roots showed significantly higher Shannon diversity at entry (Fig. 2d), indicating enhanced pathogen ingress. Together, these results demonstrate that the physical integrity of roots contributes to the root-entry bottleneck, in agreement with previous qualitative reports showing that root structure impedes early infection ^5,6,7,8^.

**Figure 2.**
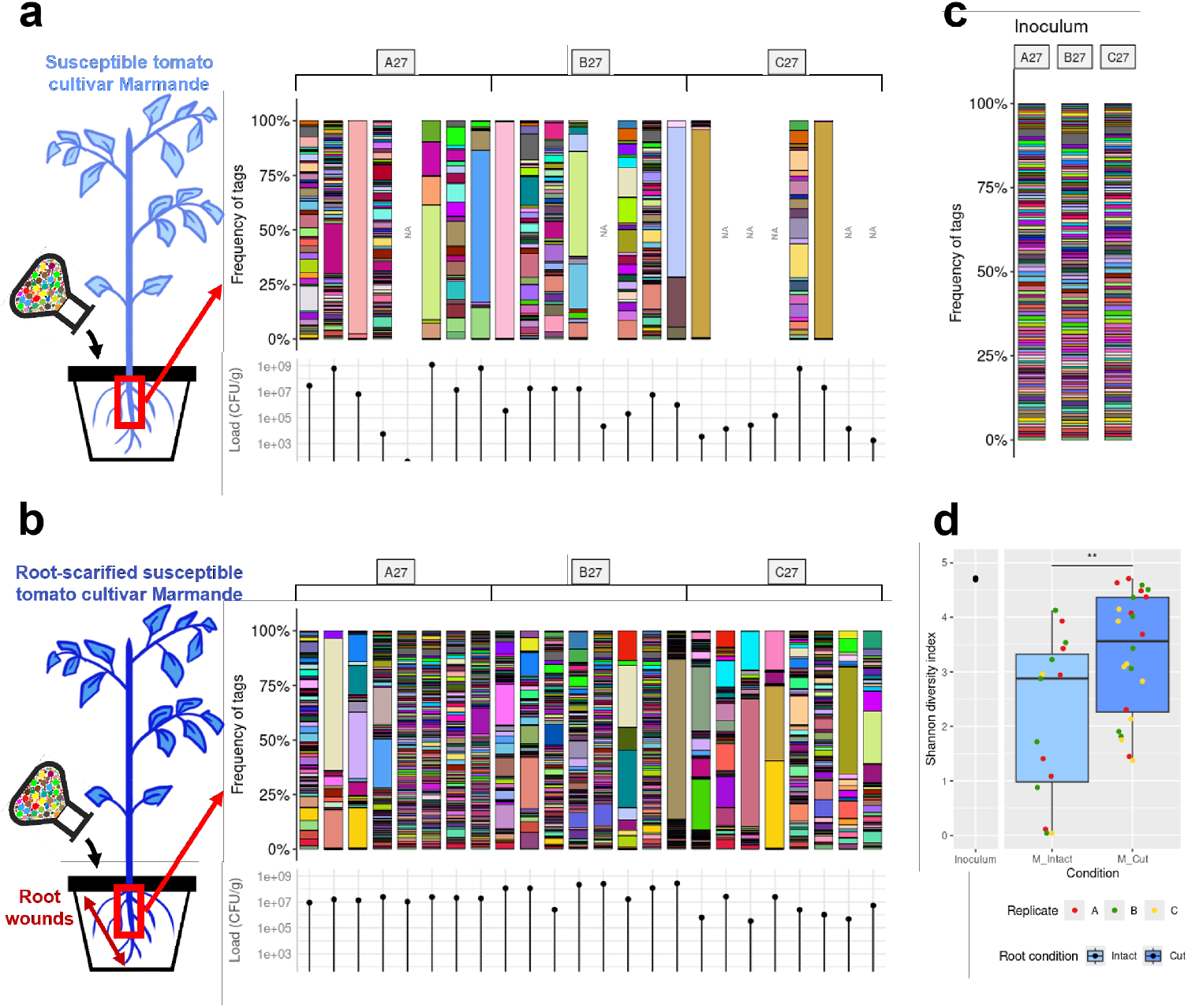
Tomato root-entry bottleneck depends on the physical integrity of the roots. **a and b.** Diversity profiles during colonisation of the roots of control intact Marmande plants (panel **a**, *n* = 24) or root-scarified Marmande plants (panel **b**, *n* = 24). Colours: relative frequency of the 119 tags. Bacterial load is a barplot under each diversity profile (log scale). “NA”: unsuccessful sequencing due to low bacterial load. **c**. Diversity profiles in the inocula (“A27”, “B27”, “C27”) of the three time-independent replicates of infection in panel a and b. **d**. Shannon diversity index in the roots of intact Marmande plants (*M Intact*) or cut Marmande plants (*M Cut*). Wilcoxon-Mann-Whitney test (two sided, **: *p* = 0.0094). Diversity in inocula: 4.70-4.72.

### 1.2 Diversity profiles encode previously unexploited information about the pathogen’s mode of entry

#### Observed diversity profiles are inconsistent with STAMP/STAMPR frameworks

As a first approach, we quantified the root-entry bottleneck using published metrics. We computed the standard bottleneck metric *N*_*b*_ from the original STAMP framework ^1^ and both corrected (*N*_*r*_) and simulated (*N*_*s*_) bottlenecks from STAMPR ^2^. Although derived differently, these metrics share the same interpretation of a bottleneck as the “number of founding pathogen cells”. To evaluate robustness, we compared each estimate to the sample richness *R*, defined here as the minimum number of founders required to generate the observed set of tags *R* in a given sample (Fig. 3a).

**Figure 3.**
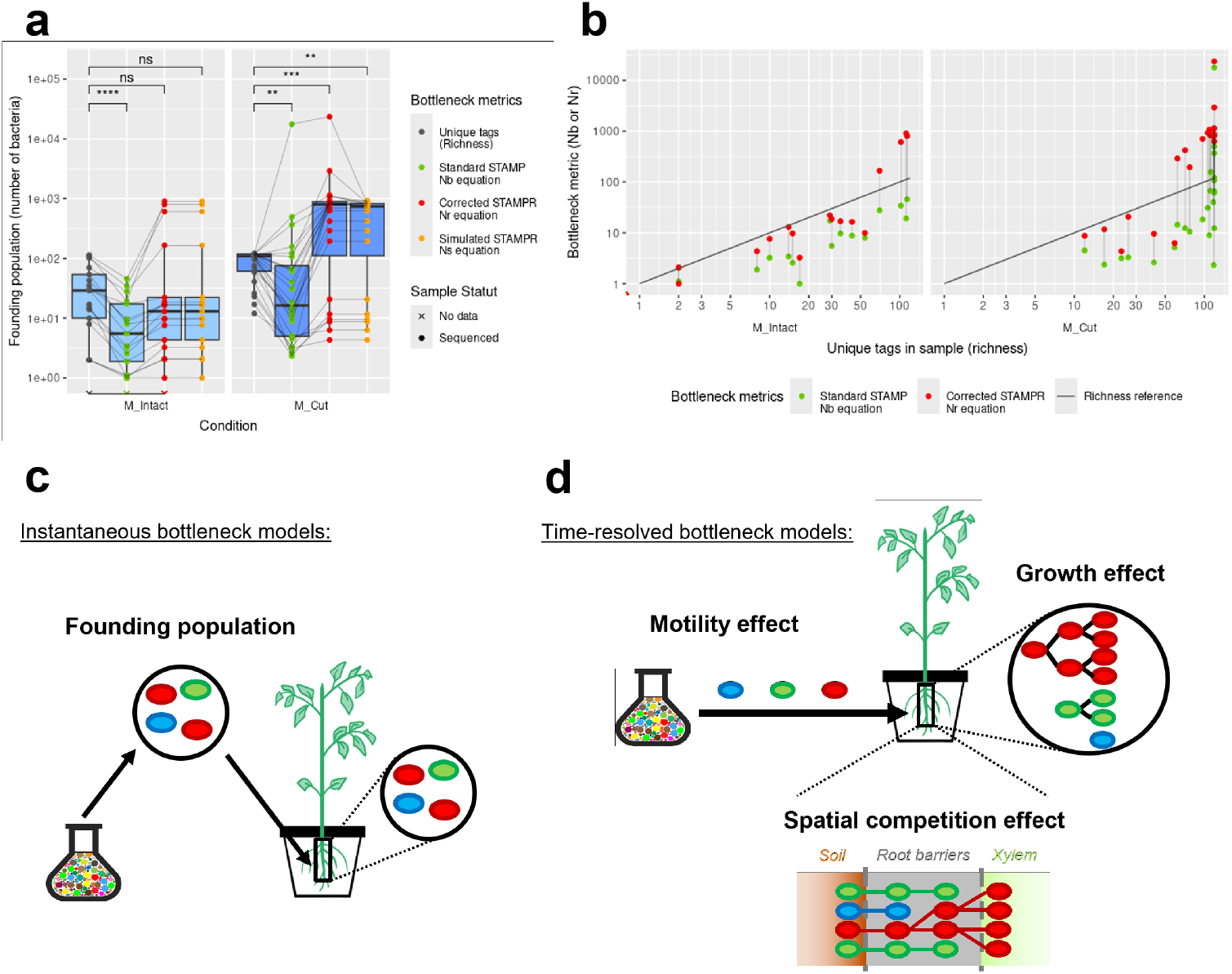
Pathogen diversity profiles better support “time-resolved” rather than “instantaneous” bottleneck models. **a.** Bottleneck estimates in intact (*M Intact*) or root-cut Marmande plants (*M Cut*) using richness, *N*_*b*_ ^1^ or, respectively *N*_*r*_ and *N*_*s*_ ^2^ (log scale). Lines link metrics from the same sample. Wilcoxon signed-rank test (two-sided, ns: *p >* 0.05; *: *p <* 0.05; ** *p <* 0.01; ****: *p <* 0.0001). **b**. *N*_*b*_ and *N*_*r*_ versus of richness (log-log scale). The diagonal *y* = *x* indicates equality; points below it are lower than richness. Line links metrics from the same sample. **c**. Instantaneous bottleneck models assume a single colonisation event from the soil to the roots ^11,12^. The STAMP model (*N*_*b*_) ^1^ or the STAMPR model (*N*_*r*_ and *N*_*s*_) ^2^ follow this assumption (details in Supplementary Fig. 3). **d**. Time-resolved bottleneck models allow entry over time, enabling tests of mechanisms shaping diversity during infection (*e.g*. migration/dispersion to the xylem, early–late growth differences, and spatial competition). Examples include the CR, BS and IR models (details in Fig. *3)*

Median richness was *R* ≈ 30 tags in intact plants and *R* ≈ 110 in cut-root plants. Unexpectedly, all tested bottleneck estimators exhibit some degree of inconsistency with the observed tag counts. The original STAMP metric *N*_*b*_ performed poorly, markedly underestimating entry given the number of detected tags (intact: median *N*_*b*_ ≈ 3; cut: median *N*_*b*_ ≈ 14). in contrast, the STAMPR estimators (*N*_*r*_ and *N*_*s*_) tracked richness more closely overall, although not uniformly across conditions (intact: median *N*_*r*_ ≈ 12; cut: median *N*_*r*_ ≈ 300). Because *N*_*r*_, appeared to be the most consistent estimator, we next tested, sample per sample, whether it could account for the observed richness (Fig. 3b). In most intact-root plants (12/17) and in a substantial subset of cut-root plants (6/24), we found *N*_*r*_ *< R* indicating that systematic discrepancies are commonly. Notably, these failures clustered below a richness threshold of roughly ∼ 60 tags, a regime that was not highlighted previously. Inspection of individual diversity profiles (Figs. 2) further suggest a biological origin: both *N*_*b*_ and *N*_*r*_ over-represent high-frequency tags and effectively discount low-frequency tags. This implies that standard bottleneck metrics capture only part of the colonisation process, and can miss biological meaningful complexity carried by the long tail of rarer lineages.

These discrepancies led us to re-evaluate the biological meaning of *N*_*b*_-derived indicators in our infection system. Revisiting the population-genetic theory underlying these estimators ^11,12^ clarified that the *N*_*b*_ equation ^1^ and the derived *N*_*r*_ and *N*_*s*_ ^2^ are rooted in a very specific regime: barrier crossing is effectively instantaneous and decoupled from subsequent within-host growth. We refer to this class as *instantaneous bottleneck models*, which implicitly assume that a single founder event occurs over a short time window and is independent of time (Fig. 3c). A concrete representative of this class is the STAMP model (ℳ_STAMP_, Supplementary Fig. 3), which is sufficiently explicit to allow mathematical analysis. By contrast, *N*_*r*_ and *N*_*s*_ are computational corrections of *N*_*b*_. Beyond empirical validation, they offer limited analytical tractability, making it difficult to probe their assumptions or failure modes more deeply.

#### Investigating time-dependent effects is necessary for a better understanding of pathogen colonisation

Unlike non-parametric summaries (*e.g*., Shannon diversity, richness), bottleneck estimators such as *N*_*b*_ and *N*_*r*_ are only meaningful in so far as their underlying model assumptions are satisfied. Yet, for *instantaneous bottleneck models*, few practical tools exist to formally assess whether these assumptions are satisfied in a given experimental system. This motivates considering alternative mechanistic descriptions that may better reproduce the observed data. In particular, prior work suggests that *R. pseudosolanacearum* entry proceeds via a sustained influx rather than a brief, short-lived founder event ^5,6,7,8^. Consistent with this view, tag frequencies within samples are not close to the homogeneous compositions expected under instantaneous entry. Instead, diversity profiles typically display pronounced gradients, with a few highly represented tags followed by a long tail of low frequency lineages (Supplementary Fig. 2), suggesting that founding and subsequent growth are intertwined over time.

If entry and growth proceed concurrently, bacteria that enter earlier should, on average, attain higher frequencies than those entering later. This motivates a class of *time-resolved bottleneck models* in which founding overlaps with within-host expansion (Fig. 3d). rather than relying on a single summary metric, our goal is to quantify the contribution of explicit time-dependent biological processes (motility, spatial competition, growth) that cannot be interrogated within the STAMP(-R) paradigm. To challenge the instantaneous-bottleneck assumption, we introduce three alternative models grounded in distinct population genetics perspectives (detailed in Supplementary Fig. 3):

- **Constant-entry-rate model** (ℳ_CR_): pathogen cells enter the root at a constant rate and subsequently grow exponentially.
- **Bolthausen-Sznitman model** (ℳ_BS_): root colonisation is framed as a stochastic range expansion that captures diffusion and spatial occupation within the root. The resulting genealogies follow a Bolthausen-Sznitman coalescent ^13^.
- **Increasing-entry-rate model** (ℳ_IR_): a generalisation of ℳ_CR_ in which entry rates increase exponentially over time; while cells still grow exponentially. This model captures the progressive breaching of the host barriers as observed in ℳ_BS_ (Supplementary Fig. 4), while retaining a simpler, more directly interpretable mechanism.

### 1.3 A model-testing framework reveals new biological insights into *in planta* infection dynamics

We present a model-based procedure for analysing neutral barcoding data in which the shape of the within-host diversity profiles is treated as a diagnostic signature of the infection dynamics. The rationale is straightforward: if diversity profiles encode the mode and timing of entry, then the most plausible bottleneck mechanism can be identified by confronting observed profiles with those predicted by mechanistic models. For each sample, we simulate tag-frequency distributions under four candidate models—ℳ_STAMP_, ℳ_CR_, ℳ_BS_, ℳ_IR_—and estimate model parameters by optimising agreement with the empirical diversity structure (Fig. 4a). Model fit is quantified by minimising the discrepancy between a sample’s Hill-numbers curve and the corresponding simulated Hill-numbers curve.

**Figure 4.**
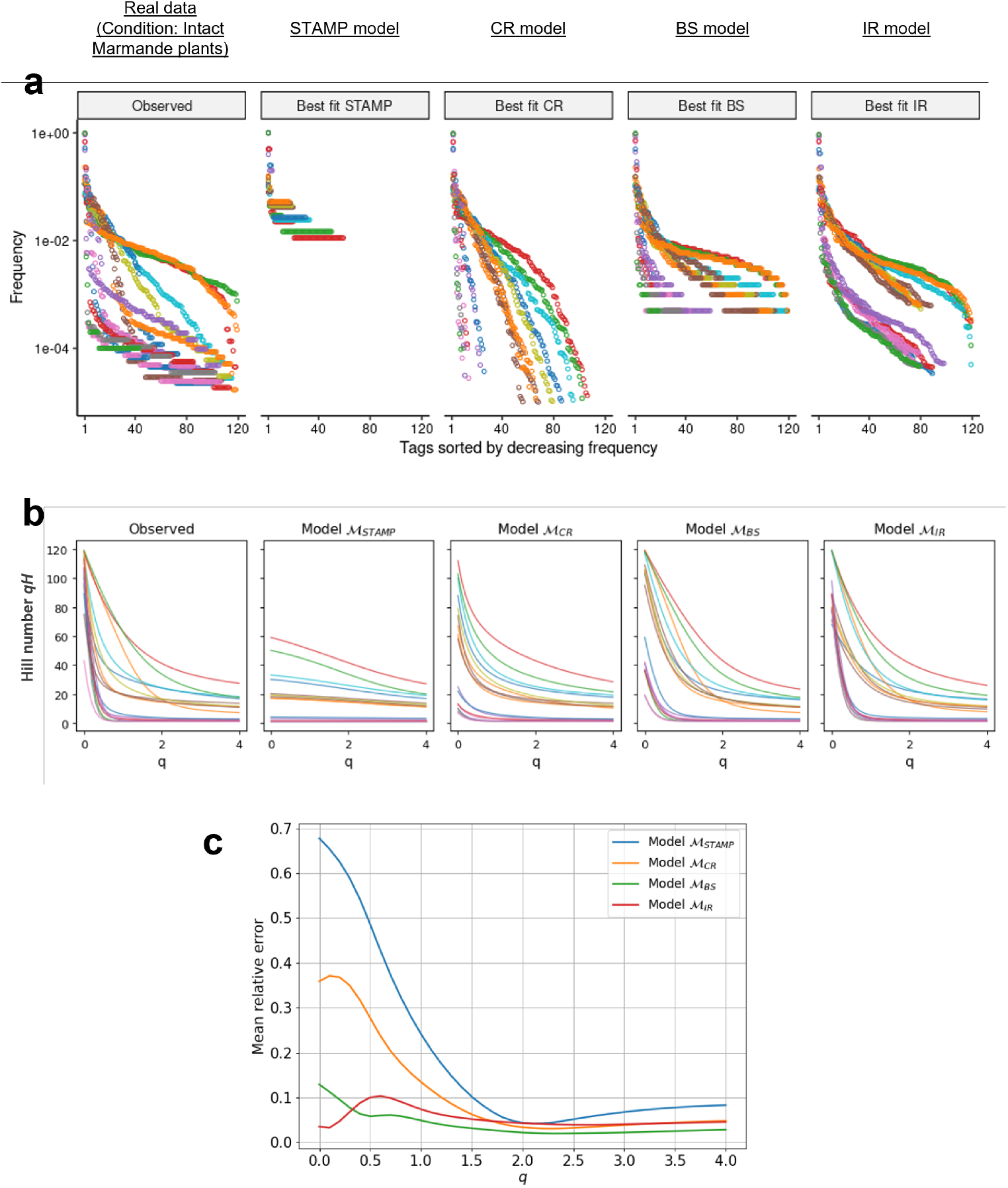
Time-resolved bottleneck models better explain tomato root colonisation. **a.** Diversity profiles observed in samples (intact Marmande plants) compared to the best fitting profiles under simulation of the models ℳ_STAMP_, ℳ_CR_, ℳ_BS_ and ℳ_IR_ (log scale). Colours: individual samples. **b**. Diversity profiles in samples or in simulations according to the Hill numbers ^*q*^*H* from order *q* = 0 (importance of richness) to order *q* = 4 (importance of evenness). As a guide, *q* = 0 equals richness, *q* = 1 is the exponential of Shannon entropy, and *q* = 2 is the inverse Simpson index (or Simpson diversity). For a given order *q*, a high ^*q*^*H* value in sample indicates a high diversity. Colours: individual samples. **c**. Error between simulations and real data in the range from *q* = 0 to *q* = 4, averaged over all infection conditions in Fig. 2 and Fig. 5.

A Hill-numbers curve provides a compact and informative representation of diversity profile structure because it jointly captures tag richness and relative abundance ^14^. Hill-numbers curves can be examined directly from empirical data or from the the best-fitting simulations produced by each model (Fig. 4b). Low orders *q* give greater weight to rare tags, whereas higher orders *q* increasingly emphasise dominant tags. As a result, the full Hill-numbers curve offers an essential one-to-one summary of a tag-frequency distribution, allowing diversity profiles to be compared unambiguously across samples and models.

Models’ accuracy was quantified using an error score defined as the deviation between observed and simulated Hill-numbers curves, computed across all samples (Fig. 4c) or separately by condition (Supplementary Fig. 5). This model-testing framework allows us to address the nature of the bottleneck directly. ℳ_STAMP_ systematically failed to reproduce the contribution of low frequency tags, indicating that an instantaneous bottleneck is not an adequate representation of the infection process. In contrast, ℳ_CR_, ℳ_BS_ and ℳ_IR_ achieved markedly better fits, simultaneously capturing both the near-equilibrium among dominant tags and the persistence of numerous low-frequency lineages. Together, these results support a time-resolved bottleneck as a more plausible explanation of roots infection dynamics.

Because each time-resolved bottleneck model corresponds to a distinct biological scenario, additional qualitative insight can be gained by comparing their relative performance. Notably, ℳ_BS_ and ℳ_IR_ outperformed ℳ_CR_ at low orders (*q* ≤ 2), where the Hill-numbers are more sensitive to low-frequency tags. The key mechanistic distinction is that ℳ_CR_ assumes a constant entry rate, whereas both ℳ_BS_ and ℳ_IR_ allow entry to accelerate over time (Supplementary Fig. 4). This comparison refines our interpretation of the bottleneck in two ways:

- First, low-frequency tags—systematically absent in instantaneous bottlenecks models—are a natural outcome of time-resolved bottlenecks. The superior performance of ℳ_BS_ is consistent with its capacity to preserve rare lineages through the genealogical structure of range expansions ^13^ whereas ℳ_CR_ tends to erode this rare tail. This implies that even under highly stringent entry, minor secondary infection events can occur and remain detectable.
- Second, the improved fit of models with accelerating entry suggests that pathogen influx into the plant increases as infection progresses, consistent with a progressive weakening of host barriers. This interpretation aligns with reports of plasmolysis of cortical cells during infection ^6,7,8^. By compromising tissue integrity and potentially widening access routes toward the vasculature, such damage would be expected to mechanically facilitate continued, and increasing, pathogen entry into the xylem.

### 1.4 Time-resolved bottlenecks quantify plant susceptibility or tolerance

To broaden inference, we extended the infection dataset across three experimental factors: *root condition* (intact *vs*. cut), *plant genotype* (susceptible *cv*. Marmande *vs*. tolerant *cv*. Hawaii7996^8,15,16^), and *temperature* (27 ^°^C *vs*. 32 ^°^C) (Fig. 5a). This factorial design yielded eight conditions, including the two reference baselines of intact or cut Marmande tomato plants infected at 27°C (*M_Intact_27, M_Cut_27*) from Fig. 2. Plants were sampled after symptom onset, as described in Materials and Methods.

**Figure 5.**
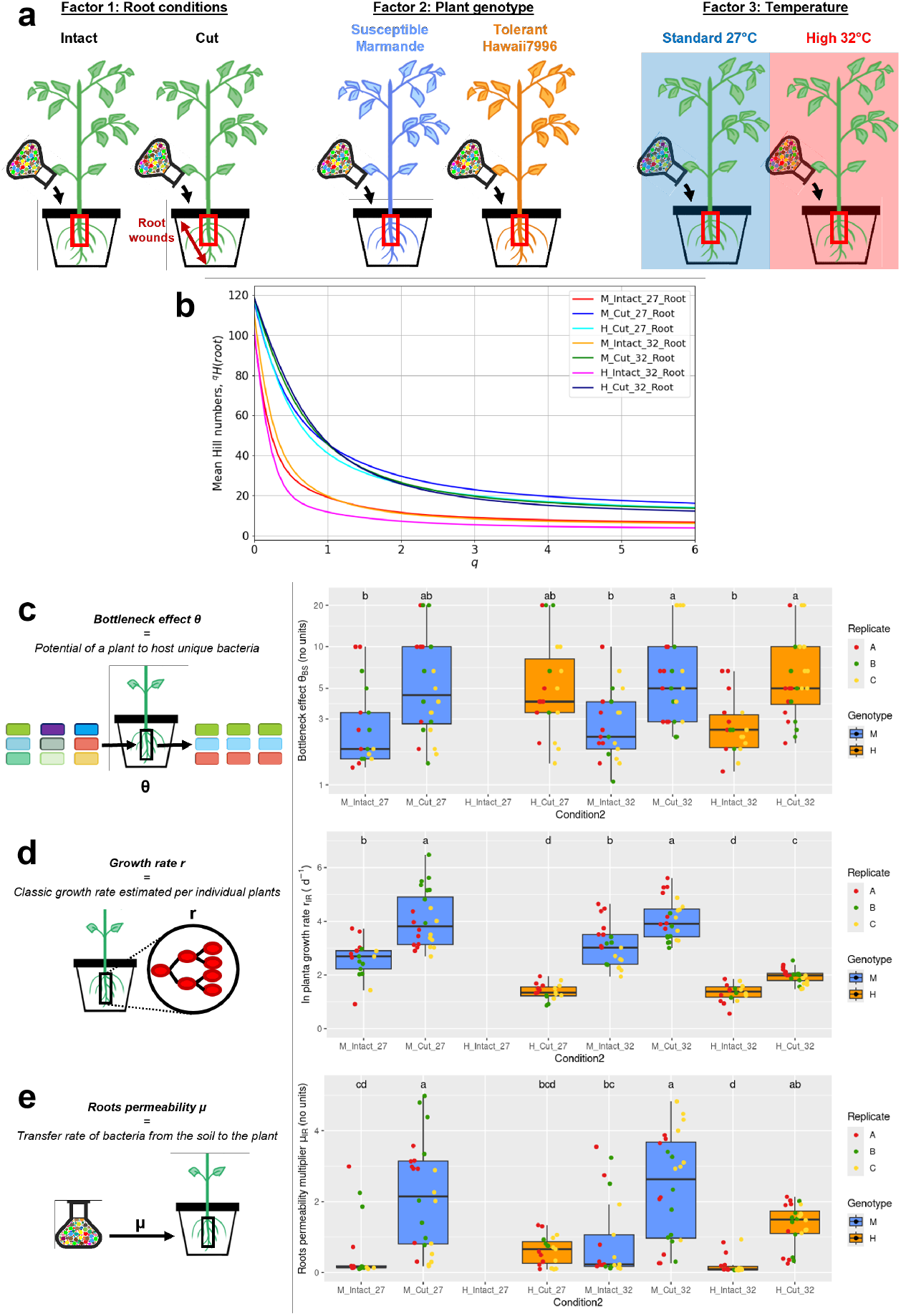
Susceptibility to *R. pseudosolanacearum* reflects time-dependent infectious dynamics. **a.** Eight tested conditions combine three factors of plant susceptibility (*n* = 24 plants/condition). “Root condition” factor: intact plants versus root-cut plants. “Plant genotype” factor: susceptible cultivar Marmande versus tolerant cultivar Hawaii7996. “Temperature” factor: standard (27°C) versus high temperature (32°C). **b**. Hill-numbers curves between *q* = 0-6 for pathogen diversity averaged over each condition (M: Marmande, H: Hawaii7996, I: Intact, C: Cut). Condition *H_Intact_27* was tolerant and did not yield sufficient colonisation for sequencing. **c**. Estimated bottleneck parameter *θ*_BS_ = *v* obtained with *ℳ*_BS_ (log scale). “Bottleneck stringency” *θ*_BS_ reflects monoclonal or diverse colonisation. Low value indicates stringent bottleneck. Kruskal-Wallis test, *post hoc* Fisher’s LSD (threshold: *p <* 0.01). **d**. “Growth rate” *r*_IR_ in individual plant roots estimated from the parameter *θ*_IR_ = *µ*_IR_*/r*_IR_ in *ℳ*_IR_. Kruskal-Wallis test, *post hoc* Fisher’s LSD test (threshold: *p <* 0.01). **e**. “Entry rate multiplier” *µ*_IR_ in each individual plant roots estimated from the parameter *θ*_IR_ = *µ*_IR_*/r*_IR_ in *ℳ*_IR_. *µ*_IR_ reflects the transfer rate of pathogen from soil to the plant roots. High value indicates faster entry of pathogen over time. Kruskal-Wallis test, *post hoc* Fisher’s LSD test (threshold: *p <* 0.01).

All three factors are known to modulate disease progression. Susceptibility-enhancing modalities (Marmande tomato plants, cut roots, 32 ^°^C, noted *M Cut 32*) led to earlier symptom onset, higher bacterial load (CFU), and consequently earlier sampling (Supplementary Fig. 1). Conversely, tolerance associated modalities (Hawaii7996 tomato plants, intact roots, 27 ^°^C, or *H_Intact_27*) limited colonisation, generating a continuum from highly susceptible (*M_Cut_32*) to strongly tolerant plants that remained asymptomatic and therefore could not be analysed (*H_Intact_27*).

#### Bottleneck stringency depends primarily on root integrity

Across conditions, the dominant driver of entry diversity was root state: intact roots admitted low diversity, whereas cut roots admitted high diversity (Fig. 5b). To compare bottleneck stringency across modems, we summarised it with a single parameter *θ* that can be extracted from each model (*θ*_STAMP_ = *N*_*b*_; *θ*_CR_ = *µ*_CR_*/r*_CR_; *θ*_BS_ = *v*; *θ*_IR_ = *µ*_IR_*/r*_IR_; Supplementary Table 1). In time-resolved models, *θ* measures how rapidly new lineages enter relative to the expansion of already established lineages, making a “number of founders” interpretation inappropriate. For the models ℳ_CR_ and ℳ_IR_, the underlying entry rate *µ* and the population growth rate *r* can be recovered from *θ*.

In ℳ_BS_, *θ*_BS_ can be interpreted as a relative speed of barrier transversal: higher values correspond to a more synchronous entry, and therefore higher diversity at the sampling site. Because ℳ_BS_ provided the best overall fit (Figs. 5), we used *θ*_BS_ as our primary comparator across conditions (Fig. 5c). Intact roots exhibit stringent bottlenecks (median *θ*_BS_ ≈ 1.5–2), whereas cut roots admitted substantially greater diversity (median *θ*_BS_ ≈ 4–5). We then used a PERMANOVA to quantify how each experimental factor contributed to variation in *θ*_BS_ (Supplementary Fig. 6). Root condition was the only significant predictor (*R*^2^ = 16.3%) indicating that *θ*_BS_ is largely insensitive to susceptible differences driven by genotype or temperature. Thus, even in a time-resolved framework, a single bottleneck stringency metric *θ* is not sufficient to characterise infection dynamics: it summarises endpoint diversity, but does not capture the total pathogen burden within the sample.

#### Growth and entry rates predict susceptibility

Although ℳ_CR_ fit the data slightly less well than ℳ_BS_, it offers a key advantage: by combining diversity with CFU and sampling time, it enables estimation, at the level of individual plants, of both the pathogen growth rate *r*_IR_ (Fig. 5d) and a root permeability (entry rate) *µ*_IR_ (Fig. 5e). Unlike *θ*_BS_, these two parameters reflect susceptibility across experimental conditions. PERMANOVA confirmed that all three factors (root condition, genotype, temperature) significantly influenced both *r*_IR_ and *µ*_IR_ (Supplementary Fig. 6). Genotype was the dominant driver of growth (*R*^2^ = 64.6%), with susceptible plants exhibiting an approximately twofold higher *r*_IR_. Root scarification and elevated temperature had additional, smaller but significant effects on *r*_IR_ (*R*^2^ = 3.0% and 3.2%, respectively). In contrast, permeability was primarily controlled by root scarification (*R*^2^ = 24.4%), with significant contributions from genotype and temperature as well (*R*^2^ = 5.1% and 2.6%).

We also examined the role of plant size by including root samples weight as a proxy for a *plant size* factor (PERMANOVA; Supplementary Fig. 6). Plant size explained a substantial fraction of the variation of *r*_IR_ (*R*^2^ = 37.3%), but this effect largely reflected confounding with genotype, because tolerant plants were typically larger than susceptible plants in our setup. For *µ*_IR_, only plant size, root condition and genotype remained significant (respectively *R*^2^ = 4.7%, 25.0%, and 1.2%). This pattern indicates that, independent of the other factors tested, larger plants tend to show higher estimated growth and entry rates, *i.e*. increased susceptibility, as captured by both *r*_IR_ and *µ*_IR_ overall indicating that fine susceptibility patterns are well-captured by pathogen influx parameter *µ* and within-host growth parameter *r*.

### 1.5 Time-resolved bottleneck models extend to animal infections

To assess whether our approach generalises beyond plants, we applied the same model-testing framework to a published dataset of *Pseudomonas aeruginosa* bacteraemia in mice (Fig. 2C in reference ^9^), which was previously reanalysed with STAMPR (Fig. 4 in reference ^2^). Following intravenous inoculation with a tagged *P. aeruginosa* library, organ-specific bottlenecks were reported using *N*_*b*_ and *N*_*r*_ (Fig. 6a; ^2,9^). Tag-relatedness analyses further suggested distinct seeding routes: lung, spleen, and liver primarily seeded by blood-circulating bacteria, whereas the gallbladder and gastrointestinal tract derive from mixtures of liver-seeded and blood-circulating populations ^2,9^.

**Figure 6.**
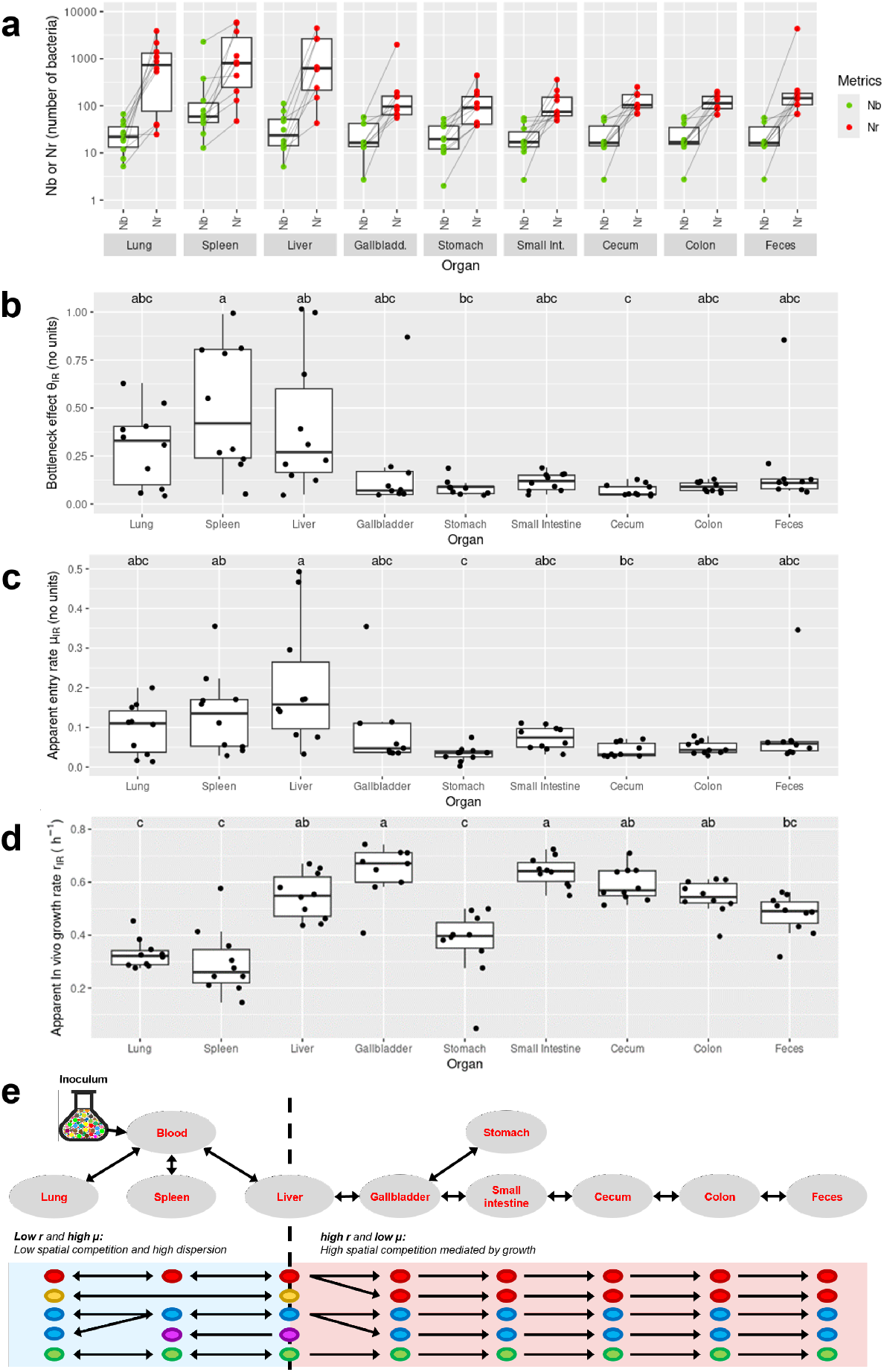
Time-resolved models capture overlooked dynamics in systemic mouse-*Pseudomonas aeruginosa* infection (original data from Fig. 2C of article ^9^, *n* = 10 mice). **a.** *N*_*b*_ and *N*_*r*_ as calculated in ^2,9^ (log scale). Line link metrics of same sample. **b**. Bottleneck parameter *θ*_IR_ obtained with *ℳ*_IR_ (log scale). Low value indicates stringent bottleneck. Kruskal-Wallis test, *post hoc* Fisher’s LSD test (threshold: *p <* 0.01). **c**. Apparent growth rate *r*_IR_ in individual organs. Whether *r*_IR_ corresponds to absolute growth rate *in vivo* depends on the assumption of sampling during exponential growth. Kruskal-Wallis test, *post hoc* Fisher’s LSD test (threshold: *p <* 0.01). **d**. Apparent entry rate multiplier *µ*_IR_ in individual organs. In this context, *µ*_IR_ represents the influx of systemic circulating bacteria into organs. High value indicates faster entry of pathogen over time. The accuracy of *µ*_IR_ is conditioned to the accuracy of *r*_IR_. Kruskal-Wallis test, *post hoc* Fisher’s LSD test (threshold: *p <* 0.01). **e**. Schematic hypothesis regarding mouse-*P. aeruginosa* infection. Lung, spleen and liver are colonised under a high dispersion mode (low *r*, high *µ*) with high connection to inoculation site (blood) promoting establishment of high pathogen diversity. Gastrointestinal organs are colonised under a high spatial competition mode (high *r*, low *µ*) with gallbladder being quickly colonised by the first arriving cells promoting the establishment of a homogeneous, low diversity.

#### Model comparison identifies time-resolved bottlenecks

The original studies already argue against an instantaneous bottleneck: the systematic underestimation of *N*_*b*_ relative to *N*_*r*_ was interpreted as evidence that clonal expansion occurs during organ colonisation ^2^. We agree that systemic infection reflects two concurrent processes: organ seeding by circulating bacteria (an entry rate *µ*) and within-organ proliferation (a growth rate *r*). Our framework makes these process explicit and directly tests the bottleneck assumption that remains implicit under STAMP(-R) metrics.

Consistent with this interpretation, error-curve comparisons show that the instantaneous model (ℳ_STAMP_) provides a poor description of organ diversity profiles, whereas time-resolved models yield substantially improved fits (Supplementary Fig. 7). Using the best-performing model (ℳ_IR_), we then extract *θ*_IR_, an interpretable measure of bottleneck stringency (Fig. 6b). As in our plant system, *θ*_IR_ captures whether colonisation proceeds in a largely monoclonal versus diversified manner under continuous pathogen fluxes.

#### Entry and growth rates resolve organ-specific dynamics

Because our approach integrates diversity, CFU and sampling time, a key advance is that it allows estimation of pathogen entry *µ*_IR_ and growth *r*_IR_ at the resolution of an individual organ in an individual mouse. Estimated entry rates *µ*_IR_ (Fig. 6c) indicate that lung, spleen, and liver are relatively permeable to circulating bacteria, whereas the gallbladder and gastrointestinal tract are comparatively impermeable. These results agree with relatedness-based inferences that the gastrointestinal tract is not directly seeded from blood ^2,9^, while adding a quantitative flow-rate interpretation through *µ*_IR_. As a additional, and previously unreported insight, the inferred growth rates *r*_IR_ (Fig. 6d) suggest that lung and spleen share similar dynamics (median *r*_IR_ ≈ 0.3 h^−1^), whereas liver, gallbladder, and most of the gastrointestinal tract (with the notable exception of the stomach) display roughly twofold higher apparent growth.

#### Revisiting the “liver filter” interpretation in the *P. aeruginosa* infection

A previous interpretation proposed that the liver acts as a filter for bacteria prior to entry into the gallbladder, based on the observed reduction in *N*_*b*_ or *N*_*r*_ from liver to gallbladder ^2,9^. Our time-resolved analysis refines this view. We likewise infer a decrease in permeability between liver and gallbladder, but additionally detect a marked increase in downstream growth. Together, these results suggest an alternative mechanism: elevated growth in the gallbladder can amplify competitive exclusion, reducing effective diversity and producing an “apparent bottleneck” signature even under continued, time-resolved entry (Fig. 6e).

#### Model-based metrics outperform STAMPR in biological meaning and statistical power

We next reanalysed a second dataset of systemic infection, tracking dissemination of *Klebsiella pneumoniae* following a pneumonia in mice (Fig. 1 in reference ^10^). As for *P. aeruginosa*, time-resolved models provided a substantial better fit to organ diversity profiles than the instantaneous model ℳ_STAMP_ (Supplementary Fig. 8). To further illustrate the added value of our framework, we compared our model-based parameters to the full set of STAMPR-derived metrics previously used to describe infection dynamics, and quantified the extent to which model-based inference improves interpretability and discriminative power (Supplementary Fig. 8).

In brief, the interpretable bottleneck metric *θ*_IR_ can replace *N*_*s*_, avoiding the conceptual ambiguity that arises when a time-resolved process is summarised with instantaneous-bottleneck indicators. The inferred growth rate *r*_IR_ also provides a mechanistic counterpart to empirical proxies of expansion that are difficult to interpret, such as *CFU/N*_*s*_ and *N*_*s*_*/N*_*b*_. Likewise, the inferred entry rate *µ*_IR_ yields a direct estimate of pathogen transfer, an essential quantity that is not explicitly addressed in STAMPR. In terms of statistical performance, these model-based parameters generally required no log-transformation for parametric testing and more clearly separate experimental effects, suggesting both improved robustness and increased biological interpretability for analysing pathogen infection dynamics.

#### *K. pneumoniae* mode of dissemination is observable through within-host growth

Two modes of *K. pneumoniae* dissemination from the lung to systemic sites have been described ^10^: *metastatic dissemination*, in which systemic infection originates from a small number of dominant lineages that first expanded in the lung, and *direct dissemination*, in which a diverse, multiclonal pathogen population spreads to peripheral organs.

To strengthen this conclusion, we analysed *r*_IR_ across multiple organs, an inference that cannot be made from CFU alone. In *direct disseminations* cases at 24h post-infection, apparent growth rates were strikingly consistent across organs, with typical values around *r*_IR_ ≈ 0.45h^−1^ (Supplementary Fig. 8). By contrast, *metastatic disseminations* was associated with an approximately 40% increase in *r*_IR_ across essentially all organs. This suggests that *metastatic dissemination* is not driven solely by enhanced replication in the lung, as originally proposed, but instead reflects a more general, host-wide increase in pathogen replication.

## 2 Discussion

Here, we propose to rethink the analysis of neutral barcoding experiments by treating diversity-profile structure as the result of a process combining successive infections and simultaneous growth of the pathogen population, rather than resulting from an instantaneous bottleneck event. Founder metrics (*N*_*b*_, *N*_*r*_, *N*_*s*_) are appealingly simple, but they assume that entry and growth are decoupled. Accordingly, elevated *N*_*s*_*/N*_*b*_ or *N*_*r*_*/N*_*b*_ from STAMPR have been interpreted as signatures of clonal expansion during colonisation ^2,10,17,18^. This interpretation is useful, yet it also highlights the limits of founder-based metrics when entry and growth co-occur.

Our results expose a counter-intuitive property of bottleneck metrics *θ* (*i.e*., *N*_*b*_, *N*_*r*_, *N*_*s*_, *θ*_CR_, *θ*_BS_, *θ*_IR_): Under time-resolved dynamics they do not quantify host defence or susceptibility. Instead, *θ* summarises the diversity among established lineages, without indicating how many bacteria (or how much flux) produced that profile. Capturing host barriers requires integrating pathogen burden and timing, via the entry and growth parameters *µ* and *r*. In our data for instance, *M Intact 32* showed rapid entry and growth with early symptoms, whereas *H Intact 32* showed slower entry and growth with delayed symptoms. Yet both display similar *θ*_BS_ values. They differ in susceptibility because (*µ*_IR_, *r*_IR_) differ. This confusion traces back to the instantaneous STAMP assumption, where *N*_*b*_ conflates diversity profile structure (a *θ*-like quantity) with entry (*µ*). In time-resolved systems, *µ* and *r* cannot be recovered from *N*_*b*_-derived indicators or *θ*_BS_. Fully characterising host defences therefore requires explicitly estimating (*µ, r*), for example, using ℳ_CR_ or ℳ_IR_.

The proposed parameters (*µ, r*) substantially extend what can be learned from neutral barcoding experiments, enabling growth-rate inference at the level of individual hosts and even at individual organs. Critically, they require only measurements that are typically already collected: diversity profile, CFU and sampling time. Because *r* reflects an exponential growth process, absolute estimations of (*µ, r*) require CFU measurements taken during the exponential phase. In that regime, inferred *r* (Fig. 5e) is consistent with standard linear regression of CFU over time (Fig. 1b), while additionally providing per-plant resolution. When CFU are measured at saturation, (*µ, r*) can still be estimated, although they become “apparent” parameters that are reliable for relative comparisons but should not be interpreted as absolute estimations.

In *R. pseudosolanacearum* infections, we leveraged contrast in host and environment to dissect susceptibility. Relative to the susceptible cultivar Marmande, the tolerant cultivar Hawaii7996 showed reduced within-plant pathogen growth, consistent with prior work ^5^, whereas differences in root permeability were less clear. Root scarification increased permeability and, unexpectedly, also increased within-plant growth, suggesting that wounding decreases internal root defences.

A plausible mechanism is that damage to root-barriers, known to regulate metabolite containment and the root-associated microbiome ^19,20,21^, promotes leakage of exudates such as glutamine. The latter is both a chemoattractant for soil bacteria ^22^ and a preferred substrate for *R. pseudosolanacearum* during infection ^23^. Likewise, cortical cell death reported during infection ^6,7,8^ could progressively widen access routes toward the xylem, enhancing both physical entry and exudate leakage over time. Elevated temperature further increased susceptibility by boosting growth and permeability. Interestingly, *r*_IR_ and *µ*_IR_ are correlated across plants (Spearman rank correlation *R*^2^ = 0.40, *p <* 0.05; Supplementary Fig. 6), suggesting a general partial coupling of the two effects during infection processes.

In *P. aeruginosa* bacteraemia in mice, our reanalysis reveals a marked within-host shift in behaviour: dissemination is strongest in blood-connected organs (lung, spleen and liver) whereas apparent growth is highest in the gastrointestinal tract (gallbladder, stomach, small intestine, caecum and colon). This pattern suggests selective pressure and competition to rapidly establish in the gastrointestinal tract, potentially because it enhances persistence and transmission through faeces shedding, or because it provides a niche for chronic infection. In line with this view, recent work links high replication potential of *P. aeruginosa* to stringent gastrointestinal tract bottlenecks ^24^, consistent with our growth-driven interpretation of “apparent bottlenecks”.

More broadly, how common are time-resolved versus instantaneous bottlenecks? The field increasingly acknowledges that time matters in neutral barcoding analysis ^2,25^. Direct evidence for truly instantaneous founding remains limited, while many frequently observed signatures, strongly heterogeneous tag frequencies, spatial and temporal declines in *N*_*b*_, abundant rare-frequency tags, or *N*_*r*_*/N*_*b*_ *>* 1, naturally arise when entry and growth overlap. This raises the possibility that instantaneous-bottleneck analysis have often been applied to systems dominated by time-dependent processes.

A notable step towards time-resolved modelling is RESTAMP for *Vibrio cholerae* in the mouse gastrointestinal tract ^3,25^, which emphasises transfer rates and growth across compartments. However, its high complexity (∼ 30 fitted parameters ^25^) and reliance on *N*_*b*_ to infer secondary parameters may hinder interpretability and increase the risk of over-fitting in a model that did not provide control of the fit against full tag frequency data.

Here, we take a different route: we compare a panel of mechanistic models directly against the observed diversity structures and select those best supported by the data. This strategy has three advantages: (*i*) assumptions need not be accepted *a priori*, poor models are rejected by fit, (*ii*) whether the bottleneck is instantaneous or time-resolved emerges from model performance, (*iii*) the selected models yield interpretable parameters that map onto explicit biological processes ^26^. More generally, Hill-numbers curves matching provides a standardised way to test any hypothesised scenario, provided it can be formalised into a mechanistic model. Our fits are not perfect, indicating that additional mechanisms could be investigated. For instance, tissue-specific heterogeneity could affect arrival times, local densities, or growth conditions but would require spatially resolved sampling and explicit spatial models.

We show that three widely applicable time-dependant mechanisms (growth, motility and spatial competition) are intertwined in bacterial infection dynamics. Because of the generality of the addressed mechanisms, the improvements in robustness and in biological insights without modifications in current experimental protocols, our findings support a need to shift from the previous *N*_*b*_-based methodology to an updated methodology accounting for time-dependant factors. We advocate for a global reanalysis of STAMP(-R) articles under a time-resolved model approach. We foresee that underappreciated biological results may be uncovered in the following cases: (*i*) studies that used only *N*_*b*_ and may have overlooked major aspects of the infection dynamics ^1,9,27,28,29^, (*ii*) studies of colonisation across continuous space (*e.g*. gastrointestinal tract colonisation) that neglected contributions of motility and growth effects by using *N*_*b*_-derived indicators ^2,10,18,24,25,30,31,32^, (*iii*) studies that empirically uncovered the importance of time dimensions using *N*_*b*_-derived indicators but were limited in their analysis ^17,18,30^, (*iv*) studies that directly addressed relations between CFU dose effects and bottleneck using *N*_*b*_-derived indicators but did not pair the two in an appropriate model ^30,33,34^.

Beyond refining earlier findings, the proposed methodology more broadly aims to connect microbial population dynamics to quantitative biological mechanisms. It opens the possibility to address questions in microbial population genetics with a unified mechanistic framework that explicitly includes time-dependant complexity. Natural extensions of this work include bringing time-resolved inference to broader microbial ecology contexts such as: tracking of complex bacterial synthetic communities ^35,36^, genealogies tracing of a bacterial population and inference of evolution rates ^13^ or mapping of complex directional pathogen fluxes at a host-wide level ^37^.

## 4 Methods

### 4.1 Construction of the STAMP collection in *R. pseudosolanacearum* and co-culture of the strains

123 oligonucleotides of a 20 bp random sequence (referred as “tag”, list in supplementary document “STAMP123 tags.fasta”) were synthesised and hybridised with their complementary sequence to form inserts with cohesive extremities BsrGI and BglII. These fragments were inserted by restriction–ligation (BsrGI / BglII) in a pRCT-GWY ^38^ derived plasmid denoted pNP413. The tag is contained inside a genetic cassette of 6 kb containing a *tetR* gene, a *mCherry* gene and two flanking regions of 1 kb used for homologous recombination with *R. pseudosolanacearum* strain GMI1000 in the chromosome ^38^.

Produced plasmids were chemically transformed into calcium-competent *E. coli* strain DH5alpha and selected using LB medium + tetracycline 10 mg/L. Clones were verified using a wild-type discriminating PCR and Sanger sequencing of the STAMP tag (Eurofins). Validated plasmids were purified in 50 *µ*L ultrapure water using 2 mL of stationary-phase culture with a column extraction kit (BS614, Bio Basic). 10 *µ*L of plasmid was linearised using SfiI enzyme and mixed with 200 *µ*L of *R. pseudo-*

*solanacearum* strain GMI1000 from a liquid culture in exponential phase, grown 1 week at 28 ^°^C in MP medium + 2% glycerol. The mix was deposited on MP medium + 2% glycerol Petri dishes and incubated 2 days at 28 ^°^C for natural transformation. Three days later, the bacterial lawn was suspended in distilled water and isolated on Phi medium + tetracycline 10 mg/L Petri dishes. The obtained *R. pseudosolanacearum* strains were validated for the correct insertion of the 6 kb cassette using a wild-type discriminating PCR and Sanger sequencing of the STAMP tag (Eurofins).

Individual clones were grown in 96-well microplates at 28 ^°^C in 250 *µ*L of Phi medium + tetracycline 10 mg/L in an Omega FLUOstar spectrophotometer (BMG Labtech) until stationary phase. An *OD*_600*nm*_-adjusted volume (50 AU_OD_ * *µ*L per well) was used to inoculate a secondary microplate culture in the same conditions. At stationary phase, wells were pooled 8-by-8 using *OD*_600*nm*_-adjusted volumes (100 AU_OD_ * *µ*L per well) for inoculation of tertiary Erlenmeyer cultures at 28 ^°^C in 20 mL Phi medium + tetracycline 10 mg/L overnight. At stationary phase, the *OD*_600*nm*_-adjusted volumes of each 8-tag pool (1400 AU_OD_ * *mL*) were mixed with 500 *µ*L glycerol 60% for storage at −70 ^°^C. The day before infection, an inoculum of 120 tags was produced by thawing the 15 tubes of 8 tags (total volume ∼ 25 mL) and growing them directly in an Erlenmeyer with 250 mL of Phi medium + tetracycline 10 mg/L at 28 ^°^C overnight. Upon validation of these final co-cultures, one tag (tag007) was found missing but an otherwise almost perfect stoichiometry was obtained for the 119 other tags. The three remaining tags (denoted tag121, tag122 and tag123) were grown and kept separately for sequencing calibrations.

Used media. LB medium: Yeast extract (Duchefa Biochimie) 5 g/L; Bacto tryptone (Becton Dickinson Difco) 10 g/L; NaCl 10 g/L in distilled water. Phi medium: Bacto peptone (Gibco) 10 g/L; Casamino acid (MP) 1 g/L; Yeast extract (Duchefa Biochimie) 1 g/L in distilled water. MP medium: KH2PO4 3.4 g/L; (NH4)2, SO4 0.5 g/L; MgSO4 (7H2O) 50 mg/L; FeSO4 (7H2O) 125 µg/L in distilled water. Adjust pH to 7.0 with KOH 10N.

### 4.2 Plant culture, plant infection and samples collection

For all experiments, tomato plants (susceptible cultivar Marmande “M” or tolerant cultivar Hawaii7996 “H”) were sowed in peat-based substrate and then cultivated in 6 × 6 cm pots until an age of approximately 3 weeks in a phytotron (25.5±1 ^°^C day 12 h / 24±1 ^°^C night 12 h, hygrometry 60±15%). The plants were then acclimated to the final culture chamber for 2 or 3 days either at a standard temperature (27±0.5 ^°^C day 12 h / 26±0.5 ^°^C night 12 h, hygrometry 60±5%) or at a high temperature (32±1 ^°^C day and night 12 h/12 h, hygrometry 70±5%). The days of infection, two technical replicates of 2 mL of the inoculum STAMP co-culture were taken for sequencing of the initial population. The co-culture was diluted using tap water (to an *OD*_600*nm*_ = 0.05) and 50 mL of diluted co-culture (∼ 2.5 × 10^9^ CFU) was used to infect plants by watering. Right before inoculation, the condition of “Root cut” plants was produced by scarifying roots of the plants using a scalpel by drawing an “L”-shaped trench in the pot. “Intact” plants were otherwise watered without scarification.

For the experiments of kinetics of infection, one infection was performed (inoculum S27) on *n* = 42 plants (Conditions M_Intact_27). *n* = 8 or *n* = 9 plants were then sampled between DPI 2 to DPI 6. The plant was unrooted and dirt removed. A root sample was collected corresponding to the whole taproot cut at the highest lateral root and removed from all lateral roots. Samples were weighed, surface-sterilised using 70% ethanol and cleaned using sterile distilled water. Samples were cut into the smallest possible pieces using a scalpel and macerated in 2.5 mL sterile distilled water for approximately 3 h. The macerate was used to determine bacterial load by plating serial dilutions on Phi medium + tetracycline 10 mg/L Petri dishes using an automatic spreader (EasySpiral Pro, Interscience). The remaining ∼ 2 mL of macerate was centrifuged and gDNA was extracted from the bacterial pellets using a lysis–isopropanol–ethanol protocol (Wizard Genomic DNA Purification Kit A1125, Promega) in 50 *µ*L final of ultrapure water.

For the experiments with single-point sampling, six temporally independent infections were performed (inocula denoted A27, B27, C27, A32, B32, C32) combining a total of 192 plants (Conditions: M_Intact_27, M_Cut_27, H_Intact_27, H_Cut_27, M_Intact_32, M_Cut_32, H_Intact_32, H_Cut_32). For each condition three replicates were performed *n* = 3 × 8 = 24. At the first symptoms of wilting (or at DPI 13 or 14 if no symptoms) within a batch of 8 plants in a replicate, the whole batch was collected. Bacterial load was quantified and DNA was extracted for sequencing as described above.

### 4.3 Library preparation and sequencing of the diversity profiles

For each purified gDNA, absorbance was measured for an *A*_260*/*280_ quality ratio. Whenever possible, samples were diluted to an empiric concentration between [*C*] = 20 to 100 ng*/µ*L (corresponding to a theoretical number of unique molecules of gDNA of 3 × 10^6^ to 1.5 × 10^7^ tags*/µ*L) satisfying a non-limiting amount of targets and an accurate measure of concentration. For samples in which concentration was below 20 ng*/µ*L or gave aberrant values in view of bacterial load (*e.g*. vegetal phenolic compounds), a value *N*_*max*_ defined as the *theoretical maximum number of tags* was derived from bacterial load. A 35-cycle adapter PCR1 was performed to amplify a 174 bp region containing the tag. This PCR was performed using primers with the advised 22 bp Nanopore adapter regions for PCR barcoding (SQK-LSK114, Oxford Nanopore Technologies). Additional 4 bp regions were included between primer hybridisation sequence and adapter sequence. These 4 bp represent a “run barcode” that allows filtering of cross-contamination when using the same Nanopore flow cell in several different sequencing runs. For this article, 6 different pairs of primers were sequentially used with different “run barcodes” in this fashion. For samples for which [*C*] *>* 20 ng*/µ*L and *N*_*max*_ *>* 2500 tags*/µ*L, 4 *µ*L of purified DNA was used in a 75 *µ*L PCR1. For samples for which [*C*] *<* 20 ng*/µ*L and *N*_*max*_ *<* 2500 tags*/µ*L or samples that did not amplify in the first condition, a larger volume of 20 *µ*L of purified DNA was used in a 75 *µ*L PCR1 to ensure an adequate number of targets. Samples that did not validate the two first conditions were repurified in the same volume using 2X magnetic bead-based mix in a SPRI protocol ^39^ before reattempting a PCR1. Samples that were validated for a ∼ 226 bp band on gel electrophoresis were then purified using 1.5X magnetic bead-based mix in a SPRI protocol ^39^ to eliminate primers and to concentrate the PCR product twofold.

2.5 *µ*L of reconcentrated PCR1 product was used in a 30-cycle barcoding PCR2 in a final volume of 25 *µ*L using primers purchased from the Nanopore 96 Barcode Expansion Pack (EXP-PBC096, Oxford Nanopore Technologies). Samples that were validated for a ∼ 274 bp band on gel were then quantified for DNA concentration using Quant-iT PicoGreen dsDNA Assay (P7589, Invitrogen) and pooled in equal stoichiometry before a purification with 1.2X magnetic bead-based mix in a SPRI protocol ^39^. End-prep and adapter ligation steps were performed according to the PCR barcoding protocol (SQKLSK114, Oxford Nanopore Technologies) with the exception of SPRI purifications that were adjusted to a 1.5X ratio in accordance with the length of the DNA fragments of interest.

A PromethION flow cell R10.4.1 was loaded with an adjusted quantity of 200 fmol of DNA and data acquisition was performed with a goal of minimum depth of sequencing of ∼ 20k unprocessed reads for all samples. The flow cell was washed according to protocol (EXP-WSH004-XL, Oxford Nanopore Technologies) to reuse it for a total of 6 runs of ∼ 96 samples. For technical simplicity, some samples went through the whole sequencing process despite not being validated in the early PCR steps. These samples were excluded from the final analysis.

### 4.4 Raw data processing

DNA sequences were basecalled using Dorado version 0.7.3 with the Super Accurate model v5.0.0 and a minimum Q score of 10. Briefly, sequences were first demultiplexed for the Nanopore barcode and the Nanopore barcodes were trimmed using the in-built Dorado tools. DNA sequences were then processed through a QIIME 2 bioinformatics pipeline for metagenomics. DNA sequences were size-filtered to keep only fragments between 160 and 200 bp. A first closed-reference mapping was performed to identify the “run barcode”. For reads which validated the correct run barcode, a second closed-reference mapping was performed against the 123 potential STAMP tags to calculate the tag abundances. Three tags exogenous tags (namely tag121, tag122, tag123) were used to calibrate barcode hopping. Barcode hopping was estimated by measuring the hopping between the 120 STAMP tags from samples toward the three control tags. A typical order of magnitude of ∼ 1% of noise reads was obtained for final depths of sequencing of *>* 10^4^ reads in all samples.

### 4.5 Validation of the 119-tags collection for STAMP experiments

All strains showed comparable growth *in vitro* and the proportions of the 119 tags remained stable in co-culture aside from tag007 lost due to a pipetting error (Supplementary Fig. 9), indicating that preparation steps did not introduce technical bottlenecks, that tagged strains exhibited indistinguishable *in vitro* growth, and that sequencing did not introduce appreciable PCR bias. To validate neutral fitness of the collection *in planta*, we assumed that all tags should have identical probability to appear across independent infections. Tags showed no consistent over-representation across infections (Supplementary Fig. 9), validating that tagged strains have similar probabilities of establishing infection and the observed diversity profiles reflect pathogen entry dynamics rather than tag-specific fitness effects or technical artifacts such as sequencing bias.

### 4.6 *N*_*b*_-derived indicators calculation and improvements of the STAMPR pipeline

For the calculation of the *N*_*b*_, *N*_*r*_ and *N*_*s*_ metrics, the code from the original STAMPR pipeline ^2^ was adapted. In short, the code was corrected to include the “resiliency algorithm” method and the calculation of *N*_*r*_ in addition to *N*_*s*_ (see below). We also included additional error estimators during the calculation of *N*_*b*_ and *N*_*r*_.

As part of the pipeline, we set the prefiltering threshold to remove noise tags at 25 or fewer reads. This threshold was selected based on the estimation of barcode hopping in our experimental setup. This threshold corresponds to a threshold to filter out at least quantile 99% of tags emerging from noise alone.). Eliminated tags typically accounted for less than 1% of tag abundance in samples and less stringent thresholds had minimal change on *N*_*b*_ and *N*_*r*_.

The standard bottleneck *N*_*b*_ in Fig. 3a, 6a was calculated based on the analytical equation formalised in the original STAMP publication ^1^ using the prefiltered data:

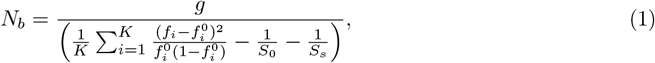

*K* is the number of tags in the inoculum, 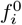 and *f*_*i*_ are the frequencies of tag *i* respectively in the inoculum and the sample, *g* = 1 is the number of generations, *S*_0_ and *S*_*s*_ are the sequencing depths respectively in the inoculum and the sample. A second way to compute *N*_*b*_ is proposed with *θ*_STAMP_ based on the fit of a multinomial model (see Section 4.8)). This second way to compute standard bottleneck can be approximated by a simpler analytical equation:

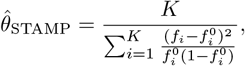

Significant discrepancies between *N*_*b*_ and *θ*_STAMP_ may appear when frequencies *f*_*i*_ do not follow a multinomial distribution.

The limit of detection of the *N*_*b*_ metric was calculated in our experiment by considering a signal-to-noise problem. Noise was considered to be the sum of technical artefacts in tag abundances originating from PCR noise, residual barcode hopping or misattribution of tags during bioinformatic analysis. An artefactual *N*_*b*_ was calculated between technical replicate of sequencing of a same inoculum. *N*_*b*_ *>* 4 × 10^3^ was obtained indicating that it is not possible to discriminate values of *N*_*b*_ above this threshold.

A corrected bottleneck value *N*_*r*_ was derived from *N*_*b*_. The original code of the STAMPR pipeline ^2^ was cleaned and annotated. Previously, only calculation of *N*_*s*_ was made available based on the “reference resample plot” of the algorithm. Instead, we corrected the code so that both *N*_*s*_ and *N*_*r*_ are systematically computed. Finally, an algorithm was added to estimate whether the number of tags in the inoculum is sufficient to accurately estimate a *N*_*b*_ or *N*_*r*_. This algorithm is similar to the simulations that were performed in the original STAMP publication ^1^. The algorithm simulates bottleneck events under the estimated *N*_*b*_ or *N*_*r*_ and the resulting standard deviation provides the confidence interval in the calculation of a given *N*_*b*_ or *N*_*r*_.

### 4.7 Measuring diversity in samples and model comparison

In this section, we construct a method to compare the observed data with the data generated by mechanistic models (see Section 4.8). This method allows us (*i*) to estimate the parameters of our models and (*ii*) to compare the ability of our models to reproduce the observed data.

#### Hill numbers and Hill curve

Our comparison methods is based on the Hill numbers. The Hill number of order *q* of a frequency vector *F* ^1^ = (*f*_1_, …, *f*_*K*_) is defined as:

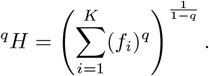

Certain values of *q* correspond to standard diversity indices: ^0^*H* represents species richness (number of types); ^1^*H* corresponds to the exponential of Shannon diversity index; ^2^*H* is related to the inverse Simpson diversity index, giving more weight to common tags. For all values of *q*, the maximum value of ^*q*^*H* is reached when *f*_*i*_ = 1*/K* for all *i*. In this case, ^*q*^*H* = *K*. More generally, for each fixed *q*, a lower ^*q*^*H* corresponds to lower diversity, and the parameter *q* controls the relative emphasis placed on common tags (large *q*) or rare tags (small *q*).

As proposed in reference ^14^, we describe the distribution of the frequencies through the full *diversity profile* ^*q*^*H* over a continuum of values of *q*. The reason for this choice is that the diversity profile fully characterises the frequencies (proposition A22 of reference ^14^), as two frequency vectors cannot lead to the same profile unless they are equal up to a permutation.

#### Models and data comparison

Observing the Hill curves allows us to directly visualise the effect of the bottleneck on the frequency distribution. These profiles also allow us to compare models and data. Consider two frequency vectors 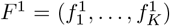 and 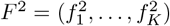. We define the discrepancy between these frequencies as the average relative difference between the Hill numbers of the two vectors over a range of *q* values:

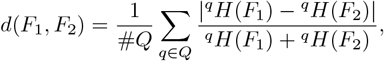

where *Q* is a representative set of values of *q*, and #*Q* is the number of elements in this set. We took *Q* to be a set of evenly spaced values ranging from 0 to 4. This range was empirically selected as experimental data typically showed the highest variations on the interval [0, 4].

Now, consider a stochastic model ℳ(*θ*), parametrised by *θ* ≥ 0, and describing the distribution of the frequencies *F* after the bottleneck event. Given the observed experimental frequencies 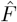, for each fixed *θ*, we define the performance *P* [ℳ(*θ*)] of ℳ(*θ*) as the 0.05 quantile of the distribution of 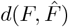 over replicate simulations. Then, the performance *P* [ℳ] of the model ℳ is obtained by choosing the parameter *θ* that leads to the smallest value of *P* [ℳ(*θ*)]. The model is more effective in reproducing 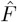 when *P* [ℳ] is small. In other words, we let *P* [ℳ(*θ*)] be the 0.05 quantile of

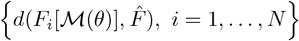

where *F*_*i*_[ℳ(*θ*)] ∈ [0, 1]^*K*^, *i* = 1, …, *N*, are the final frequencies for *N* replicate simulations simulated with model ℳ and parameter *θ*. We then let

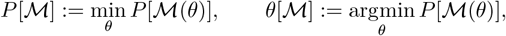

be, respectively, the performance of the model ℳ and the estimated parameter for the model ℳ.

In our situation, we have several experimental frequency vectors 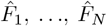, observed under the same experimental conditions. We consider independently the vectors 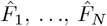. For *i* = 1, …, *N*, let *θ*_*i*_ be the parameter that leads to the smallest value of *P* [ℳ(*θ*_*i*_); *F*_*i*_]. The model performance is then the average of the *P* [ℳ(*θ*_*i*_); *F*_*i*_], *i* = 1, …, *N*, and the output parameter 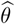 is the average of the *θ*_*i*_.

### 4.8 Mechanistic Models

We propose four stochastic mechanistic models that describe different ways of crossing the barrier. Comparing the performance of these models allows us to select the best one and infer the most probable underlying mechanism. For each model ℳ(*θ*), the parameter *θ* determines the intensity of the bottleneck. In all models, the smaller the value of *θ*, the stronger the bottleneck effect (however, *θ* may scale differently according to the model).

#### Instantaneous bottleneck model ℳ_STAMP_(*θ*_STAMP_)

This is the standard model derived from the original STAMP publication ^1^. From a mechanistic perspective, the model assumes that the founding population after the bottleneck consists of *N* bacteria that cross a barrier simultaneously. These individuals are randomly selected from the initial population according to a multinomial distribution:

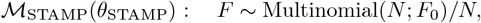

where *F*_0_ represents the pre-bottleneck frequencies and *F* stands for the post-bottleneck frequencies. The parameter of the model is the size of the founding population: *θ*_STAMP_ = *N* . It can be determined as the value 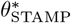 that maximises the performance *P* [ℳ_STAMP_(*θ*_STAMP_)].

#### Constant rate model ℳ_CR_(*θ*_CR_)

Bacteria cross the barrier individually and stochastically at a Poisson rate *µ >* 0, carrying tags selected according to *F*_0_, the pre-barrier frequencies. Once across the barrier, bacteria reproduce deterministically following Malthusian growth at rate *r*. Under those assumptions, the output *X*_*t*_ = (*x*_1_(*t*), …, *x*_*K*_(*t*)) represents the number of bacteria carrying each tag at time *t*. By the properties of Poisson point processes, this is equivalent to assuming that

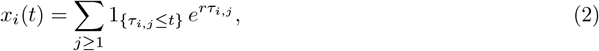

where, for each *i* ∈ {1, …, *K*}, {*τ*_*i,j*_, *j* ≥ 1} are the points of a Poisson process with intensity *µ/K*, and these processes are mutually independent. The corresponding frequencies depend on the observation time and are given by *F*_*t*_ = *X*_*t*_*/n*_*t*_, where *n*_*t*_ = ∑ *x*_*i*_(*t*). The parameter of the constant-rate model is defined as *θ*_CR_ = *µ/r*. It quantifies the relative entry rate of bacteria through the bottleneck compared to their Malthusian growth rate. We interpret *θ*_CR_ as a measure of bottleneck strength: high *θ*_CR_ (*r* ≪ *µ*) means that bacteria cross the bottleneck relatively quickly compared to their growth, making the final population composition closer to the pre-bottleneck distribution. Conversely, low *θ*_CR_ (*r* ≫ *µ*) means that bacteria grow much faster than they cross the bottleneck. Early crossers dominate, as they have more time to expand before new bacteria enter, leading to a founder effect where the first few individuals disproportionately shape the post-bottleneck population structure.

#### Bolthausen-Sznitman model ℳ_BS_(*v*)

To construct the Bolthausen–Sznitman model, we assume that the population is described by a Fisher-KPP equation. The Fisher–KPP equation is a reaction–diffusion equation commonly used to model population dynamics, particularly the evolution of neutral tags during a spatio-temporal expansion process ^40^ occurring at a constant propagation speed. To account for finite-population effects, we introduce stochasticity, leading to a noisy Fisher–KPP equation.

We see the bottleneck as an area where the population must remain very small, and outside of which the population is large. In the experiment, *n* individuals are sampled shortly after the population has crossed the barrier. *What are their genealogies?* No common ancestor can be found outside the barrier, since the population is large in these areas. However, inside the barrier, common ancestors can be found. The connection to our problem is that if two individuals located after the barrier are sampled, then they will necessarily carry the same tag if they have a common ancestor that lived after the tags were attributed. If no such common ancestor exists, the tags of the two sampled individuals are independent. Similarly, if we know the genealogies of a sample of *n* ≥ 3 individuals, then we perfectly know the joint law of their tags. Therefore, understanding how the genealogies of individuals are affected by the crossing of the barrier is sufficient to understand the frequencies of the different tags that we observe at sampling time.

There exist different standard population genetics models for such genealogies. We tested two of them, the Kingman coalescent and the Bolthausen-Sznitman coalescent, which are conjectured or shown to arise in various forms of the noisy Fisher-KPP equations ^13,41,42^. We do not show the Kingman coalescent results in the main material, as it was unable to reproduce diversity profiles similar to the experimental ones (Supplementary Fig. 5, 7). In Section 4.9, we provide a construction of the Kingman coalescent and the Bolthausen-Sznitman coalescent.

The strength of the barrier in this model is measured by the time *T* during which the coalescent process is run to obtain the genealogy of the individuals following the bottleneck, which can be interpreted as the time taken by the population to cross the barrier. We parametrise the model with the speed *θ*_BS_ = 1*/T* . A high *θ*_BS_ means that the population crosses the barrier rapidly, so that the bottleneck is weak.

#### Increasing rate model ℳ_IR_(*µ/r*)

Inspired by the Bolthausen–Sznitman model, the increasing rate model offers a simpler mechanistic interpretation while still capturing a similar behaviour in terms of arrival-time profiles (Supplementary Fig. 4). Conceptually, ℳ_IR_ is a generalisation of the constantrate model ℳ_CR_, in which the entry rate *µ* in the root is replaced by an increasing time-dependent rate function *λ*(*t*), leading to a non-homogeneous Poisson process. Once across the barrier, bacteria again reproduce deterministically following Malthusian growth at rate *r*. Equation (2) thus remains unchanged, but the {*τ*_*i,j*_, *j* ≥ 1} are now the points of an inhomogeneous Poisson point process on R_+_ with intensity *t* 1→ *λ*(*t*).

To maintain consistency with the observed arrival-time profiles in the Bolthausen–Sznitman model, we define the rate as

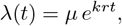

with *k* = 0.78. This value was chosen because it yields the closest match between the arrival-time distributions produced by the ℳ_IR_ model and those observed in the Bolthausen–Sznitman model, even though the underlying parameter values differ between the two models (see Supplementary Fig. 4). This exponential increase in the entry rate over time reflects a progressively intensifying inflow of individuals through the barrier. We also make this choice in order to avoid introducing an additional parameter in the model, to simplify the comparison with other models. The scaling factor *rt* ensures that the frequencies produced by this model remain independent of the Malthusian growth rate *r*.

To clarify this point, consider the non-homogeneous Poisson process with intensity

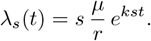

In this process, the arrival time *t*_*i,s*_ of the *i*th bacterium is proportional to 1*/s*, that is, *t*_*i,s*_ = *t*_*i*,1_*/s*. Therefore, the relative frequency of the (*i* + 1)th bacterium to the *i*th is

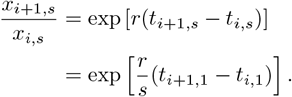

Thus, with the crossing intensity *λ*_*s*_(*t*) and Malthusian growth rate *r*, the relative frequencies are the same as with the crossing intensity *λ*_1_(*t*) and growth rate *r/s*. Taking the scaling factor *s* = *r*, we see that the model with crossing intensity *λ*(*t*) and growth rate *r* is equivalent, in terms of frequencies, to the model with crossing intensity

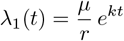

and growth rate 1.

Hence, as in the ℳ_CR_ model, only the ratio *µ/r* is identifiable from the observed frequencies *F* . Consequently, we arbitrarily fix *r* = 1 and *t* = 10, and determine the value 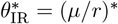 that maximises the performance *P* [ℳ_IR_(*θ*_IR_ = *µ/r*)].

#### Summary of the models

- In ℳ_STAMP_(*θ*_STAMP_ = *N*_*b*_), the founding population after the bottleneck consists of exactly *θ*_STAMP_ bacteria that cross the barrier simultaneously, selected from the initial population according to a multinomial distribution, leading to a stronger founder effect as *θ*_STAMP_ decreases.
- In ℳ_CR_(*θ*_CR_ = *µ/r*), bacteria cross the bottleneck stochastically at a Poisson rate *µ* and then grow exponentially at rate *r*. The ratio *θ*_CR_ = *µ/r* characterises the strength of the bottleneck: a low *θ*_CR_ results in strong founder effects and reduced diversity.
- In ℳ_BS_(*θ*_BS_), bottleneck crossing is modelled as a stochastic propagation process, where the parameter *θ*_BS_ corresponds to the speed at which the population crosses the bottleneck. A lower speed *θ*_BS_ increases the amount of coalescence, leading to a stronger bottleneck effect.
- In ℳ_IR_(*θ*_IR_ = *µ/r*), the bottleneck is crossed at a time-inhomogeneous rate 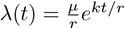, leading to progressively more frequent entries as time increases. Bacteria then grow exponentially at rate *r*. This variable-rate mechanism generalises the constant entry rate model ℳ_CR_ and reproduces arrival profiles similar to those of the ℳ_BS_ model. The parameter *θ*_IR_ = *µ/r* again governs the bottleneck strength, with smaller values indicating a stronger founder effect.

### 4.9 Description of the Kingman and Bolthausen-Sznitman coalescents

#### The Kingman coalescent

The Kingman coalescent is an object that follows the genealogy of current individuals. Let *n* current individuals be labelled by 1, …, *n*. The *n*-Kingman coalescent is a process going backward in time, which, at time *t*, contains the ancestors of the *n* current individuals living at time *t before* the present time. More specifically:

1. The initial state, denoted by Π_0_, is the set of blocks {1}, …, {*n*}. Each block corresponds to a single individual living at the present time;
2. Each *t >* 0 corresponds to a time *in the past*. Then Π_*t*_ is made of blocks of families of individuals living at the present time. For example, if Π_*t*_ = {1, 2}, {3}, {4}, this means that individuals 1 and 2 share a common ancestor that lived at time −*t*; and 3 and 4 do not share a common ancestor with 1, 2 that lived at this time;
3. The dynamics of Π_*t*_ is as follows: any given pair of blocks merges at rate 1. For example, at rate 1, {1, 2} and {3} will try to merge to form {1, 2, 3}. A merger at time *t* means that the two families have found their most recent common ancestor at time *t* before present.

#### The Bolthausen-Sznitman coalescent

The Bolthausen-Sznitman coalescent, similarly to the Kingman coalescent, is a process that goes backward in time and that, at time *t*, regroups individuals who share a common ancestor at time *t before* the present time. However, the mechanism for the merging of blocks is different from that of the Kingman coalescent. While in the Kingman coalescent, only two blocks can merge at the same time, the Bolthausen-Sznitman coalescent allows *multiple mergers*. There are several constructions of the Bolthausen-Sznitman coalescent. It can be seen as the Λ-coalescent with Λ(*dx*) = *dx* ^43^. However, we define it here in another way, which is apter to simulations, using the construction of reference ^44^ (see also the description in reference ^43^). First, construct a tree recursively in the following way. The root is indexed by 1. Once *n* vertices are positioned on the tree, the (*n* + 1)^*th*^ vertex chooses its parent uniformly at random among the *n* vertices of the tree. Each vertex of the tree corresponds to a block; initially, all blocks contain only a singleton.

The merging procedure is as follows. Each edge of the tree carries an exponential clock with rate 1. If the clock of an edge rings, the edge is “lifted”; namely, all descendants of the edge are absorbed by the parent of the edge. The procedures goes on until there is only one vertex left (see Supplementary Fig. 10).

### 4.10 Estimation of *µ* and *r* using the constant rate or increasing rate models

We consider here the models ℳ_CR_ and ℳ_IR_. The growth rate is estimated as follows. For each sample, we measure the bacterial population associated with the first arrival as *f*_1_ × (bacterial load), where *f*_1_ is the proportion of the majority tag. Assuming Malthusian growth with rate *r*, this population size equals 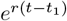, where *t* is the observation time and *t*_1_ is the arrival time of the first tag under the model. For the model ℳ_CR_, we use *t*_1_ ≈ *E*[*t*_1_] = 1*/µ* = 1*/*(*rθ*). For the model ℳ_IR_, we approximate *t*_1_ by the median value of the first arrival time, that we compute as follows. We use the cumulative intensity function:

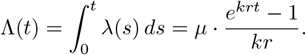

The cumulative distribution function of the first arrival time is:

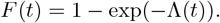

The median *t*_med_ satisfies *F* (*t*_med_) = 0.5, *i.e*.,

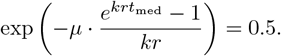

Thus,

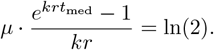

and

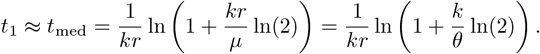

We then approximate *r* using the formula:

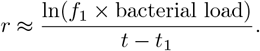

For the model ℳ_CR_, this leads to

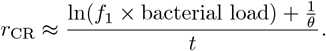

For the model ℳ_IR_, we get:

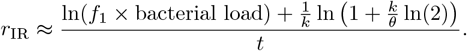

## Data availability

All processed sequence counts and all associated metadata were kept unmodified in a all-in-one Excel data sheet available in supplementary data. Outputs from the time-resolved analysis algorithms are conserved in a semi-controlled local environment (R project “SV bottleneck analysis 2024”) in supplementary documents. The raw sequences data and corresponding metadata for this study have been deposited in the European Nucleotide Archive (ENA) at EMBL-EBI under accession number PRJEB102723 (https://www.ebi.ac.uk/ena/browser/view/PRJEB102723).

## Code availability

The code for all analyses is made available in an annotated R Markdown notebook in supplementary documents. The notebook is contained in a semi-controlled local environment inside the R project “SV bottleneck analysis 2024” allowing to reproduce the integrality of the presented analyses, figures and statistical tests from the original data sheet (tested environment: Windows 11, RStudio, R version 4.2.2, Python version 3.11.2). The code for the processing of raw sequences data is also provided in the file “SV nanopore pipeline 2024” (tested environment: Shell script in Linux Ubuntu with a Conda environment, Python version 3.11.2).

## Authors contribution

S.V. and N.P. designed the plant infection experiments. S.V. and N.P. constructed and validated the biological material. S.V., I.M. and N.P. performed the plant infection experiments. S.V. sequenced and processed the sequences data. N.B., R.F. and L.R. designed and formalised the model testing approach and the time-resolved algorithms. S.V., N.B., R.F. and L.R. wrote the code of the notebook. S.V., N.B., R.F., L.R. and N.P. discussed and analysed the data. S.V. wrote an original draft. All authors corrected and contributed to the manuscript.

## Acknowledgements

The authors thank Dr. Karthik Hullahalli (ID), Dr. Kelly E. R. Bachta (ID) and Dr. Michael A. Bachman (ID) for their assistance in accessing the animal infection data reanalysed in the study. The authors thank Dr. Alice Guidot (ID), Dr. Philippe Remigi and Dr. Stéphane Genin for their insightful discussions regarding the manuscript. The authors thank Baptiste Mayjonade (ID) and Sandra Moreau for their assistance and the numerous insights during the establishment of Nanopore sequencing protocols.

This work was supported by the following grants. S.V. was supported by a PhD grant from the Ministére de l’Enseignement Supérieur, de la Recherche et de l’Innovation. N.P. and S.V. were supported by a seed funding “STAMP” from INRAE department Plant Health and Environment. N.B., R.F. and L.R. were supported by the ANR project ReaCh (ANR-23-CE40-0023-01). N.B. and R.F. were supported by the Chaire Modélisation Mathématique et Biodiversité (École Polytechnique, Muséum national d’Histoire naturelle, Fondation de l’École Polytechnique, VEOLIA Environnement).

## Supplementary Material

**Supplementary Table 1.** Score of error (mean ± half-width of the 95% confidence interval) and mean normalised bottleneck parameter *θ* estimated for each infection condition from 2 and 5 (mean ± half-width of the 95% CI). Values of *θ* are normalised by the control condition M_Intact_27. Condition *H_Intact_27* was tolerant and did not yield sufficient colonisation to allow sequencing of the diversity and estimation of *θ*.

| Model | Average error | $\theta$ M-Intact-27 | $\theta$ M-Cut-27 | $\theta$ H-Cut-27 | $\theta$ M-Intact-32 | $\theta$ M-Cut-32 | $\theta$ H-Intact-32 | $\theta$ H-Cut-32 |
| --- | --- | --- | --- | --- | --- | --- | --- | --- |
| $\mathcal{M}_{\text{STAMP}}$ | $0.20 \pm 0.01$ | $1.00 \pm 0.58$ | $3.08 \pm 1.50$ | $2.20 \pm 1.29$ | $0.91 \pm 0.54$ | $2.65 \pm 1.20$ | $0.85 \pm 0.80$ | $2.57 \pm 1.20$ |
| $\mathcal{M}_{\text{CR}}$ | $0.12 \pm 0.01$ | $1.00 \pm 0.58$ | $2.14 \pm 0.81$ | $1.72 \pm 0.77$ | $0.92 \pm 0.53$ | $2.17 \pm 0.77$ | $0.74 \pm 0.65$ | $2.18 \pm 0.79$ |
| $\mathcal{M}_{\text{BS}}$ | $0.06 \pm 0.01$ | $1.00 \pm 0.29$ | $2.20 \pm 0.90$ | $2.35 \pm 1.01$ | $0.97 \pm 0.37$ | $1.98 \pm 0.64$ | $1.02 \pm 0.39$ | $1.91 \pm 0.51$ |
| $\mathcal{M}_{\text{IR}}$ | $0.08 \pm 0.02$ | $1.00 \pm 0.74$ | $1.92 \pm 0.59$ | $1.98 \pm 0.62$ | $1.14 \pm 0.60$ | $2.11 \pm 0.51$ | $0.75 \pm 0.57$ | $2.33 \pm 0.47$ |

**Supplementary Figure 1.**
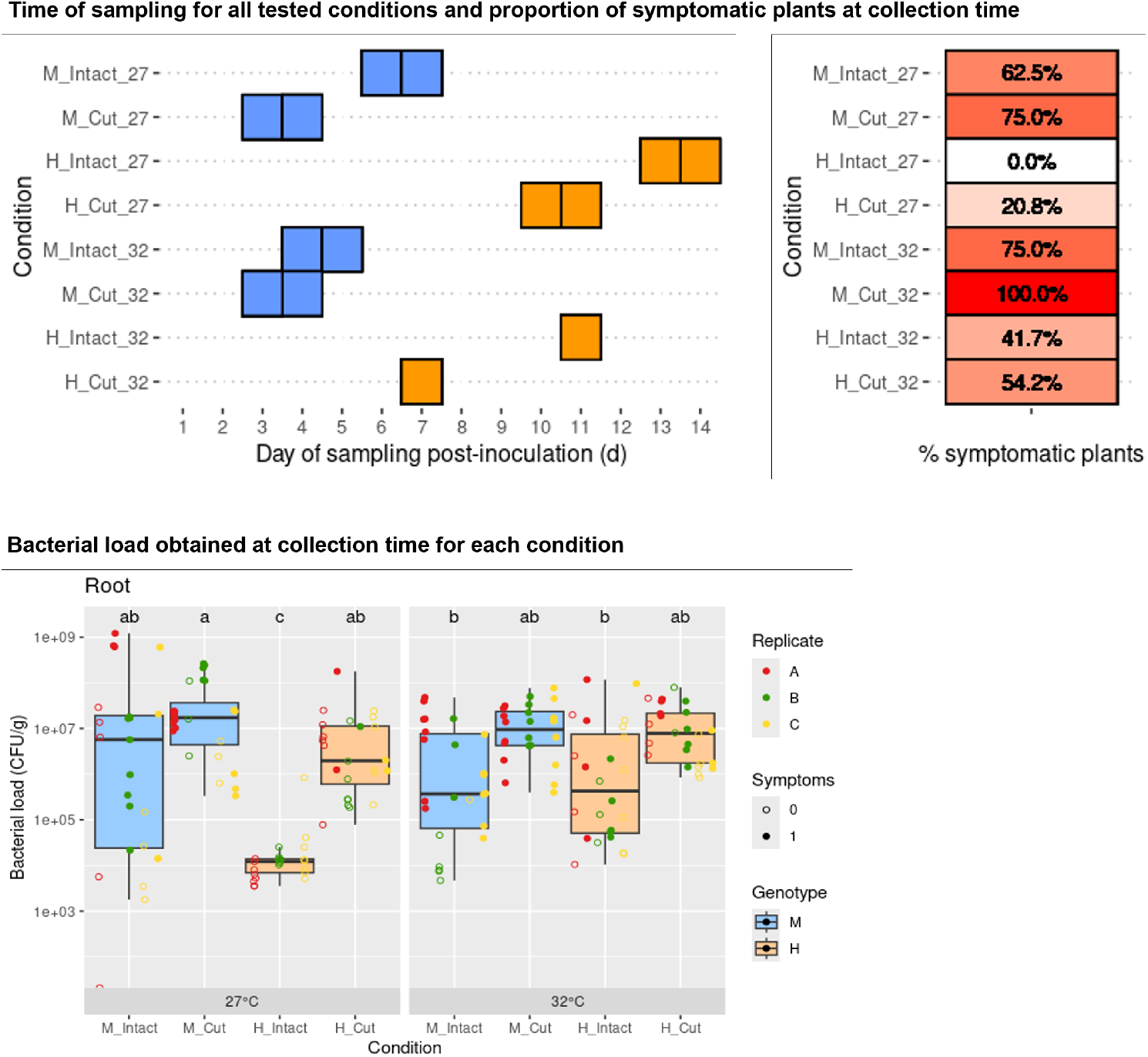
Day of sampling and proportion of symptomatic plants corresponding to the all conditions in Fig. 5. Conditions *M_Intact_27* and *M_Cut_27* correspond to the data shown in Fig. 2). At the first sign of symptoms on a batch of a replicate (replicate A, B or C), the *n* = 8 plants of the batch were collected. Batches that showed no symptoms were otherwise collected at 13-14 DPI. Bacterial load was systematically quantified. Conditions that were sampled earlier showed systematically higher or similar loads compared to conditions sampled later. Ultimately, the plants that were collected earlier also corresponded to a faster colonisation in CFU but also had faster symptoms development indicating a good representation of a gradient of susceptibility to tolerance.

**Supplementary Figure 2.**
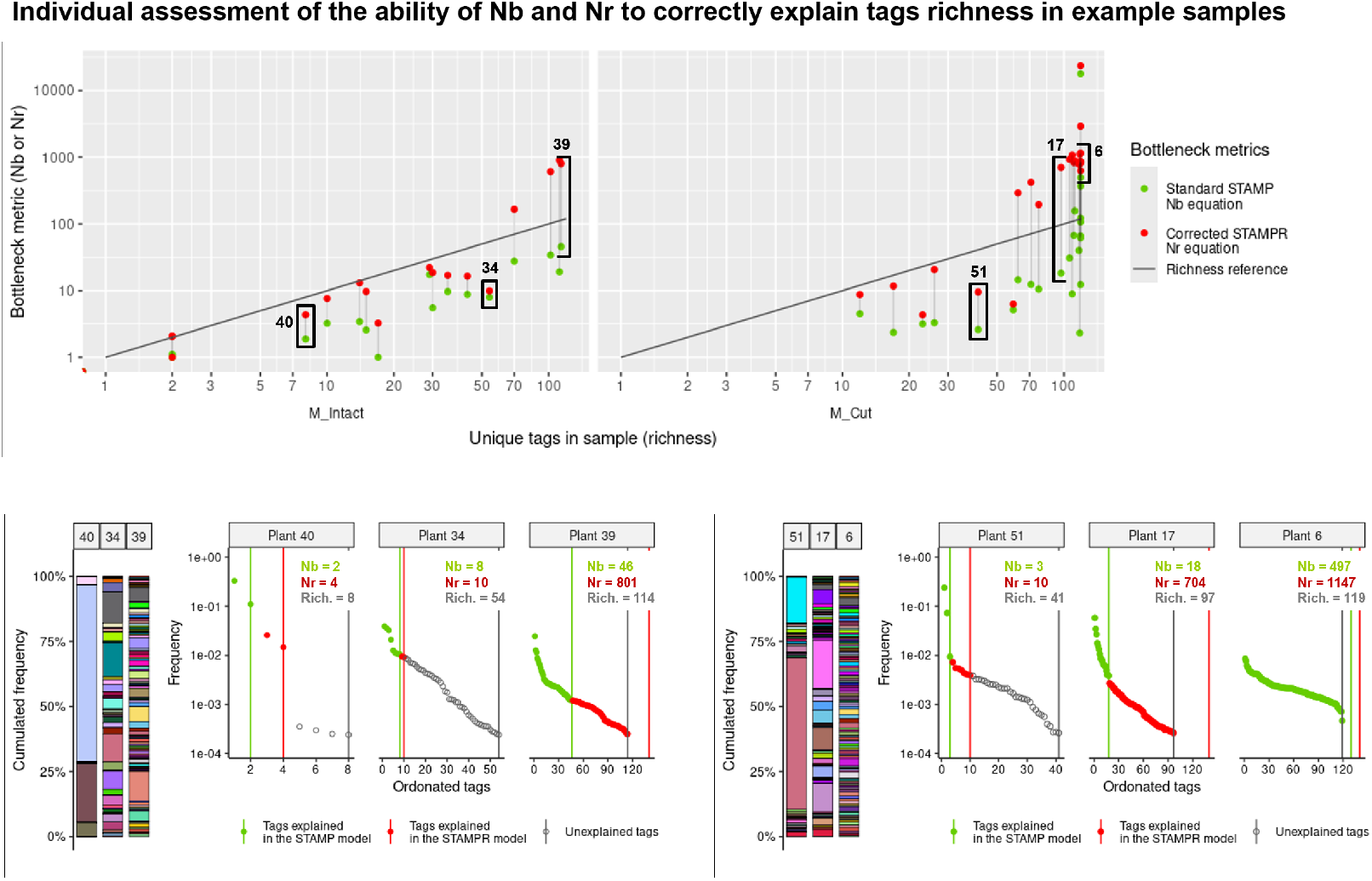
Assessment of the behaviour of *N*_*b*_ and *N*_*r*_ as a function of richness in sample (log-log scale). A reference line *y* = *x* represents richness, values of *N*_*b*_ or *N*_*r*_ under the line are lower than richness in the same sample. A line connects *N*_*b*_ and *N*_*r*_ calculated from the same sample. Illustrative examples of diversity profiles are shown in three samples of intact Marmande plants (plants 40, 34 and 39) or three samples of root-cut Marmande plants (plants 51, 17, 6) (log scale). Tags are sorted in decreasing frequency. Green and red lines represent the number of tags that are successfully represented by *N*_*b*_ and *N*_*r*_ respectively.

**Supplementary Figure 3.**
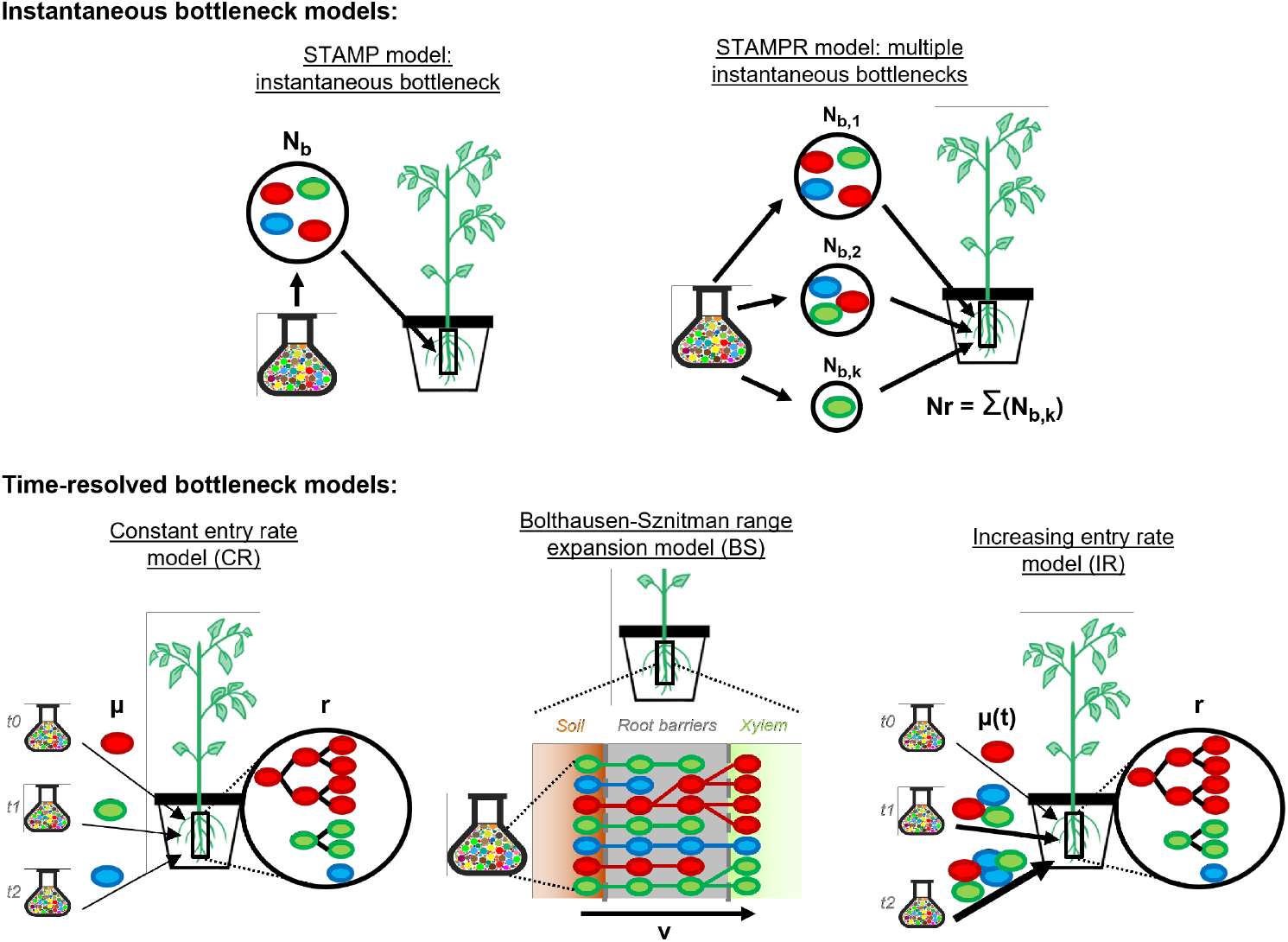
Models tested in the study. **“Instantaneous bottleneck” (STAMP) model** underlying the *N*_*b*_ equation ^1^. *N*_*b*_ represents a “number of founders”, implicitly considering a bottleneck as a single, instantaneous founder event separating two non-overlapping populations ^11,12^. Four assumptions are implied ^11^: (1) neutral diversity: all tags have identical growth and virulence fitness; (2) compartmentalised space: soil and root are distinct compartments; (3) spatial homogeneity: all tags have equal access to root, tissue complexity is ignored; (4) instantaneous barrier: a single, instantaneous entry with no secondary infections. Under these assumptions, entry is quasi-instantaneous and decoupled from time-dependant effects. Assumptions (1) and (2) are supported by knowledge of the infection and the neutral tags collection. Assumptions (3) and (4) remain untested in the literature and are contradicted by the observation of motility and growth during colonisation ^5,6,7,8^. **“Multiple instantaneous bottlenecks” (STAMPR) model** empirically interpreted from the *N*_*r*_ and *N*_*s*_ algorithms ^2^. Colonisation occurs over *k* events of instantaneous entry, each contributing to *N*_*b,k*_ new founders, each *k* subpopulation representing a proportion of the total population. Assumptions of instantaneous entry are inherited from the STAMP model. As STAMPR is an empirical correction rather than a mechanistic model, it cannot be mathematically investigated beyond empirical checks. **“Constant entry rate” (CR) model**. Plant colonisation is the combination of a constant entry process and a pure growth process. Pathogens enter the roots at a constant entry rate (or permeability) *µ*_CR_ while already present pathogens grow at an exponential rate *r*_CR_. The bottleneck metric *θ*_CR_ = *µ*_CR_*/r*_CR_ is the ratio of entry over occupation of space by growth and informs on the quantity of unique cells entering the plant before saturation. Low *θ*_CR_ indicates a colonisation that is mostly governed by growth (a stringent bottleneck). **“Bolthausen-Sznitman” (BS) model**. Root barriers breaching is modelled by a range expansion in a finite population ^13,45^. Pathogen cells migrate randomly within the barrier and divide at a density-dependent rate (cells in more crowded areas divide less often). Cells that manage to cross this barrier then colonise the plant. Resulting sampling of tags can be obtained using the Bolthausen-Sznitman coalescent (see Section 4.8). The bottleneck metric *θ*_BS_ = *v* is the relative speed at which pathogens can cross the root barriers and reach the final compartment. Low value indicates slow breaching of the roots barriers allowing for more branching of pathogen cells lines (a stringent bottleneck). **“Increasing entry rate” (IR) model**, a generalisation of the CR model. Colonisation of the plant is a combination of an exponentially increasing entry process and a pure growth process. As entry increases over time, an entry rate (or permeability) multiplier *µ*_IR_ governs the acceleration of entry of pathogen over time. Simultaneously, already present pathogens grow at an exponential rate *r*_IR_. Similarly to the CR model, low *θ*_IR_ = *µ*_IR_*/r*_IR_ represents a stringent bottleneck.

**Supplementary Figure 4.**
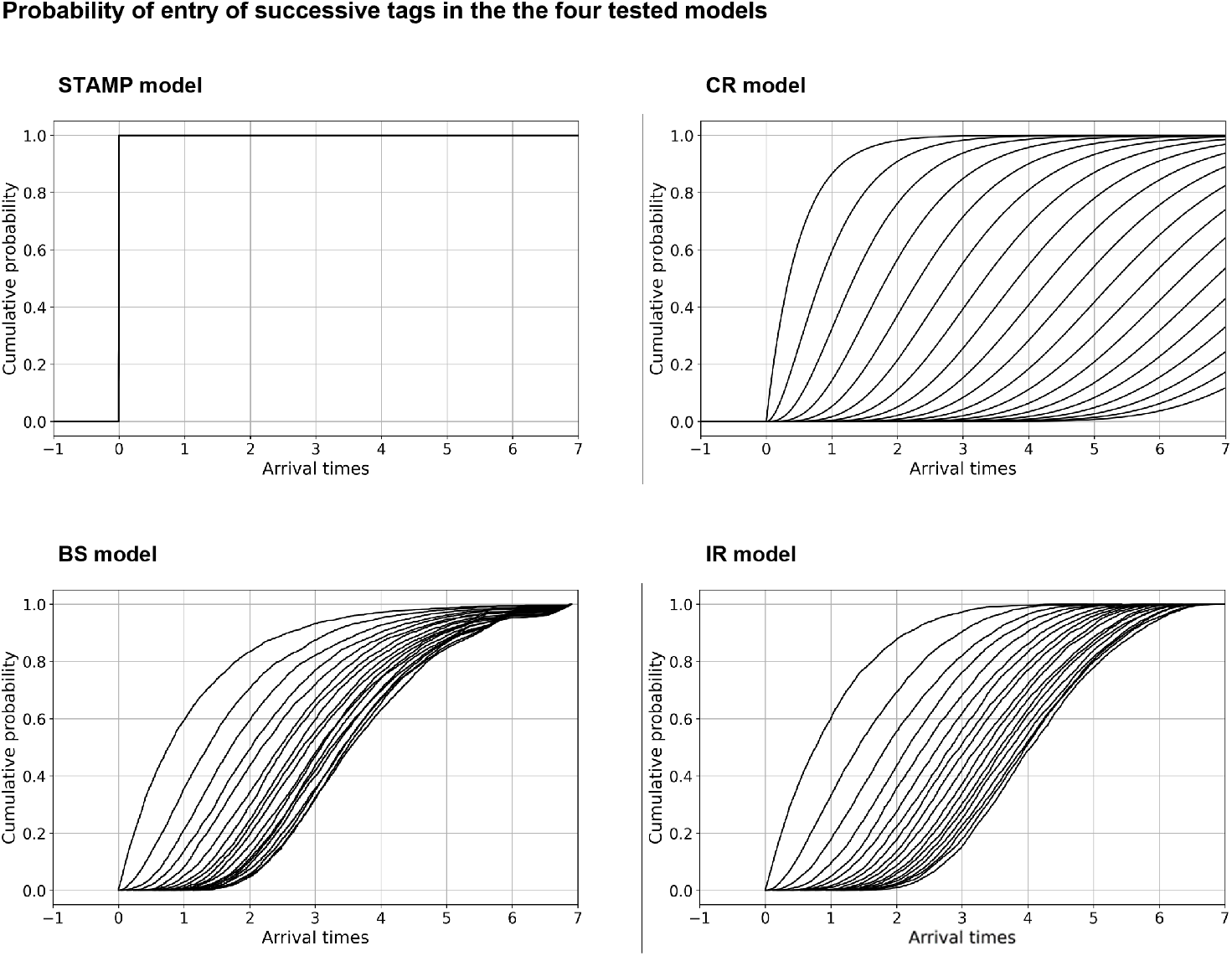
Comparison of arrival time distributions under the four tested models. The first bacterium enters the root at a normalised time 0. Each curve represents the Cumulative Distribution Function (CDF) of the arrival time of the *k*^th^ bacterium relative to the first one. In this illustration, the parameter values are: *µ*_CR_*/r* = 2 for ℳ_CR_, *v* = 10 for ℳ_BS_ and *µ*_CR_*/r* = 0.1 for ℳ_IR_.

**Supplementary Figure 5.**
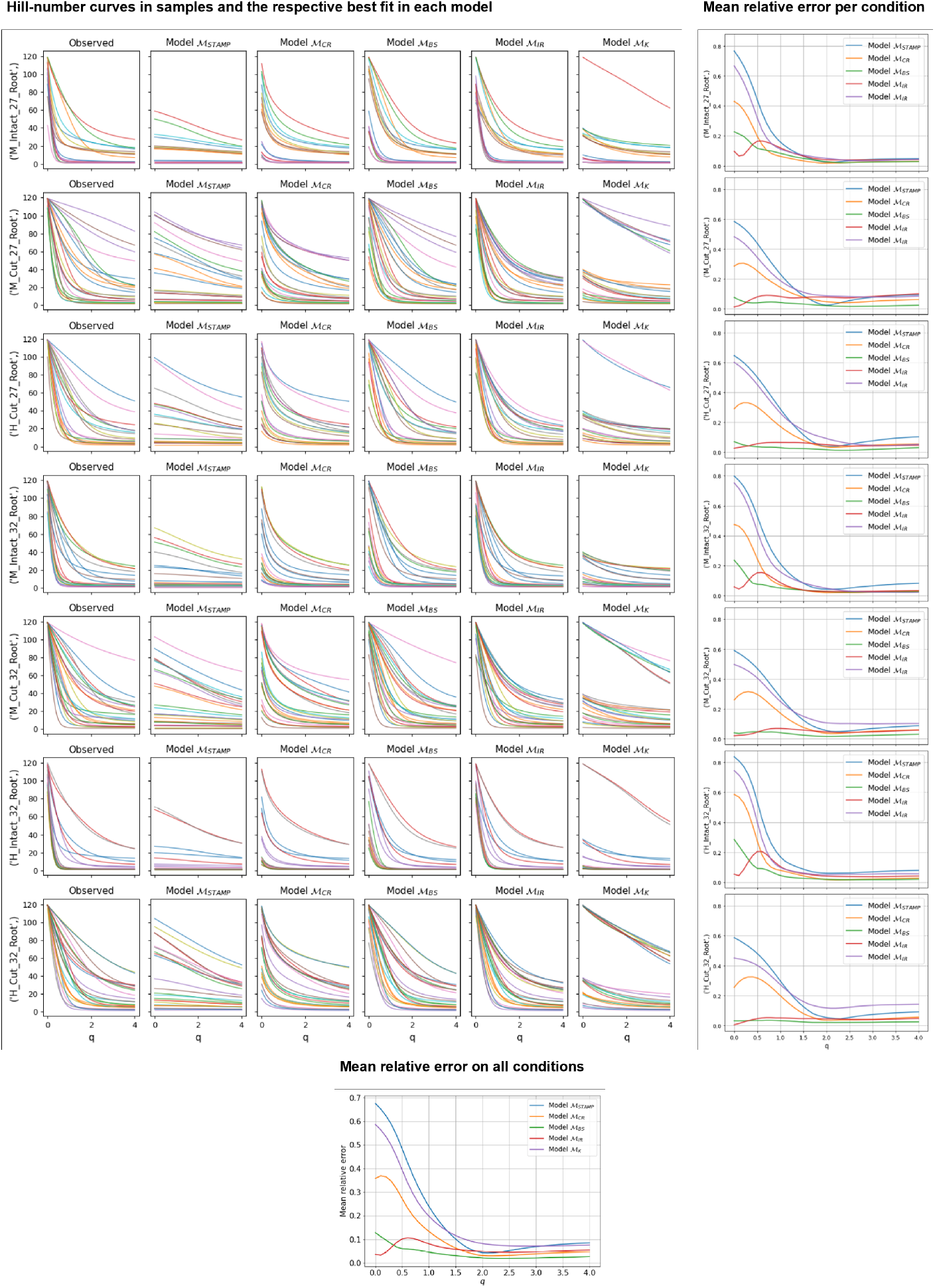
Diversity profiles in samples and simulations according to the Hill numbers from order *q* = 0 (giving more importance to richness) to order *q* = 4 (giving more importance to evenness). For a given order *q*, a high value ^*q*^*H* indicates a high diversity. Each colour corresponds to a sample profile from Fig. 5 including those of Fig. 2. An additional model ℳ_K_ (Kingman model) was tested but is not presented in the main text. ℳ_K_ provides a simpler equation to describe biological processes of ℳ_BS_ but was globally unable to accurately describe the data. For each model, a score of error represents the difference between the Hill-number curve in observed data versus in best fit from a model. The error can be averaged on each condition or on all samples.

**Supplementary Figure 6.**
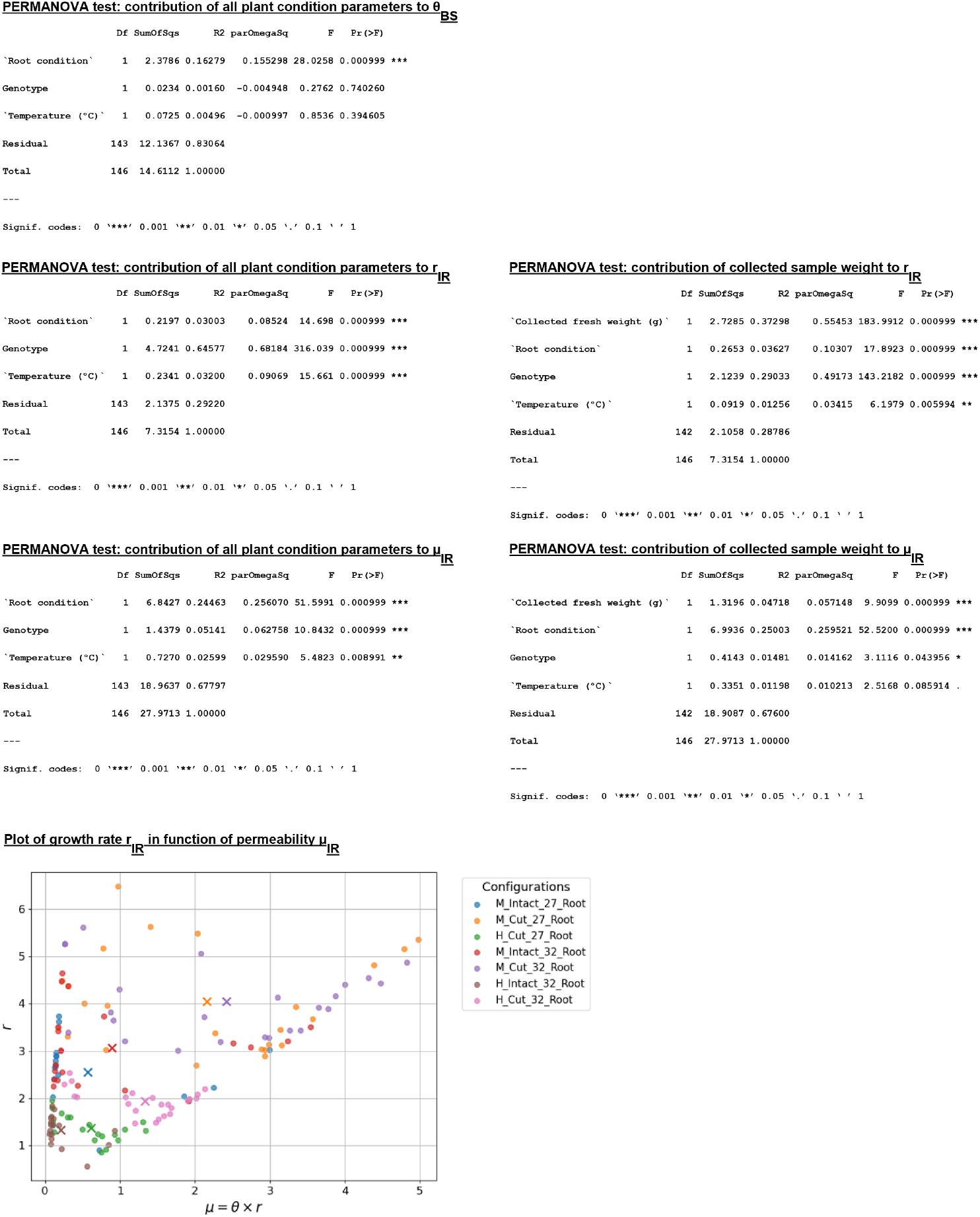
Statistics performed on the estimated parameters from Fig. 5c, 5d, 5e. Non-parametric PERMANOVA tests were performed to see the relative contribution of the three tested biological factors (*plant genotype, roots condition* and *temperature*) on *θ*_*BS*_, *µ*_IR_ and *r*_IR_. Another PERMANOVA test was performed on *µ*_IR_ and *r*_IR_ with the addition of root sample weight as a proxy for a *plant size* factor. As a guide, “Pr (¿F)” and “R2” respectively represents the p-value and the coefficient of determination *R*^2^ for each tested factor. Additionally, “parOmegaSq” represents the effect size *ω*^2^ of each factor (*>* 0.1: small effect; *>* 0, 3: medium effect, *>* 0.5 large effect). Growth rate *r*_IR_ was plotted in function of permeability *µ*_IR_. Each cross on the graph represents the average value for each condition. Values of *r*_IR_ and *µ*_IR_ correlate positively (Spearman ranks correlation, *R*^2^ = 0.40, *p* = 5.97 × 10^−7^).

**Supplementary Figure 7.**
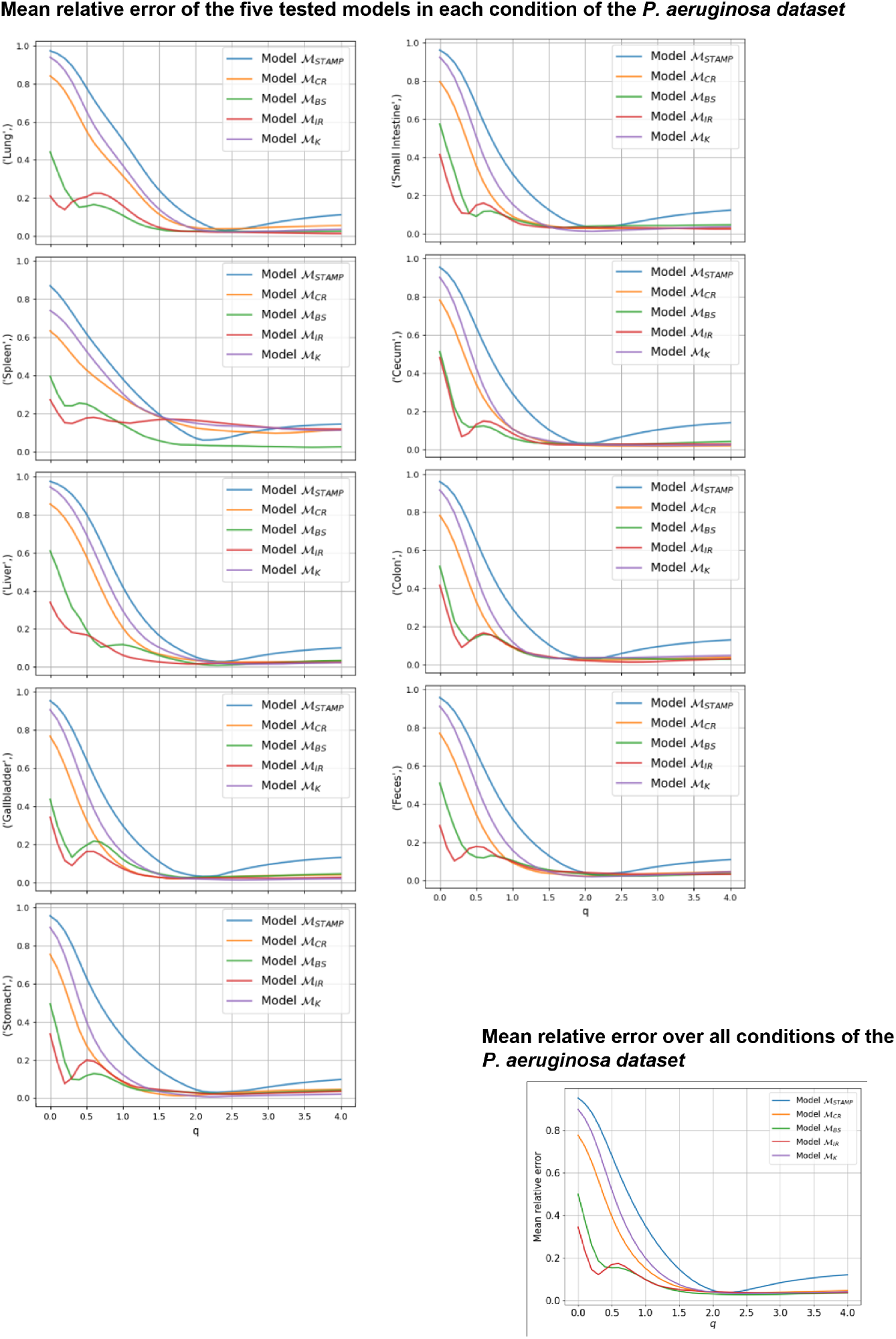
Score of error for the models in the range *q* = 0 to *q* = 4 on the *P. aeruginosa* infection dataset reanalysed in Fig. 6. An additional model ℳ_K_ (Kingman model) was tested but is not presented in the main text. ℳ_K_ provides a simpler equation to describe biological processes of ℳ_BS_ but was globally unable to accurately describe the data. For each model, a score of error represents the difference between the Hill-number curve in observed data versus in best fit from a model. The error can be averaged for each organ or on all samples.

**Supplementary Figure 8.**
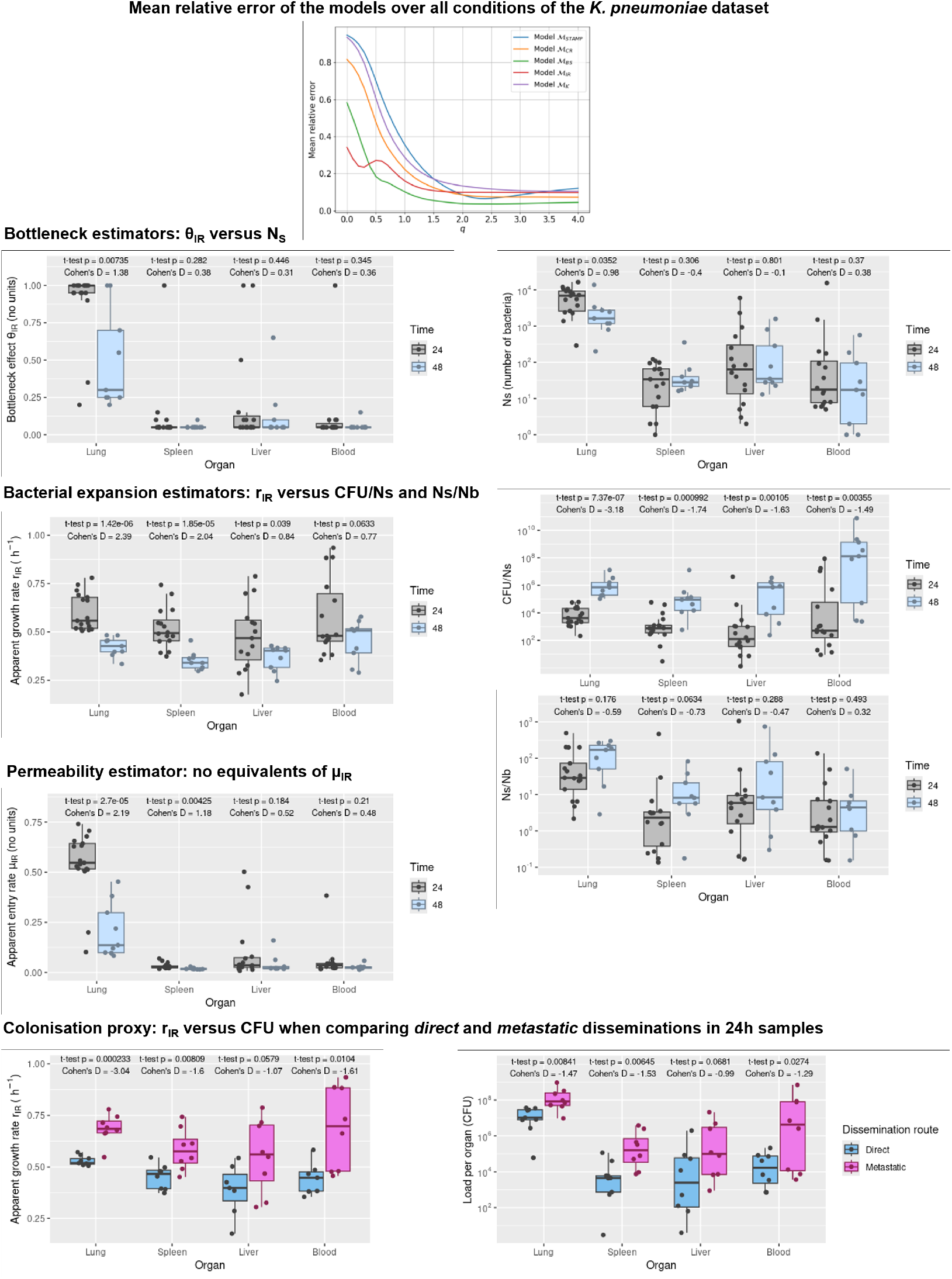
Time-resolved models outperform STAMPR to describe mouse-*K. pneumoniae* infection (original data from Fig. 1 of article ^10^ (*n* = 17 and 9 mice sampled at 24h and 48h respectively). The averaged score of error of the models over *q* = 0-4 showed a better fit of ℳ_IR_ over ℳ_STAMP_. Model-based estimators (*θ*_IR_, *µ*_IR_, *r*_IR_) were compared to their closest counterpart in STAMPR as follow: *θ*_IR_ *vs. N*_*s*_ for the bottleneck effect; *r*_IR_ *vs. CFU/Ns* and *Ns/Nb* for bacterial expansion; *µ*_IR_ had no equivalent in STAMPR. A comparison of *r*_IR_ *vs. CFU* showcases colonisation of organs at 24h between the infections characterised as *direct* or *metastatic* in article ^10^. Performances of estimators were assessed using Student’s two-sided t-test and Cohen’s D effect size between samples collected at 24h and 48h post-infection. For STAMPR metrics (*N*_*s*_, *CFU/N*_*s*_, *N*_*s*_*/N*_*b*_, *CFU*), tests were performed after a log10 transformation (as in article ^10^). Greater values of Cohen’s D (positively or negatively) indicates better statistical power at discriminating 24h and 48h.

**Supplementary Figure 9.**
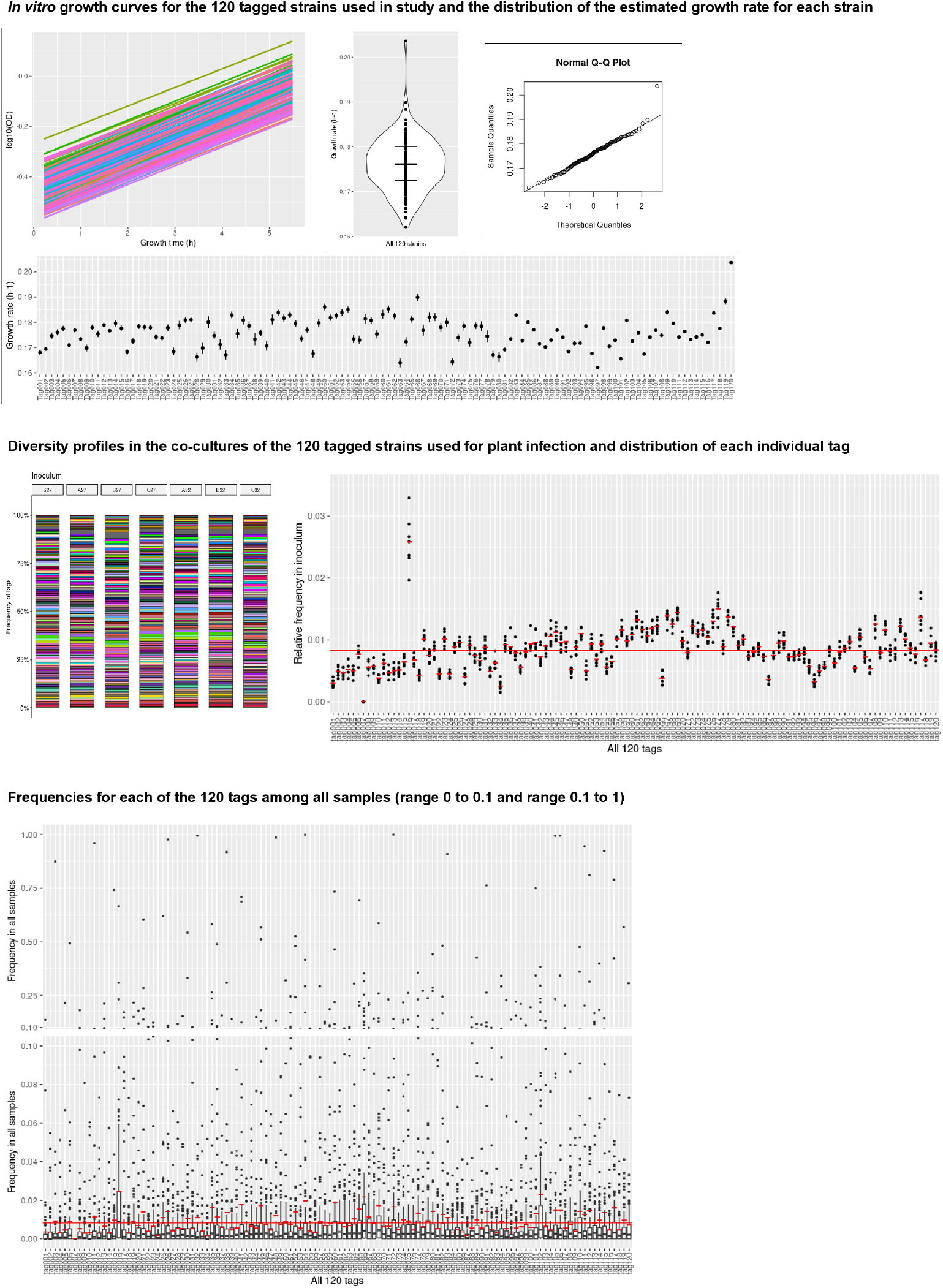
Validation of the 120 tagged strains *in vitro*. 120 isogenic tagged strains were first grown individually in microplate cultures and monitored for their growth curve through *OD*_600*nm*_ (logarithmic scale). Growth of the 120 strain was investigated in a violin plot (thick bar: median; thin bars: 1st and 3rd quartile) and in a quantile-quantile plot against a normal distribution. No visual deviation from normality could be observed suggesting a similar growth behaviour between the strains (average growth rate *r* ≈ 0.176*h*^−1^). Estimation of the growth rate was also individually investigated for each strain (error bars: standard deviation on the growth rate). **Validation of the 119-tags co-cultures *in vitro***. All co-cultures used for plant inoculation were sequenced. Visually, diversity profiles were stable across each independently prepared inoculum verifying that no technical bottlenecks were introduced at inoculum preparation. The relative frequencies at which tags appear in inocula are plotted (red bars: average across inocula; global red line: perfect frequency of 1/120). Tag007 was mistakenly forgotten from the initial pool while Tag016 was mistakenly added twice to the pool. The average frequencies did not significantly differ from the expectation of a distribution centered on a median of 1*/*120 (Wilcoxon-Mann-Whitney test, two-sided, *mu* = 1*/*120, *p* = 0.74), overall showing no indications that the 119 final tags differ in growth in co-culture. **Validation of the 119-tags collection *in planta* for the study of population dynamics**. Relative frequencies of tags in all samples pooled from Fig. 5 including samples from Fig. 2. The frequencies are observed independently of condition (total: *n* = 131 sequenced samples on 6 time-independent infections; red bars: average frequency across samples; global red line: perfect frequency of 1*/*120). For visual clarity, the frequencies are separated in two ranges (0 to 0.1 and 0.1 to 1) and the points within the boxplots are not shown. Under an expectation of neutral fitness, the frequencies averaged over a large number of events of infection should tend toward the initial inocula frequencies regardless of condition. The average frequencies did not significantly differ from the expectation of a distribution centred on a median of 1*/*120 (Wilcoxon-Mann-Whitney test, two-sided, *mu* = 1*/*120, *p* = 0.38), overall showing no detectable difference from the assumption of neutral fitness at a collection level nor systematic sequencing bias.

**Supplementary Figure 10.**
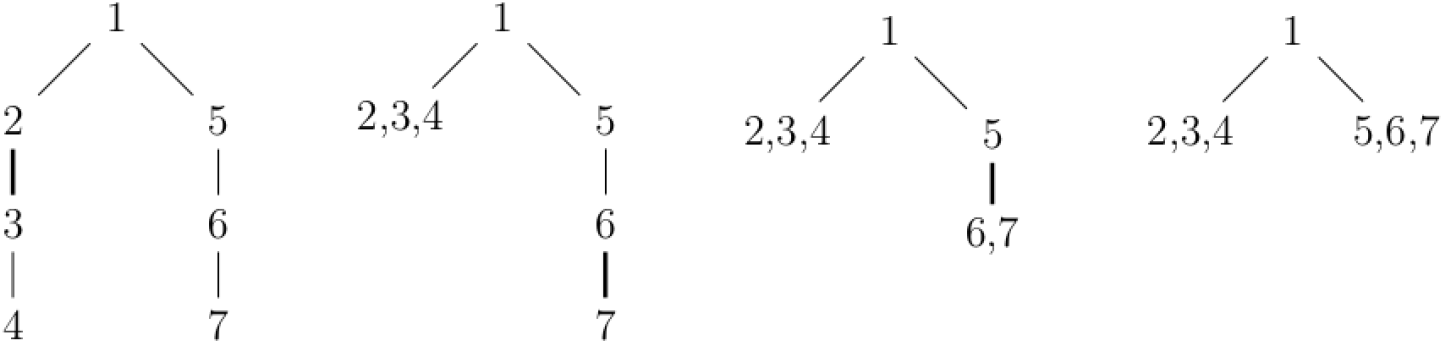
The construction of the Bolthausen-Sznitman coalescent of reference ^44^. Each vertex corresponds to an individual. The initial tree is a random tree where the individual *n* + 1 chooses its parent uniformly at random among 1, …, *n*. At each step, an edge (drawn thick on the picture) is lifted.

