## Supplementary material for "Time-resolved bottlenecks reveal concurrent pathogen entry and growth during bacterial infection dynamics": Code 1: Analysis_2024.Rmd

````
---
title: "Time-resolved bottleneck analysis in bacterial infection dynamics - Annotated notebook"
authors: "Sylvain Vicente, Nathanael Boutillon, Raphael Forien, Lionel Roques"
date: "11/2025"
output:
  pdf_document: default
  html_document: default
header-includes:
  - |
    ```{=latex}
    \usepackage{fvextra}
    \DefineVerbatimEnvironment{Highlighting}{Verbatim}{
      breaksymbolleft={}, 
      showspaces = false,
      showtabs = false,
      breaklines,
      commandchars=\\\{\}
editor_options: 
  chunk_output_type: inline
---

# General information

This code is distributed under a Creative Commons CC BY 4.0 license (https://creativecommons.org/licenses/by/4.0/). You are free to share and adapt the code for any uses. If this code substantially participated your work, please consider citing the corresponding article.

The code and data used for the analyses performed in the article "Time-resolved bottleneck analysis in bacterial infection dynamics" are compiled in a partially controlled environment (the R project "SV_bottleneck_analysis_2024"). In this project, the annotated notebook "Analysis_2024.Rmd" controls all the code used for the analyses. It also contains some complementary technical details that could not be described in the manuscript, including some development regarding the STAMPR framework and the newly proposed "time-resolved bottleneck" approach.

This notebook was co-written by Sylvain Vicente, Nathanael Boutillon, Raphael Forien and Lionel Roques.

This notebook is a combination of R and Python scripts and was initially written and run on R version 4.2.2 and Python version 3.11.2.

This notebook was made with transparency and reproducibility in mind. If you find any errors or need help to reuse this code, please consider writing to Nemo or me.

--------------
General biological context:
Experiments of infection of tomato plants were performed using a neutral barcode collection of ~120 tags in Ralstonia pseudosolanacearum. The objective was to test four factors on the infection dynamics of the pathogen inside the plant:
- Root condition at the infection (intact roots vs cut roots).
- Plant genotype (susceptible Marmande tomato plants vs. tolerant Hawaii7996 tomato plants).
- Temperature during infection (27°C vs 32°C).
- Observed zone of infection (root vs stem).

Following these infections, we developed mathematical analyses (the time-resolved bottleneck framework) to better understand the mode of entry of the pathogen in the plant and infer on quantitative metrics for the infection.

--------------
The files in the R project is divided as follow:
- The folder "Data" contains all the counts tables and metadata used in the article. All data and metadata regarding infection experiments of tomato plants by R. pseudosolanacearum are compiled in the table "Data_tomato_2024.xlsx". Data regarding growth curves of R. pseudosolancearum in vitro are contained in the table "Data_120_growth_2024.xlsx". Additionally, the subfolders "Bachta_natcomm_2020_data" and "Holmes_natcomm_2025_data" contain the datasets about infection experiments by P. aeruginosa or K. pneumoniae and were extracted from their respective articles (supplementary data from Bachta et al, Nature Communications, 2020. and GitHub https://github.com/hullahalli/stampr_rtisan from Holmes et al, Nature Communications, 2025). We thank Dr. Kelly E. R. Bachta for providing the table "2025_9_9_NatComms_CFU_Data_Bachta.xlsx" containing complementary CFU data that were not published within article Bachta et al, Nature Communications, 2020.

- The folder "Analysis_output" contains all analysed data tables. The folder mainly contains output from the calculation of Nb, Nr or Ns as well as the formatted .csv tables used for the "model testing" algorithms.

- The different folders starting with "Models_output..." contain the output of the model testing algorithms on the different datasets. The folder mainly contain the values of the estimated parameters $\theta$, $\mu$ and $r$ and the intermediate files for each simulation. In order, the folder "Models_output" corresponds to the main R. pseudosolanacearum dataset, the extension "_kinetic" relates to the plants infected by R. pseudosolanacearum and collected over time, the extension "_bachta" relates to the P. aeruginosa dataset and the extension "_holmes" relates to the K. pneumoniae dataset. An additional extension "_correct_inoc" corresponds to the same dataset but analysed with a corrected version that takes into account the starting inoculum frequencies.

--------------
About the content of the annotated notebook:
The initial setup chunk loads all the R libraries and the Python dependencies used in the different sections. It will be necessary that you install R and Python if you want to run this code. The initial setup chunk also loads and preprocesses the R. pseudosolanacearum experiments tables. This completes some very basic data preprocessing that was also performed directly on the Excel table using Excel formula (such as for the calculation of a bacterial load using counted CFU and samples weight which can be inspected on the Excel table itself). The initial setup chunk is written entirely in R. All subsequent chunks are separated in independent "sections" that can be run independently.

The first section ("# Section R analysis") is dedicated to the application of the STAMP and STAMPR framework on the R. pseudosolanacearum dataset as well as the production of most of the descriptive figures regarding the experiment (CFU, barcode frequency barplots, in vitro growth curves...). This section is written entirely in R.

All next sections are applications of the model testing algorithms on the different datasets. They contain a mix of R and Python chunks. Each section is essentially a duplication of the same algorithms and applied on each dataset, at the exception of the following differences. All datasets were analysed twice, one section without correction for the starting inoculum frequencies (assumption of a perfect equi-proportion of starting tags) and a section with updated algorithms taking into account inoculum frequencies. On the K. pneumoniae dataset specifically, the section with inoculum correction also contains additional multithreading functions because of the large size of the dataset.

--------------
Guidelines to apply the model testing approach to different dataset (taking the example of the R. pseudosolanacearum dataset):
- Prepare an explicit data table that can be extracted in the format of a R data.frame. R has packages to extract .csv, .tsv, .xlsx... into data.frame. The data table should be organised with one biological sample per row and with the different tested conditions in column (see example of "SV_bottleneck_analysis_2024/Data/Data_tomato_2024.xlsx" extracted in the data.frame "tab_tot"). If you have multiple technical replicate of sequencing of an identical biological sample, you can sum up or average the tag counts as if it was a single sequencing. If you test several factors (i.e. temperature + root conditions + genotype), combine them so that all tested configurations are contained in one single column.

- Convert the table to (or keep it as) an appropriate .csv table (example: "SV_bottleneck_analysis_2024/Analysis_output/tab_tot.csv") that can be used by the model testing algorithms. Do the same for the inoculum table if you intend to use the algorithms with inoculum correction (example: "SV_bottleneck_analysis_2024/Analysis_output/tab_inoc.csv").

- Pick one version of the model testing algorithms that you want. Each section is a version applied on a dataset (example: "# Section reanalysis of the tomato dataset including correction with the starting inoculum frequencies"). The first Python chunk is an initialisation chunk where you will need to specify, for each used variable, the corresponding column in the .csv (!!! be careful, in Python, column numbers start from 0 unlike in R !!!). If one column does not apply to your experimental system, specify a mock column with only integer 1 in it. If you use CFU instead of bacterial load (CFU/g), put the CFU values in the bacterial load column and use the mock column for sample weight. Finally, specify all tested configurations in the appropriate list.

- The subsection II ("# II. Model testing on the experiment of kinetics of infection in tomato plants") contains the bulk of the model testing algorithms. The code is in Python. The first chunk is an initialising the helper functions, don't forget to specify the appropriate target .csv for the samples table and the inoculum table. The chunks for each models can then be launched individually. Modify appropriately the model output directories and set the appropriate tested $\theta$ range and its increment. For $M_{IR}$, you can also modify the input parameter $k$. When a model is done running, the output is saved. Typical run times (processor AMD EPYC7302 CPU, single-thread) for the R. pseudosolanacearum dataset with inoculum correction was the following: $M_{STAMP}$: 1h, $M_{CR}$: 2h, $M_{K}$: 5h, $M_{BS}$: 6h, $M_{IR}$: 12h.

- When all models are done running and the outputs have been saved, the section III ("# Some graphs") contains code chunks in Python and R to analyse the models' error curve and extract all estimated parameters in .csv.


# Initial setup of the R scripts and preprocessing of the data documents (run this prior to any scripts in the document)

Retrieving documents necessary to all the notebook

```{r setup}
knitr::opts_chunk$set(echo= TRUE, tidy.opts = list(width.cutoff = 60), tidy="styler", warning= FALSE, message= FALSE)

suppressWarnings({

# R packages used in the R scripts
library(reticulate)
library(rstudioapi)
library(readxl)
library(tidyverse)
library(agricolae)
library(vegan)
library(dplyr)
library(rstatix)
library(ggsignif)
library(scales)
library(ggpubr)
library(see)
library(aplpack)
library(RColorBrewer)

# Polychrome version 1.5.1 install.packages("https://cran.r-project.org/src/contrib/Archive/Polychrome/Polychrome_1.5.1.tar.gz")
library(Polychrome)

# Retrieving the MicEco package
#library(devtools)
#install_github(repo="Russel88/MicEco")
library(MicEco)

# Python packages used in the Python scripts
py_require("numpy")
py_require("matplotlib")
py_require("os")
py_require("csv")
py_require("pickle")
  

# Setting working directory in the same folder as the notebook (not necessary in a R project)
# Server
#setwd(dirname(getActiveDocumentContext()$path))
# Local
#setwd(dirname(getSourceEditorContext()$path))

# Extracting the different tables corresponding to the different experiments

# Table of the experiment of plant infection in varying coupled conditions: temperature + genotype + roots condition
tab_tot = read_xlsx("Data/Data_tomato_2024.xlsx", range= "A1:LX193")

# Table of the experiment of plant infections collected over time
tab_kinetic = read_xlsx("Data/Data_tomato_2024.xlsx", range= "A195:HD237")

# Table of all inocula used for all infections
tab_inoc = read_xlsx("Data/Data_tomato_2024.xlsx", range= "A239:OD253")

# Table of the calibration of sequencing noise in the experiments
tab_noise_estimation = read_xlsx("Data/Data_tomato_2024.xlsx", range= "A263:ES266")

# Table containing the in vitro growth data for the 120 STAMP strains during inoculum preparation
tab_iv_growth = read_xlsx("Data/Data_120_growth_2024.xlsx", range= "A1:DQ72")


# Prefilter noise threshold: minimal number of reads considered as technical noise (index/barcode hopping)
# A value of 25 corresponds to a conservative threshold to eliminate 99% of barcodes hopping in the calibration test (see corresponding chunk)
Noise_threshold = 25


####################
# Preprocessing of the data in the experiment of infections under different conditions
####################

# Attributing a standard bottleneck Nb = 0 for samples in which CFU were detected but PCR for sequencing was not successful
# Nb and Nr are calculated using corresponding code chunk
tab_tot$`Filtered bottleneck Nb`[is.na(tab_tot$`Filtered bottleneck Nb`) & tab_tot$`Bacterial load (CFU/g)` > 0] = 0
tab_tot$`Filtered bottleneck Nb`[is.na(tab_tot$`Filtered bottleneck Nb`) & tab_tot$`Bacterial load (CFU/g)` == 0] = 0
tab_tot$`Corrected bottleneck Nr`[is.na(tab_tot$`Corrected bottleneck Nr`) & tab_tot$`Bacterial load (CFU/g)` > 0] = 0
tab_tot$`Corrected bottleneck Nr`[is.na(tab_tot$`Corrected bottleneck Nr`) & tab_tot$`Bacterial load (CFU/g)` == 0] = 0


# Rounding up standard bottlenecks Nb when the value is between 0 and 1
tab_tot$`Filtered bottleneck Nb`[tab_tot$`Filtered bottleneck Nb` > 0 & tab_tot$`Filtered bottleneck Nb` < 1] = 1


# Organising the factors in the table for graphical representations
tab_tot$Genotype = factor(tab_tot$Genotype, levels= c("M", "H"))

tab_tot = tab_tot %>%
  unite("Condition", Genotype, `Root condition`, sep= "_", remove= FALSE) %>%
  unite("Condition2", Condition, `Temperature (°C)`, sep= "_", remove= FALSE ) %>%
  unite("Condition3", Condition, Zone, sep= "_", remove= FALSE ) %>%
  unite("Condition4", Condition, `Temperature (°C)`, Zone, sep= "_", remove= FALSE ) %>%
  unite("Condition5", Genotype, Zone, sep= "_", remove= FALSE )

tab_tot$`Temperature (°C)` = factor(tab_tot$`Temperature (°C)`, levels = c(27, 32), labels = c("27°C", "32°C"))
tab_tot$Condition = factor(tab_tot$Condition, levels= c("M_Intact", "M_Cut", "H_Intact", "H_Cut"))

tab_tot$Condition2 = factor(tab_tot$Condition2, levels= c("M_Intact_27", "M_Cut_27", "H_Intact_27", "H_Cut_27", "M_Intact_32", "M_Cut_32", "H_Intact_32", "H_Cut_32"))

tab_tot$Condition3 = factor(tab_tot$Condition3, levels= c("M_Intact_Root", "M_Cut_Root", "H_Intact_Root", "H_Cut_Root"))

tab_tot$Condition4 = factor(tab_tot$Condition4, levels= c("M_Intact_27_Root", "M_Cut_27_Root", "H_Intact_27_Root", "H_Cut_27_Root", "M_Intact_32_Root", "M_Cut_32_Root", "H_Intact_32_Root", "H_Cut_32_Root"))

tab_tot$Condition5 = factor(tab_tot$Condition5, levels= c("M_Root", "H_Root"))


# Defining a logical of presence/absence of symptom at sampling
tab_tot$Symptoms = tab_tot$`Disease index`
tab_tot$Symptoms[tab_tot$Symptoms > 0] = 1
tab_tot$Symptoms[tab_tot$Symptoms == 0] = 0
tab_tot$Symptoms = as.logical(tab_tot$Symptoms)


# Defining a logical of validated/unvalidated sample for sequencing
tab_tot$Amplification = tab_tot$`PCR2 validation on gel`
tab_tot$Amplification[tab_tot$Amplification > 0] = 1
tab_tot$Amplification[tab_tot$Amplification == 0] = 0
tab_tot$Amplification[is.na(tab_tot$Amplification)] = 0
tab_tot$Amplification = as.logical(tab_tot$Amplification)


# Calculating Shannon diversity and richness for all samples
tab_temp = tab_tot

# Prefiltering using noise threshold
tab_temp[,56:175][tab_temp[,56:175] <= Noise_threshold] = 0

# Shannong diversity
tab_tot$`Shannon diversity` = diversity(tab_temp[,56:175])
tab_tot$`Shannon diversity`[tab_tot$Amplification == 0] = NA 

#Richness
tab_tot$Richness = rowSums(tab_temp[,56:175] > 0)
tab_tot$Richness[tab_tot$Amplification == 0] = NA 


####################
# Preprocessing of the data in the experiment of infections collected over time
####################

# Organising the factors in the experiments of infection collected over time
tab_kinetic$`Time post-inoculation (d)` = factor(tab_kinetic$`Time post-inoculation (d)`)

tab_kinetic$Zone = factor(tab_kinetic$Zone, levels = c("Root"))

tab_kinetic = tab_kinetic %>%
  unite("Condition", Genotype, `Root condition`, sep= "_", remove= FALSE) %>%
  unite("Condition2", Condition, `Temperature (°C)`, sep= "_", remove= FALSE ) %>%
  unite("Condition3", Condition, Zone, sep= "_", remove= FALSE ) %>%
  unite("Condition4", Condition, `Temperature (°C)`, Zone, sep= "_", remove= FALSE ) %>%
  unite("Condition5", Genotype, Zone, sep= "_", remove= FALSE )


# Setting a value of bacterial load in Marmande to 1 if no bacteria were detected
# Simplification assumption that an infection of susceptible plants is ~100% successful
tab_kinetic$`Bacterial load (CFU/g)`[tab_kinetic$`Bacterial load (CFU/g)` == 0] = 1


# Defining a logical of presence/absence of symptom at sampling
tab_kinetic$Symptoms = tab_kinetic$`Final Disease index`
tab_kinetic$Symptoms[tab_kinetic$Symptoms > 0] = 1
tab_kinetic$Symptoms[tab_kinetic$Symptoms == 0] = 0
tab_kinetic$Symptoms = as.logical(tab_kinetic$Symptoms)


# Defining a logical of validated/unvalidated sample for sequencing
tab_kinetic$Amplification = tab_kinetic$`PCR2 validation on gel`
tab_kinetic$Amplification[tab_kinetic$Amplification > 0] = 1
tab_kinetic$Amplification[tab_kinetic$Amplification == 0] = 0
tab_kinetic$Amplification[is.na(tab_kinetic$Amplification)] = 0
tab_kinetic$Amplification = as.logical(tab_kinetic$Amplification)


# Calculating Shannon diversity and richness for all samples
tab_temp = tab_kinetic

# Prefiltering using noise threshold
tab_temp[,56:175][tab_temp[,56:175] <= Noise_threshold] = 0

# Shannon diversity
tab_kinetic$`Shannon diversity` = diversity(tab_temp[,56:175])
tab_kinetic$`Shannon diversity`[tab_kinetic$Amplification == 0] = NA 

# Richness
tab_kinetic$Richness = rowSums(tab_temp[,56:175] > 0)
tab_kinetic$Richness[tab_kinetic$Amplification == 0] = NA 


# Setting a predetermined random seed for reproductibility
set.seed(123)


# Creating a colour palette of 120 discrete colourblind-friendly colours
P120 = createPalette(120,  c(brewer.pal(8, "Set1"),brewer.pal(8, "Dark2")))
P120 = unname(P120)

# Visualised colours
swatch(P120)


# Arranging tab_kinetic for the Python scripts
tab_tot$Validation = tab_tot$`PCR2 validation on gel`
tab_tot$Validation[tab_tot$Validation > 0] = 1
tab_tot$Validation[is.na(tab_tot$Validation)] = 0

# Export preprocessed tables for the Python scripts
write_csv(tab_tot,"Analysis_output/tab_tot.csv")

# Arranging tab_inoc for the Python scripts
tab_temp = tab_inoc %>%
  filter(!is.na(`Nanopore Run`)) %>%
  filter(`Inoculum ID 3` == 1)

write_csv(tab_temp,"Analysis_output/tab_inoc.csv")

# Arranging tab_kinetic for the Python scripts
tab_temp = tab_kinetic %>%
  filter(Zone == "Root") %>%
  filter(Condition2 == "M_Intact_27") %>%
  filter(`PCR2 validation on gel` > 0)

tab_temp$Validation = 1

write_csv(tab_temp,"Analysis_output/tab_kinetic.csv")

# Helper function for a non parametric test of Kruskal-Wallis with compact letter display with prearranging of the plant conditions
# p-value < 0.01 threshold for compact letter display

Graph_kruskal = function(
  
  # Table of interest
  tab,
  
  # Column name of the quantitative value (should be numeric)
  y,
  
  # Column name of the real groups to compare (should be a factor)
  x,
  
  # Skip empty factor H_Intact_27?
  skip_h = FALSE

) {
  
  tab = subset(tab, select= c(y, x))
  names(tab) = c("y", "Condition")
  
  # Performing test and retrieving the letters
  test = kruskal(tab$y, tab$Condition, group=TRUE, p.adj="bonferroni", alpha= 0.01)
  
  # Retrieving conditions and letters
  cld = as.data.frame(test$groups)
  cld = rownames_to_column(cld, "Condition")
  
  # Adding empty factor H_Intact_27 if not present
  if ((("H_Intact_27" %in% cld$Condition) == FALSE) & skip_h == FALSE) {
    cld = rbind(cld, c("H_Intact_27", NA, NA))
  }
  
  # Arranging the order of the letters by those of the condition in the table
  cld$Condition = factor(cld$Condition, levels= levels(tab$Condition))
  cld = arrange(cld, cld$Condition)
  
  return(cld)
  }

})

```

# Section R analysis (Nb, Nr, most data visualisation graphs)

# I. Upgraded STAMPR algorithm: Calculation of the bottleneck metrics Nr and FRD

## Description

The following code is an upgraded version of the algorithm proposed in the STAMPR method paper (Hullahalli et al. mSystems, 2021, code available on https://github.com/hullahalli/stampr_rtisan). The following modifications were performed:

- Correction of the the Nr value so it can pick the output of the "resiliency algorithm" (Nb_resilient) as one of the potential "best Nr" value. The previous code was not taking into account the "resiliency algorithm" even when it produced the "best Nr" according to the algorithm.
- Improvement of the annotation of the code with more explicit variable names and description of the algorithms.
- Providing access to some input parameters of the STAMPR algorithm.
- Inclusion of an analysis of the robustness of Nr with the calculation of the mathematical standard deviation on each Nr value. This analysis is based on the simulations performed in Abel et al, Nature Methods, 2016.
- Addition of a threshold in the resiliency algorithm when attaining low number of resilient tags. This threshold prevents cases of aberrantly high values of Nr that are calculated with low mathematical robustness based on a few tags.
- Improvement of the annotation of the code of the FRD calculation algorithm.

The outputs of the algorithm are the following:
1. Nb_unfiltered: Nb according to the original STAMP method without any noise filtering.
2. Nb_filtered: Nb according to the original STAMP method after prefiltered noise reads with the "Noise_threshold" parameter.
3. Nb_resilient: maximum value taken by Nb after prefiltering noise and applying of the "resiliency algorithm".
4. Ns: Nb value that is predicted when simulating multiple bottleneck events in the "reference resample plot".
5. Nr: Accepted value Nr which is the highest value between Nb_filtered, Nb_resilient and Ns.
6. Nr_STD: Mathematical error (standard deviation) on the value Nr under the assumption of a perfect STAMP model.

## Nr function

Calculation of a Nr value between a sample and its inoculum. Provide the following arguments:
- Sample index: name of the sample.
- Sample tags range: a vector of all STAMP tags read counts in sample.
- Inoculum tags range: a vector of the corresponding tags read counts in inoculum. Order and number of tags must correspond.
- Don't forget to set up the different algorithm parameters (see inside the function).

N.B: 0 reads tags in the inoculum should always be removed both in inoculum and samples as a tag that is not present in the inoculum is, in principle, non-existent. If you have tags that are confounded with noise in inoculum (i.e. under the noise threshold), I highly suggest to filter them out as they might produce bias in the final Nb and Nr calculated values.

```{r Upgraded Nr function}

# Definition of the function

Nr_calculate = function(
  # Sample name
  Sample_index,
  
  # STAMP tags reads counts in sample
  Sample_range,
  
  # STAMP tags reads counts in inoculum
  Inoc_range,
  
  # CFU reference for the maximum Nr threshold (default 10^8)
  CFU_reference = 10^8,
  
  # Minimal weight to remove non-noise threshold barcodes (default 2%)
  minweight = 0.02,
  
  # Noise threshold in number of reads per barcode (default 1)
  Noise_threshold = 1,
  
  # Minimum number of tags to ensure a non aberrant Nb_resilient value
  min_resilient_tags = 10,
  
  # Number of iterations for the local minima finder algorithm (default 10000)
  R_local_minima_finder = 10000,
  
  # Search range standard deviation for the local minima finder algorithm (default 1/10)
  # A low value will find more subpopulations
  STD_local_minima_finder = 1/10,
  
  # Plots creation
  plots = FALSE,
  
  # Number of iterations for the simulation of the error on Nb_filtered and Nr
  R_error_estimation = 1000
  
){
  
  ####################
  # 1. Calculation of the original STAMP bottleneck Nb with no filtering
  ####################
  
  # Original Nb formula
  fi0 = na.omit(Inoc_range/sum(Inoc_range))
  fis = na.omit(Sample_range / sum(Sample_range))
  sigma <- ((fis - fi0)^2) / (fi0*(1-fi0))
  sigma <- sigma[sigma!=Inf]
  F <- mean(na.omit(sigma))
  Nb_unfiltered <- 1 / (F - 1/sum(Inoc_range) - 1/sum(Sample_range))
  
  
  ####################
  # 2. Calculation of the original STAMP bottleneck Nb after prefiltering step
  ####################
  
  # Calculation of signal-to-noise (SNR) correction by considering a read threshold for noise
  one_ratio = sqrt(sum(Sample_range > Noise_threshold) / sum(Sample_range >= 1 & Sample_range <= Noise_threshold))
  
  if(one_ratio < 1) {
    one_ratio = -(one_ratio ^2)
    }
  
  Noise_correction_factor = 10 ^ (-one_ratio)
  Noise_distance = colSums(rbind(Sample_range, Inoc_range))
  
  # Producing random simulation based on the observed distributions of tags in inoculum and sample
  Resampled_inoc = rmultinom(50, round(sum(Sample_range) * Noise_correction_factor), Noise_distance)
  Resampled_inoc = t(Resampled_inoc)
  
  # Calculating a resampled table including dispersion between measured tag distribution and simulated tags
  # This parameter can be corrected by taking into account a measure of CFU load
  tab_resampled = ceiling(colMeans(Sample_range - Resampled_inoc))
  tab_resampled[tab_resampled < 0] = 0
  
  if (sum(tab_resampled > 1) >  (2 * sum(tab_resampled == 1))) {
    
    times = min(ceiling(CFU_reference), sum(tab_resampled > 0))
    } else {
      
      times = min(ceiling(CFU_reference), sum(tab_resampled > 1))
      }
  
  # Filtered Nb calculation as per orginal Nb formula
  fi0 = na.omit(Inoc_range/sum(Inoc_range))
  fis = na.omit(tab_resampled / sum(tab_resampled))
  sigma <- ((fis - fi0)^2) / (fi0*(1-fi0))
  sigma <- sigma[sigma!=Inf]
  F <- mean(na.omit(sigma))
  Nb_filtered <- 1 / (F - 1/sum(Inoc_range) - 1/sum(tab_resampled)) 

  
  # Checkpoint: stop everything if Nb is negative
  if (Nb_filtered <= 0) {
    
    output = data.frame(rbind(c(Sample_index, c(sum(tab_resampled), Nb_unfiltered, NA, NA, NA, NA, NA, NA, NA))))
    names(output) = c("Sample ID", "TotalReads", "Nb_unfiltered", "Nb_filtered", "Nb_resilient", "Ns", "Nr", "Number of barcodes", "Standard deviation on Nb_filtered", "Standard deviation on Nr")
    
    return(output)
  }
  
  
  ####################
  # 3. Calculation of the resiliency algorithm with noise prefiltering and extraction of the maximum Nb
  ####################
  
  # Ordering distribution of tags in inoculum versus in silico resampled inoculum
  tab_dif = t(rbind(Inoc_range, tab_resampled))
  tab_dif = tab_dif[order(tab_dif[,2]),]
  colnames(tab_dif) = c("Inoculum", "Samples")
    
  # Calculating bottleneck Nb resiliency range
  # Sequential rarefaction of tags from highest to lowest represented tags
  Tags_range = rev(seq(from = length(Inoc_range) - times + 1, to = length(Inoc_range), by = 1))
  
  Nb_resiliency_range = c()
    
  for (k in c(1:length(Tags_range))) {
    
    # Calculation of a range of Nb as per original Nb formula
    fi0 = na.omit(tab_dif[,1][1:Tags_range[k]] / sum(tab_dif[,1][1:Tags_range[k]]))
    fis = na.omit(tab_dif[,2][1:Tags_range[k]] / sum(tab_dif[,2][1:Tags_range[k]]))
    sigma = ( fis - fi0 ) ^ 2 / (fi0*(1-fi0))
    sigma = sigma[sigma!=Inf]
    F = mean(na.omit(sigma))
    Nb_resiliency_range[k] = 1 / (F - 1/sum(Inoc_range) - 1/sum(tab_resampled))
    }
  
  tags_at_0 = length(Inoc_range) - length(Nb_resiliency_range)
    
  # Scanning the resiliency range for local minima (breakpoints in the curves)
  # Arbitrary number of starting initiation sites set with p
  # Arbitrary search range value with STD_local_minima_finder
  p = Nb_resiliency_range[seq(1, length(na.omit(Nb_resiliency_range)), length.out = log(length(na.omit(Nb_resiliency_range))))]
    
  if (length(p) == 0) {
    p = Nb_resiliency_range[1]
  }
    
  breakpoints = c()
    
  for (k in c(1:length(p))) {
    start = p[k]
    
    for (l in c(1:R_local_minima_finder)) {
      
      where = which(Nb_resiliency_range == start)
      newlocation = abs(rnorm(1, mean = where, sd = sum(tab_resampled > 1) * STD_local_minima_finder))
    
      if(newlocation > length(na.omit(Nb_resiliency_range))) {
        newlocation = where
        }
    
      if(newlocation < 1) {
        newlocation = where
        }
    
      if(Nb_resiliency_range[round(newlocation)] < start) {
        
        start = Nb_resiliency_range[round(newlocation)]
        }
      }
    
    breakpoints[k] = which(Nb_resiliency_range == start)
  }

  if(length(breakpoints) == 0) {
    breakpoints = 1
  }
  
  
  # Sort to keep unique breakpoints and combine breakpoints next to each other
  sorted_breakpoints = sort(unique(c(breakpoints, length(na.omit(Nb_resiliency_range)))))
  sorted_breakpoints = sorted_breakpoints[sorted_breakpoints != 0]

  a = c(sorted_breakpoints, 10000000) - c(-10000000, sorted_breakpoints)
    
  if (min(a) == 1) {
    sorted_breakpoints = sorted_breakpoints[-(max(which(a == 1)) - 1)]
    }

  sorted_breakpoints[sorted_breakpoints > (max(sorted_breakpoints) - 5)] = max(sorted_breakpoints)
  sorted_breakpoints = unique(sorted_breakpoints)
    
  # Calculation of the fractions of reads contained between each breakpoint ("weights")
  sorted_breakpoints_freq = c()
    
  for (k in c(1:length(sorted_breakpoints))) {
      
    sorted_breakpoints_freq[k] = sum(tail(sort(tab_resampled), sorted_breakpoints[k])) / sum(tab_resampled)
    }

  weights = sorted_breakpoints_freq - c(0, sorted_breakpoints_freq)[1:length(sorted_breakpoints_freq)]
  indices = data.frame(sorted_breakpoints, weights)
  colnames(indices) = c("Number of barcodes", "Fraction of reads")
    
    
  # Calculation of the greatest log change between weights
  # Tags before lowest breakpoint will be attributed to noise
  log_weights = log(weights)
  weightsdif = (log_weights - c(log_weights[2:length(indices[,1])], 0))[1:length(log_weights)-1]

  if(length(weightsdif) == 0) {
    weightsdif = indices[,1]
    }

  noisestart = indices[,1][which(weightsdif == max(weightsdif))]
    
    
  # If there are no tags below the lowest breakpoint
  # Set noise to be below the next breakpoint
  if (is.na(indices[,1][min(which(indices[,2] < minweight))])) {
    noisestart = max(indices[,1])
    }
    
  
  # If the sum of the predicted non noise weights exceeds the limit defined from calibration
  # Set noise to all tags under the defined minimum threshold
  minweightcutoff <- min(which(cumsum(indices[,2])>(1-minweight)))

  if (sum(indices[,2][which(indices[,1] > noisestart)]) > minweight) {
    noisestart = indices[,1][minweightcutoff]
    }
  
  
  ####################
  # 4. Generation of the reference resample plot and computation of the simulated Ns
  #################### 
  
  FirstResample = as.numeric(rmultinom(1, sum(tab_resampled), Inoc_range/sum(Inoc_range)))
  steps1 = round(seq(from = 1, to = length(FirstResample), length.out = 100))
  steps2 = round(seq(from = length(FirstResample), to = length(FirstResample)*20, length.out = 100))
  steps = c(steps1, steps2)
  y = c()
    
  for (k in c(1:length(steps))) {
      
    vec = rmultinom(5, steps[k], FirstResample/sum(FirstResample))
    vec[vec!=0] = 1
    y[k] = mean(colSums(vec))
    }
    
    
  # Interpolation of the non noise part of the reference resample plot
  # Extraction of Ns
  interpol <- approx(x = steps, y = y, n = length(Inoc_range))
  xvals <- as.numeric(unlist(interpol[1]))
  yvals <- as.numeric(unlist(interpol[2]))
  dfxy2 <- sortedXyData(x = xvals, y = yvals)
  Ns <- NLSstClosestX(dfxy2, noisestart)
  
  ####################  
  # 5. Graphs and selection of the Nr
  ####################
    
  # Sample per sample plots of breakpoints
  
  if(plots == TRUE){
    
    converttocutoffs<- function(p) {
      cutoffposition <- length(tab_resampled) - p
      cutoffvalue <- tab_dif[cutoffposition+1,2]
      
      if(cutoffvalue == 0) {cutoffvalue <- 1}    
      cutoffvalue/sum(tab_resampled) 
      }
    
    cutoffpositions <- unlist(lapply(sorted_breakpoints, converttocutoffs))
    cutoffpositions <<- cutoffpositions
    par(mfrow = c(2,1))
  
    plot(1:times,Nb_resiliency_range, log = "y", ylim = c(.01, 2E6),
         xlim = c(1,length(as.numeric(na.omit(Nb_resiliency_range)))),
         main = "Resiliency", ylab = "Nb", xlab = "Iteration")
    abline(v= sorted_breakpoints)
    abline(v= (length(Nb_resiliency_range) - min_resilient_tags), col= "red")
      
    plot(tab_resampled/sum(tab_resampled), log = "y",
         ylim = c(1E-8, 1),
         main = "Barcodes", ylab = "Frequency", xlab = "Barcode")
    abline(h = cutoffpositions)
    }
  
  # Selecting the Nr
  # Retrieving the effective number of used barcodes in the Nr value for error estimation
  tags_threshold = min_resilient_tags - tags_at_0
    
  if (tags_threshold < 0) {
    tags_threshold = 0
  }
    
  if (length(Nb_resiliency_range) > tags_threshold) {
    Nb_resilient = max(Nb_resiliency_range[1:(length(Nb_resiliency_range) - tags_threshold)], na.rm= TRUE)
  } else {
    Nb_resilient = max(Nb_resiliency_range, na.rm= TRUE)
  }
    
  Nr = max(Nb_filtered, Nb_resilient, Ns)
  
  if (Nr == Nb_resilient) {
    Effective_tags = length(Inoc_range) - match(Nb_resilient, Nb_resiliency_range) + 1
  }
    
  if (Nr == max(Nb_filtered, Ns)) {
    Effective_tags = length(Inoc_range[Inoc_range > Noise_threshold])
  }
  
  
  ####################
  # 6. Calculation of the error on the Nb and Nr value through simulations
  ####################
  
  # Using the total number of used tags and Nb_filtered to resimulate a bottleneck under a perfect STAMP model case
  tab_simulation = rmultinom(R_error_estimation, round(Nb_filtered), rep(1/length(Inoc_range),length(Inoc_range)))
  
  # Calculating resimulated values "Ne" of Nb_filtered
  fis = tab_simulation / round(Nb_filtered)
  sigma = ((fis - 1/length(Inoc_range))^2) / (1/length(Inoc_range)*(1-1/length(Inoc_range)))
  Ne = 1 / colMeans(sigma)
  Ne[Ne == Inf] = NA
  
  Nb_filtered_STD = sd(Ne, na.rm= TRUE)
  
  
  # Same protocol for Nr
  # Using the effective number of tags and Nr to resimulate a bottleneck under a perfect STAMP model case
  tab_simulation = rmultinom(R_error_estimation, round(Nr), rep(1/Effective_tags,Effective_tags))
  
  # Calculating resimulated values "Ne" of Nr
  fis = tab_simulation / round(Nr)
  sigma = ((fis - 1/Effective_tags)^2) / (1/Effective_tags*(1-1/Effective_tags))
  Ne = 1 / colMeans(sigma)
  Ne[Ne == Inf] = NA
  
  Nr_STD = sd(Ne, na.rm= TRUE)
  

  ####################
  
  # Returning all values  
  output = data.frame(rbind(c(Sample_index, c(sum(tab_resampled), Nb_unfiltered, Nb_filtered, Nb_resilient, Ns, Nr, Effective_tags, Nb_filtered_STD, Nr_STD))))
  names(output) = c("Sample ID", "TotalReads", "Nb_unfiltered", "Nb_filtered", "Nb_resilient", "Ns", "Nr", "Number of barcodes", "Standard deviation on Nb_filtered", "Standard deviation on Nr")
  
  return(output)
  }

```

## FRD function

For two paired samples, calculation of a FRD to estimate their relatedness. Provide the following arguments:
- Samples tags ranges: two vectors of STAMP tags read counts of identical dimension to be compared.

```{r Upgraded FRD function}

# Cavalli-Sforza chord distance function
getGD <- function(f1, f2) {
    f1 <- f1/sum(f1)
    f2 <- f2/sum(f2)
    cosTheta = sum(sqrt(f1*f2))
    chorddistance = 2*sqrt(2)/pi*sqrt(1-cosTheta)
    chorddistance
    }

# FRD function
FRD_calculate = function(
  
  # Name of the couple of sample
  Couple_index,
  
  # First sample
  vec1,
  
  # Second sample
  vec2,
  
  # Plot creation
  plots = FALSE
) {
  
  tab_dif = as.data.frame(t(rbind(vec1, vec2, sqrt((vec1/sum(vec1))*(vec2/sum(vec2))))))
  
  #names(tab_dif) = c("Sample 1", "Sample 2", "Co-occurence" )  
  
  tab_dif_sorted = tab_dif[order(tab_dif[,3], decreasing= TRUE),]
  
  
  # Calculation of resilient genetic distance RD
  # Removal of the less co-abundant tags and calculation of the GD
  tab_temp = tab_dif_sorted
  GD_resiliency = c()

  for (j in c(1:min(sum(vec1>0), sum(vec2>0)))) {
    GD_temp = getGD(tab_temp[,1], tab_temp[,2])
    GD_resiliency = c(GD_resiliency, GD_temp)
    tab_temp = tab_temp[1:dim(tab_temp)[1]-1,] 
    }
  
  # Summary of statistics
  GD = GD_resiliency[1]
  GD_resiliency = GD_resiliency[2:length(GD_resiliency)]
  RD = sum(na.omit(GD_resiliency) < 0.8)
  FRD1to2 = log(RD+1)/log(sum(vec2!=0)+1)
  FRD2to1 = log(RD+1)/log(sum(vec1!=0+1))
  
  # Plotting and displaying statistics
  if (plots == TRUE) {
    par(mfrow = c(1,1))
    plot(GD_resiliency, xlab = "Iteration", ylab = "Genetic Distance", ylim = c(0,1))
    print(paste("RD =", RD))
    print(paste("GD =", GD))
    print(paste("MinGD=", min(na.omit(GD_resiliency))))
    print(paste("Sample1 to Sample2 =", FRD1to2))
    print(paste("Sample2 to Sample1 =", FRD2to1))
    }
  
  # Retrieving values
  output = data.frame(rbind(c(Couple_index, GD, RD, FRD1to2, FRD2to1)))
  names(output) = c("Couple ID", "Genetic distance GD", " Resilient distance RD", "FRD Sample1 to Sample2", "FRD Sample2 to Sample1")
  
  return(output)
}

```

## Use of the Nr algorithm to analyse plant samples

Application of Nr algorithm to the experiment of the infections in different conditions (tab_tot).

```{r Nr function application 1}

# Setting random seed
set.seed(123)

# Defining table to collect results
tab_final = data.frame(matrix(nrow=0, ncol=10))

# Scanning all samples
for (i in tab_tot$`Plant ID`) {

  # Verifying if sample is validated, skipping to the next if invalid
  if (tab_tot$Amplification[i] == FALSE) {
    tab_final = rbind(tab_final, c(i,NA,NA,NA,NA,NA,NA,NA,NA,NA))
    
    next
  }
  
  # Extracting STAMP tags range from sample
  tab_tags = tab_tot %>%
    filter(`Plant ID` == i)
  tab_tags = tab_tags[,56:175]
  tab_tags = data.frame(tab_tags)
  
  # Extracting tags from corresponding inoculum
  tab_tags_inoc = tab_inoc %>%
    filter(`Inoculum ID 2` == tab_tot$`Inoculum ID`[tab_tot$`Plant ID` == i])
  tab_tags_inoc = tab_tags_inoc[,147:266]
  names(tab_tags_inoc) = names(tab_tags)
    
  # Curate non-analysable tags that are under the noise prefiltering threshold value in inoculum
  tab_tags = tab_tags[colSums(tab_tags_inoc) > Noise_threshold]
  tab_tags_inoc = tab_tags_inoc[colSums(tab_tags_inoc) > Noise_threshold]
  
  # Renaming tab_final to match Nr_calculate because rbind messes it at each iteration
  names(tab_final) = c("Sample ID", "TotalReads", "Nb_unfiltered", "Nb_filtered", "Nb_resilient", "Ns", "Nr", "Number of barcodes", "Standard deviation on Nb_filtered", "Standard deviation on Nr")
  
  # Using the Nr function and collection values
  # Setting minweight to 0.001 because reads are already prefiltered
  tab_final = rbind(tab_final,
                    Nr_calculate(Sample_index= i, Sample_range= tab_tags, Inoc_range= tab_tags_inoc, Noise_threshold= Noise_threshold, minweight= 0.001))
  }
    
# Final names labelling
names(tab_final) = c("Plant ID", "TotalReads", "Unfiltered bottleneck Nb", "Filtered bottleneck Nb", "Resilient bottleneck Nb", "Simulated bottleneck Ns", "Corrected bottleneck Nr", "Effective number of tags in Nr", "Standard deviation on filtered Nb", "Standard deviation on Nr")

# Results can be added to the dataset
write_csv2(tab_final, "Analysis_output/Nr_calculation_tab_tot.csv")

```

Application of Nr algorithm to the infections with sampling over time (tab_kinetic).

```{r Nr function application 2}

# Setting random seed
set.seed(123)

# Defining table to collect results
tab_final = data.frame(matrix(nrow=0, ncol=10))

# Scanning all samples
for (i in tab_kinetic$`Plant ID`) {

  # Verifying if sample is validated, skipping to the next if invalid
  if (tab_kinetic$Amplification[i] == FALSE) {
    tab_final = rbind(tab_final, c(i,NA,NA,NA,NA,NA,NA,NA,NA,NA))
    
    next
  }
  
  # Extracting STAMP tags range from sample
  tab_tags = tab_kinetic %>%
    filter(`Plant ID` == i)
  tab_tags = tab_tags[,56:175]
  tab_tags = data.frame(tab_tags)
  
  # Extracting tags from corresponding inoculum
  tab_tags_inoc = tab_inoc %>%
    filter(`Inoculum ID 2` == tab_kinetic$`Inoculum ID`[tab_kinetic$`Plant ID` == i])
  tab_tags_inoc = tab_tags_inoc[,147:266]
  names(tab_tags_inoc) = names(tab_tags)
    
  # Curate non-analysable tags that are under the noise prefiltering threshold value in inoculum
  tab_tags = tab_tags[colSums(tab_tags_inoc) > Noise_threshold]
  tab_tags_inoc = tab_tags_inoc[colSums(tab_tags_inoc) > Noise_threshold]
  
  # Renaming tab_final to match Nr_calculate because rbind messes it at each iteration
  names(tab_final) = c("Sample ID", "TotalReads", "Nb_unfiltered", "Nb_filtered", "Nb_resilient", "Ns", "Nr", "Number of barcodes", "Standard deviation on Nb_filtered", "Standard deviation on Nr")
  
  # Using the Nr function and collection values
  # Setting minweight to 0.001 because reads are already prefiltered
  tab_final = rbind(tab_final,
                    Nr_calculate(Sample_index= i, Sample_range= tab_tags, Inoc_range= tab_tags_inoc, Noise_threshold= Noise_threshold, minweight= 0.001))
  
  }
    
# Final names labelling
names(tab_final) = c("Plant ID", "TotalReads", "Unfiltered bottleneck Nb", "Filtered bottleneck Nb", "Resilient bottleneck Nb", "Simulated bottleneck Ns", "Corrected bottleneck Nr", "Effective number of tags in Nr", "Standard deviation on filtered Nb", "Standard deviation on Nr")

# Results can be added to the dataset
write_csv2(tab_final, "Analysis_output/Nr_calculation_tab_kinetic.csv")

```

# II. (unused)

# III. Initial analysis of the R. pseudosolanacearum infection dataset using Nb and Nr

## Description of the experiment of infection in varying conditions (tab_tot)

### Visualisation of bacteria load

Bacterial load data between all conditions

```{r Graph}

####################
# Bacterial load in root
####################

# Retrieving table and filtering data of interest
tab = tab_tot %>%
  filter(Zone == "Root")

# Kruskal-wallis statistical test
test = Graph_kruskal(tab= tab, y= "Bacterial load (CFU/g)", x= "Condition2")

# Display helper
values = tab %>% 
  group_by(Condition) %>% 
  summarise(mean = mean("Bacterial load (CFU/g)"), sd = sd("Bacterial load (CFU/g)"))

# Graph v1
ggplot(data= tab, aes(x= Condition, y= `Bacterial load (CFU/g)`, fill= Genotype)) +
  
  geom_boxplot(outlier.shape= NA, width=0.9) +
  geom_point(aes(fill= Genotype, colour= Replicate, shape= Symptoms),
             position= position_jitterdodge(jitter.width= 0.1, dodge.width= 1), alpha= 1) +
  geom_text(data = values,  y = 1, aes(label = test$groups , x = Condition), vjust = -30, inherit.aes = FALSE) +
  
  facet_wrap(~ `Temperature (°C)`, scales= "free_x", strip.position="bottom") +
  
  scale_fill_manual(values= c("#99CCFF", "#FFCC99")) +
  scale_colour_manual(values= c("#E41A1C", "#339900", "#FFD92F")) +
  scale_y_continuous(trans = log10_trans(), limits=c(50, 2*10^9)) +
  scale_shape_manual(values= c(1,19), labels= c("No", "Yes")) +
  labs(title= "Root", y= "Bacterial load in root (CFU/g)", shape= "Symptoms") +
  theme_gray()

# Graph v2
ggplot(data= tab, aes(x= Condition, y= `Bacterial load (CFU/g)`, fill= Genotype)) +
  
  geom_violin(data= tab, width= 1, position= position_nudge(x= -0.1, y= 0)) +
  stat_boxplot(aes(ymin = after_stat(lower), ymax = after_stat(upper)),
               geom="errorbar", width=.25, position= position_nudge(x= -0.1, y= 0)) + 
  stat_boxplot(aes(ymin = after_stat(middle), ymax = after_stat(middle)),
               geom="errorbar", linewidth=1, width=.25, position= position_nudge(x= -0.1, y= 0)) +
  geom_point(aes(fill= Genotype, colour= Replicate, shape= Symptoms),
             position= position_nudge(x= 0.2, y= 0), alpha= 1) +
  geom_text(data = values,  y = 1, aes(label = test$groups , x = Condition), vjust = -30, inherit.aes = FALSE) +
  
  facet_wrap(~ `Temperature (°C)`, scales= "free_x", strip.position="bottom") +
  
  scale_fill_manual(values= c("#99CCFF", "#FFCC99")) +
  scale_colour_manual(values= c("#E41A1C", "#339900", "#FFD92F")) +
  scale_y_continuous(trans = log10_trans(), limits=c(50, 2*10^9)) +
  scale_shape_manual(values= c(1,19), labels= c("No", "Yes")) +
  labs(title= "Root", y= "Bacterial load in root (CFU/g)", shape= "Symptoms") +
  theme_gray()


####################
# Comparison in load between M_Intact_27 and M_Cut_27
####################

tab = tab_tot %>%
  filter(Condition2 %in% c("M_Intact_27", "M_Cut_27"))

tab$`Root condition` = factor(tab$`Root condition`, levels= c("Intact", "Cut"))

# Graph 2 v3: Boxplots of bacterial load in Marmande plants at 27°C

ggplot(data= tab, aes(x= Zone, y= `Bacterial load (CFU/g)`)) +
  
  geom_boxplot(aes(fill = `Root condition`), outlier.shape= NA, width=0.9, show.legend= FALSE) +
  geom_point(aes(colour= Replicate, shape= Symptoms),
             position= position_jitterdodge(jitter.width= 0.1, dodge.width= 1), alpha= 1) +
  
  facet_wrap(~ Condition, scales= "free_x", strip.position="bottom") +
  scale_fill_manual(values = c("#99CCFF", "#6699FF")) +
  scale_colour_manual(values= c("#E41A1C", "#339900", "#FFD92F")) +
  scale_y_continuous(trans = log10_trans(), limits=c(50, 2*10^9)) +
  scale_shape_manual(values= c(1,19), labels= c("No", "Yes")) +
  labs(y= "Bacterial load in root (CFU/g)", shape= "Symptoms") +
  theme_gray() +
  theme(strip.background= element_blank(), strip.placement= "outside")

```

### Visualisation of Nb or Nr

Filtered bottleneck Nb in all conditions

```{r Graph}

####################
# Nb in root
####################

# Retrieving table and filtering data of interest
tab = tab_tot %>%
  filter(Zone == "Root")

# Kruskal-wallis statistical test
test = Graph_kruskal(tab= tab, y= "Filtered bottleneck Nb", x= "Condition2")

# Display helper
values = tab %>% 
  group_by(Condition) %>% 
  summarise(mean = mean("Filtered bottleneck Nb"), sd = sd("Filtered bottleneck Nb")) 


# Graph v1
ggplot(data= tab, aes(x= Condition, y= `Filtered bottleneck Nb`, fill= Genotype)) +
  
  geom_boxplot(outlier.shape= NA, width=0.9) +
  geom_pointrange(aes(fill= Genotype, colour= Replicate,
                    ymax= (`Filtered bottleneck Nb` + `Nb mathematical standard deviation STD_Nb`), ymin= (`Filtered bottleneck Nb` - `Nb mathematical standard deviation STD_Nb`)),
                  position= position_jitterdodge(jitter.width= 0.3, dodge.width= 1), size= 0.2, alpha= 1) +
  geom_text(data = values, aes(label = test$groups, y = 1 , x = Condition), vjust = -27.5, inherit.aes = FALSE) +

  facet_wrap(~ `Temperature (°C)`, scales= "free_x", strip.position="bottom") +
  scale_fill_manual(values= c("#99CCFF", "#FFCC99")) +
  scale_colour_manual(values= c("#E41A1C", "#339900", "#FFD92F")) +
  scale_y_continuous(limits= c(0, NA)) +
  coord_cartesian(ylim = c(0,175)) +
  labs(title= "Root", y = "Root entry bottleneck Nb (in number of bacteria)") +
  theme_gray()


# Graph v2
ggplot(data= tab, aes(x= Condition, y= `Filtered bottleneck Nb`, fill= Genotype)) +
  
  geom_boxplot(outlier.shape= NA, width=0.9) +
  geom_point(aes(fill= Genotype, colour= Replicate, shape= Amplification),
             position= position_jitterdodge(jitter.width= 0.1, dodge.width= 1), alpha= 1) +
  geom_text(data = values, aes(label = test$groups, y = 1 , x = Condition), vjust = -27.5, inherit.aes = FALSE) +
  
  facet_wrap(~ `Temperature (°C)`, scales= "free_x", strip.position="bottom") +
  scale_fill_manual(values= c("#99CCFF", "#FFCC99")) +
  scale_colour_manual(values= c("#E41A1C", "#339900", "#FFD92F")) +
  scale_y_log10(limits = c(1, 25000)) +
  scale_shape_manual(values= c(4,19)) +
  labs(title= "Root", y = "Root entry bottleneck Nb (in number of bacteria)") +
  theme_gray()


####################
# Nb in all samples
####################

tab = tab_tot

# Graph v1
ggplot(data= tab, aes(x= Condition3, y= `Filtered bottleneck Nb`, fill= Condition5)) +
  
  geom_boxplot(outlier.shape= NA, width=0.9) +
  geom_point(aes(colour= Replicate, shape= Amplification), alpha= 1) +
  geom_line(aes(group= `Plant ID`, colour= Replicate), alpha= 1, size= 0.3, linetype= 1) + 
  
  facet_wrap(~ `Temperature (°C)`, scales= "free_x", strip.position="bottom") +
  scale_fill_manual(values = c("#99CCFF", "#6699FF", "#FFCC99", "#FF9900")) +
  scale_colour_manual(values= c("#E41A1C", "#339900", "#FFD92F")) +
  scale_x_discrete(guide = guide_axis(angle = 90)) +
  scale_y_continuous(trans = log10_trans(), limits=c(1, 25000)) +
  scale_shape_manual(values= c(4,16)) +
  labs(y = "Bottleneck Nb between inoculum and compartment (in number of bacteria)") +
  theme_gray()

```

Corrected bottleneck Nr in all conditions

```{r Graph}

####################
# Nr in root
####################

tab = tab_tot %>%
  filter(Zone == "Root")

# Adding error bars to the Nr value
tab$`Nr Errorbar max` = tab$`Corrected bottleneck Nr` + tab$`Nr mathematical standard deviation STD_Nr`
tab$`Nr Errorbar min` = tab$`Corrected bottleneck Nr` - tab$`Nr mathematical standard deviation STD_Nr`
tab$`Nr Errorbar min`[tab$`Nr Errorbar min` < 1] = 1

# Kruskal-wallis statistical test
test = Graph_kruskal(tab= tab, y= "Bacterial load (CFU/g)", x= "Condition2")

# Display helper
values = tab %>% 
  group_by(Condition) %>% 
  summarise(mean = mean(`Corrected bottleneck Nr`), sd = sd(`Corrected bottleneck Nr`)) 


# Graph v1
ggplot(data= tab, aes(x= Condition, y= `Corrected bottleneck Nr`, fill= Genotype)) +
  
  geom_boxplot(outlier.shape= NA, width=0.9) +
  geom_point(aes(fill= Genotype, colour= Replicate, shape= Amplification),
             position= position_jitterdodge(jitter.width= 0.1, dodge.width= 1), alpha= 1) +
  geom_text(data = values, aes(label = test$groups, y = 1 , x = Condition), vjust = -27.5, inherit.aes = FALSE) +
  
  facet_wrap(~ `Temperature (°C)`, scales= "free_x", strip.position="bottom") +
  scale_fill_manual(values = c("#99CCFF", "#FFCC99")) +
  scale_colour_manual(values= c("#E41A1C", "#339900", "#FFD92F")) +
  scale_y_log10(limits = c(1, 25000)) +
  scale_shape_manual(values= c(4,19), name= "Sample Statut", labels= c("No data","Sequenced")) +
  labs(title= "Root", y = "Corrected bottleneck Nr in root (number of bacteria)") +
  theme_gray()


# Graph v2
ggplot(data= tab, aes(x= Condition, y= `Corrected bottleneck Nr`, fill= Genotype)) +
  
  geom_boxplot(outlier.shape= NA, width=0.9) +
  geom_pointrange(aes(fill= Genotype, colour= Replicate, shape= Amplification,
                      ymax= `Nr Errorbar max`,
                      ymin= `Nr Errorbar min`),
                  position= position_jitterdodge(jitter.width= 0.7, dodge.width= 1), size= 0.2, alpha= 1) +
  geom_text(data = values, aes(label = test$groups, y = 1 , x = Condition), vjust = -27.5, inherit.aes = FALSE) +
  
  facet_wrap(~ `Temperature (°C)`, scales= "free_x", strip.position="bottom") +
  scale_fill_manual(values = c("#99CCFF", "#FFCC99")) +
  scale_colour_manual(values= c("#E41A1C", "#339900", "#FFD92F")) +
  scale_y_log10(limits = c(1, 25000)) +
  scale_shape_manual(values= c(4,19), name= "Sample Statut", labels= c("No data","Sequenced")) +
  labs(title= "Root", y = "Corrected bottleneck Nr in root (number of bacteria)") +
  theme_gray()

```

Comparisons of Nb, Nr and Shannon diversity

```{r Graph}

####################
# Comparison Nb vs Nr in root
####################

tab = tab_tot %>%
  filter(Zone == "Root")

# Joining together Nb and Nr values to compare them
tab = pivot_longer(tab, c(`Filtered bottleneck Nb`,`Corrected bottleneck Nr`))

tab$name[tab$name == "Filtered bottleneck Nb"] = "Nb"
tab$name[tab$name == "Corrected bottleneck Nr"] = "Nr"

tab = tab %>%
  unite("Condition3", Condition, name, sep= "_", remove= FALSE)

tab$Condition3 = factor(tab$Condition3, levels= c("M_Intact_Nb", "M_Intact_Nr", "M_Cut_Nb", "M_Cut_Nr", "H_Intact_Nb", "H_Intact_Nr", "H_Cut_Nb", "H_Cut_Nr"))


# Producing dummy data for the empty condition H_Intact_27
tab_temp = tab
tab_temp$value[tab_temp$Condition2 == "H_Intact_27"] = sample(48)

# Non-parametric test of Wilcoxon-Mann-Whitney
test = tab_temp %>%
  group_by(`Temperature (°C)`) %>%
  wilcox_test(formula = value ~ Condition3,
                      comparisons= list(c("M_Intact_Nb", "M_Intact_Nr"), c("M_Cut_Nb", "M_Cut_Nr"),
                                        c("H_Intact_Nb", "H_Intact_Nr"), c("H_Cut_Nb", "H_Cut_Nr")),
                      paired= FALSE, p.adjust.method = "none") %>%
  add_xy_position(x= "Condition3", fun= "max")

# Organising conditions
test$p.adj[3] = NA
test$p.adj.signif[3] = ""
test$Genotype = c("M", "M", "H", "H", "M", "M", "H", "H")
test$y.position = log10(test$y.position)


# Graph v1
ggplot(data= tab, aes(x= Condition3, y= value, fill= Genotype)) +
  
  geom_boxplot(outlier.shape= NA, width=0.9) +
  geom_point(aes(colour= Replicate, shape= Amplification), alpha= 1) +
  #geom_line(aes(group= `Plant ID`, colour= Replicate), alpha= 1, size= 0.3, linetype= 1) + 
  stat_pvalue_manual(test, y.position= 4.6) +
  
  facet_wrap(~ `Temperature (°C)`, scales= "free_x", strip.position="bottom") +
  scale_fill_manual(values = c("#99CCFF", "#FFCC99")) +
  scale_colour_manual(values= c("#E41A1C", "#339900", "#FFD92F")) +
  scale_x_discrete(guide = guide_axis(angle = 90)) +
  scale_y_continuous(trans = log10_trans(), limits=c(1, 40000)) +
  scale_shape_manual(values= c(4,16)) +
  labs(title= "Root", x= "Condition", y = "Nb or Nr in root (number of bacteria)") +
  theme_gray()


# Selecting only a comparison of M_Intact_27 and M_Cut_27
tab = tab %>%
  filter(Genotype == "M") %>%
  filter(`Temperature (°C)` == "27°C")

test = test %>%
  filter(Genotype == "M") %>%
  filter(`Temperature (°C)` == "27°C")

tab$`Root condition` = factor(tab$`Root condition`, levels= c("Intact", "Cut"))
test$`Root condition` = factor(c("Intact", "Cut"), levels= c("Intact", "Cut"))

# Graph v2
ggplot(data= tab, aes(x= name, y= value)) +
  
  geom_boxplot(aes(fill = `Root condition`), outlier.shape= NA, width=0.9, show.legend= FALSE) +
  geom_point(aes(colour= name, shape= Amplification),
             position= position_jitter(width= 0.1), alpha= 1) +
  stat_compare_means(label= "p.signif", method= "wilcox.test", paired= FALSE,
                     comparisons= list(c(1,2)),
                     tip.length= 0, step.increase= 0) +
  
  facet_wrap(~ Condition, scales= "free_x", strip.position="bottom") +
  scale_fill_manual(values = c("#99CCFF", "#6699FF")) +
  #scale_colour_manual(values= c("grey35", "red1"), name= "Bottleneck metrics",
  #                    labels= c("\nSTAMP Nb \n(one-step model) ", "\nSTAMPR Nr \n(multiple one-step model)")) +
  scale_colour_manual(values= c("chartreuse3", "red1"), name= "Bottleneck metrics",
                      labels= c("\nStandard STAMP \nNb equation", "\nCorrected STAMPR \nNr equation")) +
  scale_y_continuous(trans = log10_trans(), limits=c(1, 50000)) +
  scale_shape_manual(values= c(4,19), name= "Sample Statut", labels= c("No data","Sequenced")) +
  labs(title= "Root entry bottleneck", x= "Condition", y= "Founding population (number of bacteria)") +
  theme_gray() +
  theme(axis.text.x= element_blank(), axis.ticks.x= element_blank(), strip.background= element_blank())


####################
# Comparison Nb vs Nr vs richness in root
####################

tab = tab_tot %>%
  filter(Zone == "Root")

tab$Richness[is.na(tab$Richness)] = 0

# Joining Nb, Nr, Ns and richness together to compare them
tab = pivot_longer(tab, c(`Filtered bottleneck Nb`,`Corrected bottleneck Nr`, `Simulated bottleneck Ns`, Richness))

tab$name[tab$name == "Filtered bottleneck Nb"] = "Nb"
tab$name[tab$name == "Corrected bottleneck Nr"] = "Nr"
tab$name[tab$name == "Simulated bottleneck Ns"] = "Ns"

tab$name = factor(tab$name, levels= c("Richness", "Nb", "Nr", "Ns"))

tab = tab %>%
  unite("Condition3", Condition, name, sep= "_", remove= FALSE) %>%
  filter(Genotype == "M") %>%
  filter(`Temperature (°C)` == "27°C")

# Organising conditions
tab$Condition3 = factor(tab$Condition3, levels= c("M_Intact_Richness", "M_Intact_Nb", "M_Intact_Nr", "M_Intact_Ns", "M_Cut_Richness", "M_Cut_Nb", "M_Cut_Nr", "M_Cut_Ns"))
tab$`Root condition` = factor(tab$`Root condition`, levels= c("Intact", "Cut"))


# Graph v1
ggplot(data= tab, aes(x= name, y= value)) +
  
  geom_boxplot(aes(fill = `Root condition`), outlier.shape= NA, width=0.9, show.legend= FALSE) +
  geom_point(aes(colour= name, shape= Amplification), alpha= 1) +
  geom_line(aes(group= `Plant ID`), alpha= 0.3, size= 0.3, linetype= 1) +
  stat_compare_means(label= "p.signif", method= "wilcox.test", paired= TRUE,
                     comparisons= list(c(1,2), c(1,3),c(1,4)), step.increase= 0.07) +
  
  facet_wrap(~ Condition, scales= "free_x", strip.position="bottom") +
  scale_fill_manual(values = c("#99CCFF", "#6699FF")) +
  scale_colour_manual(values= c("grey35", "chartreuse3", "red1", "orange1"), name= "Bottleneck metrics",
                      labels= c("\nUnique tags \n(Richness)", "\nStandard STAMP \nNb equation",
                                "\nCorrected STAMPR \nNr equation", "\nSimulated STAMPR \nNs equation")) +
  scale_y_continuous(trans = log10_trans(), limits=c(1, 180000), breaks= c(1,10,100,1000,10000,100000)) +
  scale_shape_manual(values= c(4,19), name= "Sample Statut", labels= c("No data","Sequenced")) +
  labs(x= "Condition", y= "Founding population (number of bacteria)") +
  theme_gray() +
  theme(axis.text.x= element_blank(), axis.ticks.x= element_blank(), strip.background= element_blank())


# Second visualisation
tab = tab_tot %>%
  filter(Zone == "Root")

tab$Richness[is.na(tab$Richness)] = 0

# Joining Nb and Nr together to compare them
tab = pivot_longer(tab, c(`Filtered bottleneck Nb`,`Corrected bottleneck Nr`))

tab$name[tab$name == "Filtered bottleneck Nb"] = "Nb"
tab$name[tab$name == "Corrected bottleneck Nr"] = "Nr"
tab$name = factor(tab$name, levels= c("Nb", "Nr"))

tab = tab %>%
  unite("Condition3", Condition, name, sep= "_", remove= FALSE) %>%
  filter(Genotype == "M") %>%
  filter(`Temperature (°C)` == "27°C")

# Organising conditions
tab$Condition3 = factor(tab$Condition3, levels= c("M_Intact_Nb", "M_Intact_Nr", "M_Cut_Nb", "M_Cut_Nr"))
tab$`Root condition` = factor(tab$`Root condition`, levels= c("Intact", "Cut"))


# Creating a secondary table to connect Nb and Nr from a same sample on the graph
tab_temp = tab_tot %>%
  filter(Genotype == "M") %>%
  filter(`Temperature (°C)` == "27°C") %>%
  filter(Zone == "Root")

tab_temp$Richness[is.na(tab_temp$Richness)] = 0

# Graph v2
ggplot(data= tab, aes(x= Richness, y= value, colour = name)) +
  
  geom_point() +
  geom_function(fun= ~.x+0, aes(colour= "a")) +
  geom_segment(data= tab_temp, aes(x= Richness, y= `Filtered bottleneck Nb`,
                                   xend= Richness, yend= `Corrected bottleneck Nr`),
               colour= "black", alpha= 0.15, show.legend = FALSE) +
  
  facet_wrap(~ Condition, scales= "free_x", strip.position="bottom") +
  scale_x_continuous(trans = log10_trans(), limits=c(1, 120), n.breaks = 12) +
  scale_y_continuous(trans = log10_trans(), limits=c(1, 24000), breaks= c(1,10,100,1000,10000,100000)) +
  scale_colour_manual(values= c("chartreuse3", "red1", "grey35"), name= "Bottleneck metrics",
                      labels= c("\nStandard STAMP \nNb equation", "\nCorrected STAMPR \nNr equation", "Richness reference")) +
  labs(x= "Unique tags in sample (richness)", y= "Bottleneck metric (Nb or Nr)") +
  theme_gray() +
  theme(strip.background= element_blank(), strip.placement = "outside")


# Graph v3
ggplot(data= tab, aes(x= Richness, y= value, colour = name)) +
  
  geom_point() +
  geom_function(fun= ~.x+0, aes(colour= "a")) +
  geom_segment(data= tab_temp, aes(x= Richness, y= `Filtered bottleneck Nb`,
                                   xend= Richness, yend= `Corrected bottleneck Nr`),
               colour= "black", alpha= 0.15, show.legend = FALSE) +
  
  facet_wrap(~ Condition, scales= "free_x", strip.position="bottom") +
  scale_x_continuous(trans = log10_trans(), limits=c(1, 120), n.breaks = 10) +
  scale_y_continuous(trans = log10_trans(), limits=c(1, 24000), breaks= c(1,10,100,1000,10000,100000)) +
  scale_colour_manual(values= c("chartreuse3", "red1", "grey35"), name= "Bottleneck metrics",
                      labels= c("\nStandard STAMP \nNb equation", "\nCorrected STAMPR \nNr equation", "Richness reference")) +
  labs(x= "Unique tags in sample (richness)", y= "Bottleneck metric (Nb or Nr)") +
  theme_gray() +
  theme(strip.background= element_blank(), strip.placement = "outside", legend.position= "bottom")


####################
# Comparison of Shannon diversity in root
####################

tab = tab_tot

# Filtering out invalid samples
tab$`Shannon diversity`[tab$Amplification == 0 | is.na(tab$Amplification)] = NA

# Selection only the root samples of conditions M_Intact_27 and M_Cut_27
tab_temp = tab %>%
  filter(Genotype == "M") %>%
  filter(`Temperature (°C)` == "27°C") %>%
  filter(Zone == "Root")

tab_temp$`Root condition` = factor(tab_temp$`Root condition`, levels= c("Intact", "Cut"))


# Graph v1
ggplot(data= tab_temp, aes(x= Condition, y= `Shannon diversity`, fill= `Root condition`)) +
  
  geom_boxplot(outlier.shape= NA, width=0.9) +
  geom_point(aes(colour= Replicate), position =
               position_jitter(width = 0.2), alpha= 1) +
  stat_compare_means(label= "p.signif", method= "wilcox.test", paired= FALSE,
                     comparisons= list(c(1,2)),
                     tip.length= 0, step.increase= 0.1) +
  
  scale_fill_manual(values = c("#99CCFF", "#6699FF")) +
  scale_colour_manual(values= c("#E41A1C", "#339900", "#FFD92F")) +
  scale_y_continuous(limits=c(0,5), breaks= c(0:5)) +
  labs(x= "Condition", y = "Shannon diversity index") +
  theme_gray() +
  theme(legend.position= "bottom", legend.box= "vertical")


####################
# Shannon diversity in all inocula from the experiment
####################

# Shannon diversity in inocula
tab = tab_inoc %>%
  filter(`Inoculum ID 2` %in% c("A27","B27","C27","A32","B32","C32")) %>%
  filter(`Inoculum ID 3` == 1)

# Prefiltering using noise threshold
tab[,147:266][tab[,147:266] <= Noise_threshold] = 0
tab$`Shannon diversity` = diversity(tab[,147:266])


# Graph v1
ggplot(data= tab, aes(x = "Inoculum", y= `Shannon diversity`)) +

  geom_point(alpha= 1) +
  
  #scale_x_discrete(guide = guide_axis(angle = 90)) +
  scale_y_continuous(limits=c(0,5), breaks = c(0:5)) +
  labs(y = "Shannon diversity index", x= NULL) +
  theme_gray()

```

Statistical test of the contribution of each condition to Nr (tab_tot) 

```{r Calculation}

####################
# Effect quantification in Nr
####################

tab = tab_tot %>%
  filter(Amplification == TRUE)

tab = select(tab, c("Genotype", "Temperature (°C)", "Root condition", "Zone", "Corrected bottleneck Nr"))

# Encode each condition to logical
tab$Genotype = as.integer(tab$Genotype == "M")
tab$`Temperature (°C)` = as.integer(tab$`Temperature (°C)` == "27°C")
tab$`Root condition` = as.integer(tab$`Root condition` == "Intact")


# Non-parametric PERMANOVA for multivariate data
a = adonis2(formula = tab$`Corrected bottleneck Nr` ~ (Genotype + `Temperature (°C)` + `Root condition`)^2,
                       data = tab, permutations= 1000, by= "terms")
print(a)


# Spearman correlation between bacterial load in root and symptoms
tab = tab_tot %>%
  filter(Zone == "Root")

a = cor.test(tab$`Disease index`, tab$`Bacterial load (CFU/g)`, method= "spearman")
print(a)

```

### Additional graphs

Extracting day of sampling and symptoms development

```{r Graph}

####################
# Symptoms and day of sampling in the experiment with varying conditions (tab_tot)
####################

tab = unique(tab_tot)

tab = arrange(tab, by= tab$Condition2)
tab$`Time post-inoculation (d)` = factor(tab$`Time post-inoculation (d)`, levels= c(1:14))
tab$Condition2 = factor(tab$Condition2, levels= rev(levels(tab$Condition2)))

# Counting the proportion of symptomatic plants (Disease index > 1) in each condition
tab$`Symptomatic plant proportion` = 0

for (i in unique(tab$Condition2)) {
  tab_temp = tab_tot %>%
    filter(Condition2 == i)
  
  a = sum(tab_temp$Symptoms == 1) / length(tab_temp$`Sample ID`)
  
  tab$`Symptomatic plant proportion`[tab$Condition2 == i] = a
}

# Graph v1
ggplot(data= tab, aes(x= `Time post-inoculation (d)`, y= Condition2, fill= Condition)) +
  
  geom_tile(colour= 1, linewidth= 0.5, show.legend= FALSE) +
  
  scale_colour_manual(values= c("black")) +
  scale_x_discrete(drop= FALSE) +
  scale_fill_manual(values = c("#6699FF", "#6699FF", "#FF9900", "#FF9900"), na.value= "white") +
  labs(x= "Day of sampling post-inoculation (d)", y= "Condition") +
  theme_pubclean()

# Graph v2
ggplot(data= tab, aes(x = "", y= Condition2, fill= `Symptomatic plant proportion`)) +
  
  geom_tile(colour= 1, linewidth= 0.5, show.legend= FALSE) +
  geom_text(aes(label= scales::percent(`Symptomatic plant proportion`))) +
  
  scale_fill_gradient2(high= "red") +
  scale_x_discrete(drop= FALSE) +
  labs(x= "% symptomatic plants", y= "Condition") +
  theme_pubclean()

```

## Description of the infection with sampling over time (tab_kinetic)

### Visualisation of the condition M_Intact_27

Nb, Nr and internal growth rate curve

```{r Graph}

####################
# Bacterial load over time
####################

tab = tab_kinetic %>%
  filter(Condition2 == "M_Intact_27")

# Graph v1
ggplot(data= tab, aes(x= `Time post-inoculation (d)`, y= `Bacterial load (CFU/g)`, fill= Genotype)) +
  
  geom_violin(data= tab, width= 0.95, show.legend= FALSE) +
  stat_boxplot(aes(ymin = after_stat(lower), ymax = after_stat(upper)), geom="errorbar", width=.25) + 
  stat_boxplot(aes(ymin = after_stat(middle), ymax = after_stat(middle)), geom="errorbar", linewidth=1.5, width=.25) +
  geom_point(aes(fill= Genotype, colour= Replicate),
             position= position_jitterdodge(jitter.width= 0.6, dodge.width= 1), alpha= 0.7, show.legend= FALSE) +
  
  scale_fill_manual(values= c("#6699FF")) +
  scale_colour_manual(values= c("#E41A1C")) +
  scale_y_continuous(trans = log10_trans(), limits=c(1, NA)) +
  theme_gray()


# Formating data for linear regression
tab_temp = data.frame(Zone= factor(), N0= numeric(), std_error_N0= numeric(),
                      Rate= numeric(), std_error_Rate= numeric(), Adj.rr= numeric())

a = summary(lm(data= filter(tab, Zone == "Root"), log(`Bacterial load (CFU/g)`) ~ as.integer(`Time post-inoculation (d)`)))
tab_temp = rbind(tab_temp, c("Root", a$coefficients[1], a$coefficients[3],
                             a$coefficients[2], a$coefficients[4], a$adj.r.squared))

names(tab_temp) = c("Zone", "N0", "std_error_N0", "Rate", "std_error_Rate", "Adj.rr")

tab_temp[,2:6] = lapply(tab_temp[,2:6], as.numeric)

tab_temp$Equation = paste(tab_temp$Zone,
                          "   y= exp(",round(tab_temp$Rate, 2), "t",
                          ifelse(tab_temp$N0 >= 0, "+",""),
                          round(tab_temp$N0, 2) , ")", sep= "")

tab_temp$Equation2 = paste("y= exp(",round(tab_temp$Rate, 2), "t",
                          ifelse(tab_temp$N0 >= 0, "+",""),
                          round(tab_temp$N0, 2) , ")", sep= "")

# Graph v3 
ggplot(data= tab, aes(x= as.numeric(as.character(`Time post-inoculation (d)`)), y= `Bacterial load (CFU/g)`, colour= Zone)) +
  
  geom_point() +
  geom_smooth(method= "lm", formula= y~x, se= FALSE) +
  #stat_regline_equation(aes(label= paste(after_stat(eq.label), after_stat(rr.label), sep= "~~~~")), formula = I(log(10^y))~x) +
  annotate("text", colour= rev(c("#F8766D")), x=2.95, y= c(10e8),
           label= rev(paste(tab_temp$Equation, "   R²= ", round(as.numeric(tab_temp$Adj.rr), 2), sep= ""))) +
  
  scale_y_continuous(trans= "log10",
                     breaks = trans_breaks("log10", function(x) 10^x),
                     labels = trans_format("log10", math_format(10^.x))) +
  labs(x= "Time post-inoculation (d)") +
  theme_gray() +
  theme(legend.position="bottom")


# Calculation of residual effect not explained by exponential growth
tab$`Residual Load (CFU/g)` = NA

for (i in levels(tab$Zone)) {
  tab$`Residual Load (CFU/g)`[tab$Zone == i] = (tab$`Bacterial load (CFU/g)`[tab$Zone == i] -
                                                      exp(tab_temp$Rate[tab_temp$Zone == i] *
                                                            as.integer(tab$`Time post-inoculation (d)`[tab$Zone == i]) +
                                                            tab_temp$N0[tab_temp$Zone == i]))
  }

tab_temp2 = tab_temp

# Graph v4
ggplot(data= tab, aes(x= as.numeric(as.character(`Time post-inoculation (d)`)), y= `Residual Load (CFU/g)`, colour= Zone)) +
  
  geom_point() +
  labs(x= "Time post-inoculation (d)") +
  theme_gray() +
  theme(legend.position="bottom")


####################
# Nb and Nr over time
####################

tab = tab_kinetic %>%
  filter(Condition2 == "M_Intact_27")

tab$`Time post-inoculation (d)` = factor(tab$`Time post-inoculation (d)`, levels= c(1,2,3,4,5,6))

tab = tab %>%
  filter(!is.na(`Filtered bottleneck Nb`))


# Graph v1

ggplot(data= tab, aes(x= Zone, y= `Filtered bottleneck Nb`, fill= Zone)) +
  
  geom_pointrange(aes(fill= Zone, colour= Replicate,
                    ymax= (`Filtered bottleneck Nb` + `Nb mathematical standard deviation STD_Nb`), ymin= (`Filtered bottleneck Nb` - `Nb mathematical standard deviation STD_Nb`)),
                  size= 0.2, alpha= 1, show.legend= FALSE) +
  geom_line(aes(group= `Plant ID`, colour= Replicate), alpha= 1, size= 0.5, linetype= 2, show.legend= FALSE) +
  facet_grid(~ `Time post-inoculation (d)`, scales= "free_x", switch= "x",
             labeller= labeller(`Time post-inoculation (d)` = ~ paste("DPI ", .x)), drop= FALSE) +
  
  scale_fill_manual(values= c("#CCFFFF", "#99FFFF", "#66FFFF", "#33FFFF", "#00FFFF")) +
  scale_colour_manual(values= c("#E41A1C")) +
  scale_x_discrete(guide = guide_axis(angle = 90)) +
  scale_y_continuous(trans = log10_trans(), limits=c(1, NA)) +
  labs(y = "Bottleneck Nb (number bacteria)           ") +
  theme_gray()


# Graph v2

ggplot(data= tab, aes(x= Zone, y= `Corrected bottleneck Nr`, fill= Zone)) +
  
  geom_pointrange(aes(fill = Zone, colour= Replicate,
                      ymax= `Corrected bottleneck Nr` + `Nr mathematical standard deviation STD_Nr`,
                      ymin= `Corrected bottleneck Nr` - `Nr mathematical standard deviation STD_Nr`),
                  size= 0.2, alpha= 1, show.legend= FALSE) +
  geom_line(aes(group= `Plant ID`, colour= Replicate), alpha= 1, size= 0.5, linetype= 2, show.legend= FALSE) +
  
  facet_grid(~ `Time post-inoculation (d)`, scales= "free_x", switch= "x",
             labeller= labeller(`Time post-inoculation (d)` = ~ paste("DPI ", .x)), drop= FALSE) +
  scale_fill_manual(values= c("#CCFFFF", "#99FFFF", "#66FFFF", "#33FFFF", "#00FFFF")) +
  scale_colour_manual(values= c("#E41A1C")) +
  scale_x_discrete(guide = guide_axis(angle = 90)) +
  scale_y_continuous(trans = log10_trans(), limits=c(1, NA)) +
  labs(y = "Corrected bottleneck Nr (number bacteria)           ") +
  theme_gray()


####################
# Kinetics of bacteria load and shannon diversity over time
####################

tab = tab_kinetic %>%
  filter(Condition2 == "M_Intact_27")

tab$`Time post-inoculation (d)` = as.numeric(as.character(tab$`Time post-inoculation (d)`))
tab$Zone = factor(tab$Zone, levels= c("High_Stem", "Mid_Stem", "Low_Stem", "Root"))

# Graph v1
ggplot(data= tab, aes(x= `Time post-inoculation (d)`)) +
  
  geom_point(aes(y= `Bacterial load (CFU/g)`), colour= "blue") +
  geom_point(aes(y= `Filtered bottleneck Nb`), colour= "red") +
  
  #geom_smooth(aes(y= `Bacterial load (CFU/g)`), method= "loess", colour= "blue", se= FALSE) +
  stat_summary(aes(y= `Bacterial load (CFU/g)`), fun.y=mean, geom="line", colour = "blue" , alpha=0.8) +
  stat_summary(aes(y= `Bacterial load (CFU/g)`), fun.data=mean_sd, geom="ribbon", fill = "blue" , alpha=0.15) + 
  
  #geom_smooth(aes(y= `Filtered bottleneck Nb`), method= "loess", colour= "red", se= FALSE) +
  stat_summary(aes(y= `Filtered bottleneck Nb`), fun.y=mean, geom="line", colour = "red" , alpha=0.8) +
  stat_summary(aes(y= `Filtered bottleneck Nb`), fun.data=mean_sd, geom="ribbon", fill = "red" , alpha=0.15) +
  
  facet_wrap(~ Zone, scales= "free_x", ncol= 1) +
  scale_y_continuous(trans= "log10", name= "Bacterial load (CFU/g)",
                     breaks = trans_breaks("log10", function(x) 10^x),
                     labels = trans_format("log10", math_format(10^.x)),
                     sec.axis = sec_axis(~.*1, name= "Bottleneck Nb (number bacteria)",
                                         breaks = trans_breaks("log10", function(x) 10^x),
                                         labels = trans_format("log10", math_format(10^.x))) ) +
  
  scale_x_continuous(breaks= c(1:6)) +
  coord_cartesian(xlim= c(1,6), ylim= c(1,NA)) +
  labs(x= "Time post-inoculation (d)") +
  theme_minimal() +
  theme(axis.title.y.left = element_text( color="blue"),
        axis.line.y.left = element_line(colour = "blue"),
        axis.text.y.left = element_text(color="blue"),
        axis.title.y.right = element_text( color="red"),
        axis.line.y.right = element_line(colour = "red"),
        axis.text.y.right = element_text(color="red"))


# Comparing only the root samples
tab_temp = tab %>%
  filter(Zone == "Root")

# Graph v2
ggplot(data= tab_temp, aes(x= `Time post-inoculation (d)`, y= `Bacterial load (CFU/g)`)) +
  
  geom_point(aes(y= `Bacterial load (CFU/g)`), colour= "blue") +
  
  geom_smooth(aes(y= `Bacterial load (CFU/g)`), method= "lm", colour= "black", se= FALSE, linetype= "dashed") +
  annotate("text", colour= c("black"), x=2.8, y= c(10e7),
           label= paste(tab_temp2$Equation2[1], "   R²= ", round(as.numeric(tab_temp2$Adj.rr[1]), 2), sep= "")) +
  stat_summary(aes(y= `Bacterial load (CFU/g)`), fun.y=mean, geom="line", colour = "blue" , alpha=0.8) +
  stat_summary(aes(y= `Bacterial load (CFU/g)`), fun.data=mean_sd, geom="ribbon", fill = "blue" , alpha=0.15) + 
  
  scale_y_continuous(trans= "log10", name= "Bacterial load (CFU/g)",
                     breaks = trans_breaks("log10", function(x) 10^x),
                     labels = trans_format("log10", math_format(10^.x))) +
  
  scale_x_continuous(breaks= c(1:6)) +
  coord_cartesian(xlim= c(1,6), ylim= c(1,NA)) +
  labs(x= "Time post-inoculation (d)") +
  theme_minimal() +
  theme(axis.title.y.left = element_text( color="blue"),
        axis.line.y.left = element_line(colour = "blue"),
        axis.text.y.left = element_text(color="blue"))


# Comparing Shannon diversity in valid root samples
tab_temp = tab %>%
  filter(Zone == "Root") %>%
  filter(Amplification == TRUE)

# Graph v3
ggplot(data= tab_temp, aes(x= `Time post-inoculation (d)`)) +
  
  geom_point(aes(y= `Shannon diversity`), colour= "red") +
  
  #geom_smooth(aes(y= `Shannon diversity`), method= "loess", colour= "red", se= FALSE) +
  stat_summary(aes(y= `Shannon diversity`), fun.y=mean, geom="line", colour = "red" , alpha=0.8) +
  stat_summary(aes(y= `Shannon diversity`), fun.data=mean_sd, geom="ribbon", fill = "red" , alpha=0.15) +
  
  scale_y_continuous(name= "Shannon diversity\nindex") +
  scale_x_continuous(breaks= c(1:6)) +
  coord_cartesian(xlim= c(1,6), ylim= c(0,5)) +
  labs(x= "Time post-inoculation (d)") +
  theme_minimal() +
  theme(axis.title.y.left = element_text( color="red"),
        axis.line.y.left = element_line(colour = "red"),
        axis.text.y.left = element_text(color="red"))


# Complementary Shannon diversity reference in inoculum
tab_temp = tab_inoc %>%
  filter(`Inoculum ID 2` == "S27") %>%
  filter(`Inoculum ID 3` == 1)

# Prefiltering using noise threshold
tab_temp[,147:266][tab_temp[,147:266] <= Noise_threshold] = 0
tab_temp$`Shannon diversity` = diversity(tab_temp[,147:266])

# Graph v4
ggplot(data= tab_temp, aes(x= "")) +
  
  geom_point(aes(y= `Shannon diversity`)) +
  
  scale_y_continuous(name= "Shannon diversity\nindex") +
  coord_cartesian(ylim= c(0,5)) +
  labs(x= "Inoculum") +
  theme_minimal() +
  theme(axis.title.y.left = element_text( color="red"),
        axis.line.y.left = element_line(colour = "red"),
        axis.text.y.left = element_text(color="red"))

```

## General exploratory analyses on the dataset

Exploratory tests of predictions of initial timings of infection based on bacterial load and Nb

```{r Graph}

####################
# Prediction of the initial bacterial load under the assumption of a STAMP model
####################

tab = tab_tot %>%
  filter(Zone == "Root") %>%
  filter(`Filtered bottleneck Nb` < 10000)

# Calculation the predicted initial load at t0 under the assumption of a STAMP model of entry
tab$`Putative initial load (CFU/g)` = tab$`Filtered bottleneck Nb` / tab$`Collected fresh weight (g)`

# Graph v1

ggplot(data= tab, aes(x= Condition, y= `Putative initial load (CFU/g)`, fill= Genotype)) +
  
  geom_boxplot(outlier.shape= NA, width=0.9) +
  geom_point(aes(fill= Genotype, colour= Replicate),
             position= position_jitterdodge(jitter.width= 0.1, dodge.width= 1), alpha= 1) +
  
  facet_wrap(~ `Temperature (°C)`, scales= "free_x", strip.position="bottom") +
  scale_fill_manual(values = c("#99CCFF", "#FFCC99")) +
  scale_colour_manual(values= c("#E41A1C", "#339900", "#FFD92F")) +
  scale_y_log10(limits = c(1, 2000)) +
  labs(title= "Root", y = "Predicted load at moment of root infection (bacteria/g)") +
  theme_gray()


# Prediction of the growth curve between the initial load (determined through Nb) and the final load
tab_temp = pivot_longer(tab, c(`Putative initial load (CFU/g)`,`Bacterial load (CFU/g)`))
tab_temp$name[tab_temp$name == "Putative initial load (CFU/g)"] = 1
tab_temp$name[tab_temp$name == "Bacterial load (CFU/g)"] = tab_temp$`Time post-inoculation (d)`[tab_temp$name == "Bacterial load (CFU/g)"]


# Graph v2
ggplot(data= tab_temp, aes(x= as.numeric(name), y= value, colour= Zone)) +
  
  geom_point() +
  geom_smooth(method= "lm", formula= y~x, se= FALSE) +
  stat_regline_equation(aes(label= paste(after_stat(eq.label), after_stat(rr.label), sep= "~~~~")), formula = I(log(10^y))~x) +
  
  facet_wrap(~ Condition2, scales= "free_x", strip.position="bottom") +
  scale_y_continuous(trans= "log10",
                     breaks = trans_breaks("log10", function(x) 10^x),
                     labels = trans_format("log10", math_format(10^.x))) +
  labs(x= "Time post-inoculation (d)") +
  theme_gray() +
  theme(legend.position="bottom")


# Graph v3
ggplot(data= tab, aes(x= `Putative initial load (CFU/g)`, y= `Bacterial load (CFU/g)`)) +
  
  geom_point() +
  geom_smooth(method= "lm", formula=y~x+0) +
  stat_regline_equation(aes(label= paste(after_stat(eq.label), after_stat(rr.label), sep= "~~~~")), formula = y~x) +
  
  facet_wrap(~ Condition2, scales= "free_x", strip.position="bottom") +
  #scale_y_continuous(trans= "log10",
  #                   breaks = trans_breaks("log10", function(x) 10^x),
  #                   labels = trans_format("log10", math_format(10^.x))) +
  theme_gray() +
  theme(legend.position="bottom")


####################
# Correlation of entry vs final observed load
####################

# Spearman correlation: initial load vs final load
a = cor.test(tab$`Putative initial load (CFU/g)`, tab$`Bacterial load (CFU/g)`, method= "spearman")
print(a)

# Spearman correlation: initial load vs final load per individual conditions
for (i in levels(tab$Condition2)) {
  
  tab_temp = tab %>%
    filter(Condition2 == i)
  
  a = cor.test(tab_temp$`Putative initial load (CFU/g)`, tab_temp$`Bacterial load (CFU/g)`, method= "spearman")
  print(a)
}

# Spearman correlation: Shannon vs final load
a = cor.test(tab$`Shannon diversity`, tab$`Bacterial load (CFU/g)`, method= "spearman")
print(a)

# Spearman correlation: Shannon diversity vs net number of CFU
tab$`Net Number of CFU` = tab$`Bacterial load (CFU/g)` * tab$`Collected fresh weight (g)`

a = cor.test(tab$`Shannon diversity`, tab$`Net Number of CFU`, method= "spearman")
print(a)


####################
# Prediction of the initial bacterial load in the infection with sampling over time in M_Intact_27 (tab_kinetic)
####################

tab = tab_kinetic %>%
  filter(Condition2 == "M_Intact_27") %>%
  filter(Zone == "Root")

# Calculation the predicted initial load at t0 under the assumption of a STAMP model of entry
tab$`Putative initial load (CFU/g)` = tab$`Filtered bottleneck Nb` / tab$`Collected fresh weight (g)`

# Calculating a normalised load at t0 based on theparameter r=2.23 found by fitting an exponential growth on data
tab$`Normalised load t0 (CFU/g)` = tab$`Bacterial load (CFU/g)` / (exp(2.23 * as.numeric(tab$`Time post-inoculation (d)`)))

# Graph v1
ggplot(data= tab, aes(x= `Putative initial load (CFU/g)`, y= log(`Normalised load t0 (CFU/g)`))) +
  
  geom_point() +
  geom_smooth(method= "lm", formula=y~x+0) +
  stat_regline_equation(aes(label= paste(after_stat(eq.label), after_stat(rr.label), sep= "~~~~")), formula = y~x) +
  
  #scale_y_continuous(trans= "log10",
  #                   breaks = trans_breaks("log10", function(x) 10^x),
  #                   labels = trans_format("log10", math_format(10^.x))) +
  theme_gray() +
  theme(legend.position="bottom")


# Spearman correlation: initial load (based on Nb) and normalised bacterial load in root
a = cor.test(tab$`Putative initial load (CFU/g)`, tab$`Normalised load t0 (CFU/g)`, method= "spearman")
print(a)

```

Exploratory simulations regarding the behaviour of a STAMP model bottleneck

```{r Simulation}

####################
# Simulation of a perfect infinite population passing through two successive bottlenecks Nb1 and Nb2
####################

# Number of tags k in the study
k = 119

# Range of tested primary bottleneck Nb1
Nb1_range = c(1,10,50)

# Tested secondary bottleneck Nb2
Nb2_range = c(100,10000)

# Number of iterations
R = 1000

# Initialisation of some variables for later
Ne1_range = data.frame(Nb1= integer(), Ne1= integer())
Ne2_range = data.frame(Nb1= integer(), Nb2= integer(), Ne2= integer(), Netot= integer())
STD1_range = data.frame(Nb1= integer(), STD_Ne1= integer())
STD2_range = data.frame(Nb1= integer(), Nb2= integer(), STD_Ne2= integer(), STD_Netot= integer())
tags2 = matrix(nrow= k, ncol= R)

# Simulation of the first bottleneck
for (i in Nb1_range) {
  
  # Random sampling for the primary bottleneck Nb1
  tags1 = rmultinom(R, i, rep(1/k,k))
  
  # Calculation of the estimation of the bottleneck Ne1 based on the equation
  f1 = tags1 / i
  F_ratio = ((f1 - 1/k)^2) / (1/k*(1-1/k))
  Ne1 = 1 / colMeans(F_ratio)
  Ne1[Ne1 == Inf] = NA
  Ne1_range = rbind(Ne1_range, data.frame(Nb1= i, Ne1= Ne1))
  STD1_range = rbind(STD1_range, data.frame(Nb1= i, STD_Ne1= sd(Ne1)))
  
  # Simulation of a secondary bottleneck
  for (j in Nb2_range) {
    
    # Random sampling for the secondary bottleneck Nb2 
    for (l in c(1:R)) {
    tags2[,l] = rmultinom(1, j, f1[,l])
    }
  
    # Calculation of the estimation of the bottleneck Ne2 based on the equation
    f2 = tags2 / j
    F_ratio = ((f2 - f1)^2) / (f1*(1-f1))
    F_ratio[is.nan(F_ratio)] = NA
    Ne2 = 1 / colMeans(F_ratio, na.rm = TRUE)
    Ne2[Ne2 == Inf] = NA
  
    # Calculation of the total bottleneck Netot when considering both primary and secondary bottleneck simultaneously
    F_ratio = ((f2 - 1/k)^2) / (1/k*(1-1/k))
    Netot = 1 / colMeans(F_ratio)
    Ne2_range = rbind(Ne2_range, data.frame(Nb1= i, Nb2= j, Ne2= Ne2, Netot))
    STD2_range = rbind(STD2_range, data.frame(Nb1= i, Nb2= j, STD_Ne2= sd(Ne2), STD_Netot= sd(Netot)))
  }
}


# Extracting standard deviation on Ne1
simul_fit = summary(lm(data = STD1_range, STD_Ne1 ~ Nb1+0))
print(simul_fit)


# Simulation of the mathematical error on a single bottleneck event Nb1
ggplot(data= Ne1_range, aes(x= Nb1, y= Ne1)) +
  geom_point(fill = "gray", alpha= 0.1) +
  stat_summary(fun.data=mean_sd, geom="ribbon", fill = "red" , alpha=0.25) +
  geom_smooth(method= "lm", formula=y~x+0) +
  labs(title= "Mathematical error on one bottleneck Nb1")
  
ggplot(data= STD1_range, aes(x= Nb1, y= STD_Ne1)) +
  geom_point() +
  geom_smooth(method= "lm", formula=y~x+0) +
  stat_regline_equation(formula = y~x+0, ) +
  labs(title= "Mathematical error on one bottleneck Nb1 (standard deviation)")


# Simulation of the mathematical error caused by Nb1 on Nb2
ggplot(data= Ne2_range, aes(x= factor(Nb2), y= Ne2)) +
  facet_wrap(~ Nb1, scales = "free_y", strip.position= "bottom", labeller= labeller(Nb1 = ~ paste("Nb1=", .x))) +
  geom_boxplot() +
  scale_y_continuous(limits= c(0,15000)) +
  labs(title= "Mathematical error on a second bottleneck Nb2")

ggplot(data= STD2_range, aes(x= Nb2, y= STD_Ne2)) +
  facet_wrap(~ Nb1, scales = "free_y", strip.position= "bottom", labeller= labeller(Nb1 = ~ paste("Nb1=", .x))) +
  geom_point() +
  geom_smooth(method= "lm", formula= y ~ x+0 ) +
  stat_regline_equation(formula = y ~ x+0, ) +
  labs(title= "Mathematical error on a second bottleneck Nb2 (standard deviation")


# Estimation of the cumulated bottleneck Netot after the double bottleneck event
ggplot(data= Ne2_range, aes(x= factor(Nb2), y= Netot)) +
  facet_wrap(~ Nb1, scales = "free_y", strip.position= "bottom", labeller= labeller(Nb1 = ~ paste("Nb1=", .x))) +
  geom_boxplot() +
  labs(title= "Cumulated bottleneck Nbtot after two events of bottleneck Nb1 and Nb2")


ggplot(data= Ne2_range, aes(x= Nb2, y= Netot)) +
  facet_wrap(~ Nb1, scales = "free_y", strip.position= "bottom", labeller= labeller(Nb1 = ~ paste("Nb1=", .x))) +
  geom_point(fill = "gray", alpha= 0.1) +
  stat_summary(fun.data=mean_sd, geom="ribbon", fill = "red" , alpha=0.25) +
  labs(title= "Cumulated bottleneck Nbtot after two events of bottleneck Nb1 and Nb2")


# Distribution of tag counts in simulations
for (i in c(1:R)) {
  tags1[,i] = sort(tags1[,i])
}

tab_temp = data.frame(Tags= c(1:k), count= rowSums(tags1))
tab_temp$count = tab_temp$count / sum(tab_temp$count)

ggplot(tab_temp, aes(x= Tags, y= count)) +
    geom_hline(aes(yintercept= 1/k), colour= "blue") +
    geom_col()


####################
# # Simulation of single bottleneck Nb in function of the number of tags k
####################

# Range of tested tags k in the study
k_range = c(5,10,20,30,40,50,60,70,80,90,100)

# Tested primary bottleneck Nb1
Nb1 = 200

# Number of simulations
R = 10000

# Initialisation of some variables for later
Ne1_range = data.frame(Nb1= integer(), k= integer(), Ne1= integer())
STD1_range = data.frame(Nb1= integer(), k= integer(), STD_Ne1= integer())


# Testing each condition of tags
for (i in k_range) {
  
  # Random sampling for the primary bottleneck Nb1 
  tags1 = rmultinom(R, Nb1, rep(1/i,i))
  
  # Calculation of the estimation of the bottleneck Ne1 according to equation
  f1 = tags1 / Nb1
  F_ratio = ((f1 - 1/i)^2) / (1/i*(1-1/i))
  Ne1 = 1 / colMeans(F_ratio)
  Ne1[Ne1 == Inf] = NA
  Ne1_range = rbind(Ne1_range, data.frame(Nb1= Nb1, k= i, Ne1= Ne1))
  STD1_range = rbind(STD1_range, data.frame(Nb1= Nb1, k= i, STD_Ne1= sd(Ne1)))
}


# Simulation of the mathematical error on a single bottleneck event Nb1
ggplot(data= Ne1_range, aes(x= k, y= Ne1)) +
  geom_point(fill = "gray", alpha= 0.1) +
  stat_summary(fun.data=mean_sd, geom="ribbon", fill = "red" , alpha=0.25) +
  geom_smooth(method= "lm", formula=y~x+0) +
  labs(title= paste0("Simulation of the mathematical from number of tags using tested Nb1 =", Nb1))

ggplot(data= Ne1_range, aes(x= factor(k), y= Ne1)) +
  geom_boxplot() +
  geom_hline(aes(yintercept= Nb1, colour= "red")) +
  scale_y_continuous(limits= c(0,1000)) +
  labs(title= paste0("Simulation of the mathematical from number of tags using tested Nb1 =", Nb1))


####################
# Simulation of a cluster effect on a bottleneck Nb1
# Simulation of a temporal effect on a bottleneck Nb1
####################

# Range of tested tags k in the study
k = 119

# Tested primary bottleneck Nb1
Nb1 = 10000

# Number of simulations
R = 1000

# Sets of tested founder effect and clustering effect parameters (always enter trio)
# F_strength: standard deviation of normal noise added to initial 1/k distribution
# C_probability: probability that a tag becomes a cluster of bacteria
# C_strength: standard deviation of normal noise added to cluster tags
Param_range = data.frame(F_strength= c(0, 0.004, 0.04, 0.4),
                         C_probability= c(0, 0, 0, 0),
                         C_strength= c(0, 0, 0, 0),
                         Parameter= c("0", "1", "2", "3"))

Param_range$Parameter = factor(Param_range$Parameter, levels= Param_range$Parameter)

# Initialisation of some variables for later
Ne1_range = data.frame()
STD1_range = data.frame()

# Testing all i couples of parameters
for (i in c(1:nrow(Param_range))) {
  
  tags1 = matrix(nrow= k, ncol= 0)
  
  for (j in c(1:R)) {
    
    # Founder effect simulated as normal noise added to perfect equiprobability 1/k
    effects = data.frame(Tags= c(1:k), F_effect= 1/k + 1/k * abs(rnorm(1:k, 0, Param_range$F_strength[i])))
    effects$F_effect = effects$F_effect / sum(effects$F_effect)

    # Cluster effect simulated as a few tags with a very high probability to appear
    effects$cluster_success = as.logical(sample(c(0,1), k, replace= TRUE,
                                                prob= c(1-Param_range$C_probability[i], Param_range$C_probability[i])))
    effects$C_effect = (1/k + abs(rnorm(1:k, 0, Param_range$C_strength[i]))) * effects$cluster_success

    # Combinating effects
    effects$p = effects$F_effect + effects$C_effect
    effects$p = effects$p / sum(effects$p)
    
    # Random sampling for the primary bottleneck Nb1 and calculation of its estimation Ne1
    tags1 = cbind(tags1, rmultinom(1, Nb1, effects$p))
  }
  
  # Estimation of the bottleneck Ne1 according to equation
  f1 = tags1 / Nb1
  F_ratio = ((f1 - 1/k)^2) / (1/k*(1-1/k))
  Ne1 = 1 / (colMeans(F_ratio))
  Ne1[Ne1 == Inf] = NA
  
  Ne1_range = rbind(Ne1_range, data.frame(Nb1= Nb1, k= k, Ne1= Ne1, Param_ID= i,
                                          Parameter= Param_range$Parameter[i],
                                          F_strength= Param_range$F_strength[i],
                                          C_probability= Param_range$C_probability[i],
                                          C_strength= Param_range$C_strength[i]))
  
  STD1_range = rbind(STD1_range, data.frame(Nb1= Nb1, k= k, STD_Ne1= sd(Ne1), Ne1= Ne1, Param_ID= i,
                                          Parameter= Param_range$Parameter[i],
                                          F_strength= Param_range$F_strength[i],
                                          C_probability= Param_range$C_probability[i],
                                          C_strength= Param_range$C_strength[i]))
}


# Graphs of the last simulated founder and cluster effects
  ggplot(arrange(effects, F_effect), aes(x= c(1:k), y= F_effect)) +
    geom_hline(aes(yintercept= 1/k), colour= "blue") +
    geom_col()

  ggplot(arrange(effects, C_effect), aes(x= c(1:k), y= C_effect)) +
    geom_hline(aes(yintercept= 1/k), colour= "blue") +
    geom_col()

  ggplot(arrange(effects, p), aes(x= c(1:k), y= p)) +
    geom_hline(aes(yintercept= 1/k), colour= "blue") +
    geom_col()


# Simulation of the mathematical error with noise effect
ggplot(data= Ne1_range, aes(x= Parameter, y= Ne1)) +
  geom_boxplot() +
  geom_hline(aes(yintercept= Nb1), colour= "red") +
  #coord_cartesian(ylim= c(0,2000)) +
  labs(title= paste0("Simulation of the mathematical error for an imperfect bottleneck Nb1 =", Nb1))

ggplot(data= Ne1_range, aes(x= C_strength, y= Ne1)) +
  geom_point(fill = "gray", alpha= 0.1) +
  stat_summary(fun.data=mean_sd, geom="ribbon", fill = "red" , alpha=0.25) +
  #geom_smooth(method= "lm", formula=y~x) +
  labs(title= paste0("Simulation of the mathematical error for an imperfect bottleneck Nb1 =", Nb1))

ggplot(data= Ne1_range, aes(x= F_strength, y= Ne1)) +
  geom_point(fill = "gray", alpha= 0.1) +
  stat_summary(fun.data=mean_sd, geom="ribbon", fill = "red" , alpha=0.25) +
  #geom_smooth(method= "lm", formula=y~x) +
  labs(title= paste0("Simulation of the mathematical error for an imperfect bottleneck Nb1 =", Nb1))


# Distribution of tag counts in simulations
for (i in c(1:R)) {
  tags1[,i] = sort(tags1[,i])
}

tab_temp = data.frame(Tags= c(1:k), count= rowSums(tags1))
tab_temp$count = tab_temp$count / sum(tab_temp$count)

ggplot(tab_temp, aes(x= Tags, y= count)) +
    geom_hline(aes(yintercept= 1/k), colour= "blue") +
    geom_col()


####################
# Simulation of a perfect STAMP bottleneck but using real inoculum frequency
####################

# Iteration
R = 3

# Simulated bottleneck strength
Nb1 = 200

# Used inoculum frequencies (drawn from inoculum table tab_inoc)
f1 = tab_inoc[1,147:266] / sum(tab_inoc[1,147:266])

# Random sampling for the bottleneck
tags1 = rmultinom(R, Nb1, f1)


# Adding simulation index
tab_temp = as.data.frame(t(tags1[,1:R]))
tab_temp$ID = factor(c(1:3))
colnames(tab_temp)[1:120] = c(1:120)

# Ordering frequencies from lowest to highest
for (i in c(1:nrow(tab_temp))) {
  tab_temp[i,1:119] = tab_temp[i,1:119][rev(order(tab_temp[i,1:119]))]
}

tab_temp = pivot_longer(tab_temp, cols = c(1:119))
tab_temp$name = factor(tab_temp$name, levels= c(1:119))

# ordered graph
ggplot(data= tab_temp, aes(x= ID, y= value, fill = name)) +
 
  #facet_grid( ~ ID, scales= "free_x") +
  geom_col(position="fill", colour= "black", size= 0.2, show.legend = FALSE) +
  
  scale_y_continuous(labels= scales::percent) +
  scale_x_discrete(guide= NULL) +
  coord_cartesian(ylim = c(0,1.05)) +
  labs(title= "Nb = 10", x= "", y= "Frequency of tags") +
  theme_pubr() +
  theme(strip.background= element_blank(), strip.text= element_blank(), panel.spacing= unit(0, "lines"))

```

Extraction of stackplots to illustrate STAMP diversity

N.B.: for simplicity, some stackplots were generated despite corresponding to invalid samples. The corresponding stackplots were manually curated afterward.

```{r Graph}

####################
# Diversity in all inocula
####################

tab = tab_inoc %>%
  filter(`Inoculum ID 3` == 1) %>%
  filter(!is.na(`PCR2 validation on gel`))

tab$`Inoculum ID 2` = factor(tab$`Inoculum ID 2`, levels= c("S27","A27","B27","C27","A32","B32","C32"))

tab_temp = pivot_longer(tab, cols = c(147:266))
tab_temp$name = factor(tab_temp$name)


# Prefiltering using noise threshold
tab_temp$value[tab_temp$value <= Noise_threshold] = 0

# Graph
ggplot(data= tab_temp, aes(x= `Inoculum ID 2`, y= value, fill = name)) +
  facet_grid( ~ `Inoculum ID 2`, scales= "free_x") +
  geom_col(position="fill", colour= "black", size= 0.2, show.legend = FALSE) +
  scale_fill_manual(values= P120) +
  scale_y_continuous(labels= scales::percent) +
  scale_x_discrete(guide = NULL) +
  coord_cartesian(ylim = c(0,1.05)) +
  labs(title= "Inoculum", x= "", y= "Frequency of tags") +
  theme_pubr()


####################
# Diversity in inocula from tab_tot
####################

tab = tab_inoc %>%
  filter(`Inoculum ID 3` == 1) %>%
  filter(`Inoculum ID 2` %in% c("A27","B27","C27","A32","B32","C32"))

tab$`Inoculum ID 2` = factor(tab$`Inoculum ID 2`, levels= c("A27","B27","C27","A32","B32","C32"))

# Producing individual graph of each inoculum
for (i in tab$`Inoculum ID 2`) {
  
  # Organising data
  tab_temp = tab %>%
    filter(`Inoculum ID 2` == i)
  tab_temp = pivot_longer(tab_temp, cols = c(147:266))
  tab_temp$name = factor(tab_temp$name)

  # Prefiltering using noise threshold
  tab_temp$value[tab_temp$value <= Noise_threshold] = 0
  
  # Graph
  g = ggplot(data= tab_temp, aes(x= `Inoculum ID 2`, y= value, fill = name)) +
    facet_grid( ~ `Inoculum ID 2`, scales= "free_x") +
    geom_col(position="fill", colour= "black", size= 0.2, show.legend = FALSE) +
    scale_fill_manual(values= P120) +
    scale_y_continuous(labels= scales::percent) +
    scale_x_discrete(guide = NULL) +
    coord_cartesian(ylim = c(0,1.05)) +
    labs(title= "Inoculum", x= "", y= "Frequency of tags") +
    theme_gray()
  
  print(g)
}


####################
# Extracting all samples from a specific condition
####################

# Selecting condition M_Intact_27
tab = tab_tot %>%
  filter(`Temperature (°C)` == "27°C") %>%
  filter(Condition == "M_Intact") %>%
  filter(Zone == "Root")


# Graph of the bacterial load in each sample
ggplot(data= tab, aes(x= Zone, y= `Bacterial load (CFU/g)`)) +
  facet_grid( ~ `Plant ID`, scales= "free_x") +
  geom_point(colour= "black", show.legend = FALSE) +
  geom_segment(aes(x= Zone, y= 0,
                   xend= Zone, yend= `Bacterial load (CFU/g)`), colour= "black", show.legend = FALSE) +
  scale_x_discrete(guide= NULL) +
  scale_y_continuous(trans = log10_trans(), limits=c(10^2, 1.5*10^9)) +
  labs(x= "", y= "") +
  labs(x= "", y= "Load (CFU/g)") +
  theme_lucid() +
  theme(strip.background= element_blank(), strip.text= element_blank(), panel.spacing= unit(0, "lines"))

# ordering tags by frequency
tab_temp = tab

for (i in c(1:nrow(tab_temp))) {
  tab_temp[i,56:175] = tab_temp[i,56:175][,rev(order(tab_temp[i,56:175]))]
}

# ordering stackplot by Nb value
tab_temp$`Filtered bottleneck Nb`[tab_temp$Amplification == FALSE] = -1
tab_temp = tab_temp[rev(order(tab_temp$`Filtered bottleneck Nb`)),]
tab_temp$`Order ID` = factor(tab_temp$`Plant ID`, levels = tab_temp$`Plant ID`)
tab_temp = pivot_longer(tab_temp, cols = c(56:175))
tab_temp$name = factor(tab_temp$name)
tab_temp$value[tab_temp$value <= Noise_threshold] = 0

tab = pivot_longer(tab, cols = c(56:175))
tab$name = factor(tab$name)
tab$value[tab$value <= Noise_threshold] = 0


# Graph diversity per sample
ggplot(data= tab, aes(x= Zone, y= value, fill = name)) +
  facet_grid( ~ `Plant ID`, scales= "free_x") +
  geom_col(position="fill", colour= "black", size= 0.2, show.legend = FALSE) +
  #geom_text(aes(Zone, label= paste("Nr=",round(`Corrected bottleneck Nr`)), fill= NULL),
  #          stat = "summary", fun = NULL, vjust = -0.6, position = "fill") +
  #geom_text(aes(Zone, label= round(`Nr/Nb ratio`,3), fill= NULL),
  #          stat = "summary", fun = NULL, vjust = -0.6, position = "fill") +
  stat(aes(Zone, label= `Corrected bottleneck Nr`)) +
  scale_fill_manual(values= P120) +
  scale_y_continuous(labels= scales::percent) +
  scale_x_discrete(guide= NULL) +
  coord_cartesian(ylim = c(0,1.05)) +
  labs(x= "", y= "") +
  labs(x= "", y= "Frequency of tags") +
  theme_pubr() +
  theme(strip.background= element_blank(), strip.text= element_blank(), panel.spacing= unit(0, "lines"))


# Graph diversity ordonated by frequency and Nb
ggplot(data= tab_temp, aes(x= Zone, y= value, fill = name)) +
  facet_grid( ~ `Order ID`, scales= "free_x") +
  geom_col(position="fill", colour= "black", size= 0.2, show.legend = FALSE) +
  geom_text(aes(Zone, label= paste("",round(`Filtered bottleneck Nb`)), fill= NULL),
            stat = "summary", fun = NULL, vjust = -0.6, position = "fill") +
  stat(aes(Zone, label= `Corrected bottleneck Nr`)) +
  scale_y_continuous(labels= scales::percent) +
  scale_x_discrete(guide= NULL) +
  coord_cartesian(ylim = c(0,1.05)) +
  labs(x= "", y= "") +
  labs(x= "", y= "Sorted tags frequencies") +
  theme_pubr() +
  theme(strip.background= element_blank(), strip.text= element_blank(), panel.spacing= unit(0, "lines"))


####################
# Diversity in inocula from tab_kinetic
####################

tab = tab_inoc %>%
  filter(`Inoculum ID 3` == 1) %>%
  filter((`Inoculum ID 2` %in% c("S27"))) %>%
  filter(!is.na(`PCR2 validation on gel`))

tab$`Inoculum ID 2` = factor(tab$`Inoculum ID 2`, levels= c("S27"))

tab = pivot_longer(tab, cols = c(147:266))
tab$name = factor(tab$name)

# Prefiltering using noise threshold
tab$value[tab$value <= Noise_threshold] = 0


# Graph
ggplot(data= tab, aes(x= `Inoculum ID 2`, y= value, fill = name)) +
  facet_grid( ~ `Inoculum ID 2`, scales= "free_x") +
  geom_col(position="fill", colour= "black", size= 0.2, show.legend = FALSE) +
  scale_fill_manual(values= P120) +
  scale_y_continuous(labels= scales::percent) +
  scale_x_discrete(guide = NULL) +
  coord_cartesian(ylim = c(0,1.05)) +
  labs(title= "Inoculum", x= "", y= "Frequency of tags") +
  theme_pubr()


####################
# Extracting example samples for visualisation in tab_kinetic (all valid root samples)
####################

tab = tab_kinetic %>%
  filter(Genotype == "M") %>%
  filter(`Root condition` == "Intact") %>%
  filter(`Temperature (°C)` == "27") %>%
  filter(Zone == "Root") %>%
  filter(Amplification == TRUE)  %>%
  filter(`Time post-inoculation (d)` %in% c(3,4,5,6) )

tab = pivot_longer(tab, cols = c(56:175))
tab$name = factor(tab$name)

# Graph
ggplot(data= tab, aes(x= Zone, y= value, fill = name)) +
  facet_grid(~`Sample ID`, scales= "free_x") +
  geom_col(position="fill", colour= "black", size= 0.2, show.legend = FALSE) +
  #geom_text(aes(Zone, label= paste("Nr=",round(`Corrected bottleneck Nr`)), fill= NULL),
  #          stat = "summary", fun = NULL, vjust = -0.6, position = "fill") +
  #geom_text(aes(Zone, label= round(`Nr/Nb ratio`,3), fill= NULL),
  #          stat = "summary", fun = NULL, vjust = -0.6, position = "fill") +
  stat(aes(Zone, label= `Corrected bottleneck Nr`)) +
  scale_fill_manual(values= P120) +
  scale_y_continuous(labels= scales::percent) +
  scale_x_discrete(guide= NULL) +
  coord_cartesian(ylim = c(0,1.05)) +
  labs(x= "", y= "") +
  labs(x= "", y= "Frequency of tags") +
  theme_pubr() +
  theme(strip.background= element_blank(), strip.text= element_blank(), panel.spacing= unit(0, "lines"))


####################
# Extracting individual samples with stackplot and frequency graph
####################

tab = tab_tot %>%
  filter(Zone == "Root") %>%
  filter(`Plant ID` %in% c(40,34,39))

tab$`Plant ID` = factor(tab$`Plant ID`, levels= c(40,34,39))


# ordering by frequency value
tab_temp = tab

for (i in c(1:nrow(tab_temp))) {
  tab_temp[i,56:175] = tab_temp[i,56:175][,rev(order(tab_temp[i,56:175]))]
}

tab_temp = pivot_longer(tab_temp, cols = c(56:175))

tab = pivot_longer(tab, cols = c(56:175))


# Prefiltering using noise threshold
tab_temp$value[tab_temp$value <= Noise_threshold] = 0
tab_temp$value = tab_temp$value / sum(tab_temp$value)

# Converting tag ID to number
tab_temp$name = as.numeric(substring(tab_temp$name, 4))

# Removing zero values for graph
tab_temp = tab_temp %>%
  filter(value > 0)


# Rounding Nb and Nr values
tab_temp$`Corrected bottleneck Nr` = round(tab_temp$`Corrected bottleneck Nr`)
tab_temp$`Filtered bottleneck Nb` = round(tab_temp$`Filtered bottleneck Nb`)


# graph raw profile
ggplot(data= tab, aes(x= Zone, y= value, fill = name)) +
  facet_grid(~`Plant ID`, scales= "free_x") +
  geom_col(position="fill", colour= "black", size= 0.2, show.legend = FALSE) +
  #geom_text(aes(Zone, label= paste("Nr=",round(`Corrected bottleneck Nr`)), fill= NULL),
  #          stat = "summary", fun = NULL, vjust = -0.6, position = "fill") +
  #geom_text(aes(Zone, label= round(`Nr/Nb ratio`,3), fill= NULL),
  #          stat = "summary", fun = NULL, vjust = -0.6, position = "fill") +
  stat(aes(Zone, label= `Corrected bottleneck Nr`)) +
  scale_fill_manual(values= P120) +
  scale_y_continuous(labels= scales::percent) +
  scale_x_discrete(guide= NULL) +
  coord_cartesian(ylim = c(0,1.05)) +
  labs(x= "", y= "Cumulated frequency") +
  theme_pubr() +
  theme(panel.spacing= unit(0, "lines"), title = element_text(size= 11), axis.text = element_text(size= 9), legend.text = element_text(size= 9))


# Graph frequency graph
ggplot(data= tab_temp, aes(x= name, y= value, colour= name)) +
  facet_grid(~`Plant ID`, scales= "free", labeller = labeller(`Plant ID` = ~paste("Plant", .x))) +
  #geom_point(shape= 1) +
  geom_vline(data= group_by(tab_temp, by= tab_temp$`Plant ID`),
             aes(xintercept= ifelse(`Filtered bottleneck Nb` < 120, `Filtered bottleneck Nb`, 130), colour= "a")) +
  geom_point(data= group_by(tab_temp, by= tab_temp$`Plant ID`),
             aes(x= name, y= ifelse(tab_temp$name <= tab_temp$`Filtered bottleneck Nb`, value, NA),
             colour= "a")) +
  geom_vline(data= group_by(tab_temp, by= tab_temp$`Plant ID`),
             aes(xintercept= ifelse(tab_temp$Richness >= tab_temp$`Corrected bottleneck Nr`,
                                    `Corrected bottleneck Nr`, 140), colour= "b")) +
  geom_point(data= group_by(tab_temp, by= tab_temp$`Plant ID`),
             aes(x= name, y= ifelse(tab_temp$name <= tab_temp$`Corrected bottleneck Nr` &
                                      tab_temp$name > tab_temp$`Filtered bottleneck Nb`, value, NA),
             colour= "b")) +
  geom_vline(data= group_by(tab_temp, by= tab_temp$`Plant ID`),
             aes(xintercept= Richness, colour= "c")) +
  geom_point(data= group_by(tab_temp, by= tab_temp$`Plant ID`),
             aes(x= name, y= ifelse(tab_temp$name > tab_temp$`Corrected bottleneck Nr`, value, NA),
             colour= "c"),
             shape= 1) +
  scale_x_continuous(limits= c(1,NA), n.breaks= 6) +
  scale_y_continuous(trans= "log10", limits= c(10^-3.9,1)) +
  scale_colour_manual(values= c("chartreuse3", "red1", "grey35"), name= "",
                      labels= c("Tags explained \nin the STAMP model", "Tags explained \nin the STAMPR model", "Unexplained tags")) +
  labs(x= "Ordonated tags", y= "Frequency") +
  theme_pubr() +
  theme(panel.spacing= unit(1.3, "lines"), legend.position = "bottom", title = element_text(size= 11), axis.text = element_text(size= 9), legend.text = element_text(size= 9))
  


```

Exploratory tests on STAMP tags distributions

```{r Calculation}

####################
# Examination of tags distribution in some samples
####################

tab = tab_tot %>%
  filter(`Plant ID` %in% c(1:20)) %>%
  filter(Zone == "Root")


# Ordering tags by frequency
tab_tags = tab[,56:175]

for (i in c(1:nrow(tab_tags))) {
  tab_tags[i,] = sort(tab_tags[i,], decreasing = TRUE, na.last = TRUE)
  tab_tags[i,][tab_tags[i,] <= 25] = NA
}

tab_tags = tab_tags / rowSums(tab_tags, na.rm = TRUE)
tab_tags = data.frame(t(tab_tags))
colnames(tab_tags) = tab$`Plant ID`
tab_tags = cbind(tab_tags, Order= c(1:120))
tab_tags = pivot_longer(tab_tags, cols= c(1:(length(tab_tags)-1)))
tab_tags$name = factor(tab_tags$name)


# Graph of the frequency distribution of the tags
ggplot(data= tab_tags, aes(x= value)) +
  facet_wrap( ~ name, scales= "free") +
  geom_density() +
  geom_vline(xintercept= 1/119, colour= "red") +
  labs(title = "Frequency distribution", x= "Abundancy levels", y= "Number of tags") +
  theme_gray()


# Producing an ordered sum of the tags by frequency
tab_temp = tab[,56:175]

for (i in c(1:nrow(tab_temp))) {
  tab_temp[i,] = sort(tab_temp[i,], decreasing = FALSE, na.last = TRUE)
  tab_temp[i,][tab_temp[i,] <= 25] = 0
}

tab_temp = tab_temp / rowSums(tab_temp, na.rm = TRUE)

for (i in c(2:length(tab_temp))) {
  tab_temp[,i] = tab_temp[,i] + tab_temp[,i-1]
}

tab_temp = data.frame(t(tab_temp))
colnames(tab_temp) = tab$`Plant ID`
tab_temp = cbind(tab_temp, Order= c(1:120))
tab_temp = pivot_longer(tab_temp, cols= c(1:(length(tab_temp)-1)))
tab_temp$name = factor(tab_temp$name)


# Graph of the cumulative frequency distribution of the tags
ggplot(data= tab_temp, aes(x= Order, y= value)) +
  facet_wrap( ~ name, scales= "free") +
  geom_col() +
  labs(title = "Cumulative frequency graph", x= "Cumulative ordered tags", y= "Cumulated relative abundancy (%)") +
  theme_gray()


####################
# Testing a normalised comparison of tags between all samples
####################

tab = tab_tot
tab_final = data.frame()

# Scanning all conditions
for (i in levels(tab$Condition4)) {
  tab_temp = tab %>%
    filter(Condition4 == i) %>%
    filter(Amplification == TRUE)
  
  # Prefiltering using noise threshold
  tab_temp[56:175][tab_temp[56:175] <= Noise_threshold] = 0
  
  # Normalising
  if (nrow(tab_temp) > 0) {
    for (j in c(1:nrow(tab_temp))) {
  
      # Normalising tag abundancy by depth of sequencing
      tab_temp[j,56:175] = tab_temp[j,56:175] / sum(tab_temp[j,56:175])
      
      # Normalising tag abundancy by weighted sum of present tag in a sample
      tab_temp[j,56:175] = tab_temp[j,56:175] * length(tab_temp[j,56:175][tab_temp[j,56:175] > 0])
      }
    }

  # Relative sum of all tags
  
  tab_temp = as.data.frame(t(colSums(tab_temp[,56:175])))
  tab_temp = pivot_longer(tab_temp, cols = c(1:120))
  tab_temp$name = factor(tab_temp$name)
  tab_temp$Condition = i
  
  tab_final = rbind(tab_final, tab_temp)
  }

tab_final$Condition = factor(tab_final$Condition, levels= levels(tab$Condition4))


# Graph
ggplot(data= tab_final, aes(x= Condition, y= value, fill = name)) +
  facet_grid( ~ Condition, scales= "free_x") +
  geom_col(position="fill", colour= "black", size= 0.2, show.legend = FALSE) +
  scale_fill_manual(values= P120) +
  scale_y_continuous(labels= scales::percent) +
  scale_x_discrete(guide = NULL) +
  coord_cartesian(ylim = c(0,1.05)) +
  labs(title= "Normalised sum of samples", x= "", y= "Frequency of tags") +
  theme_gray() +
  theme(strip.text= element_text(angle= 90))

```

## Technical validation of the sequencing protocol

Estimation of the technical sequencing noise in the experimental protocol

- Individual culture of three exogenous tags (tag121, tag122 and tag123) were amplified and sequenced following the same protocol of plant samples and in parallel to plant samples.
- The postulate is that all tags in the range 1 to 120 present in these samples are contamination from "barcode hopping" produced during the preparation of the library.
- The baseline of sequencing noise is estimated by measuring the presence of the 120 biological tags contaminating samples with an exogenous tag.

```{r Graph}

####################
# barcode hopping noise analysis
####################

tab = tab_noise_estimation[,21:140]


# Calculating quantiles of noise tags averaged over the three control samples
Noise_quantiles = c(rowMeans(cbind(quantile(tab[1,], 0.5), quantile(tab[2,], 0.5), quantile(tab[3,], 0.5))),
                    rowMeans(cbind(quantile(tab[1,], 0.75), quantile(tab[2,], 0.75), quantile(tab[3,], 0.75))),
                    rowMeans(cbind(quantile(tab[1,], 0.95), quantile(tab[2,], 0.95), quantile(tab[3,], 0.95))),
                    rowMeans(cbind(quantile(tab[1,], 0.99), quantile(tab[2,], 0.99), quantile(tab[3,], 0.99))))

tab = rbind(tab, colMeans(tab))
tab = data.frame(t(tab))
colnames(tab) = c("V1", "V2", "V3", "Mean")

# Graph v1
ggplot(data= tab) +
  geom_histogram(aes(x= Mean), binwidth = 1, fill= "lightblue", colour= "darkgray") +
  geom_vline(xintercept= Noise_quantiles) +
  scale_x_continuous(limits= c(0,30), breaks= c(0,5,10,15,20,25,30)) +
  labs(x= "Noise read counts", y= "Tags")

```

Estimation of the limit of detection of the STAMP approach using inoculum data

- A highly stringent bottleneck will strongly affect artificial diversity and can be easily detected with a low number of tags. At the contrary, a less stringent bottleneck will need more tags to be detected because of a less important loss of diversity.
- Given this behaviour, the high limit of detection of Nb can be assessed by considering a noise-to-signal problem. An aberrant Nb will correspond to cases when sequencing noise (a difference in tag counts between the final population and the inoculum caused by barcode hopping) will be greater than signal (a difference in tag counts between the final population and the inoculum provoked by a biological bottleneck).
- As inocula are systematically sequenced in two technical replicate, we can give an estimation of high limit of detection of Nb by calculating Nb values that are produced simply by noise differences between two technical replicate. 

```{r Calculation}

####################
# Estimation of the high limit of detection of Nb
####################

tab = tab_inoc %>%
  filter(!is.na(`Initial nanopore count`))

# Retrieving the pairs of technical replicates of sequencing of inocula
tab1 = tab[c(1:nrow(tab)%%2) == 1,][,21:140]
tab2 = tab[c(1:nrow(tab)%%2) == 0,][,21:140]

# Removing non-analysable tags that are under the noise prefiltering threshold value in root
tab1 = tab1[colSums(tab2) > Noise_threshold]
tab2 = tab2[colSums(tab2) > Noise_threshold]


# Calculation of STAMP bottleneck from first replicate to second replicate
fi0 = na.omit(tab1 / rowSums(tab1))
fis = na.omit(tab2 / rowSums(tab2))
sigma <- ((fis - fi0)^2) / (fi0*(1-fi0))
F <- rowMeans(na.omit(sigma))
Nb_noise1 <- 1 / (F - 1/rowSums(tab2) - 1/rowSums(tab1))

# Calculation of STAMP bottleneck from second replicate to first replicate
fi0 = na.omit(tab2 / rowSums(tab2))
fis = na.omit(tab1 / rowSums(tab1))
sigma <- ((fis - fi0)^2) / (fi0*(1-fi0))
F <- rowMeans(na.omit(sigma))
Nb_noise2 <- 1 / (F - 1/rowSums(tab2) - 1/rowSums(tab1))


print(Nb_noise1)
print(Nb_noise2)

```

## Technical validation of the 120 strains STAMP collection

Visualisation of the in vitro growth curves for each strain

```{r Graph}

####################
# In vitro growth curves for the 120 STAMP tag strains
####################

tab = pivot_longer(tab_iv_growth, cols = c(2:121))

# Removing lag phase with an empirical 0.2h threshold and stationnary phase with an empirical 5.5h threshold
tab = tab %>%
  filter(`Time [h]` < 5.5 & `Time [h]` > 0.2)


# Graph growth curves
ggplot(data= tab, aes(x= `Time [h]`, y= log10(value), colour= name)) +
  geom_point() +
  labs(x= "Growth time (h)", y= "log10(OD)") +
  theme_gray()


# Graph growth curves
ggplot(data= tab, aes(x= `Time [h]`, y= log10(value), colour= name)) +
  geom_smooth(method= "lm", formula= y~x, se= FALSE, show.legend= FALSE) +
  labs(x= "Growth time (h)", y= "log10(OD)") +
  theme_gray()


# Formating data for linear regression
tab_final = data.frame(Tag= factor(), r= numeric(), r_std_error= numeric(), Adj.r.squared= numeric())

for (i in unique(tab$name)) {
  
  tab_temp = tab %>%
    filter(name == i)
  
  a = summary(lm(data= tab_temp, log(value) ~ `Time [h]`))
  tab_final = rbind(tab_final, list(i, a$coefficients[2], a$coefficients[4], a$adj.r.squared))
}

names(tab_final) = c("Tag", "r", "r_std_error", "Adj.r.squared")


# Graph distribution of the growth rates
ggplot(data= tab_final, aes(x= Tag, y= r)) +
  geom_pointrange(aes(ymax= (r + r_std_error), ymin= (r - r_std_error)), size= 0.2, alpha= 1) +
  scale_x_discrete(name = c(), guide= guide_axis(angle = 90)) +
  labs(x= "Tagged strain", y= "Growth rate (h-1)") +
  theme(axis.text.x= element_text(size=8))


# Graph violin plot of growth rates for the 120 tags
ggplot(data= tab_final, aes(x = "", y= r)) +
  geom_violin() +
  geom_point(alpha= 1) +
  stat_boxplot(aes(ymin = after_stat(lower), ymax = after_stat(upper)),
               geom="errorbar", width=.25) + 
  stat_boxplot(aes(ymin = after_stat(middle), ymax = after_stat(middle)),
               geom="errorbar", linewidth=1, width=.25) +
  scale_x_discrete(guide= guide_axis(angle = 90)) +
  labs(x= "All 120 strains", y= "Growth rate (h-1)") +
  theme(axis.text.x= element_text(size=8))


# Shapiro-Wilk test for normality
shapiro.test(tab_final$r)

# Graph quantile-quantile plot
qqnorm(tab_final$r)
qqline(tab_final$r)

```

Visualisation of the fitness of each tagged strain in co-culture conditions (tab_inoc)

```{r Graph}

####################
# Distribution of each tags in all inocula data (tab_inoc)
####################

tab = tab_inoc %>%
  filter(`Inoculum ID 3` == 1) %>%
  filter(!is.na(`Valid length count`))

# Prefiltering using noise threshold
tab[,147:266][tab[,147:266] <= Noise_threshold] = 0

# Normalising
for (i in c(1:nrow(tab))) {
    # Normalising tag abundancy by depth of sequencing
    tab[i,147:266] = tab[i,147:266] / sum(tab[i,147:266])
    }


# Wilcoxon-Mann-Whitney test on the average values of frequencies
a = as.data.frame(colSums(tab[,147:266], na.rm = TRUE)) / nrow(tab)
print(wilcox.test(a$`colSums(tab[, 147:266], na.rm = TRUE)`, mu= 1/120))


# Graph distribution of average frequency of each tags in all inocula
ggplot(data= a, aes(x= "", y= `colSums(tab[, 147:266], na.rm = TRUE)`)) +
  geom_violin() +
  geom_point(size= 0.8) +
  geom_hline(yintercept= 1/120)


# Preparing table for graph
tab = pivot_longer(tab, cols = c(147:266))
tab$name = substring(tab$name, 5,10)
tab$name = factor(tab$name)


# Graph distribution of average frequency of each tags in all inocula
ggplot(data= tab, aes(x = name, y= value)) +
  geom_point(size= 0.8) +
  stat_mean(colour= "red", size= 7, shape= 45) +
  geom_hline(yintercept = 1/120, colour= "red") +
  scale_x_discrete(guide= guide_axis(angle = 90)) +
  labs(x= "All 120 tags", y= "Relative frequency in inoculum") +
  coord_cartesian(ylim= c(0,0.035)) +
  theme(axis.text.x= element_text(size=8))

```

Visualisation of the fitness of each tagged strain in plant infections (tab_tot)

```{r Graph}

####################
# Distribution of each tags in all root samples data from tab_tot
####################

tab = tab_tot %>%
  filter(Zone == "Root") %>%
  filter(Amplification == TRUE)

# Prefiltering use noise threshold
tab[,56:175][tab[,56:175] <= Noise_threshold] = 0


# Normalising data
for (i in c(1:nrow(tab))) {
  
    # Normalising by ponderating by tag variation in inoculum
    #tab_temp = tab_inoc %>%
    #  filter(`Inoculum ID 2` == tab[i,]$`Inoculum ID`)
    
    #tab_temp[,147:266] = tab_temp[,147:266] / sum(tab_temp[,147:266])
    
    #tab[i,56:175] = tab[i,56:175] / tab_temp[,147:266]
    
    #tab[i,56:175][is.nan(unlist(tab[i,56:175]))] = 0
    
    # Normalising tag abundancy by depth of sequencing
    tab[i,56:175] = tab[i,56:175] / sum(tab[i,56:175])
    }


# Wilcoxon-Mann-Whitney test on the average values of frequencies
a = as.data.frame(colSums(tab[,56:175], na.rm = TRUE)) / nrow(tab)
print(wilcox.test(a$`colSums(tab[, 56:175], na.rm = TRUE)`, mu= 1/120))


# Graph distribution of average frequency of each tags in samples
ggplot(data= a, aes(x= "", y= `colSums(tab[, 56:175], na.rm = TRUE)`)) +
  geom_violin() +
  geom_point(size= 0.8) +
  geom_hline(yintercept= 1/120)


tab = pivot_longer(tab, cols = c(56:175))
tab$name = factor(tab$name)


# Distribution of each tag in samples in the rage 0 to 0.1
ggplot(data= tab, aes(x = name, y= value)) +
  geom_boxplot(outliers = TRUE, outlier.size = 0.6) +
  #geom_point(size= 0.8) +
  stat_mean(colour= "red", size= 7, shape= 45) +
  geom_hline(yintercept = 1/120, colour= "red", ) +
  scale_x_discrete(guide= guide_axis(angle = 90)) +
  labs(x= "All 120 tags", y= "Frequency in all samples") +
  coord_cartesian(ylim= c(0,0.1)) +
  scale_y_continuous(breaks= c(0,0.02,0.04,0.06,0.08,0.1)) +
  theme(axis.text.x= element_text(size=8))


# Distribution of each tag in samples in the rage 0.1 to 1
ggplot(data= tab, aes(x = name, y= value)) +
  geom_boxplot(outliers = TRUE, outlier.size = 0.6) +
  scale_x_discrete(guide= guide_axis(angle = 90)) +
  labs(x= "All 120 tags", y= "Frequency in all samples") +
  coord_cartesian(ylim= c(0.1,1)) +
  scale_y_continuous(breaks= c(0.1,0.25,0.5,0.75,1)) +
  theme(axis.text.x= element_text(size=8))


# Test distribution in inocula


tab = tab_inoc %>%
  filter(`Inoculum ID 3` == 1)

tab[,147:266][tab[,147:266] <= Noise_threshold] = 0

for (i in c(1:nrow(tab))) {
  
    # Normalising tag abundancy by depth of sequencing
    tab[i,147:266] = tab[i,147:266] / sum(tab[i,147:266])
    }


# Wilcoxon-Mann-Whitney test on the average values of frequencies

a = as.data.frame(colSums(tab[,147:266], na.rm = TRUE)) / nrow(tab)

wilcox.test(a$`colSums(tab[, 147:266], na.rm = TRUE)`, mu= 1/120)

# Student t test (inadequate test that does not verify hypothesis)
#t.test(a, mu = 1/120)


ggplot(data= a, aes(x= "", y= `colSums(tab[, 147:266], na.rm = TRUE)`)) +
  
  geom_violin() +
  
  geom_point(size= 0.8) +
  
  geom_hline(yintercept= 1/120)


# Preparing samples for graph

tab = pivot_longer(tab, cols = c(147:266))


tab$name = substring(tab$name, 5,10)

tab$name = factor(tab$name)


ggplot(data= tab, aes(x = name, y= value)) +
  
  geom_point(size= 0.8) +
  
  stat_mean(colour= "red", size= 7, shape= 45) +
  
  geom_hline(yintercept = 1/120, colour= "red") +
  
  scale_x_discrete(guide= guide_axis(angle = 90)) +
  
  labs(x= "All 120 tags", y= "Relative frequency in inoculum") +
  
  coord_cartesian(ylim= c(0,0.035)) +
  
  theme(axis.text.x= element_text(size=8))


```


# Section of the newly developped model testing approach in Python (run this prior to any python scripts)

Launch this at the beginning of the session to load data

N.B.: don't forget to set up your Python environment in Rstudio.

```{python}

import numpy as np
import matplotlib.pyplot as plt
import csv
import os
import time
from matplotlib.animation import FuncAnimation
from matplotlib import pylab
import math
import pickle
import multiprocessing as mp
from scipy.stats import gamma
import sympy as sp
from scipy.stats import t
from scipy.optimize import minimize_scalar
import matplotlib.cm as cm
from scipy.stats import spearmanr


# Info retrieved from document "Analysis_output/tab_tot.csv" for the functions "read_file"
Ntypes = 120 # Number of different tags
tags_range = 55 # Starting column for tags
sample_index_column = 1 # Column with sample names
configuration_column = 5 # Column that contains all tested samples conditions
time_of_sampling_column = 17 # Column with DPI of sampling
bacterial_load_column = 33
sample_weight_column = 19 #only necessary if the input is bacterial load to convert it to number of CFU
validation_column = 343 # Column to check that the sample is validated
inoculum_index_column = 214 # Index of the inoculum that was used for the infection
q_values = np.linspace(0, 8, 100)  # Range of q-values for Hill numbers

# Configurations
lconfigurations = [
    ['M_Intact_27_Root'],
    ['M_Cut_27_Root'],
    ['H_Intact_27_Root'],
    ['H_Cut_27_Root'],
    ['M_Intact_32_Root'],
    ['M_Cut_32_Root'],
    ['H_Intact_32_Root'],
    ['H_Cut_32_Root'],
    ]

```


# I. Evalutation of the Hill diversity in the data

To capture the diversity and distribution patterns of proportions, we use **Hill numbers**. Hill numbers provide a range of diversity measures depending on the parameter $q$. For a set of proportions $p_i$, the Hill number of order $q$ is given by:

$$
{}^{q}D = \left( \sum_{i=1}^{N} p_i^q \right)^{\frac{1}{1 - q}}
$$

Classical cases:
- For $q = 0$: Hill number represents species richness (number of types).
- For $q = 1$: Hill number corresponds to the exponential of Shannon entropy, capturing diversity while accounting for species abundance.
- For $q = 2$: Hill number is related to the inverse Simpson index, giving more weight to common species.

## Average Hill Curves

```{python, fig.width=10, fig.height=6}
# @title

file_path = "Analysis_output/tab_tot.csv"  # adjust if needed

if not os.path.exists(file_path):
    print(f"File '{file_path}' not found! Please check the path.")
else:
    with open(file_path, newline='') as file:
        data = list(csv.reader(file, delimiter=','))
        print(f"Successfully loaded '{file_path}' with {len(data)} rows.")


# Configurations for root and stem
lconfigurations_root = [
    ['M_Intact_27_Root'],
    ['M_Cut_27_Root'],
    ['H_Intact_27_Root'],
    ['H_Cut_27_Root'],
    ['M_Intact_32_Root'],
    ['M_Cut_32_Root'],
    ['H_Intact_32_Root'],
    ['H_Cut_32_Root'],
    ]

# Define colors for different experiments
colors = ['red', 'blue', 'black', 'cyan', 'orange', 'green', 'magenta', 'navy']


# Helper functions

def compute_hill_number(p, q):
    if q == 1:
        # Special case for q=1 (Shannon entropy)
        return np.exp(-np.sum(p * np.log(p)))
    elif q == 0:
        # Special case for q=0 (species richness)
        return np.sum(p > 0)
    else:
        # General case for q != 1
        return np.power(np.sum(np.power(p, q)), 1 / (1 - q))

def compute_hill_curve(X, q_values, Tags_range):
    # Normalize the counts to probabilities
    p = X / np.sum(X)
    return [compute_hill_number(p, q) for q in q_values]

def read_file(data, configuration, tags_range, configuration_column, validation_column, sample_index_column, time_of_sampling_column, bacterial_load_column, Ntypes):
    list_alpha = []
    conditions = []
    for row in data:
        if row[configuration_column] == configuration[0]: # Check condition
            if row[validation_column] == '1': # Check sequencing validation
              
              list_alpha.append([int(row[j]) for j in range(tags_range, Ntypes+tags_range)])
              conditions.append((row[sample_index_column], int(row[time_of_sampling_column]), float(row[bacterial_load_column])))

    return np.array(list_alpha), np.array(conditions)


# Main analysis

plt.figure(figsize=(10, 6))

# Iterate over root configurations and plot all on the same graph
for i, (config_root) in enumerate(zip(lconfigurations_root)):
    # Read data for root configurations
    list_alpha_root, _ = read_file(data, config_root[0], tags_range, configuration_column, validation_column, sample_index_column, time_of_sampling_column, bacterial_load_column, Ntypes)

    if len(list_alpha_root) == 0:
        print(f"No data found for configuration: {config_root}")
        continue

    # Compute Hill numbers for each replicate
    hill_curves_root = [compute_hill_curve(alpha, q_values, tags_range) for alpha in list_alpha_root]

    # Average Hill numbers over replicates
    hill_curve_mean_root = np.mean(hill_curves_root, axis=0)
    
    # Plot the mean Hill number curves for root (plain line)
    plt.plot(q_values, hill_curve_mean_root, color=colors[i % len(colors)], lw=2,
             label=f"{config_root[0]}")

# Increase font sizes
plt.xlabel("$q$", fontsize=16)
plt.ylabel("Mean Hill numbers, ${}^{q}H(root)$", fontsize=16)
plt.legend(fontsize=14)
plt.xticks(fontsize=16)
plt.yticks(fontsize=16)
plt.xlim(0,6)
plt.grid()
plt.tight_layout()
#plt.savefig("mean_hill_numbers_root.png", dpi=300, bbox_inches='tight')
plt.show()

```

# II. Simulation of the mechanistic models CR and IR

## II.1 Simulation of CR model

```{python}
# @title

T = 10
mu = 1
Ntypes = 120
r = 1

# Simulate arrival times tau
tau_list = []
current_time = 0.0
while True:
    inter_arrival = np.random.exponential(scale=1/mu)
    current_time += inter_arrival
    if current_time > T:
        break
    tau_list.append(current_time)

tau = np.array(tau_list)
n = len(tau)

# Assign each arrival a random type between 1 and Ntypes
types = np.random.randint(1, Ntypes+1, size=n)

# Group arrival times by type
tau_by_type = [tau[types == i] for i in range(1, Ntypes+1)]

# Define a function to compute f(t) = [f_1(t), ..., f_Ntypes(t)]
def f_of_t(t):
    f_values = np.zeros(Ntypes)
    for i in range(Ntypes):
        arrivals_i = tau_by_type[i]
        valid_arrivals = arrivals_i[arrivals_i <= t]
        f_values[i] = np.sum(np.exp(r * (t - valid_arrivals)))
    return f_values

# Choose time points for plotting
t_points = np.linspace(0, T, 200)  # 200 points between 0 and T
f_matrix = np.array([f_of_t(t) for t in t_points])  # shape = (200, Ntypes)

# Sort the processes by their final values at the last time point
final_values = f_matrix[-1]            # final values at t = T
order = np.argsort(final_values)       # indices that sort the final values

# Reorder f_matrix and labels according to the sorted order
f_matrix_sorted = f_matrix[:, order]
labels_sorted = [f"Type {i+1}" for i in order]

# Create a stacked area plot with sorted order
plt.figure(figsize=(10,6))
plt.stackplot(t_points, f_matrix_sorted.T, labels=labels_sorted)

plt.xlabel("Time t")
plt.ylabel("f_i(t)")
plt.title("Stacked area plot of all f_i(t), sorted by final value")

# With 120 types, the legend might be large. Omit or show selectively.
# plt.legend(loc='upper left')

plt.show()

```

## II.2 Simulation of IR model

```{python}
# @title

# NHPP inverse cumulative intensity function
def nhpp_exponential_inverse(s, mu, k):
    return (1.0 / k) * np.log(k * s / mu + 1)

# Simulation des temps d'arrivée NHPP
def simulate_nhpp(mu, k, T):
    tau = []
    s = 0.0
    while True:
        s += np.random.exponential(1.0)
        t_candidate = nhpp_exponential_inverse(s, mu, k)
        if t_candidate > T:
            break
        tau.append(t_candidate)
    return np.array(tau)

# Générer les fonctions f_i(t)
def compute_f_matrix(tau, types, Ntypes, r, t_points):
    tau_by_type = [tau[types == i] for i in range(Ntypes)]
    f_matrix = []

    for t in t_points:
        f_t = np.zeros(Ntypes)
        for i in range(Ntypes):
            arrivals = tau_by_type[i]
            arrivals_t = arrivals[arrivals <= t]
            f_t[i] = np.sum(np.exp(r * (t - arrivals_t)))
        f_matrix.append(f_t)

    return np.array(f_matrix)

# Paramètres
mu = 0.1
k = 0.78
T = 10
Ntypes = 120
r = 1.0
t_points = np.linspace(0, T, 200)

# Étapes
tau = simulate_nhpp(mu, k, T)
n = len(tau)
types = np.random.randint(0, Ntypes, size=n)
f_matrix = compute_f_matrix(tau, types, Ntypes, r, t_points)

# Tri des types selon la valeur finale
final_values = f_matrix[-1]
order = np.argsort(final_values)
f_matrix_sorted = f_matrix[:, order]
labels_sorted = [f"Type {i}" for i in order]

# Visualisation
plt.figure(figsize=(10, 6))
plt.stackplot(t_points, f_matrix_sorted.T, labels=labels_sorted)
plt.xlabel("Time t")
plt.ylabel("f_i(t)")
plt.title("Stacked area plot des f_i(t) - modèle NHPP exponentiel")
# plt.legend(loc='upper left')  # désactivé par défaut
plt.show()

```

# III. Parameter estimation based on frequencies (do not relaunch unless necessary; outputs are already saved)

```{python, fig.width=8, fig.height=6}
# @title Main functions common to all models. Launch before the estimation

def sample_theta(K, theta_range, distribution_model):
    theta = np.random.uniform(*theta_range)
    V = distribution_model(K, theta)
    return V / np.sum(V)

def quantile_distance(theta_range, frequencies, distribution_model, parameter_name="theta", Nsim=100, quantile=0.05):
    K = len(frequencies)
    Q = np.arange(0, 4.1, 0.1)

    def hill_number(V, q):
        if q == 1:
            return np.exp(-np.sum(V[V > 0] * np.log(V[V > 0])))
        elif q == 0:
            return np.sum(V > 0)
        else:
            return (np.sum(V ** q)) ** (1 / (1 - q))

    def calculate_distance(V1, V2):
        DS = 0
        for q in Q:
            Hq1 = hill_number(V1, q)
            Hq2 = hill_number(V2, q)
            DS += abs(Hq1 - Hq2) / (Hq1 + Hq2)
        return DS / len(Q)

    distances = []
    for _ in range(Nsim):
        V = sample_theta(K, theta_range, distribution_model)
        D = calculate_distance(V, frequencies)
        distances.append(D)

    return np.quantile(distances, quantile)

def find_best_theta(frequencies, theta_range, distribution_model, distance_function, parameter_name="theta", step=1):
    distances = []
    theta_values = np.arange(theta_range[0], theta_range[1] + step, step)

    for theta in theta_values:
        dist = distance_function((theta, theta), frequencies, distribution_model, parameter_name)
        distances.append(dist)

    best_theta = theta_values[np.argmin(distances)]
    min_distance = min(distances)
    return best_theta, min_distance
  

def read_file(data, configuration, tags_range, configuration_column, validation_column, sample_index_column, time_of_sampling_column, bacterial_load_column, Ntypes):
    frequencies = []
    conditions = []
    for row in data:
        if row[configuration_column] == configuration[0]: # Check condition
            if row[validation_column] == '1': # Check sequencing validation
                counts = [int(row[j]) for j in range(tags_range, Ntypes + tags_range)]
                counts_sum = sum(counts)
                if counts_sum > 0:
                    frequencies.append([c / counts_sum for c in counts])
                    time = float(row[17])
                    conditions.append((row[sample_index_column], int(row[time_of_sampling_column]), float(row[bacterial_load_column])))

    return np.array(frequencies), np.array(conditions)

def process_configurations(data, lconfigurations, distribution_model, theta_range, step, distance_function=quantile_distance, parameter_name="theta", model_name="Adaptable Frequency Model"):
    theta_values = {}  # Dictionary to store theta lists, mean values, and performance
    results = []

    for configuration in lconfigurations:
        frequencies, _ = read_file(data, configuration, tags_range, configuration_column, validation_column, sample_index_column, time_of_sampling_column, bacterial_load_column, Ntypes)
        if len(frequencies) == 0:
            print(f"No data for configuration: {configuration}")
            continue

        mle_theta = []
        dist_s = []
        for freq in frequencies:
            best_theta, dist_ = find_best_theta(freq, theta_range, distribution_model, distance_function, parameter_name, step)
            mle_theta.append(best_theta)
            dist_s.append(dist_)

        # Compute mean theta and mean distance across all replicates
        mean_mle_theta = np.mean(mle_theta)
        mean_distance = np.mean(dist_s)  # Performance metric

        # Store both all best theta values, mean theta, and mean performance
        theta_values[tuple(configuration)] = {
            "theta_list": mle_theta,  # List of all best theta values
            "mean_theta": mean_mle_theta,  # Mean of best theta values
            "mean_distance": mean_distance  # Mean distance (performance metric)
        }

        # Store and print results (only mean values)
        results.append((configuration, mean_mle_theta, mean_distance))
        print(f"Modèle: {model_name}, Configuration: {configuration}, Mean theta: {mean_mle_theta:.2f}, Performance (Mean Distance): {mean_distance:.4f}")

    return theta_values  # Return both detailed and summarized theta values
  

with open('Analysis_output/tab_tot.csv', newline='') as file:
    data = list(csv.reader(file, delimiter=','))

# Functions to plot the Hill curves
def hill_number(V, q):
    """Compute Hill numbers for a given frequency vector V and order q."""
    if q == 1:
        return np.exp(-np.sum(V[V > 0] * np.log(V[V > 0])))
    elif q == 0:
        return np.sum(V > 0)
    else:
        return (np.sum(V ** q)) ** (1 / (1 - q))

def calculate_distance_per_q(V1, V2, Q):
    """Compute the diversity-based distance for each q."""
    distances = []
    for q in Q:
        Hq1 = hill_number(V1, q)
        Hq2 = hill_number(V2, q)
        distance = abs(Hq1 - Hq2) / (Hq1 + Hq2) if (Hq1 + Hq2) != 0 else 0
        distances.append(distance)
    return distances

def generate_best_frequency(K, theta, distribution_model, real_frequency, num_sim=100):
    """Generate 100 frequency vectors and return the closest one to the real frequency."""
    simulated_vectors = [sample_theta(K, (theta, theta), distribution_model) for _ in range(num_sim)]
    distances = [np.mean(calculate_distance_per_q(V, real_frequency, np.arange(0, 4.1, 0.1))) for V in simulated_vectors]
    best_index = np.argmin(distances)  # Index of lowest distance
    return simulated_vectors[best_index]  # Return the best-matching frequency

def plot_hill_curves(real_frequencies, best_simulated_frequencies):
    """Plot Hill curves for real and best simulated frequencies in separate subplots."""
    Q = np.arange(0, 4.1, 0.1)
    num_freqs = len(real_frequencies)
    color_map = plt.colormaps["tab10"]

    fig, axes = plt.subplots(1, 2, figsize=(12, 6), sharey=True)

    # Plot each real frequency in the left subplot
    for i, freq in enumerate(real_frequencies):
        hill_curve = [hill_number(freq, q) for q in Q]
        color = color_map(i / max(1, num_freqs - 1))
        axes[0].plot(Q, hill_curve, color=color, alpha=0.9, linewidth=2)

    # Plot each best simulated frequency in the right subplot
    for i, best_freq in enumerate(best_simulated_frequencies):
        hill_curve = [hill_number(best_freq, q) for q in Q]
        color = color_map(i / max(1, num_freqs - 1))
        axes[1].plot(Q, hill_curve, color=color, alpha=0.9, linewidth=2)

    # Formatting subplots
    axes[0].set_title("Observed Frequencies")
    axes[1].set_title("Best Simulated Frequencies")
    for ax in axes:
        ax.set_xlabel("q")
        ax.set_ylabel("Hill Number")
        ax.set_ylim(0, 120)

    plt.tight_layout()
    plt.show()

def process_with_plots(data, lconfigurations, distribution_model, theta_values):
    """Process configurations, generate the best frequency vectors, and plot Hill curves and global mean distance."""
    Q = np.arange(0, 4.1, 0.1)
    global_mean_distances = np.zeros(len(Q))  # Accumulate mean distances over all configurations
    total_samples = 0  # Count total frequencies processed
    all_configurations_mean_distances = [] # Mean distances for all individual configuration
    configuration_mean_distances = np.zeros(len(Q))  # Accumulate mean distances within each configuration
    configuration_samples = 0 # Count frequencies processed within each configuration

    all_real_frequencies = {}  # Store real frequencies for all configurations
    all_best_simulated_frequencies = {}  # Store best simulated frequencies for all configurations

    for configuration in lconfigurations:
        real_frequencies, _ = read_file(data, configuration, tags_range, configuration_column, validation_column, sample_index_column, time_of_sampling_column, bacterial_load_column, Ntypes)
        if len(real_frequencies) == 0:
            print(f"No data for configuration: {configuration}")
            continue

        K = len(real_frequencies[0])  # Number of species/types
        config_key = tuple(configuration)
        theta_list = theta_values.get(config_key, {}).get("theta_list", None)

        if theta_list is None or len(theta_list) != len(real_frequencies):
            print(f"Missing or mismatched theta values for configuration: {configuration}")
            continue

        best_simulated_frequencies = []
        for real_freq, theta in zip(real_frequencies, theta_list):
            best_freq = generate_best_frequency(K, theta, distribution_model, real_freq)
            best_simulated_frequencies.append(best_freq)

            # Compute distance for this frequency
            distances = calculate_distance_per_q(real_freq, best_freq, Q)
            global_mean_distances += np.array(distances)
            total_samples += 1
            
            # Distance retriever for the configuration
            configuration_mean_distances += np.array(distances)
            configuration_samples += 1

        # Compute and append distance for the configuration
        all_configurations_mean_distances.append(configuration_mean_distances / configuration_samples)

        # Reset configuration distance and sample count
        configuration_mean_distances = np.zeros(len(Q))
        configuration_samples = 0
        
        # Store results for this configuration
        all_real_frequencies[config_key] = real_frequencies
        all_best_simulated_frequencies[config_key] = best_simulated_frequencies

        # Plot Hill curves for this configuration
        plot_hill_curves(real_frequencies, best_simulated_frequencies)

    # Compute final averaged distances
    if total_samples > 0:
        global_mean_distances /= total_samples

    # Plot global error figure
    plt.figure(figsize=(8, 6))
    plt.plot(Q, global_mean_distances, color="black", linewidth=2)
    plt.xlabel("q")
    plt.ylabel("Mean Distance")
    plt.title("Mean Distance Averaged Over All Configurations")
    plt.show()

    return Q, global_mean_distances, all_real_frequencies, all_best_simulated_frequencies, all_configurations_mean_distances

# functions to Save and load the results
def save_pickle(theta_values, filename="theta_values1.pkl"):
    """
    Save theta_values dictionary as a Pickle file.
    """
    with open(filename, "wb") as f:
        pickle.dump(theta_values, f)
    print(f"Saved to {filename}")

def load_pickle(filename="theta_values1.pkl"):
    """
    Load theta_values dictionary from a Pickle file.
    """
    with open(filename, "rb") as f:
        return pickle.load(f)

```

## III.1 Multinomial model

**Model:** $\mathcal{M}(N_b; 1/K,\ldots,1/K)$ with $K=120$.

---

Here, we wind the sample size $N_b$ that minimizes the average distance between a simulated frequency vector (drawn from a multinomial distribution) and the data, among integer values of $N_b$ between 1 and 200.

The distance $D$ is defined as the average relative difference between the Hill indices of the two vectors over a range of $q$ values: $$ D = \frac{1}{|Q|} \sum_{q \in Q} \frac{|H_q(Z) - H_q(\hbox{ frequencies})|}{H_q(Z) + H_q(\hbox{frequencies})},$$where $H_q$ (Hill index) is calculated as:
- For $q = 1$, $H_q = \exp\left(-\sum_{i} V_i \log(V_i)\right)$ (Shannon entropy exponentiated).
- For $q \neq 1$, $H_q = \left(\sum_{i} V_i^q\right)^{\frac{1}{1-q}}$

---

**Main steps**:
- Simulates frequency distributions $N_{sim}$ times using a multinomial distribution with sample size $N_b$.
- Computes the average distance between the data and the simulated frequency distributions.
- Finds the value of $N_b$ which minimizes the average distance.


```{python}
# define the model
def distribution_modelM1(K, theta):
    Nb = theta
    return np.random.multinomial(Nb, [1 / K] * K)
```

```{python}

# define the range for parameter values
theta_range = (1, 200)
step = 1


print('Model performance')
# Run the processing function
theta_values1 = process_configurations(data, lconfigurations, distribution_modelM1, theta_range, step, quantile_distance, parameter_name="Nb", model_name = "STAMP Model")
save_pickle(theta_values1,"Models_output/theta_values1.pkl")
```

#### Hill Curves

```{python}
theta_values1 = load_pickle("Models_output/theta_values1.pkl")  # Loads it back
Q1, D1, RF, BF1, all_D1 = process_with_plots(data, lconfigurations, distribution_modelM1, theta_values1)
np.save("Models_output/Q1.npy", Q1)
np.save("Models_output/D1.npy", D1)
np.save("Models_output/all_D1.npy", all_D1)
save_pickle(RF, filename="Models_output/RF.pkl")
save_pickle(BF1, filename="Models_output/BF1.pkl")

```

## III.3 Mechanistic model CR

We may fix $r=1$ and $t=10$ here, as we only use the frequencies and not the total population sizes, we can only estimate $r/\mu$

```{python}
# define the model
def distribution_modelM3(K, theta):
    mu = theta
    Ntypes = K
    r = 1
    T = 10
    # on simule les temps d'entrée :
    tau = [ np.random.exponential( 1/mu ) ]
    while tau[-1] < T:
        tau += [ tau[-1] + np.random.exponential( 1/mu ) ]
    qt_prov = [ np.exp( r * (T-tau[i]) ) for i in range(len(tau)) ]
    types = [ np.random.randint(0, Ntypes) for i in range(len(tau))]
    qt_finale = []
    for i in range(len(tau)):
        if types[i] != -1:
            qt_cur = qt_prov[i]
            for j in range(i+1, len(tau)):
                if types[j] == types[i] and types[j] != -1:
                    qt_cur += qt_prov[j]
                    types[j] = -1
            types[i] = -1
            qt_finale += [ qt_cur ]
    if len(qt_finale) > Ntypes: # impossible, normalement
        print(len(qt_finale), "types différents sont passés")
        qt_finale = qt_finale[:Ntypes]
    else:
        qt_finale += (Ntypes - len(qt_finale)) * [0]
    res = np.array(qt_finale) #/ np.sum(np.array(qt_finale))

    return np.sort(res)[::-1]
```

```{python}
# define the range for parameter values
theta_range = (1, 50)
step = 1

print('Model performance')
# Run the processing function
theta_values3 = process_configurations(data, lconfigurations, distribution_modelM3, theta_range, step, quantile_distance, parameter_name="Nb", model_name = "CR1 Model")
save_pickle(theta_values3,"Models_output/theta_values3.pkl")
```

#### Hill Curves

```{python}
theta_values3 = load_pickle("Models_output/theta_values3.pkl")
Q3, D3, RF, BF3, all_D3 = process_with_plots(data, lconfigurations, distribution_modelM3, theta_values3)
np.save("Models_output/Q3.npy", Q3)
np.save("Models_output/D3.npy", D3)
np.save("Models_output/all_D3.npy", all_D3)
save_pickle(BF3, filename="Models_output/BF3.pkl")
```

## III.4 Bolthausen-Sznitman coalescent

To construct the Bolthausen-Sznitman coalescent, we use the procedure of Goldschmidt and Martin (*Random recursive trees and the Bolthausen-Sznitman coalescent. Electron. J. Probab. 10, 718–745*). See also section 6.1 of Berestycki, *Recent Progress in Coalescent Theory, 2009*.

### Construction of the model

```{python}
#**Step 1**: construct the initial random recursive tree.
def update_edges( l, k ):
    current = k
    while current != 0:
        current = l[current][0]
        l[current][2] += [k]


def init_bs( n ):
    l_edges = [ [0,[0],[]] ]
    new_edge = [0,[0],[]]
    for k in range(1,n):
        parent_new = np.random.randint(k)
        new_edge = [ parent_new, [k], []]
        l_edges += [ new_edge ]
        update_edges( l_edges, k )
    return l_edges
#**Step 2**: each edge is "lifted" at rate $1$ (in the terminology of *Berestycki 2009*).
def step_bs( tree, k ): # lift edge k of tree
    loss = 0
    parent = tree[k][0]
    tree[parent][1] += [k] + tree[k][2] # the edge is lifted to the parent
    for i in [k] + tree[k][2]:
        if tree[i][1][0] != -1:
            tree[i][1] = [-1] # unactivated
            loss += 1
    return tree, loss # tree : resulting tree; loss: number of blocks lost
#**Step 3:** the following function runs a Bolthausen-Sznitman coalescent for a time [t], starting from an initial tree [tree] with specified number of blocks [nblocks]).

def run_bs( nblocks, t, tree ):

    t_cur = np.random.exponential(1/nblocks)

    while t_cur < t and nblocks > 1:


        i = np.random.randint( 1, nblocks ) # choose the edge to be lifted
        edge = 0
        while i > -1:
            if tree[ edge ][1][0] != -1:
                edge += 1
                i -= 1
            else:
                edge += 1
        tree, loss = step_bs( tree, edge - 1 )
        nblocks -= loss

        t_cur += np.random.exponential(1/nblocks)

    return tree, nblocks


#run_bs(10000,0.5)
def display( tree ):
    for k in range( len(tree) ):
        if tree[k][1][0] != -1:
            print(tree[k][1], end='  ')
    print("")
def get_prop_bs( N, t ):
    tree,_ = run_bs( N, t, init_bs( N ) )
    prop = [] # get proportions
    for e in tree:
        if e[1][0] != -1:
            prop += [ len(e[1]) ]
    prop = prop / np.sum( prop )
    return prop
def sample_tags(prop, Ntags):
    prop_tags = np.zeros( Ntags )
    for p in prop:
        tag = np.random.randint( Ntags )
        prop_tags[ tag ] += p
    return prop_tags
```

```{python}
# define the model
def distribution_modelM4(K, theta):
    t = theta
    return sample_tags( get_prop_bs(2000, t), K )
```

### Optimization

```{python}
# define the range for parameter values
theta_range = (0.05, 1)
step = 0.05

print('Model performance')
# Run the processing function
theta_values4 = process_configurations(data, lconfigurations, distribution_modelM4, theta_range, step, quantile_distance, parameter_name="t", model_name = "BS Model")
save_pickle(theta_values4,"Models_output/theta_values4.pkl")
```

#### Hill Curves

```{python}
theta_values4 = load_pickle("Models_output/theta_values4.pkl")
Q4, D4, RF, BF4, all_D4 = process_with_plots(data, lconfigurations, distribution_modelM4, theta_values4)
np.save("Models_output/Q4.npy", Q4)
np.save("Models_output/D4.npy", D4)
np.save("Models_output/all_D4.npy", all_D4)
save_pickle(BF4, filename="Models_output/BF4.pkl")
```

## III.5 Kingman coalescent

The Kingman coalescent is simulated in the most natural way.

The following function runs a Kingman coalescent for time [t], where the initial blocks correspond to proportions [prop].

```{python}
def run_km( t, prop = [] ):

    nblocks = len(prop)
    t_cur = 0

    if nblocks > 1:
        t_cur = np.random.exponential(2/(nblocks * (nblocks-1)))

    while t_cur < t and nblocks > 1:
        t_cur += np.random.exponential(2/(nblocks * (nblocks-1)))

        i,j=[0,0]
        while i == j:
            i,j = np.random.randint( 0, nblocks, 2)
        prop[i] += prop[j] # j gets to i
        prop.pop(j) # remove j
        nblocks -= 1
    return prop

def sample_tags(prop, Ntags):
    prop_tags = np.zeros( Ntags )
    for p in prop:
        tag = np.random.randint( Ntags )
        prop_tags[ tag ] += p
    return prop_tags
```

```{python}
# define the model
def distribution_modelM5(K, theta):
    t = theta
    return sample_tags( run_km(t, [1/1000] * 1000), K )
```

```{python}
# define the range for parameter values
theta_range = (0.05, 1)
step = 0.05
print('Model performance')
# Run the processing function
theta_values5 = process_configurations(data, lconfigurations, distribution_modelM5, theta_range, step, quantile_distance, parameter_name="t", model_name = "K Model")
save_pickle(theta_values5,"Models_output/theta_values5.pkl")
```

#### Hill Curves

```{python}
theta_values5 = load_pickle("Models_output/theta_values5.pkl")
Q5, D5, RF, BF5, all_D5 = process_with_plots(data, lconfigurations, distribution_modelM5, theta_values5)
np.save("Models_output/Q5.npy", Q5)
np.save("Models_output/D5.npy", D5)
np.save("Models_output/all_D5.npy", all_D5)
save_pickle(BF5, filename="Models_output/BF5.pkl")
```

## III.6 Mechanistic model with variable rate (IR model)

```{python}

# NHPP inverse cumulative intensity function for λ(t) = mu * exp(k * t)
def nhpp_exponential_inverse(s, mu, k):
    return (1.0 / k) * np.log(k * s / mu + 1)

# define the model
def distribution_modelM6(K, theta):
    mu = theta
    k = 0.78
    Ntypes = K
    T = 10

    # Step 1: Simulate NHPP arrival times
    tau = []
    s = 0.0
    while True:
        s += np.random.exponential(1.0)  # Homogeneous Poisson interarrival
        t_candidate = nhpp_exponential_inverse(s, mu, k)
        if t_candidate > T:
            break
        tau.append(t_candidate)

    # Step 2: Compute provisional quantity
    qt_prov = [np.exp((T - t)) for t in tau]

    # Step 3: Assign and merge types
    types = [np.random.randint(0, Ntypes) for _ in tau]
    qt_finale = []
    for i in range(len(tau)):
        if types[i] != -1:
            qt_cur = qt_prov[i]
            for j in range(i + 1, len(tau)):
                if types[j] == types[i] and types[j] != -1:
                    qt_cur += qt_prov[j]
                    types[j] = -1
            types[i] = -1
            qt_finale.append(qt_cur)

    # Step 4: Padding or trimming
    if len(qt_finale) > Ntypes:
        print(len(qt_finale), "types différents sont passés")
        qt_finale = qt_finale[:Ntypes]
    else:
        qt_finale += (Ntypes - len(qt_finale)) * [0]

    # Return sorted descending quantities
    return np.sort(np.array(qt_finale))[::-1]

```

```{python}
# define the range for parameter values
theta_range = (0.05, 1)
step = 0.05

print('Model performance')
# Run the processing function
theta_values6 = process_configurations(data, lconfigurations, distribution_modelM6, theta_range, step, quantile_distance, parameter_name="mu", model_name = "VR Model")
save_pickle(theta_values6,"Models_output/theta_values6.pkl")
```

#### Hill Curves

```{python}
theta_values6 = load_pickle("Models_output/theta_values6.pkl")
Q6, D6, RF, BF6, all_D6 = process_with_plots(data, lconfigurations, distribution_modelM6, theta_values6)
np.save("Models_output/Q6.npy", Q6)
np.save("Models_output/D6.npy", D6)
np.save("Models_output/all_D6.npy", all_D6)
save_pickle(BF6, filename="Models_output/BF6.pkl")
```


# IV. Some figures and analysis of models results

## Barcodes: experiments vs models

```{python, fig.width=10, fig.height=6}

# =======================
# Load pickle files
# =======================

def load_pickle(filename):
    if not os.path.exists(filename):
        raise FileNotFoundError(f"'{filename}' not found in repository.")
    with open(filename, "rb") as f:
        return pickle.load(f)

# Load data
RF = load_pickle("Models_output/RF.pkl")
BF1 = load_pickle("Models_output/BF1.pkl")
BF3 = load_pickle("Models_output/BF3.pkl")
BF4 = load_pickle("Models_output/BF4.pkl")
BF6 = load_pickle("Models_output/BF6.pkl")
theta_values = load_pickle("Models_output/theta_values1.pkl")  # Adjust the file name if needed

# =======================
# Helper function
# =======================

def plot_stacked_bars(ax, frequencies_list, title=None):
    n_replicates = len(frequencies_list)
    x_positions = np.arange(n_replicates)

    for i, freqs in enumerate(frequencies_list):
        freqs_sorted = np.sort(freqs)[::-1]
        bottom = 0
        for freq in freqs_sorted:
            ax.bar(x_positions[i], freq, bottom=bottom, width=0.5, edgecolor='black')
            bottom += freq

    # Clean axes
    ax.set_xticks([])
    ax.set_yticks([])
    for spine in ax.spines.values():
        spine.set_visible(False)

    ax.set_ylim(0, 1)

    if title:
        ax.set_title(title, fontsize=10)

# =======================
# Row names (no K model)
# =======================

row_names = [
    "Observed",
    "Model $\mathcal{M}_{STAMP}$",
    "Model $\mathcal{M}_{CR}$",
    "Model $\mathcal{M}_{BS}$",
    "Model $\mathcal{M}_{IR}$",
]

# =======================
# Select experiments
# =======================

# Extract experiment names
experiment_names = list(RF.keys())

# Select experiments to process
# selected_experiments = [experiment_names[0], experiment_names[2]]
selected_experiments = [experiment_names[0]]  # <-- change if needed

# =======================
# Main loop over experiments
# =======================

for exp_name in selected_experiments:
    observed_list = RF[exp_name]
    bf1_list = BF1[exp_name]
    bf3_list = BF3[exp_name]
    bf4_list = BF4[exp_name]
    bf6_list = BF6[exp_name]

    n_replicates = len(observed_list)

    # === Sorting according to descending theta_list ===
    theta_list = theta_values[exp_name]['theta_list']
    sorted_indices = np.argsort(theta_list)[::-1]  # Indices of descending theta

    # Reorder all replicate lists based on sorted indices
    observed_list = [observed_list[i] for i in sorted_indices]
    bf1_list = [bf1_list[i] for i in sorted_indices]
    bf3_list = [bf3_list[i] for i in sorted_indices]
    bf4_list = [bf4_list[i] for i in sorted_indices]
    bf6_list = [bf6_list[i] for i in sorted_indices]

    # Create figure: 5 rows, 1 column
    fig, axes = plt.subplots(
        nrows=5, ncols=1,
        figsize=(max(6, n_replicates * 0.7), 10),
        sharex=True, sharey=True
    )

    fig.suptitle(f"Configuration: {exp_name}", fontsize=18)

    all_data = [
        observed_list,
        bf1_list,
        bf3_list,
        bf4_list,
        bf6_list,
    ]

    for row_idx, data_list in enumerate(all_data):
        ax = axes[row_idx] if n_replicates > 1 else axes
        plot_stacked_bars(ax, data_list)

        ax.set_ylabel(row_names[row_idx], fontsize=12, rotation=0, labelpad=40, ha='right')

        # Top axis: add replicate numbers
        if row_idx == 0:
            ax.set_xticks(np.arange(n_replicates))
            ax.set_xticklabels([f"Rep {i}" for i in sorted_indices], fontsize=8)
            ax.tick_params(axis='x', which='both', length=0)

    plt.tight_layout(rect=[0, 0, 1, 0.95])

    # Save the figure
    #filename = f"{exp_name}_results_compact_sorted.png"
    #fig.savefig(filename, dpi=300, bbox_inches='tight')

    # Show the figure
    plt.show()


# Extraction of the simulated barcodes frequencies from the selected experiment into .csv files
np.savetxt("Models_output/Barcodes_best_prediction_MI27/observed_list.csv",
        observed_list,
        delimiter =";",
        fmt ='% s')

np.savetxt("Models_output/Barcodes_best_prediction_MI27/STAMP_list.csv",
        bf1_list,
        delimiter =";",
        fmt ='% s')

np.savetxt("Models_output/Barcodes_best_prediction_MI27/CR_list.csv",
        bf3_list,
        delimiter =";",
        fmt ='% s')

np.savetxt("Models_output/Barcodes_best_prediction_MI27/BS_list.csv",
        bf4_list,
        delimiter =";",
        fmt ='% s')

np.savetxt("Models_output/Barcodes_best_prediction_MI27/IR_list.csv",
        bf6_list,
        delimiter =";",
        fmt ='% s')

```

## Global figure with Hill numbers for models STAMP, CR, BS, IR

```{python, fig.width=20, fig.height=18}

# Function to load pickle files
def load_pickle(filename):
    with open(filename, "rb") as f:
        return pickle.load(f)

# Load the saved data
RF = load_pickle("Models_output/RF.pkl")
BF1 = load_pickle("Models_output/BF1.pkl")
BF3 = load_pickle("Models_output/BF3.pkl")
BF4 = load_pickle("Models_output/BF4.pkl")
BF6 = load_pickle("Models_output/BF6.pkl")

# Extract experiment names from RF
experiment_names = list(RF.keys())

# Define Q values (x-axis for Hill numbers)
Q = np.arange(0, 4.1, 0.1)
#Q = np.load("Q1.npy")
# Function to compute Hill numbers
def hill_number(V, q):
    """Compute Hill numbers for a given frequency vector V and order q."""
    if q == 1:
        return np.exp(-np.sum(V[V > 0] * np.log(V[V > 0])))
    elif q == 0:
        return np.sum(V > 0)
    else:
        return (np.sum(V ** q)) ** (1 / (1 - q))

# Function to compute Hill curves for multiple frequency distributions
def compute_hill_curves(frequency_data):
    """Compute Hill curves for all frequency distributions in a configuration."""
    hill_curves = []
    for freq_vector in frequency_data:
        curve = [hill_number(freq_vector, q) for q in Q]
        hill_curves.append(curve)
    return np.array(hill_curves)

# Number of configurations and models
num_configurations = 7
num_models = 4  # Models 1, 3, 4, 6


# Create figure with subplots (7 rows, 5 columns)
fig, axes = plt.subplots(num_configurations, num_models + 1, figsize=(20, 18), sharex=True, sharey=True)

# Iterate over configurations and plot Hill curves
for i, exp_name in enumerate(experiment_names):
    # Compute Hill curves for observed and models
    observed_hill = compute_hill_curves(RF[exp_name])
    bf1_hill = compute_hill_curves(BF1[exp_name])
    bf3_hill = compute_hill_curves(BF3[exp_name])
    bf4_hill = compute_hill_curves(BF4[exp_name])
    bf6_hill = compute_hill_curves(BF6[exp_name])

    # Plot observed frequencies (first column)
    for curve in observed_hill:
        axes[i, 0].plot(Q, curve, alpha=0.8, linewidth=1)
    axes[i, 0].set_title("Observed", fontsize=12)

    # Plot Model 1 (second column)
    for curve in bf1_hill:
        axes[i, 1].plot(Q, curve, alpha=0.8, linewidth=1)
    axes[i, 1].set_title("Model $\mathcal{M}_{STAMP}$", fontsize=12)

    # Plot Model 3 (third column)
    for curve in bf3_hill:
        axes[i, 2].plot(Q, curve, alpha=0.8, linewidth=1)
    axes[i, 2].set_title("Model $\mathcal{M}_{CR}$", fontsize=12)

    # Plot Model 4 (fourth column)
    for curve in bf4_hill:
        axes[i, 3].plot(Q, curve, alpha=0.8, linewidth=1)
    axes[i, 3].set_title("Model $\mathcal{M}_{BS}$", fontsize=12)


    # Plot Model 6 (5th column)
    for curve in bf6_hill:
        axes[i, 4].plot(Q, curve, alpha=0.8, linewidth=1)
    axes[i, 4].set_title("Model $\mathcal{M}_{IR}$", fontsize=12)

# Set labels for rows with experiment names
for i, exp_name in enumerate(experiment_names):
    axes[i, 0].set_ylabel(f"{exp_name}", fontsize=12)

# Set shared x-label
for ax in axes[-1, :]:
    ax.set_xlabel("q", fontsize=12)

plt.tight_layout()
#plt.savefig("all_models_hill.png", dpi=300, bbox_inches='tight')

plt.show()
```

## Error curves

```{python, fig.width=8, fig.height=6}

# Initialize plot
plt.figure(figsize=(8, 6))

i = 1
Q = np.load(f"Models_output/Q{i}.npy")  # Load Qi
D = np.load(f"Models_output/D{i}.npy")  # Load Di
plt.plot(Q, D, linewidth=2, label="Model $\mathcal{{M}}_{STAMP}$")  # Plot with label
i = 3
Q = np.load(f"Models_output/Q{i}.npy")  # Load Qi
D = np.load(f"Models_output/D{i}.npy")  # Load Di
plt.plot(Q, D, linewidth=2, label="Model $\mathcal{{M}}_{CR}$")  # Plot with label
i = 4
Q = np.load(f"Models_output/Q{i}.npy")  # Load Qi
D = np.load(f"Models_output/D{i}.npy")  # Load Di
plt.plot(Q, D, linewidth=2, label="Model $\mathcal{{M}}_{BS}$")  # Plot with label
i = 6
Q = np.load(f"Models_output/Q{i}.npy")  # Load Qi
D = np.load(f"Models_output/D{i}.npy")  # Load Di
plt.plot(Q, D, linewidth=2, label="Model $\mathcal{{M}}_{IR}$")  # Plot with label


# Formatting the plot
plt.xlabel("$q$", fontsize=16)
plt.ylabel("Mean relative error", fontsize=16)
plt.legend(fontsize=14)
plt.xticks(fontsize=16)
plt.yticks(fontsize=16)
plt.grid()
plt.tight_layout()
#Save the figure as a PNG file
#plt.savefig("mean_error.png", dpi=300, bbox_inches='tight')

plt.show()
```

```{python, fig.width=6, fig.height=32}

# Plot mean error for each condition
# Load the saved data
RF = load_pickle("Models_output/RF.pkl")

# Extract experiment names from RF
experiment_names = list(RF.keys())

Q = np.arange(0, 4.1, 0.1)

# Create figure with subplots (7 rows, 1 columns)
fig, axes = plt.subplots(7, 1, figsize=(6, 32), sharex=True, sharey=True)

# Iterate over configurations and plot Hill curves
for j, exp_name in enumerate(experiment_names):

  i = 1
  D = np.load(f"Models_output/all_D{i}.npy")
  D = D[j]
  axes[j].plot(Q, D, alpha=0.8, linewidth=2, label="Model $\mathcal{{M}}_{STAMP}$")

  i = 3
  D = np.load(f"Models_output/all_D{i}.npy")
  D = D[j]
  axes[j].plot(Q, D, alpha=0.8, linewidth=2, label="Model $\mathcal{{M}}_{CR}$")

  i = 4
  D = np.load(f"Models_output/all_D{i}.npy")
  D = D[j]
  axes[j].plot(Q, D, alpha=0.8, linewidth=2, label="Model $\mathcal{{M}}_{BS}$")

  i = 6
  D = np.load(f"Models_output/all_D{i}.npy")
  D = D[j]
  axes[j].plot(Q, D, alpha=0.8, linewidth=2, label="Model $\mathcal{{M}}_{IR}$")

  #SV Set experiment label
  axes[j].set_ylabel(f"{exp_name}", fontsize=12)
  axes[j].grid()
  axes[j].legend(fontsize=14)

axes[j].set_xlabel("q", fontsize=12)

plt.tight_layout()
plt.show()

```

## Distribution of arrival times

### BS model

```{python}
# This is just the BS model
#**Step 1**: construct the initial random recursive tree.
def update_edges( l, k ):
    current = k
    while current != 0:
        current = l[current][0]
        l[current][2] += [k]


def init_bs( n ):
    l_edges = [ [0,[0],[]] ]
    new_edge = [0,[0],[]]
    for k in range(1,n):
        parent_new = np.random.randint(k)
        new_edge = [ parent_new, [k], []]
        l_edges += [ new_edge ]
        update_edges( l_edges, k )
    return l_edges
#**Step 2**: each edge is "lifted" at rate $1$ (in the terminology of *Berestycki 2009*).
def step_bs( tree, k ): # lift edge k of tree
    loss = 0
    parent = tree[k][0]
    tree[parent][1] += [k] + tree[k][2] # the edge is lifted to the parent
    for i in [k] + tree[k][2]:
        if tree[i][1][0] != -1:
            tree[i][1] = [-1] # unactivated
            loss += 1
    return tree, loss # tree : resulting tree; loss: number of blocks lost
#**Step 3:** the following function runs a Bolthausen-Sznitman coalescent for a time [t], starting from an initial tree [tree] with specified number of blocks [nblocks]).

def run_bs( nblocks, t, tree ):

    t_cur = np.random.exponential(1/nblocks)

    while t_cur < t and nblocks > 1:


        i = np.random.randint( 1, nblocks ) # choose the edge to be lifted
        edge = 0
        while i > -1:
            if tree[ edge ][1][0] != -1:
                edge += 1
                i -= 1
            else:
                edge += 1
        tree, loss = step_bs( tree, edge - 1 )
        nblocks -= loss

        t_cur += np.random.exponential(1/nblocks)

    return tree, nblocks


#run_bs(10000,0.5)
def display( tree ):
    for k in range( len(tree) ):
        if tree[k][1][0] != -1:
            print(tree[k][1], end='  ')
    print("")
def get_prop_bs( N, t ):
    tree,_ = run_bs( N, t, init_bs( N ) )
    prop = [] # get proportions
    for e in tree:
        if e[1][0] != -1:
            prop += [ len(e[1]) ]
    prop = prop / np.sum( prop )
    return prop
def sample_tags(prop, Ntags):
    prop_tags = np.zeros( Ntags )
    for p in prop:
        tag = np.random.randint( Ntags )
        prop_tags[ tag ] += p
    return prop_tags


```

```{python}

# Assuming you have your get_prop_bs(p1, t) function defined elsewhere

def compute_samples(args):
    t, i, p1, nreps = args
    samples = []
    for _ in range(nreps):
        prop_sample = get_prop_bs(p1, t)
        prop_sorted_sample = np.sort(prop_sample)[::-1]
        arrival_sample = -np.log(prop_sorted_sample)
        if len(arrival_sample) >= i:
            samples.append(arrival_sample[i-1] - arrival_sample[0])
    return (t, i, np.array(samples))

if __name__ == "__main__":
    p1 = 1000
    nreps = 4000
    t_values = np.arange(0.1, 1.0, 0.1)

    # Prepare tasks for multiprocessing
    tasks = []
    for t in t_values:
        for i in range(2, 21):
            tasks.append((t, i, p1, nreps))

    # Use 10 processes
    with mp.Pool(processes=10) as pool:
        results = pool.map(compute_samples, tasks)

    # Organize results into a dictionary for easier use later
    output_data = {}
    for t, i, samples in results:
        samples = np.sort(samples)
        cdf = np.arange(len(samples)) / len(samples)
        if t not in output_data:
            output_data[t] = {}
        output_data[t][i] = {'samples': samples, 'cdf': cdf}

    # Save output_data to a pickle file
    with open('Models_output/arrival_times_data_bolthausen.pkl', 'wb') as f:
        pickle.dump(output_data, f)

    print("Data saved to 'arrival_times_data_bolthausen.pkl'. Ready for plotting later!")
```

```{python, fig.width=10, fig.height=6}

# Load the saved data
with open('Models_output/arrival_times_data_bolthausen.pkl', 'rb') as f:
    output_data = pickle.load(f)

# Now plot
for t, data_by_i in output_data.items():
    if abs(t - 0.1) < 1e-8:  # Only plot when t == 0.1
        plt.figure(figsize=(10, 6))

        for i in sorted(data_by_i.keys()):
            samples = data_by_i[i]['samples']
            cdf = data_by_i[i]['cdf']
            plt.plot(samples, cdf, color='black')  # All curves in black

        # Set larger fonts
        plt.xlabel('Arrival times', fontsize=18)
        plt.ylabel('Cumulative probability', fontsize=18)
        plt.xticks(fontsize=16)
        plt.yticks(fontsize=16)

        # Remove title and legend
        # (So we don't call plt.title() or plt.legend())

        # Set x-axis limits
        plt.xlim([-1, 7])

        plt.grid(True)

        # Save the figure
        #plt.savefig('arrival_times_plot_t0.1_bs.png', dpi=300, bbox_inches='tight')

        #plt.close()  # Close the figure after saving
        
        plt.show()
        
```

### CR model

```{python, fig.width=10, fig.height=6}

# Define the Poisson rate
mu = 2.0  # You can adjust this if needed

# Set up x-axis range
x = np.linspace(-1, 7, 1000)  # As you asked: from -1 to 7

# Create the figure
plt.figure(figsize=(10, 6))

# Loop over i from 2 to 20
for i in range(2, 21):
    if i > 1:
        # For t_i - t_1, the distribution is Gamma(shape=i-1, rate=mu)
        shape = i - 1
        scale = 1 / mu  # scipy gamma uses scale = 1 / rate
        cdf = gamma.cdf(x, a=shape, scale=scale)

        plt.plot(x, cdf, color='black')  # All curves black

# Set labels and style
plt.xlabel('Arrival times', fontsize=18)
plt.ylabel('Cumulative probability', fontsize=18)
plt.xticks(fontsize=16)
plt.yticks(fontsize=16)
plt.xlim([-1, 7])
plt.grid(True)

# No legend, no title as you asked

# Save the figure
#plt.savefig('arrival_times_poisson_model.png', dpi=300, bbox_inches='tight')

plt.show()

```

### Multinomial model

```{python, fig.width=10, fig.height=6}

# Set up x-axis range
x = np.linspace(-1, 7, 1000)  # x from -1 to 7

# CDF is 0 for x < 0, and 1 for x >= 0
cdf = np.where(x >= 0, 1.0, 0.0)

# Create the figure
plt.figure(figsize=(10, 6))

# Plot same curve for all i=2 to 20
for i in range(2, 21):
    plt.plot(x, cdf, color='black')  # All curves black

# Set labels and style
plt.xlabel('Arrival times', fontsize=18)
plt.ylabel('Cumulative probability', fontsize=18)
plt.xticks(fontsize=16)
plt.yticks(fontsize=16)
plt.xlim([-1, 7])
plt.grid(True)

# No legend, no title

# Save the figure
#plt.savefig('arrival_times_all_at_zero.png', dpi=300, bbox_inches='tight')

plt.show()

```

### Kingman model

```{python}
def run_km( t, prop = [] ):

    nblocks = len(prop)
    t_cur = 0

    if nblocks > 1:
        t_cur = np.random.exponential(2/(nblocks * (nblocks-1)))

    while t_cur < t and nblocks > 1:
        t_cur += np.random.exponential(2/(nblocks * (nblocks-1)))

        i,j=[0,0]
        while i == j:
            i,j = np.random.randint( 0, nblocks, 2)
        prop[i] += prop[j] # j gets to i
        prop.pop(j) # remove j
        nblocks -= 1
    return prop
```

```{python}

def compute_samples(args):
    t, i, p1, nreps = args
    samples = []
    for _ in range(nreps):
        prop_sample = run_km(t, [1/p1] * p1)
        prop_sorted_sample = np.sort(prop_sample)[::-1]
        arrival_sample = -np.log(prop_sorted_sample)
        if len(arrival_sample) >= i:
            samples.append(arrival_sample[i-1] - arrival_sample[0])
    return (t, i, np.array(samples))

if __name__ == "__main__":
    p1 = 1000
    nreps = 4000
    t_values = np.arange(0.1, 1.0, 0.1)

    # Prepare tasks for multiprocessing
    tasks = []
    for t in t_values:
        for i in range(2, 21):
            tasks.append((t, i, p1, nreps))

    # Use 10 processes
    with mp.Pool(processes=10) as pool:
        results = pool.map(compute_samples, tasks)

    # Organize results into a dictionary for easier use later
    output_data = {}
    for t, i, samples in results:
        samples = np.sort(samples)
        cdf = np.arange(len(samples)) / len(samples)
        if t not in output_data:
            output_data[t] = {}
        output_data[t][i] = {'samples': samples, 'cdf': cdf}

    # Save output_data to a pickle file
    with open('Models_output/arrival_times_data_kingman.pkl', 'wb') as f:
        pickle.dump(output_data, f)

    print("Data saved to 'arrival_times_data_kingman.pkl'. Ready for plotting later!")

```

```{python, fig.width=10, fig.height=6}

# Load the saved data
with open('Models_output/arrival_times_data_kingman.pkl', 'rb') as f:
    output_data = pickle.load(f)

# Now plot
for t, data_by_i in output_data.items():
    if abs(t - 0.1) < 1e-8:  # Only plot when t == 0.1
        plt.figure(figsize=(10, 6))

        for i in sorted(data_by_i.keys()):
            samples = data_by_i[i]['samples']
            cdf = data_by_i[i]['cdf']
            plt.plot(samples, cdf, color='black')  # All curves in black

        # Set larger fonts
        plt.xlabel('Arrival times', fontsize=18)
        plt.ylabel('Cumulative probability', fontsize=18)
        plt.xticks(fontsize=16)
        plt.yticks(fontsize=16)

        # Remove title and legend
        # (So we don't call plt.title() or plt.legend())

        # Set x-axis limits
        plt.xlim([-1, 7])

        plt.grid(True)

        # Save the figure
        #plt.savefig('arrival_times_plot_t0.1_kingman.png', dpi=300, bbox_inches='tight')

        #plt.close()  # Close the figure after saving
        
        plt.show()

```

### Increasing rate model IR

```{python, fig.width=10, fig.height=6}

scale_t = 1
# === Step 1: Define symbolic intensity function λ(t) ===
t = sp.Symbol('t', real=True, positive=True)
s = sp.Symbol('s', real=True, positive=True)

# Example: λ(t) = mu * t ** k
mu, k = sp.symbols('mu k', real=True, positive=True)
lambda_t = scale_t * mu * sp.exp( k * t * scale_t)  # <-- change this to any intensity function
#lambda_t = mu * t ** k  # <-- change this to any intensity function
# === Step 2: Compute cumulative intensity Λ(t) ===
Lambda_t = sp.integrate(lambda_t, (t, 0, t))

# === Step 3: Solve for inverse: Λ(t) = s  =>  t = Λ⁻¹(s) ===
inv_Lambda = sp.solve(sp.Eq(Lambda_t, s), t)
if not inv_Lambda:
    raise ValueError("Could not compute symbolic inverse of Λ(t).")
inv_Lambda_expr = inv_Lambda[0]

# === Step 4: Convert to numerical function ===
inv_Lambda_func = sp.lambdify((s, mu, k), inv_Lambda_expr, modules='numpy')

# === Step 5: Simulation Parameters ===
mu_val = 0.1
k_val = 0.78
n_samples = 4000
max_i = 20
x_plot = np.linspace(-1, 7, 1000)

# === Step 6: Plotting NHPP simulation ===
plt.figure(figsize=(10, 6))

for i in range(2, max_i + 1):
    # Step 1: Homogeneous Poisson arrivals (Exponential inter-arrival times)
    inter_arrivals = np.random.exponential(scale=1.0, size=(n_samples, i))
    S_times = np.cumsum(inter_arrivals, axis=1)  # Homogeneous arrival times

    # Step 2: Transform using inverse cumulative intensity function
    T_times = inv_Lambda_func(S_times, mu_val, k_val)

    # Step 3: Compute time spread between first and last arrival
    delta_t = T_times[:, i-1]  - T_times[:, 0]

    # Step 4: Plot empirical CDF
    delta_t_sorted = np.sort(delta_t)
    cdf = np.linspace(0, 1, len(delta_t_sorted))
    plt.plot(delta_t_sorted * scale_t, cdf, color='black')

# Style
plt.xlabel('Difference in arrival time', fontsize=18)
plt.ylabel('Cumulative probability', fontsize=18)
plt.xticks(fontsize=16)
plt.yticks(fontsize=16)
plt.xlim([-1, 7])
plt.grid(True)

# Save figure
# plt.savefig('arrival_times_nhpp_symbolic.png', dpi=300, bbox_inches='tight')
plt.show()
```

```{python, fig.width=10, fig.height=6}
# This new code uses the simulation algorithm to produce the curves. It serves as a check


# Simulate NHPP with λ(t) = mu * exp(k * t), return arrival times
def simulate_nhpp_arrivals(mu, k, T, max_arrivals):
    tau = []
    s = 0.0
    while len(tau) < max_arrivals:
        s += np.random.exponential(1.0)
        t_candidate = (1.0 / k) * np.log(k * s / mu + 1)
        if t_candidate > T:
            break
        tau.append(t_candidate)
    return tau

# Parameters
mu = 0.1
k = 0.7
T = 10
n_samples = 4000
max_i = 20

# Storage for each i (2 to max_i)
delta_ts_by_i = {i: [] for i in range(2, max_i + 1)}

# Run simulations
for _ in range(n_samples):
    tau = simulate_nhpp_arrivals(mu, k, T, max_i)
    if len(tau) < 2:
        continue
    for i in range(2, min(len(tau), max_i + 1)):
        delta = tau[i - 1] - tau[0]
        delta_ts_by_i[i].append(delta)

# Plot CDFs
plt.figure(figsize=(10, 6))

for i in range(2, max_i + 1):
    delta_t = np.array(delta_ts_by_i[i])
    if len(delta_t) == 0:
        continue
    delta_t_sorted = np.sort(delta_t)
    cdf = np.linspace(0, 1, len(delta_t_sorted))
    plt.plot(delta_t_sorted, cdf, color='blue', alpha=0.5)

# Style
plt.xlabel('Difference in arrival time ($t_i - t_1$)', fontsize=18)
plt.ylabel('Cumulative probability', fontsize=18)
plt.title('CDFs of NHPP Arrival Time Differences (ModelM6)', fontsize=18)
plt.xticks(fontsize=16)
plt.yticks(fontsize=16)
plt.grid(True)
plt.xlim([-1, 7])
plt.tight_layout()
#plt.savefig('arrival_times_modelM6_nhpp.png', dpi=300)
plt.show()

```

## Build Latex Table

```{python}

# Define model numbers and experiment names
model_numbers = [1, 3, 4, 6]
model_labels = {
    1: "$\\mathcal{M}_{\\text{STAMP}}$",
    3: "$\\mathcal{M}_{\\text{CR}}$",
    4: "$\\mathcal{M}_{\\text{BS}}$",
    6: "$\\mathcal{M}_{\\text{IR}}$"
}

experiment_names = [f"Exp. {i}" for i in range(1, 9)]

experiment_mapping = {
    "Exp. 1": ('M_Intact_27_Root',),
    "Exp. 2": ('M_Cut_27_Root',),
    "Exp. 3": ('H_Intact_27_Root',),
    "Exp. 4": ('H_Cut_27_Root',),
    "Exp. 5": ('M_Intact_32_Root',),
    "Exp. 6": ('M_Cut_32_Root',),
    "Exp. 7": ('H_Intact_32_Root',),
    "Exp. 8": ('H_Cut_32_Root',),
}


latex_rows = []

for i in model_numbers:
    filename = f"Models_output/theta_values{i}.pkl"
    if i == 6:
        filename = f"Models_output/theta_values{i}.pkl"
    
    with open(filename, "rb") as f:
        theta_values = pickle.load(f)
    
    # Performance
    mean_distances = [v["mean_distance"] for v in theta_values.values()]
    n_perf = len(mean_distances)
    mean_perf = np.mean(mean_distances)
    std_perf = np.std(mean_distances, ddof=1)
    t_val_perf = t.ppf(0.975, df=n_perf - 1)
    ci_perf = t_val_perf * std_perf / np.sqrt(n_perf)

    # Get normalization factor (mean θ of Exp. 1)
    key_exp1 = experiment_mapping["Exp. 1"]
    theta_list_exp1 = theta_values[key_exp1]["theta_list"]
    if i == 4:
        inverted_exp1 = [1/theta if theta != 0 else 0 for theta in theta_list_exp1]
        norm_mean_theta = np.mean(inverted_exp1)
    else:
        norm_mean_theta = np.mean(theta_list_exp1)

    theta_strings = []
    for exp in experiment_names:
        key = experiment_mapping[exp]
        
        if key in theta_values:
            theta_list = theta_values[key]["theta_list"]
            n_theta = len(theta_list)
            t_val_theta = t.ppf(0.975, df=n_theta - 1)
            
            if i == 4:
                inverted = [1/theta if theta != 0 else 0 for theta in theta_list]
                mean_theta = np.mean(inverted)
                std_theta = np.std(inverted, ddof=1)
            else:
                mean_theta = np.mean(theta_list)
                std_theta = np.std(theta_list, ddof=1)

            ci_theta = t_val_theta * std_theta / np.sqrt(n_theta)

            # Normalize both mean and CI by Exp. 1 mean
            mean_rel = mean_theta / norm_mean_theta
            ci_rel = ci_theta / norm_mean_theta
        else:
            mean_rel = np.nan
            ci_rel = np.nan

        theta_strings.append(f"{mean_rel:.2f} ± {ci_rel:.2f}")

    model_label = model_labels[i]
    row = f"{model_label} & {mean_perf:.2f} ± {ci_perf:.2f} & " + " & ".join(theta_strings) + r" \\"
    latex_rows.append(row)

# Assemble LaTeX code
latex_code = r"""\begin{table}[h]
\centering
\tiny{
\begin{tabular}{c|c|cccccccc}
\hline
Model & Avg Performance & """ + " & ".join(experiment_names) + r""" \\
\hline
""" + "\n".join(latex_rows) + r"""
\hline
\end{tabular}}
\caption{
    Model performance (mean ± half-width of the 95\% confidence interval) and relative mean $\theta$ values (normalized by Exp. 3 for each model; mean ± half-width of the 95\% CI) across experimental conditions:
    Exp. 1: M, Intact, 27; Exp. 2: M, Cut, 27; Exp. 3: H, Intact, 27;
    Exp. 4: H, Cut, 27; Exp. 5: M, Intact, 32; Exp. 6: M, Cut, 32;
    Exp. 7: H, Intact, 32; Exp. 8: H, Cut, 32.
}
\label{tab:model_performance}
\end{table}"""

with open("Models_output/latex_table.tex", "w") as f:
    f.write(latex_code)

latex_file_path = "Models_output/latex_table.tex"


```

# V. Build csv tables with the parameters estimated from the models

In the CR model and the IR models, we measure $r$ on each replicate as follows: we  assume that the first bacteria crossed the barrier at $t=t_1$. The corresponding population size at time $t$ is equal to $\exp(r \,(t-t_1))$ and to $p_1(t) \times \text{load}$. Thus, $r \approx \frac{\ln(p_1(t) \times \text{load})}{t-t_1}$. For model $\mathcal{M}_{CR}$, we have $E[t_1] = 1/\mu = 1/ (r \theta)$, while for model $\mathcal{M}_{IR}$, we compute the median of $t_1$, see main text.

Note that the "load" value is the net number of bacteria (CFU) is computed as the product between the baterial load (CFU/g) and the weight (g).

```{python}
# STAMP Model This code builds a csv file with the data. The last column corresponds to theta

with open('Analysis_output/tab_tot.csv', newline='') as file:
    data = list(csv.reader(file, delimiter=','))

def read_file(data, configuration, tags_range, configuration_column, validation_column, time_of_sampling_column, bacterial_load_column, sample_weight_column, sample_index_column, Ntypes):
    frequencies = []
    times = []
    loads = []
    weights = []
    indexes = []

    for row in data:
        if row[configuration_column] == configuration[0]: # Check condition
            if row[validation_column] == '1': # Check sequencing validation
                counts = [int(row[j]) for j in range(tags_range, Ntypes + tags_range)]  # Adjust for weight column
                counts_sum = sum(counts)
                if counts_sum > 0:
                    frequencies.append([c / counts_sum for c in counts])
                    times.append(float(row[time_of_sampling_column]))
                    loads.append(float(row[bacterial_load_column]))
                    weights.append(float(row[sample_weight_column]))
                    indexes.append(row[sample_index_column])

    return np.array(frequencies), np.array(times), np.array(loads), np.array(weights), np.array(indexes)


def load_pickle(filename):
    with open(filename, "rb") as f:
        return pickle.load(f)

# Load theta values
theta_values = pickle.load(open('Models_output/theta_values1.pkl', 'rb'))

# Load precomputed rates (r) per config
# Here we recompute the rates so we can align them


# Read CSV into list
with open('Analysis_output/tab_tot.csv', newline='') as file:
    data = list(csv.reader(file))

# Compute r per valid row to match theta
# Compute rates for all configs
rates = {}
all_indexes = {}
for configuration in lconfigurations:
    config_key = tuple(configuration)
    frequencies, times, loads, weights, indexes = read_file(data, configuration, tags_range, configuration_column, validation_column, time_of_sampling_column, bacterial_load_column, sample_weight_column, sample_index_column, Ntypes)
    
    if config_key in theta_values:
        theta_list = np.array(theta_values[config_key]['theta_list'])
        p1 = np.max(frequencies, axis=1)
        r_ = np.full_like(p1, np.nan)
        mask = (p1>0) & (loads>0) & (times!=0) & (theta_list>0)
        r_[mask] = (np.log(p1[mask]*loads[mask]*weights[mask])+(1.0/theta_list[mask]))/times[mask]
        rates[config_key] = r_
        all_indexes[config_key] = indexes


# Prepare output file with header
#input_file = "Models_output/Data_py.csv"
input_file = "Analysis_output/tab_tot.csv"
output_file = "Models_output/data_and_estimators_model_STAMP.csv"
with open(input_file, newline='') as infile:
    reader = list(csv.reader(infile))

# Header with only theta
header = ['index'] + ['theta_STAMP']

# Track per-config index to select correct theta
idx_per_config = {tuple(c): 0 for c in lconfigurations}

with open(output_file, mode='w', newline='') as outfile:
    writer = csv.writer(outfile)
    writer.writerow(header)
    for row in reader[1:]:
        configuration = (row[configuration_column],)
        index_val = ""
        theta_val = ""  # default blank
        if configuration in theta_values and row[validation_column] == '1':
            i = idx_per_config[configuration]
            theta_list = np.array(theta_values[configuration]['theta_list'])
            index_list = all_indexes[configuration]
            if i < len(theta_list):
                index_val = index_list[i]
                theta_val = theta_list[i]
                idx_per_config[configuration] += 1
        writer.writerow([index_val] + [theta_val])

print(f"OK Saved file to {output_file}")

```

```{python}
# CR Model This code builds a csv file with the data. The last three columns correspond to theta, r, mu

with open('Analysis_output/tab_tot.csv', newline='') as file:
    data = list(csv.reader(file, delimiter=','))

def read_file(data, configuration, tags_range, configuration_column, validation_column, time_of_sampling_column, bacterial_load_column, sample_weight_column, sample_index_column, Ntypes):
    frequencies = []
    times = []
    loads = []
    weights = []
    indexes = []

    for row in data:
        if row[configuration_column] == configuration[0]: # Check condition
            if row[validation_column] == '1': # Check sequencing validation
                counts = [int(row[j]) for j in range(tags_range, Ntypes + tags_range)]  # Adjust for weight column
                counts_sum = sum(counts)
                if counts_sum > 0:
                    frequencies.append([c / counts_sum for c in counts])
                    times.append(float(row[time_of_sampling_column]))
                    loads.append(float(row[bacterial_load_column]))
                    weights.append(float(row[sample_weight_column]))
                    indexes.append(row[sample_index_column])

    return np.array(frequencies), np.array(times), np.array(loads), np.array(weights), np.array(indexes)


def load_pickle(filename):
    with open(filename, "rb") as f:
        return pickle.load(f)

# Load theta values
theta_values = pickle.load(open('Models_output/theta_values3.pkl', 'rb'))

# Load precomputed rates (r) per config
# Here we recompute the rates so we can align them


# Read CSV into list
with open('Analysis_output/tab_tot.csv', newline='') as file:
    data = list(csv.reader(file))

# Compute r per valid row to match theta
# Compute rates for all configs
rates = {}
all_indexes = {}
for configuration in lconfigurations:
    config_key = tuple(configuration)
    frequencies, times, loads, weights, indexes = read_file(data, configuration, tags_range, configuration_column, validation_column, time_of_sampling_column, bacterial_load_column, sample_weight_column, sample_index_column, Ntypes)
    
    if config_key in theta_values:
        theta_list = np.array(theta_values[config_key]['theta_list'])
        p1 = np.max(frequencies, axis=1)
        r_ = np.full_like(p1, np.nan)
        mask = (p1>0) & (loads>0) & (times!=0) & (theta_list>0)
        r_[mask] = (np.log(p1[mask]*loads[mask]*weights[mask])+(1.0/theta_list[mask]))/times[mask]
        rates[config_key] = r_
        all_indexes[config_key] = indexes


# Prepare output file with header
#input_file = "Models_output/Data_py.csv"
input_file = "Analysis_output/tab_tot.csv"
output_file = "Models_output/data_and_estimators_model_CR.csv"
with open(input_file, newline='') as infile:
    reader = list(csv.reader(infile))

# Header with only theta
header = ['index', 'theta_CR', 'r', 'mu']

# Track per-config index to select correct theta
idx_per_config = {tuple(c): 0 for c in lconfigurations}

with open(output_file, mode='w', newline='') as outfile:
    writer = csv.writer(outfile)
    writer.writerow(header)
    for row in reader[1:]:
        configuration = (row[configuration_column],)
        index_val = ""
        theta_val = ""
        r_val = ""
        mu_val = ""
        if configuration in rates and row[validation_column] == '1':
            i = idx_per_config[configuration]
            theta_list = np.array(theta_values[configuration]['theta_list'])
            r_list = rates[configuration]
            index_list = all_indexes[configuration]
            if i < len(theta_list):
                index_val = index_list[i]
                theta_val = theta_list[i]
                r_val = r_list[i]
                mu_val = theta_val * r_val
                idx_per_config[configuration] += 1
        writer.writerow([index_val, theta_val, r_val, mu_val])

print(f"OK Saved file to {output_file}")

```

```{python}
# IR Model This code builds a csv file with the data. The last three columns correspond to theta, r, mu

k = 0.78

with open('Analysis_output/tab_tot.csv', newline='') as file:
    data = list(csv.reader(file, delimiter=','))

def read_file(data, configuration, tags_range, configuration_column, validation_column, time_of_sampling_column, bacterial_load_column, sample_weight_column, sample_index_column, Ntypes):
    frequencies = []
    times = []
    loads = []
    weights = []
    indexes = []

    for row in data:
        if row[configuration_column] == configuration[0]: # Check condition
            if row[validation_column] == '1': # Check sequencing validation
                counts = [int(row[j]) for j in range(tags_range, Ntypes + tags_range)]  # Adjust for weight column
                counts_sum = sum(counts)
                if counts_sum > 0:
                    frequencies.append([c / counts_sum for c in counts])
                    times.append(float(row[time_of_sampling_column]))
                    loads.append(float(row[bacterial_load_column]))
                    weights.append(float(row[sample_weight_column]))
                    indexes.append(row[sample_index_column])

    return np.array(frequencies), np.array(times), np.array(loads), np.array(weights), np.array(indexes)


def load_pickle(filename):
    with open(filename, "rb") as f:
        return pickle.load(f)

# Load theta values
theta_values = pickle.load(open('Models_output/theta_values6.pkl', 'rb'))

# Load precomputed rates (r) per config
# Here we recompute the rates so we can align them


# Read CSV into list
with open('Analysis_output/tab_tot.csv', newline='') as file:
    data = list(csv.reader(file))

# Compute r per valid row to match theta
# Compute rates for all configs
rates = {}
all_indexes = {}
for configuration in lconfigurations:
    config_key = tuple(configuration)
    frequencies, times, loads, weights, indexes = read_file(data, configuration, tags_range, configuration_column, validation_column, time_of_sampling_column, bacterial_load_column, sample_weight_column, sample_index_column, Ntypes)
    
    if config_key in theta_values:
        theta_list = np.array(theta_values[config_key]['theta_list'])
        p1 = np.max(frequencies, axis=1)
        r_ = np.full_like(p1, np.nan)
        mask = (p1>0) & (loads>0) & (times!=0) & (theta_list>0)
        r_[mask] = (np.log(p1[mask]*loads[mask]*weights[mask])+(1/k * np.log(1 + np.log(2) * k / theta_list[mask])))/times[mask]
        rates[config_key] = r_
        all_indexes[config_key] = indexes


# Prepare output file with header
#input_file = "Models_output/Data_py.csv"
input_file = "Analysis_output/tab_tot.csv"
output_file = "Models_output/data_and_estimators_model_IR.csv"
with open(input_file, newline='') as infile:
    reader = list(csv.reader(infile))

# Header with only theta
header = ['index', 'theta_IR', 'r', 'mu']

# Track per-config index to select correct theta
idx_per_config = {tuple(c): 0 for c in lconfigurations}

with open(output_file, mode='w', newline='') as outfile:
    writer = csv.writer(outfile)
    writer.writerow(header)
    for row in reader[1:]:
        configuration = (row[configuration_column],)
        index_val = ""
        theta_val = ""
        r_val = ""
        mu_val = ""
        if configuration in rates and row[validation_column] == '1':
            i = idx_per_config[configuration]
            theta_list = np.array(theta_values[configuration]['theta_list'])
            r_list = rates[configuration]
            index_list = all_indexes[configuration]
            if i < len(theta_list):
                index_val = index_list[i]
                theta_val = theta_list[i]
                r_val = r_list[i]
                mu_val = theta_val * r_val
                idx_per_config[configuration] += 1
        writer.writerow([index_val, theta_val, r_val, mu_val])

print(f"OK Saved file to {output_file}")
```

```{python}
# BS Model This code builds a csv file with the data. The last column corresponds to theta

with open('Analysis_output/tab_tot.csv', newline='') as file:
    data = list(csv.reader(file, delimiter=','))

def read_file(data, configuration, tags_range, configuration_column, validation_column, time_of_sampling_column, bacterial_load_column, sample_weight_column, sample_index_column, Ntypes):
    frequencies = []
    times = []
    loads = []
    weights = []
    indexes = []

    for row in data:
        if row[configuration_column] == configuration[0]: # Check condition
            if row[validation_column] == '1': # Check sequencing validation
                counts = [int(row[j]) for j in range(tags_range, Ntypes + tags_range)]  # Adjust for weight column
                counts_sum = sum(counts)
                if counts_sum > 0:
                    frequencies.append([c / counts_sum for c in counts])
                    times.append(float(row[time_of_sampling_column]))
                    loads.append(float(row[bacterial_load_column]))
                    weights.append(float(row[sample_weight_column]))
                    indexes.append(row[sample_index_column])

    return np.array(frequencies), np.array(times), np.array(loads), np.array(weights), np.array(indexes)


def load_pickle(filename):
    with open(filename, "rb") as f:
        return pickle.load(f)

# Load theta values
theta_values = pickle.load(open('Models_output/theta_values4.pkl', 'rb'))

# Load precomputed rates (r) per config
# Here we recompute the rates so we can align them


# Read CSV into list
with open('Analysis_output/tab_tot.csv', newline='') as file:
    data = list(csv.reader(file))

# Compute r per valid row to match theta
# Compute rates for all configs
rates = {}
all_indexes = {}
for configuration in lconfigurations:
    config_key = tuple(configuration)
    frequencies, times, loads, weights, indexes = read_file(data, configuration, tags_range, configuration_column, validation_column, time_of_sampling_column, bacterial_load_column, sample_weight_column, sample_index_column, Ntypes)
    
    if config_key in theta_values:
        theta_list = np.array(theta_values[config_key]['theta_list'])
        p1 = np.max(frequencies, axis=1)
        r_ = np.full_like(p1, np.nan)
        mask = (p1>0) & (loads>0) & (times!=0) & (theta_list>0)
        r_[mask] = (np.log(p1[mask]*loads[mask]*weights[mask])+(1.0/theta_list[mask]))/times[mask]
        rates[config_key] = r_
        all_indexes[config_key] = indexes


# Prepare output file with header
#input_file = "Models_output/Data_py.csv"
input_file = "Analysis_output/tab_tot.csv"
output_file = "Models_output/data_and_estimators_model_BS.csv"
with open(input_file, newline='') as infile:
    reader = list(csv.reader(infile))

# Header with only theta
header = ['index'] + ['1 / theta_BS']

# Track per-config index to select correct theta
idx_per_config = {tuple(c): 0 for c in lconfigurations}

with open(output_file, mode='w', newline='') as outfile:
    writer = csv.writer(outfile)
    writer.writerow(header)
    for row in reader[1:]:
        configuration = (row[configuration_column],)
        index_val = ""
        theta_val = ""  # default blank
        if configuration in theta_values and row[validation_column] == '1':
            i = idx_per_config[configuration]
            theta_list = np.array(theta_values[configuration]['theta_list'])
            index_list = all_indexes[configuration]
            if i < len(theta_list):
                index_val = index_list[i]
                theta_val = theta_list[i]
                idx_per_config[configuration] += 1
        writer.writerow([index_val] + [theta_val])

print(f"OK Saved file to {output_file}")

```

# VI. Analysis of some correlations between permeability, bottleneck and growth

In the CR model and the BS model, we measure $r$ on each replicate as follows: we  assume that the first bacteria crossed the barrier at $t=t_1$. The corresponding population size at time $t$ is equal to $\exp(r \,(t-t_1))$ and to $p_1(t) \times \text{load}$. Thus, $r \approx \frac{\ln(p_1(t) \times \text{load})}{t-t_1}$. For model $\mathcal{M}_{CR}$, we have $E[t_1] = 1/\mu = 1/ (r \theta)$, while for model $\mathcal{M}_{IR}$, we compute the median of $t_1$, see main text.

Note that the "load" value is the net number of bacteria (CFU) is computed as the product between the baterial load (CFU/g) and the weight (g).

## Helper functions

```{python}

with open('Analysis_output/tab_tot.csv', newline='') as file:
    data = list(csv.reader(file, delimiter=','))

def read_file(data, configuration, tags_range, configuration_column, validation_column, time_of_sampling_column, bacterial_load_column, sample_weight_column, sample_index_column, Ntypes):
    frequencies = []
    times = []
    loads = []
    weights = []
    indexes = []

    for row in data:
        if row[configuration_column] == configuration[0]: # Check condition
            if row[validation_column] == '1': # Check sequencing validation
                counts = [int(row[j]) for j in range(tags_range, Ntypes + tags_range)]  # Adjust for weight column
                counts_sum = sum(counts)
                if counts_sum > 0:
                    frequencies.append([c / counts_sum for c in counts])
                    times.append(float(row[time_of_sampling_column]))
                    loads.append(float(row[bacterial_load_column]))
                    weights.append(float(row[sample_weight_column]))
                    indexes.append(row[sample_index_column])

    return np.array(frequencies), np.array(times), np.array(loads), np.array(weights), np.array(indexes)

def load_freq_load(data, lconfigurations):
    all_real_frequencies = {}  # Stores frequencies per config
    all_times = {}             # Stores times per config
    all_loads = {}             # Stores loads per config
    all_weights = {}
    all_indexes = {}

    for configuration in lconfigurations:
        real_frequencies, times, loads, weights, indexes = read_file(data, configuration, tags_range, configuration_column, validation_column, time_of_sampling_column, bacterial_load_column, sample_weight_column, sample_index_column, Ntypes)
        if len(real_frequencies) == 0:
            print(f"No data for configuration: {configuration}")
            continue

        config_key = tuple(configuration)

        all_real_frequencies[config_key] = real_frequencies
        all_times[config_key] = times
        all_loads[config_key] = loads
        all_weights[config_key] = weights
        all_indexes[config_key] = indexes

    return all_real_frequencies, all_times, all_loads, all_weights, all_indexes


def load_pickle(filename):
    with open(filename, "rb") as f:
        return pickle.load(f)

```

## CR model

```{python}
# Build the r values with theta-dependent expression
RF, all_times, all_loads, all_weights, all_indexes = load_freq_load(data, lconfigurations)

theta_values = load_pickle("Models_output/theta_values3.pkl")

results = {}

for configuration in RF:
    frequencies = RF[configuration]     # Shape: (n_samples, 120)
    times = all_times[configuration]  # Shape: (n_samples,)
    loads = all_loads[configuration]  # Shape: (n_samples,)
    weights = all_weights[configuration]  # Shape: (n_samples,)
    indexes = all_indexes[configuration]
    theta_list = np.array(theta_values[configuration]['theta_list'])  # Shape: (n_samples,)

    # Compute p1: max frequency per sample (row-wise max)
    p1 = np.max(frequencies, axis=1)  # Shape: (n_samples,)

    # Initialize r values with NaNs
    r_ = np.full_like(p1, np.nan, dtype=float)

    # Apply mask to avoid division by zero or invalid log
    safe_mask = (p1 > 0) & (loads > 0) & (times != 0) & (theta_list > 0)
    # Compute updated r using the new formula
    f1_load_product = p1[safe_mask] * loads[safe_mask] * weights[safe_mask]
    r_[safe_mask] = (
        np.log(f1_load_product) + (1.0 / theta_list[safe_mask])
    ) / times[safe_mask]

    # Store results
    results[configuration] = {
        'p1': p1,
        'r_': r_
    }

```

Correlation between $\mu$ and r

```{python, fig.width=10, fig.height=6}

# Load data
theta_values = load_pickle("Models_output/theta_values3.pkl")

# Create the scatter plot
plt.figure(figsize=(10, 6))

# Font configuration
plt.rcParams.update({
    'font.size': 14,
    'axes.labelsize': 16,
    'axes.titlesize': 18,
    'legend.fontsize': 12,
    'xtick.labelsize': 13,
    'ytick.labelsize': 13
})

# Store global lists for Spearman
all_mu = []
all_r = []

# Color cycle
color_cycle = plt.rcParams['axes.prop_cycle'].by_key()['color']

for i, configuration in enumerate(RF):
    theta_list = theta_values[configuration]['theta_list']
    r_values = results[configuration]['r_']

    # Convert and align
    theta_array = np.array(theta_list)
    r_array = np.array(r_values)
    min_len = min(len(theta_array), len(r_array))
    theta_array = theta_array[:min_len]
    r_array = r_array[:min_len]

    # Filter invalid r
    valid_mask = np.isfinite(r_array)
    theta_array = theta_array[valid_mask]
    r_array = r_array[valid_mask]
    mu_array = theta_array * r_array

    # Append to global Spearman lists
    all_mu.extend(mu_array)
    all_r.extend(r_array)

    # Clean config label (remove 'Root')
    label = " - ".join([x for x in configuration if x.lower() != 'root'])

    # Plot
    color = color_cycle[i % len(color_cycle)]
    plt.scatter(mu_array, r_array, label=label, alpha=0.7, color=color)
    plt.plot(np.mean(mu_array), np.mean(r_array), marker='x', markersize=10, markeredgewidth=2, color=color)

# Labels and style
plt.xlabel(r"$\mu = \theta \times r$")
plt.ylabel("$r$")
plt.legend(bbox_to_anchor=(1.05, 1), loc='upper left', title="Configurations")
plt.grid(True)
plt.tight_layout()

# Save and show
#plt.savefig("mu_vs_r_CR.png", dpi=300)
plt.show()

# --- Spearman correlation analysis ---
all_mu = np.array(all_mu)
all_r = np.array(all_r)

spearman_corr, spearman_pval = spearmanr(all_mu, all_r)

print(f"Spearman rank correlation coefficient (μ vs r): {spearman_corr:.4f}")
print(f"P-value: {spearman_pval:.4e}")

```

Correlation between $\mu$ and weight

```{python, fig.width=10, fig.height=6}

# Load data
theta_values = load_pickle("theta_values3.pkl")

# Create the scatter plot
plt.figure(figsize=(10, 6))

# Font configuration
plt.rcParams.update({
    'font.size': 14,
    'axes.labelsize': 16,
    'axes.titlesize': 18,
    'legend.fontsize': 12,
    'xtick.labelsize': 13,
    'ytick.labelsize': 13
})

# Store global lists for Spearman
all_mu = []
all_weights2 = []

# Color cycle
color_cycle = plt.rcParams['axes.prop_cycle'].by_key()['color']

for i, configuration in enumerate(RF):
    theta_list = theta_values[configuration]['theta_list']
    r_values = results[configuration]['r_']
    weight_values = all_weights[configuration]

    # Convert and align
    theta_array = np.array(theta_list)
    r_array = np.array(r_values)
    weight_array = np.array(weight_values)
    min_len = min(len(theta_array), len(r_array), len(weight_array))
    theta_array = theta_array[:min_len]
    r_array = r_array[:min_len]
    weight_array = weight_array[:min_len]

    # Filter valid data
    valid_mask = np.isfinite(r_array) & np.isfinite(weight_array)
    theta_array = theta_array[valid_mask]
    r_array = r_array[valid_mask]
    weight_array = weight_array[valid_mask]
    mu_array = theta_array * r_array

    # Append to global lists
    all_mu.extend(mu_array)
    all_weights2.extend(weight_array)

    # Clean config label (remove 'Root')
    label = " - ".join([x for x in configuration if x.lower() != 'root'])

    # Plot
    color = color_cycle[i % len(color_cycle)]
    plt.scatter(mu_array, weight_array, label=label, alpha=0.7, color=color)
    plt.plot(np.mean(mu_array), np.mean(weight_array), marker='x', markersize=10, markeredgewidth=2, color=color)

# Labels and style
plt.xlabel(r"$\mu = \theta \times r$")
plt.ylabel("Weight")
plt.legend(bbox_to_anchor=(1.05, 1), loc='upper left', title="Configurations")
plt.grid(True)
plt.tight_layout()

# Save and show
#plt.savefig("mu_vs_weight.png", dpi=300)
plt.show()

# --- Spearman correlation analysis ---
all_mu = np.array(all_mu)
all_weights2 = np.array(all_weights2)

spearman_corr, spearman_pval = spearmanr(all_mu, all_weights2)

print(f"Spearman rank correlation coefficient (μ vs weight): {spearman_corr:.4f}")
print(f"P-value: {spearman_pval:.4e}")

```

```{python}
# --- Spearman correlation analysis: theta vs weight ---
all_theta = np.array([theta for configuration in RF for theta in theta_values[configuration]['theta_list']])
all_weights2 = np.concatenate([all_weights[configuration][:len(theta_values[configuration]['theta_list'])] for configuration in RF])

# Filter out NaNs or mismatched lengths
valid_mask = np.isfinite(all_theta) & np.isfinite(all_weights2)
theta_valid = all_theta[valid_mask]
weight_valid = all_weights2[valid_mask]

# Compute Spearman
spearman_corr_theta, spearman_pval_theta = spearmanr(theta_valid, weight_valid)

print(f"Spearman rank correlation coefficient (θ vs weight): {spearman_corr_theta:.4f}")
print(f"P-value: {spearman_pval_theta:.4e}")

```


## IR model

```{python}

k = 0.78

# Build the r values with theta-dependent expression
RF, all_times, all_loads, all_weights, all_indexes = load_freq_load(data, lconfigurations)

theta_values = load_pickle("Models_output/theta_values6.pkl")

results = {}

for configuration in RF:
    frequencies = RF[configuration]     # Shape: (n_samples, 120)
    times = all_times[configuration]  # Shape: (n_samples,)
    loads = all_loads[configuration]  # Shape: (n_samples,)
    weights = all_weights[configuration]  # Shape: (n_samples,)
    indexes = all_indexes[configuration]
    theta_list = np.array(theta_values[configuration]['theta_list'])  # Shape: (n_samples,)

    # Compute p1: max frequency per sample (row-wise max)
    p1 = np.max(frequencies, axis=1)  # Shape: (n_samples,)

    # Initialize r values with NaNs
    r_ = np.full_like(p1, np.nan, dtype=float)

    # Apply mask to avoid division by zero or invalid log
    safe_mask = (p1 > 0) & (loads > 0) & (times != 0) & (theta_list > 0)

    # Compute updated r using the new formula
    f1_load_product = p1[safe_mask] * loads[safe_mask] * weights[safe_mask]
    r_[safe_mask] = (
        np.log(f1_load_product) + 1/k * np.log(1 + np.log(2) * k / theta_list[safe_mask])
    ) / times[safe_mask]

    # Store results
    results[configuration] = {
        'p1': p1,
        'r_': r_
    }

```

Correlation between $\mu$ and r

```{python, fig.width=10, fig.height=6}

# Load data
theta_values = load_pickle("theta_values6.pkl")

# Create the scatter plot
plt.figure(figsize=(10, 6))

# Font configuration
plt.rcParams.update({
    'font.size': 14,
    'axes.labelsize': 16,
    'axes.titlesize': 18,
    'legend.fontsize': 12,
    'xtick.labelsize': 13,
    'ytick.labelsize': 13
})

# Store global lists for Spearman
all_mu = []
all_r = []

# Color cycle
color_cycle = plt.rcParams['axes.prop_cycle'].by_key()['color']

for i, configuration in enumerate(RF):
    theta_list = theta_values[configuration]['theta_list']
    r_values = results[configuration]['r_']

    # Convert and align
    theta_array = np.array(theta_list)
    r_array = np.array(r_values)
    min_len = min(len(theta_array), len(r_array))
    theta_array = theta_array[:min_len]
    r_array = r_array[:min_len]

    # Filter invalid r
    valid_mask = np.isfinite(r_array)
    theta_array = theta_array[valid_mask]
    r_array = r_array[valid_mask]
    mu_array = theta_array * r_array

    # Append to global Spearman lists
    all_mu.extend(mu_array)
    all_r.extend(r_array)

    # Clean config label (remove 'Root')
    label = " - ".join([x for x in configuration if x.lower() != 'root'])

    # Plot
    color = color_cycle[i % len(color_cycle)]
    plt.scatter(mu_array, r_array, label=label, alpha=0.7, color=color)
    plt.plot(np.mean(mu_array), np.mean(r_array), marker='x', markersize=10, markeredgewidth=2, color=color)

# Labels and style
plt.xlabel(r"$\mu = \theta \times r$")
plt.ylabel("$r$")
plt.legend(bbox_to_anchor=(1.05, 1), loc='upper left', title="Configurations")
plt.grid(True)
plt.tight_layout()

# Save and show
#plt.savefig("mu_vs_r_IR.png", dpi=300)
plt.show()

# --- Spearman correlation analysis ---
all_mu = np.array(all_mu)
all_r = np.array(all_r)

spearman_corr, spearman_pval = spearmanr(all_mu, all_r)

print(f"Spearman rank correlation coefficient (μ vs r): {spearman_corr:.4f}")
print(f"P-value: {spearman_pval:.4e}")

```

Correlation between $\mu$ and weight

```{python, fig.width=10, fig.height=6}

# Load data
theta_values = load_pickle("theta_values6.pkl")

# Create the scatter plot
plt.figure(figsize=(10, 6))

# Font configuration
plt.rcParams.update({
    'font.size': 14,
    'axes.labelsize': 16,
    'axes.titlesize': 18,
    'legend.fontsize': 12,
    'xtick.labelsize': 13,
    'ytick.labelsize': 13
})

# Store global lists for Spearman
all_mu = []
all_weights2 = []

# Color cycle
color_cycle = plt.rcParams['axes.prop_cycle'].by_key()['color']

for i, configuration in enumerate(RF):
    theta_list = theta_values[configuration]['theta_list']
    r_values = results[configuration]['r_']
    weight_values = all_weights[configuration]

    # Convert and align
    theta_array = np.array(theta_list)
    r_array = np.array(r_values)
    weight_array = np.array(weight_values)
    min_len = min(len(theta_array), len(r_array), len(weight_array))
    theta_array = theta_array[:min_len]
    r_array = r_array[:min_len]
    weight_array = weight_array[:min_len]

    # Filter valid data
    valid_mask = np.isfinite(r_array) & np.isfinite(weight_array)
    theta_array = theta_array[valid_mask]
    r_array = r_array[valid_mask]
    weight_array = weight_array[valid_mask]
    mu_array = theta_array * r_array

    # Append to global lists
    all_mu.extend(mu_array)
    all_weights2.extend(weight_array)

    # Clean config label (remove 'Root')
    label = " - ".join([x for x in configuration if x.lower() != 'root'])

    # Plot
    color = color_cycle[i % len(color_cycle)]
    plt.scatter(mu_array, weight_array, label=label, alpha=0.7, color=color)
    plt.plot(np.mean(mu_array), np.mean(weight_array), marker='x', markersize=10, markeredgewidth=2, color=color)

# Labels and style
plt.xlabel(r"$\mu = \theta \times r$")
plt.ylabel("Weight")
plt.legend(bbox_to_anchor=(1.05, 1), loc='upper left', title="Configurations")
plt.grid(True)
plt.tight_layout()

# Save and show
#plt.savefig("mu_vs_weight.png", dpi=300)
plt.show()

# --- Spearman correlation analysis ---
all_mu = np.array(all_mu)
all_weights2 = np.array(all_weights2)

spearman_corr, spearman_pval = spearmanr(all_mu, all_weights2)

print(f"Spearman rank correlation coefficient (μ vs weight): {spearman_corr:.4f}")
print(f"P-value: {spearman_pval:.4e}")

```

```{python}
# --- Spearman correlation analysis: theta vs weight ---
all_theta = np.array([theta for configuration in RF for theta in theta_values[configuration]['theta_list']])
all_weights2 = np.concatenate([all_weights[configuration][:len(theta_values[configuration]['theta_list'])] for configuration in RF])

# Filter out NaNs or mismatched lengths
valid_mask = np.isfinite(all_theta) & np.isfinite(all_weights2)
theta_valid = all_theta[valid_mask]
weight_valid = all_weights2[valid_mask]

# Compute Spearman
spearman_corr_theta, spearman_pval_theta = spearmanr(theta_valid, weight_valid)

print(f"Spearman rank correlation coefficient (θ vs weight): {spearman_corr_theta:.4f}")
print(f"P-value: {spearman_pval_theta:.4e}")

```

# VII. New analysis of the R. pseudosolanacearum infection dataset in R using the results of the model testing approach (CR, BS and IR models)

## Visualisation of models outputs for the experiment of infection in varying conditions (tab_tot)

Visualisation of the output parameters theta, mu and r

```{r Graph}

tab = tab_tot

tab_temp = read.csv("Models_output/data_and_estimators_model_STAMP.csv", header= TRUE, check.names= FALSE)
tab$Theta_STAMP = tab_temp$theta_STAMP
tab_temp = read.csv("Models_output/data_and_estimators_model_CR.csv", header= TRUE, check.names= FALSE)
tab$Theta_CR = tab_temp$theta_CR
tab$Mu_CR = tab_temp$mu
tab$r_CR = tab_temp$r
tab_temp = read.csv("Models_output/data_and_estimators_model_BS.csv", header= TRUE, check.names= FALSE)
tab$Theta_BS = tab_temp$`1 / theta_BS`
tab$Theta_BS = 1 / tab$Theta_BS
tab_temp = read.csv("Models_output/data_and_estimators_model_IR.csv", header= TRUE, check.names= FALSE)
tab$Theta_IR = tab_temp$theta
tab$Mu_IR = tab_temp$mu
tab$r_IR = tab_temp$r

tab = tab %>%
  filter(Zone == "Root") %>%
  filter(`PCR2 validation on gel` > 0)


####################
# Visualisation of the STAMP model (Nb)
####################

# Kruskal-wallis statistical test
test = Graph_kruskal(tab= tab, y= "Theta_STAMP", x= "Condition2")

# Display helper
values = tab %>% 
  group_by(Condition2, .drop= FALSE) %>% 
  summarise(mean = mean(Theta_STAMP), sd = sd(Theta_STAMP))


# Graph
ggplot(data= tab, aes(x= Condition2, y= Theta_STAMP)) +
  
  geom_boxplot(outlier.shape= NA, width=0.9, fill= "lightblue") +
  geom_point(aes(colour= Replicate),
                  position= position_jitterdodge(jitter.width= 0.3, dodge.width= 0.4), alpha= 1) +
  
  scale_colour_manual(values= c("#E41A1C", "#339900", "#FFD92F")) +
  geom_text(data = values, aes(label = test$groups, y = 1 , x = Condition2), vjust = -22, inherit.aes = FALSE) +
  scale_x_discrete(drop= FALSE) +
  scale_y_continuous(trans= "log10", limits= c(1,280)) +
  labs(title= "Instantaneous bottleneck STAMP model", y = "Bottleneck Nb (Number of bacteria)") +
  theme_gray()


####################
# Visualisation of the CR model (theta, mu and r)
####################

# Theta CR
# Kruskal-wallis statistical test
test = Graph_kruskal(tab= tab, y= "Theta_CR", x= "Condition2")

# Display helper
values = tab %>% 
  group_by(Condition2, .drop= FALSE) %>% 
  summarise(mean = mean(Theta_CR), sd = sd(Theta_CR))

# Graph theta CR
ggplot(data= tab, aes(x= Condition2, y= Theta_CR)) +
  
  geom_boxplot(outlier.shape= NA, width=0.9, fill= "pink") +
  geom_point(aes(colour= Replicate),
                  position= position_jitterdodge(jitter.width= 0.3, dodge.width= 0.4), alpha= 1) +
  
  scale_colour_manual(values= c("#E41A1C", "#339900", "#FFD92F")) +
  geom_text(data = values, aes(label = test$groups, y = 1 , x = Condition2), vjust = -21.7, inherit.aes = FALSE) +
  scale_x_discrete(drop= FALSE) +
  scale_y_continuous(limits= c(0,52)) +
  labs(title= "Constant entry rate CR model", y = "Bottleneck effect mu/r (no units)") +
  theme_gray()


# Mu CR
# Kruskal-wallis statistical test
test = Graph_kruskal(tab= tab, y= "Mu_CR", x= "Condition2")

# Display helper
values = tab %>% 
  group_by(Condition2, .drop= FALSE) %>% 
  summarise(mean = mean(Mu_CR), sd = sd(Mu_CR))

# Graph mu CR
ggplot(data= tab, aes(x= Condition2, y= Mu_CR), ) +
  
  geom_boxplot(outlier.shape= NA, width=0.9, fill= "pink") +
  geom_point(aes(colour= Replicate),
                  position= position_jitterdodge(jitter.width= 0.3, dodge.width= 0.4), alpha= 1) +
  
  scale_colour_manual(values= c("#E41A1C", "#339900", "#FFD92F")) +
  geom_text(data = values, aes(label = test$groups, y = 1 , x = Condition2), vjust = -21.7, inherit.aes = FALSE) +
  scale_x_discrete(drop= FALSE) +
  scale_y_continuous(limits= c(0,250)) +
  labs(title= "Constant entry rate CR model", y = "Permeability mu (CFU/d)") +
  theme_gray()


# r CR
# Kruskal-wallis statistical test
test = Graph_kruskal(tab= tab, y= "r_CR", x= "Condition2")

# Display helper
values = tab %>% 
  group_by(Condition2, .drop= FALSE) %>% 
  summarise(mean = mean(r_CR), sd = sd(r_CR))

# Graph r CR
ggplot(data= tab, aes(x= Condition2, y= r_CR), ) +
  
  geom_boxplot(outlier.shape= NA, width=0.9, fill= "pink") +
  geom_point(aes(colour= Replicate),
                  position= position_jitterdodge(jitter.width= 0.3, dodge.width= 0.4), alpha= 1) +
  
  scale_colour_manual(values= c("#E41A1C", "#339900", "#FFD92F")) +
  geom_text(data = values, aes(label = test$groups, y = 1 , x = Condition2), vjust = -20.7, inherit.aes = FALSE) +
  scale_x_discrete(drop= FALSE) +
  scale_y_continuous(limits= c(0,6.5)) +
  labs(title= "Constant entry rate CR model", y = "Growth rate r (d-1)") +
  theme_gray()

####################
# Visualisation of the BS model (theta)
####################

# Theta BS
# Kruskal-wallis statistical test
test = Graph_kruskal(tab= tab, y= "Theta_BS", x= "Condition2")

# Display helper
values = tab %>% 
  group_by(Condition2, .drop= FALSE) %>% 
  summarise(mean = mean(Theta_BS), sd = sd(Theta_BS))

# Graph theta BS
ggplot(data= tab, aes(x= Condition2, y= Theta_BS), ) +
  
  geom_boxplot(outlier.shape= NA, width=0.9, fill= "lightgreen") +
  geom_point(aes(colour= Replicate),
                  position= position_jitterdodge(jitter.width= 0.3, dodge.width= 0.4), alpha= 1) +
  
  scale_colour_manual(values= c("#E41A1C", "#339900", "#FFD92F")) +
  geom_text(data = values, aes(label = test$groups, y = 1 , x = Condition2), vjust = -22, inherit.aes = FALSE) +
  scale_x_discrete(drop= FALSE) +
  scale_y_continuous(trans= "log10", limits= c(1,23), breaks= c(1,3,5,10,20)) +
  labs(title= "Bolthausen-Sznitsman BS model", y = "Normalised entry speed v (no units)") +
  theme_gray()


# Graph theta BS v2
ggplot(data= tab, aes(x= Condition2, y= Theta_BS, fill= Genotype), ) +
  
  geom_boxplot(outlier.shape= NA, width=0.8) +
  geom_point(aes(colour= Replicate),
                  position= position_jitterdodge(jitter.width= 0.3, dodge.width= 0.4), alpha= 1) +
  
  scale_fill_manual(values= c("#6699FF", "#FF9900")) +
  scale_colour_manual(values= c("#E41A1C", "#339900", "#FFD92F")) +
  geom_text(data = values, aes(label = test$groups, y = 1 , x = Condition2), vjust = -23.5, inherit.aes = FALSE) +
  scale_x_discrete(drop= FALSE) +
  scale_y_continuous(trans= "log10", limits= c(1,23), breaks= c(1,3,5,10,20)) +
  labs(y = bquote("Bottleneck effect"~theta[BS]*" (no units)")) +
  theme_gray()


####################
# Visualisation of the IR model (theta, mu and r)
####################

# Theta IR
# Kruskal-wallis statistical test
test = Graph_kruskal(tab= tab, y= "Theta_IR", x= "Condition2")

# Display helper
values = tab %>% 
  group_by(Condition2, .drop= FALSE) %>% 
  summarise(mean = mean(Theta_IR), sd = sd(Theta_IR))

# Graph theta IR
ggplot(data= tab, aes(x= Condition2, y= Theta_IR), ) +
  
  geom_boxplot(outlier.shape= NA, width=0.9, fill= "tomato") +
  geom_point(aes(colour= Replicate),
                  position= position_jitterdodge(jitter.width= 0.3, dodge.width= 0.4), alpha= 1) +
  
  scale_colour_manual(values= c("#E41A1C", "#339900", "#FFD92F")) +
  geom_text(data = values, aes(label = test$groups, y = 1 , x = Condition2), vjust = -0.9, inherit.aes = FALSE) +
  scale_x_discrete(drop= FALSE) +
  scale_y_continuous(limits= c(0,1.05)) +
  labs(title= "Increasing entry rate IR model", y = "Bacterial capacity index mu/r (no units)") +
  theme_gray()


# Mu IR
# Kruskal-wallis statistical test
test = Graph_kruskal(tab= tab, y= "Mu_IR", x= "Condition2")

# Display helper
values = tab %>% 
  group_by(Condition2, .drop= FALSE) %>% 
  summarise(mean = mean(Mu_IR), sd = sd(Mu_IR))

# Graph mu IR
ggplot(data= tab, aes(x= Condition2, y= Mu_IR), ) +
  
  geom_boxplot(outlier.shape= NA, width=0.9, fill= "tomato") +
  geom_point(aes(colour= Replicate),
                  position= position_jitterdodge(jitter.width= 0.3, dodge.width= 0.4), alpha= 1) +
  
  scale_colour_manual(values= c("#E41A1C", "#339900", "#FFD92F")) +
  geom_text(data = values, aes(label = test$groups, y = 1 , x = Condition2), vjust = -17.8, inherit.aes = FALSE) +
  scale_x_discrete(drop= FALSE) +
  scale_y_continuous(limits= c(0,5.3)) +
  labs(title= "Increasing entry rate IR model", y = "Permeability mu (CFU/d)") +
  theme_gray()

# Graph mu IR v2
ggplot(data= tab, aes(x= Condition2, y= Mu_IR, fill= Genotype), ) +
  
  geom_boxplot(outlier.shape= NA, width=0.8) +
  geom_point(aes(colour= Replicate),
                  position= position_jitterdodge(jitter.width= 0.3, dodge.width= 0.4), alpha= 1) +
  
  scale_fill_manual(values= c("#6699FF", "#FF9900")) +
  scale_colour_manual(values= c("#E41A1C", "#339900", "#FFD92F")) +
  geom_text(data = values, aes(label = test$groups, y = 1 , x = Condition2), vjust = -18.9, inherit.aes = FALSE) +
  scale_x_discrete(drop= FALSE) +
  scale_y_continuous(limits= c(0,5.3)) +
  labs(y = bquote("Roots permeability multiplier"~mu[IR]*" (no units)")) +
  theme_gray()


# r IR
# Kruskal-wallis statistical test
test = Graph_kruskal(tab= tab, y= "r_IR", x= "Condition2")

# Display helper
values = tab %>% 
  group_by(Condition2, .drop= FALSE) %>% 
  summarise(mean = mean(r_IR), sd = sd(r_IR))

# Graph r IR
ggplot(data= tab, aes(x= Condition2, y= r_IR), ) +
  
  geom_boxplot(outlier.shape= NA, width=0.9, fill= "tomato") +
  geom_point(aes(colour= Replicate),
                  position= position_jitterdodge(jitter.width= 0.3, dodge.width= 0.4), alpha= 1) +
  
  scale_colour_manual(values= c("#E41A1C", "#339900", "#FFD92F")) +
  geom_text(data = values, aes(label = test$groups, y = 1 , x = Condition2), vjust = -18.5, inherit.aes = FALSE) +
  scale_x_discrete(drop= FALSE) +
  scale_y_continuous(limits= c(0,7)) +
  labs(title= "Increasing entry rate IR model", y = "Growth rate r (d-1)") +
  theme_gray()

# Graph r IR v2
ggplot(data= tab, aes(x= Condition2, y= r_IR, fill= Genotype), ) +
  
  geom_boxplot(outlier.shape= NA, width=0.9) +
  geom_point(aes(colour= Replicate),
                  position= position_jitterdodge(jitter.width= 0.3, dodge.width= 0.4), alpha= 1) +
  
  scale_fill_manual(values= c("#6699FF", "#FF9900")) +
  scale_colour_manual(values= c("#E41A1C", "#339900", "#FFD92F")) +
  geom_text(data = values, aes(label = test$groups, y = 1 , x = Condition2), vjust = -20, inherit.aes = FALSE) +
  scale_x_discrete(drop= FALSE) +
  scale_y_continuous(limits= c(0,7)) +
  labs(y = bquote("In planta growth rate"~r[IR]*" ("~d^-1*")")) +
  theme_gray()  

```

Statistical tests to see the contribution of theta, mu and r to the infection

```{r Calculation}

tab = tab_tot

tab_temp = read.csv("Models_output/data_and_estimators_model_STAMP.csv", header= TRUE, check.names= FALSE)
tab$Theta_STAMP = tab_temp$theta_STAMP
tab_temp = read.csv("Models_output/data_and_estimators_model_CR.csv", header= TRUE, check.names= FALSE)
tab$Theta_CR = tab_temp$theta_CR
tab$Mu_CR = tab_temp$mu
tab$r_CR = tab_temp$r
tab_temp = read.csv("Models_output/data_and_estimators_model_BS.csv", header= TRUE, check.names= FALSE)
tab$Theta_BS = tab_temp$`1 / theta_BS`
tab$Theta_BS = 1 / tab$Theta_BS
tab_temp = read.csv("Models_output/data_and_estimators_model_IR.csv", header= TRUE, check.names= FALSE)
tab$Theta_IR = tab_temp$theta
tab$Mu_IR = tab_temp$mu
tab$r_IR = tab_temp$r

tab = tab %>%
  filter(Zone == "Root") %>%
  filter(`PCR2 validation on gel` > 0)

tab_temp = select(tab, c("Replicate","Genotype", "Temperature (°C)", "Root condition", "Collected fresh weight (g)", "Theta_STAMP", "Theta_CR", "Theta_BS", "Theta_IR", "r_CR", "r_IR", "Mu_CR", "Mu_IR", "Shannon diversity", "Bacterial load (CFU/g)", "Time post-inoculation (d)"))

# Encode each condition to logical
# M = 0, H = 1
tab_temp$Genotype = as.integer(tab_temp$Genotype != "M")
# 27°C = 0, 32°C = 1
tab_temp$`Temperature (°C)` = as.integer(tab_temp$`Temperature (°C)` != "27°C")
# Intact = 0, Cut = 1
tab_temp$`Root condition` = as.integer(tab_temp$`Root condition` != "Intact")


####################
# Testing homoscedasticity between conditions
####################

a = levene_test(tab, Theta_BS ~ Condition4)
print(a)

a = levene_test(tab, Mu_IR ~ Condition4)
print(a)

a = levene_test(tab, r_IR ~ Condition4)
print(a)


####################
# Multiple linear regression
# Contribution of the different experimental conditions to the models outputs
####################

# Multiple linear regression test with principal effects only then principal effect + secondary interactions

# Bottleneck NB in the STAMP model
# Principal effects
a = lm(Theta_STAMP ~ (`Root condition` + Genotype + `Temperature (°C)`), data = tab_temp)
print(summary(a))
# Principal effects + secondary interactions
a = lm(Theta_STAMP ~ (`Root condition` + Genotype + `Temperature (°C)`)^2, data = tab_temp)
print(summary(a))


# Bottleneck theta in the BS model
# Principal effects
a = lm(Theta_BS ~ (`Root condition` + Genotype + `Temperature (°C)`), data = tab_temp)
print(summary(a))
# Principal effects + secondary interactions
a = lm(Theta_BS ~ (`Root condition` + Genotype + `Temperature (°C)`)^2, data = tab_temp)
print(summary(a))


# Growth rate in the IR model
# Principal effects
a = lm(r_IR ~ (`Root condition` + Genotype + `Temperature (°C)`), data = tab_temp)
print(summary(a))
# Principal effects + secondary interactions
a = lm(r_IR ~ (`Root condition` + Genotype + `Temperature (°C)`)^2, data = tab_temp)
print(summary(a))


# Permeability in the IR model
# Principal effects
a = lm(Mu_IR ~ (`Root condition` + Genotype + `Temperature (°C)`), data = tab_temp)
print(summary(a))
# Principal effects + secondary interactions
a = lm(Mu_IR ~ (`Root condition` + Genotype + `Temperature (°C)`)^2, data = tab_temp)
print(summary(a))


####################
# Non-parametric PERMANOVA test
# Contribution of the different experimental conditions to the models outputs
####################

# Bottleneck Nb in the STAMP model
#Principal effects
test = adonis2(formula = tab_temp$Theta_STAMP ~ (`Root condition` + Genotype + `Temperature (°C)`),
                       data = tab_temp, permutations= 1000, by= "terms")
print(test)
# Principal effects + secondary interactions
test = adonis2(formula = tab_temp$Theta_STAMP ~ (`Root condition` + Genotype + `Temperature (°C)`)^2,
                       data = tab_temp, permutations= 1000, by= "terms")
print(test)


# Bottleneck theta in the BS model
# Principal effects
test = adonis2(formula = tab_temp$Theta_BS ~ (`Root condition` + Genotype + `Temperature (°C)`),
                       data = tab_temp, permutations= 1000, by= "terms")
print(test)
# Principal effects + secondary interactions
test = adonis2(formula = tab_temp$Theta_BS ~ (`Root condition` + Genotype + `Temperature (°C)`)^2,
                       data = tab_temp, permutations= 1000, by= "terms")
print(test)


# Growth rate r in the IR model
# Principal effects
test = adonis2(formula = tab_temp$r_IR ~ (`Root condition` + Genotype + `Temperature (°C)`),
                       data = tab_temp, permutations= 1000, by= "terms")
print(test)
# Principal effects + secondary interactions
test = adonis2(formula = tab_temp$r_IR ~ (`Root condition` + Genotype + `Temperature (°C)`)^2,
                       data = tab_temp, permutations= 1000, by= "terms")
print(test)


# Permeability mu in the IR model
# Principal effects
test = adonis2(formula = tab_temp$Mu_IR ~ (`Root condition` + Genotype + `Temperature (°C)`),
                       data = tab_temp, permutations= 1000, by= "terms")
print(test)
# Principal effects + secondary interactions
test = adonis2(formula = tab_temp$Mu_IR ~ (`Root condition` + Genotype + `Temperature (°C)`)^2,
                       data = tab_temp, permutations= 1000, by= "terms")
print(test)


# Contribution of plant weight to mu and r in the IR model
# r
test = adonis2(formula = tab_temp$r_IR ~ (`Collected fresh weight (g)` +`Root condition` + Genotype + `Temperature (°C)`),
                       data = tab_temp, permutations= 1000, by= "terms")
print(test)
test = adonis2(formula = tab_temp$r_IR ~ (`Collected fresh weight (g)` +`Root condition` + Genotype + `Temperature (°C)`)^2,
                       data = tab_temp, permutations= 1000, by= "terms")
print(test)

# Mu
test = adonis2(formula = tab_temp$Mu_IR ~ (`Collected fresh weight (g)` +`Root condition` + Genotype + `Temperature (°C)`),
                       data = tab_temp, permutations= 1000, by= "terms")
print(test)
test = adonis2(formula = tab_temp$Mu_IR ~ (`Collected fresh weight (g)` +`Root condition` + Genotype + `Temperature (°C)`)^2,
                       data = tab_temp, permutations= 1000, by= "terms")
print(test)

```

## Visualisation of the shape of diversity according to each model

```{r Graph}

# Retrieving table of simulated barcodes according to each model (for the M_Intact_27 condition)

tab_predict = read_xlsx("Models_output/Barcodes_best_prediction_MI27/Manual_all_in_one_table.xlsx")

tab_predict$`Data type` = factor(tab_predict$`Data type`,
                                 levels= c("Observed", "STAMP", "CR", "BS", "IR"),
                                 labels = c("Observed", "Best fit STAMP", "Best fit CR", "Best fit BS", "Best fit IR"))

tab_predict$`Plant ID` = factor(tab_predict$`Plant ID`)

tab = tab_predict

tab_temp = tab

####################
# Visualising of the real data versus the best fitting simulations attempting to reproduce them according to each model
####################

# Ordering by frequency value
for (i in c(1:nrow(tab_temp))) {
  tab_temp[i,5:124] = tab_temp[i,5:124][,rev(order(tab_temp[i,5:124]))]
}

tab_temp = pivot_longer(tab_temp, cols = c(5:124))
tab = pivot_longer(tab, cols = c(5:124))

# Removing zero values for graph
tab_temp = tab_temp %>%
  filter(value > 0)

# Converting tag ID to number
tab_temp$name = as.numeric(substring(tab_temp$name, 4))

# retrieving the default pyplot colour palette
colour_tab10 = c("#1f77b4", "#ff7f0e", "#2ca02c", "#d62728", "#9467bd", "#8c564b", "#e377c2", "#7f7f7f", "#bcbd22", "#17becf", "#1f77b4", "#ff7f0e", "#2ca02c", "#d62728", "#9467bd", "#8c564b", "#e377c2")


# Graph
ggplot(data= tab_temp, aes(x= name, y= value, colour= `Plant ID`)) +
  
  geom_point(shape= 1, size= 1) +
  
  facet_grid(~ `Data type`) +
  scale_colour_discrete(type= colour_tab10) +
  scale_x_continuous(limits= c(1,120), breaks= c(1,40,80,120)) +
  scale_y_continuous(trans= "log10", limits= c(10^-5,1)) +
  labs(x= "Ordonated tags", y= "Frequency") +
  theme_pubr() +
  theme(panel.spacing= unit(0.7, "lines"), legend.position = "none",
        title = element_text(size= 11), axis.text = element_text(size= 9), legend.text = element_text(size= 9))
  
```

# Section reanalysis of the tomato dataset including correction with the starting inoculum frequencies

Launch this at the beginning of the session to load data

N.B.: don't forget to set up your Python environment in Rstudio.

```{python}

import numpy as np
import matplotlib.pyplot as plt
import csv
import os
import time
from matplotlib.animation import FuncAnimation
from matplotlib import pylab
import math
import pickle
import multiprocessing as mp
from scipy.stats import gamma
import sympy as sp
from scipy.stats import t
from scipy.optimize import minimize_scalar
import matplotlib.cm as cm
from scipy.stats import spearmanr


# Info retrieved from document "Analysis_output/tab_tot.csv" for the functions "read_file"
Ntypes = 120 # Number of different tags
tags_range = 55 # Starting column for tags
sample_index_column = 1 # Column with sample names
configuration_column = 5 # Column that contains all tested samples conditions
time_of_sampling_column = 17 # Column with DPI of sampling
bacterial_load_column = 33
sample_weight_column = 19 #only necessary if the input is bacterial load to convert it to number of CFU
validation_column = 343 # Column to check that the sample is validated
inoculum_index_column = 214 # Index of the inoculum that was used for the infection
q_values = np.linspace(0, 8, 100)  # Range of q-values for Hill numbers

# Configurations
lconfigurations = [
    ['M_Intact_27_Root'],
    ['M_Cut_27_Root'],
    ['H_Intact_27_Root'],
    ['H_Cut_27_Root'],
    ['M_Intact_32_Root'],
    ['M_Cut_32_Root'],
    ['H_Intact_32_Root'],
    ['H_Cut_32_Root'],
    ]

# Info regarding the inoculum table document "Analysis_output/tab_inoc.csv"
Ntypes = Ntypes # Samples and inoculum should have identical number of tags
corresponding_inoculum_index_column = 144 # Index of the inoculum that was used for the infection
corresponding_inoculum_tags_range = 146 # Starting column for tags in inoculum
inoculum_validation_column = 145 # Column to check that the inoculum is validated

```

# I. NULL

# II. Reanalysis of the tomato dataset with inoculum correction (do not relaunch unless necessary; outputs are already saved)

## Initialisation of the helper functions

```{python, fig.width=8, fig.height=6}
# @title Main functions common to all models. Launch before the estimation

def sample_theta(K, theta_range, starting_frequencies, distribution_model):
    theta = np.random.uniform(*theta_range)
    V = distribution_model(K, theta, starting_frequencies)
    return V / np.sum(V)

def quantile_distance(theta_range, frequencies, starting_frequencies, distribution_model, parameter_name="theta", Nsim=100, quantile=0.05):
    K = len(frequencies)
    Q = np.arange(0, 4.1, 0.1)

    def hill_number(V, q):
        if q == 1:
            return np.exp(-np.sum(V[V > 0] * np.log(V[V > 0])))
        elif q == 0:
            return np.sum(V > 0)
        else:
            return (np.sum(V ** q)) ** (1 / (1 - q))

    def calculate_distance(V1, V2):
        DS = 0
        for q in Q:
            Hq1 = hill_number(V1, q)
            Hq2 = hill_number(V2, q)
            DS += abs(Hq1 - Hq2) / (Hq1 + Hq2)
        return DS / len(Q)

    distances = []
    for _ in range(Nsim):
        V = sample_theta(K, theta_range, starting_frequencies, distribution_model)
        D = calculate_distance(V, frequencies)
        distances.append(D)

    return np.quantile(distances, quantile)

def find_best_theta(frequencies, starting_frequencies, theta_range, distribution_model, distance_function, parameter_name="theta", step=1):
    distances = []
    theta_values = np.arange(theta_range[0], theta_range[1] + step, step)

    for theta in theta_values:
        dist = distance_function((theta, theta), frequencies, starting_frequencies, distribution_model, parameter_name)
        distances.append(dist)

    best_theta = theta_values[np.argmin(distances)]
    min_distance = min(distances)
    return best_theta, min_distance


def read_file(data, configuration, tags_range, configuration_column, validation_column, sample_index_column, time_of_sampling_column, bacterial_load_column, inoculum_index_column, Ntypes):
    frequencies = []
    conditions = []
    for row in data:
        if row[configuration_column] == configuration[0]: # Check condition
            if row[validation_column] == '1': # Check sequencing validation
                counts = [int(row[j]) for j in range(tags_range, Ntypes + tags_range)]
                counts_sum = sum(counts)
                if counts_sum > 0:
                    frequencies.append([c / counts_sum for c in counts])
                    time = float(row[17])
                    conditions.append((row[sample_index_column], int(row[time_of_sampling_column]),float(row[bacterial_load_column]), row[inoculum_index_column]))

    return np.array(frequencies), np.array(conditions)


def read_starting_freq(inoc, corresponding_inoculum_tags_range, corresponding_inoculum_index_column, inoculum_validation_column, Ntypes):
    frequencies = []
    conditions = []
    for row in inoc:
      if row[inoculum_validation_column] == '1': # Check sequencing validation
          counts = [int(row[j]) for j in range(corresponding_inoculum_tags_range, Ntypes + corresponding_inoculum_tags_range)]
          counts_sum = sum(counts)
          if counts_sum > 0:
              frequencies.append([c / counts_sum for c in counts])
              time = float(row[17])
              conditions.append((row[corresponding_inoculum_index_column]))
            
    return np.array(frequencies), np.array(conditions)

def process_configurations(data, inoc, lconfigurations, distribution_model, theta_range, step, distance_function=quantile_distance, parameter_name="theta", model_name="Adaptable Frequency Model"):
    theta_values = {}  # Dictionary to store theta lists, mean values, and performance
    results = []
    
    starting_frequencies, inoculum_index = read_starting_freq(inoc, corresponding_inoculum_tags_range, corresponding_inoculum_index_column, inoculum_validation_column, Ntypes)

    for configuration in lconfigurations:
        frequencies, metadata = read_file(data, configuration, tags_range, configuration_column, validation_column, sample_index_column, time_of_sampling_column, bacterial_load_column, inoculum_index_column, Ntypes)
        
        if len(frequencies) == 0:
            print(f"No data for configuration: {configuration}")
            continue
        
        mle_theta = []
        dist_s = []
        
        if len(frequencies) > 0:
            for i, freq in enumerate(frequencies):
            
                # get the starting frequencies corresponding to the inoculum of the sample
                meta = metadata[i]
                start_freq = starting_frequencies[np.searchsorted(inoculum_index, meta[3])]
            
                best_theta, dist_ = find_best_theta(freq, start_freq, theta_range, distribution_model, distance_function, parameter_name, step)
                mle_theta.append(best_theta)
                dist_s.append(dist_)

        # Compute mean theta and mean distance across all replicates
        mean_mle_theta = np.mean(mle_theta)
        mean_distance = np.mean(dist_s)  # Performance metric

        # Store both all best theta values, mean theta, and mean performance
        theta_values[tuple(configuration)] = {
            "theta_list": mle_theta,  # List of all best theta values
            "mean_theta": mean_mle_theta,  # Mean of best theta values
            "mean_distance": mean_distance  # Mean distance (performance metric)
        }

        # Store and print results (only mean values)
        results.append((configuration, mean_mle_theta, mean_distance))
        print(f"Modèle: {model_name}, Configuration: {configuration}, Mean theta: {mean_mle_theta:.2f}, Performance (Mean Distance): {mean_distance:.4f}")

    return theta_values  # Return both detailed and summarized theta values
  

with open('Analysis_output/tab_tot.csv', newline='') as file:
    data = list(csv.reader(file, delimiter=','))

with open('Analysis_output/tab_inoc.csv', newline='') as file:
    inoc = list(csv.reader(file, delimiter=','))

# Functions to plot the Hill curves
def hill_number(V, q):
    """Compute Hill numbers for a given frequency vector V and order q."""
    if q == 1:
        return np.exp(-np.sum(V[V > 0] * np.log(V[V > 0])))
    elif q == 0:
        return np.sum(V > 0)
    else:
        return (np.sum(V ** q)) ** (1 / (1 - q))

def calculate_distance_per_q(V1, V2, Q):
    """Compute the diversity-based distance for each q."""
    distances = []
    for q in Q:
        Hq1 = hill_number(V1, q)
        Hq2 = hill_number(V2, q)
        distance = abs(Hq1 - Hq2) / (Hq1 + Hq2) if (Hq1 + Hq2) != 0 else 0
        distances.append(distance)
    return distances

def generate_best_frequency(K, theta, starting_frequencies, distribution_model, real_frequency, num_sim=100):
    """Generate 100 frequency vectors and return the closest one to the real frequency."""
    simulated_vectors = [sample_theta(K, (theta, theta), starting_frequencies, distribution_model) for _ in range(num_sim)]
    distances = [np.mean(calculate_distance_per_q(V, real_frequency, np.arange(0, 4.1, 0.1))) for V in simulated_vectors]
    best_index = np.argmin(distances)  # Index of lowest distance
    return simulated_vectors[best_index]  # Return the best-matching frequency

def plot_hill_curves(real_frequencies, best_simulated_frequencies):
    """Plot Hill curves for real and best simulated frequencies in separate subplots."""
    Q = np.arange(0, 4.1, 0.1)
    num_freqs = len(real_frequencies)
    color_map = plt.colormaps["tab10"]

    fig, axes = plt.subplots(1, 2, figsize=(12, 6), sharey=True)

    # Plot each real frequency in the left subplot
    for i, freq in enumerate(real_frequencies):
        hill_curve = [hill_number(freq, q) for q in Q]
        color = color_map(i / max(1, num_freqs - 1))
        axes[0].plot(Q, hill_curve, color=color, alpha=0.9, linewidth=2)

    # Plot each best simulated frequency in the right subplot
    for i, best_freq in enumerate(best_simulated_frequencies):
        hill_curve = [hill_number(best_freq, q) for q in Q]
        color = color_map(i / max(1, num_freqs - 1))
        axes[1].plot(Q, hill_curve, color=color, alpha=0.9, linewidth=2)

    # Formatting subplots
    axes[0].set_title("Observed Frequencies")
    axes[1].set_title("Best Simulated Frequencies")
    for ax in axes:
        ax.set_xlabel("q")
        ax.set_ylabel("Hill Number")
        ax.set_ylim(0, 120)

    plt.tight_layout()
    plt.show()

def process_with_plots(data, inoc, lconfigurations, distribution_model, theta_values):
    """Process configurations, generate the best frequency vectors, and plot Hill curves and global mean distance."""
    Q = np.arange(0, 4.1, 0.1)
    global_mean_distances = np.zeros(len(Q))  # Accumulate mean distances over all configurations
    total_samples = 0  # Count total frequencies processed
    all_configurations_mean_distances = [] # Mean distances for all individual configuration
    configuration_mean_distances = np.zeros(len(Q))  # Accumulate mean distances within each configuration
    configuration_samples = 0 # Count frequencies processed within each configuration

    all_real_frequencies = {}  # Store real frequencies for all configurations
    all_best_simulated_frequencies = {}  # Store best simulated frequencies for all configurations
    
    starting_frequencies, inoculum_index = read_starting_freq(inoc, corresponding_inoculum_tags_range, corresponding_inoculum_index_column, inoculum_validation_column, Ntypes)
    
    
    for configuration in lconfigurations:
        real_frequencies, metadata = read_file(data, configuration, tags_range, configuration_column, validation_column, sample_index_column, time_of_sampling_column, bacterial_load_column, inoculum_index_column, Ntypes)
        
        if len(real_frequencies) == 0:
            print(f"No data for configuration: {configuration}")
            continue

        K = len(real_frequencies[0])  # Number of species/types
        config_key = tuple(configuration)
        theta_list = theta_values.get(config_key, {}).get("theta_list", None)

        if theta_list is None or len(theta_list) != len(real_frequencies):
            print(f"Missing or mismatched theta values for configuration: {configuration}")
            continue

        best_simulated_frequencies = []
        i = 0
        
        if len(real_frequencies) > 0:
            for real_freq, theta in zip(real_frequencies, theta_list):
            
                # get the starting frequencies corresponding to the inoculum of the sample
                meta = metadata[i]
                start_freq = starting_frequencies[np.searchsorted(inoculum_index, meta[3])]
                i = i + 1
                
                best_freq = generate_best_frequency(K, theta, start_freq, distribution_model, real_freq)
                best_simulated_frequencies.append(best_freq)

                # Compute distance for this frequency
                distances = calculate_distance_per_q(real_freq, best_freq, Q)
                global_mean_distances += np.array(distances)
                total_samples += 1
            
                # Distance retriever for the configuration
                configuration_mean_distances += np.array(distances)
                configuration_samples += 1

        # Compute and append distance for the configuration
        all_configurations_mean_distances.append(configuration_mean_distances / configuration_samples)

        # Reset configuration distance and sample count
        configuration_mean_distances = np.zeros(len(Q))
        configuration_samples = 0
        
        # Store results for this configuration
        all_real_frequencies[config_key] = real_frequencies
        all_best_simulated_frequencies[config_key] = best_simulated_frequencies

        # Plot Hill curves for this configuration
        plot_hill_curves(real_frequencies, best_simulated_frequencies)

    # Compute final averaged distances
    if total_samples > 0:
        global_mean_distances /= total_samples

    # Plot global error figure
    plt.figure(figsize=(8, 6))
    plt.plot(Q, global_mean_distances, color="black", linewidth=2)
    plt.xlabel("q")
    plt.ylabel("Mean Distance")
    plt.title("Mean Distance Averaged Over All Configurations")
    plt.show()

    return Q, global_mean_distances, all_real_frequencies, all_best_simulated_frequencies, all_configurations_mean_distances

# functions to Save and load the results
def save_pickle(theta_values, filename="theta_values1.pkl"):
    """
    Save theta_values dictionary as a Pickle file.
    """
    with open(filename, "wb") as f:
        pickle.dump(theta_values, f)
    print(f"Saved to {filename}")

def load_pickle(filename="theta_values1.pkl"):
    """
    Load theta_values dictionary from a Pickle file.
    """
    with open(filename, "rb") as f:
        return pickle.load(f)

```

## Multinomial model

```{python}
# define the model
def distribution_modelM1(K, theta, starting_frequencies):
    Nb = theta
    return np.random.multinomial(Nb, starting_frequencies)
```

```{python}

# define the range for parameter values
theta_range = (1, 200)
step = 1

print('Model performance')
# Run the processing function
theta_values1 = process_configurations(data, inoc, lconfigurations, distribution_modelM1, theta_range, step, quantile_distance, parameter_name="Nb", model_name = "STAMP Model")
save_pickle(theta_values1,"Models_output_correct_inoc/theta_values1.pkl")
```

```{python}
theta_values1 = load_pickle("Models_output_correct_inoc/theta_values1.pkl")  # Loads it back
Q1, D1, RF, BF1, all_D1 = process_with_plots(data, inoc, lconfigurations, distribution_modelM1, theta_values1)
np.save("Models_output_correct_inoc/Q1.npy", Q1)
np.save("Models_output_correct_inoc/D1.npy", D1)
np.save("Models_output_correct_inoc/all_D1.npy", all_D1)
save_pickle(RF, filename="Models_output_correct_inoc/RF.pkl")
save_pickle(BF1, filename="Models_output_correct_inoc/BF1.pkl")

```

## Constant rate model CR

```{python}
# define the model
def distribution_modelM3(K, theta, starting_frequencies):
    mu = theta
    r = 1
    T = 10
    # simulate entry times :
    tau = [ np.random.exponential( 1/mu ) ]
    while tau[-1] < T:
        tau += [ tau[-1] + np.random.exponential( 1/mu ) ]
    qt_prov = [ np.exp( r * (T-tau[i]) ) for i in range(len(tau)) ]
    res = sample_tags( qt_prov, K, starting_frequencies )
    return np.sort(res)[::-1]

def sample_tags(prop, Ntags, starting_frequencies):
    prop_tags = np.zeros( Ntags )
    for pp in prop:
        tag = np.random.choice( range(Ntags), p = starting_frequencies) 
        prop_tags[ tag ] += pp
    return prop_tags

```

```{python}
# define the range for parameter values
theta_range = (1, 50)
step = 1

print('Model performance')
# Run the processing function
theta_values3 = process_configurations(data, inoc, lconfigurations, distribution_modelM3, theta_range, step, quantile_distance, parameter_name="Nb", model_name = "CR1 Model")

save_pickle(theta_values3,"Models_output_correct_inoc/theta_values3.pkl")
```

```{python}
theta_values3 = load_pickle("Models_output_correct_inoc/theta_values3.pkl")
Q3, D3, RF, BF3, all_D3 = process_with_plots(data, inoc, lconfigurations, distribution_modelM3, theta_values3)
np.save("Models_output_correct_inoc/Q3.npy", Q3)
np.save("Models_output_correct_inoc/D3.npy", D3)
np.save("Models_output_correct_inoc/all_D3.npy", all_D3)
save_pickle(BF3, filename="Models_output_correct_inoc/BF3.pkl")
```

## Bolthausen-Sznitman coalescent model BS

```{python}
#**Step 1**: construct the initial random recursive tree.
def update_edges( l, k ):
    current = k
    while current != 0:
        current = l[current][0]
        l[current][2] += [k]


def init_bs( n ):
    l_edges = [ [0,[0],[]] ]
    new_edge = [0,[0],[]]
    for k in range(1,n):
        parent_new = np.random.randint(k)
        new_edge = [ parent_new, [k], []]
        l_edges += [ new_edge ]
        update_edges( l_edges, k )
    return l_edges
#**Step 2**: each edge is "lifted" at rate $1$ (in the terminology of *Berestycki 2009*).
def step_bs( tree, k ): # lift edge k of tree
    loss = 0
    parent = tree[k][0]
    tree[parent][1] += [k] + tree[k][2] # the edge is lifted to the parent
    for i in [k] + tree[k][2]:
        if tree[i][1][0] != -1:
            tree[i][1] = [-1] # unactivated
            loss += 1
    return tree, loss # tree : resulting tree; loss: number of blocks lost
#**Step 3:** the following function runs a Bolthausen-Sznitman coalescent for a time [t], starting from an initial tree [tree] with specified number of blocks [nblocks]).

def run_bs( nblocks, t, tree ):

    t_cur = np.random.exponential(1/nblocks)

    while t_cur < t and nblocks > 1:


        i = np.random.randint( 1, nblocks ) # choose the edge to be lifted
        edge = 0
        while i > -1:
            if tree[ edge ][1][0] != -1:
                edge += 1
                i -= 1
            else:
                edge += 1
        tree, loss = step_bs( tree, edge - 1 )
        nblocks -= loss

        t_cur += np.random.exponential(1/nblocks)

    return tree, nblocks


#run_bs(10000,0.5)
def display( tree ):
    for k in range( len(tree) ):
        if tree[k][1][0] != -1:
            print(tree[k][1], end='  ')
    print("")
def get_prop_bs( N, t ):
    tree,_ = run_bs( N, t, init_bs( N ) )
    prop = [] # get proportions
    for e in tree:
        if e[1][0] != -1:
            prop += [ len(e[1]) ]
    prop = prop / np.sum( prop )
    return prop
def sample_tags(prop, Ntags, starting_frequencies):
    prop_tags = np.zeros( Ntags )
    for pp in prop:
        tag = np.random.choice( range(Ntags), p = starting_frequencies) 
        prop_tags[ tag ] += pp
    return prop_tags
```

```{python}
# define the model
def distribution_modelM4(K, theta, starting_frequencies):
    t = theta
    return sample_tags( get_prop_bs(2000, t), K, starting_frequencies)
```

```{python}
# define the range for parameter values
theta_range = (0.05, 1)
step = 0.05

print('Model performance')
# Run the processing function
theta_values4 = process_configurations(data, inoc, lconfigurations, distribution_modelM4, theta_range, step, quantile_distance, parameter_name="t", model_name = "BS Model")
save_pickle(theta_values4,"Models_output_correct_inoc/theta_values4.pkl")
```

```{python}
theta_values4 = load_pickle("Models_output_correct_inoc/theta_values4.pkl")
Q4, D4, RF, BF4, all_D4 = process_with_plots(data, inoc, lconfigurations, distribution_modelM4, theta_values4)
np.save("Models_output_correct_inoc/Q4.npy", Q4)
np.save("Models_output_correct_inoc/D4.npy", D4)
np.save("Models_output_correct_inoc/all_D4.npy", all_D4)
save_pickle(BF4, filename="Models_output_correct_inoc/BF4.pkl")
```

## Kingman coalescent

```{python}
def run_km( t, prop = [] ):

    nblocks = len(prop)
    t_cur = 0

    if nblocks > 1:
        t_cur = np.random.exponential(2/(nblocks * (nblocks-1)))

    while t_cur < t and nblocks > 1:
        t_cur += np.random.exponential(2/(nblocks * (nblocks-1)))

        i,j=[0,0]
        while i == j:
            i,j = np.random.randint( 0, nblocks, 2)
        prop[i] += prop[j] # j gets to i
        prop.pop(j) # remove j
        nblocks -= 1
    return prop
def sample_tags(prop, Ntags, starting_frequencies):
    prop_tags = np.zeros( Ntags )
    for pp in prop:
        tag = np.random.choice( range(Ntags), p = starting_frequencies) 
        prop_tags[ tag ] += pp
    return prop_tags

```

```{python}
# define the model
def distribution_modelM5(K, theta, starting_frequencies):
    t = theta
    return sample_tags( run_km(t, [1/1000] * 1000), K, starting_frequencies)
```

```{python}
# define the range for parameter values
theta_range = (0.05, 1)
step = 0.05
print('Model performance')
# Run the processing function
theta_values5 = process_configurations(data, inoc, lconfigurations, distribution_modelM5, theta_range, step, quantile_distance, parameter_name="t", model_name = "K Model")
save_pickle(theta_values5,"Models_output_correct_inoc/theta_values5.pkl")
```

```{python}
theta_values5 = load_pickle("Models_output_correct_inoc/theta_values5.pkl")
Q5, D5, RF, BF5, all_D5 = process_with_plots(data, inoc, lconfigurations, distribution_modelM5, theta_values5)
np.save("Models_output_correct_inoc/Q5.npy", Q5)
np.save("Models_output_correct_inoc/D5.npy", D5)
np.save("Models_output_correct_inoc/all_D5.npy", all_D5)
save_pickle(BF5, filename="Models_output_correct_inoc/BF5.pkl")
```

## Increasing entry rate model IR

```{python}

# NHPP inverse cumulative intensity function for λ(t) = mu * exp(k * t)
def nhpp_exponential_inverse(s, mu, k):
    return (1.0 / k) * np.log(k * s / mu + 1)

def distribution_modelM6(K, theta, starting_frequencies):
    mu = theta
    k = 0.78
    Ntypes = K
    T = 10

    tau = []
    s = 0.0
    while True:
        s += np.random.exponential(1.0)  # Homogeneous Poisson interarrival
        t_candidate = nhpp_exponential_inverse(s, mu, k)
        if t_candidate > T:
            break
        tau.append(t_candidate)

    qt_prov = [np.exp((T - t)) for t in tau]

    res = sample_tags( qt_prov, K, starting_frequencies )

    return np.sort(res)[::-1]

def sample_tags(prop, Ntags, starting_frequencies):
    prop_tags = np.zeros( Ntags )
    for pp in prop:
        tag = np.random.choice( range(Ntags), p = starting_frequencies) 
        prop_tags[ tag ] += pp
    return prop_tags

```

```{python}
# define the range for parameter values
theta_range = (0.05, 1)
step = 0.05

print('Model performance')
# Run the processing function
theta_values6 = process_configurations(data, inoc, lconfigurations, distribution_modelM6, theta_range, step, quantile_distance, parameter_name="mu", model_name = "VR Model")
save_pickle(theta_values6,"Models_output_correct_inoc/theta_values6.pkl")
```

```{python}
theta_values6 = load_pickle("Models_output_correct_inoc/theta_values6.pkl")
Q6, D6, RF, BF6, all_D6 = process_with_plots(data, inoc, lconfigurations, distribution_modelM6, theta_values6)
np.save("Models_output_correct_inoc/Q6.npy", Q6)
np.save("Models_output_correct_inoc/D6.npy", D6)
np.save("Models_output_correct_inoc/all_D6.npy", all_D6)
save_pickle(BF6, filename="Models_output_correct_inoc/BF6.pkl")
```

# III. Some graphs

## Helper function

```{python, fig.width=10, fig.height=6}

# =======================
# Load pickle files
# =======================

def load_pickle(filename):
    if not os.path.exists(filename):
        raise FileNotFoundError(f"'{filename}' not found in repository.")
    with open(filename, "rb") as f:
        return pickle.load(f)

# Load data
RF = load_pickle("Models_output_correct_inoc/RF.pkl")
BF1 = load_pickle("Models_output_correct_inoc/BF1.pkl")
BF3 = load_pickle("Models_output_correct_inoc/BF3.pkl")
BF4 = load_pickle("Models_output_correct_inoc/BF4.pkl")
BF6 = load_pickle("Models_output_correct_inoc/BF6.pkl")
BF5 = load_pickle("Models_output_correct_inoc/BF5.pkl")
theta_values = load_pickle("Models_output_correct_inoc/theta_values1.pkl")  # Adjust the file name if needed


```

## Error curves

```{python, fig.width=8, fig.height=6}

# Initialize plot
plt.figure(figsize=(8, 6))

i = 1
Q = np.load(f"Models_output_correct_inoc/Q{i}.npy")  # Load Qi
D = np.load(f"Models_output_correct_inoc/D{i}.npy")  # Load Di
plt.plot(Q, D, linewidth=2, label="Model $\mathcal{{M}}_{STAMP}$")  # Plot with label
i = 3
Q = np.load(f"Models_output_correct_inoc/Q{i}.npy")  # Load Qi
D = np.load(f"Models_output_correct_inoc/D{i}.npy")  # Load Di
plt.plot(Q, D, linewidth=2, label="Model $\mathcal{{M}}_{CR}$")  # Plot with label
i = 4
Q = np.load(f"Models_output_correct_inoc/Q{i}.npy")  # Load Qi
D = np.load(f"Models_output_correct_inoc/D{i}.npy")  # Load Di
plt.plot(Q, D, linewidth=2, label="Model $\mathcal{{M}}_{BS}$")  # Plot with label
i = 6
Q = np.load(f"Models_output_correct_inoc/Q{i}.npy")  # Load Qi
D = np.load(f"Models_output_correct_inoc/D{i}.npy")  # Load Di
plt.plot(Q, D, linewidth=2, label="Model $\mathcal{{M}}_{IR}$")  # Plot with label
i = 5
Q = np.load(f"Models_output_correct_inoc/Q{i}.npy")  # Load Qi
D = np.load(f"Models_output_correct_inoc/D{i}.npy")  # Load Di
plt.plot(Q, D, linewidth=2, label="Model $\mathcal{{M}}_{K}$")  # Plot with label

# Formatting the plot
plt.xlabel("$q$", fontsize=16)
plt.ylabel("Mean relative error", fontsize=16)
plt.legend(fontsize=14)
plt.xticks(fontsize=16)
plt.yticks(fontsize=16)
plt.grid()
plt.tight_layout()
#Save the figure as a PNG file
#plt.savefig("mean_error.png", dpi=300, bbox_inches='tight')

plt.show()
```

```{python, fig.width=6, fig.height=26}

# Plot mean error for each condition
# Load the saved data
RF = load_pickle("Models_output_correct_inoc/RF.pkl")

# Extract experiment names from RF
experiment_names = list(RF.keys())

Q = np.arange(0, 4.1, 0.1)

# Create figure with subplots (7 rows, 1 columns)
fig, axes = plt.subplots(7, 1, figsize=(6, 32), sharex=True, sharey=True)

# Iterate over configurations and plot Hill curves
for j, exp_name in enumerate(experiment_names):

  i = 1
  D = np.load(f"Models_output_correct_inoc/all_D{i}.npy")
  D = D[j]
  axes[j].plot(Q, D, alpha=0.8, linewidth=2, label="Model $\mathcal{{M}}_{STAMP}$")

  i = 3
  D = np.load(f"Models_output_correct_inoc/all_D{i}.npy")
  D = D[j]
  axes[j].plot(Q, D, alpha=0.8, linewidth=2, label="Model $\mathcal{{M}}_{CR}$")

  i = 4
  D = np.load(f"Models_output_correct_inoc/all_D{i}.npy")
  D = D[j]
  axes[j].plot(Q, D, alpha=0.8, linewidth=2, label="Model $\mathcal{{M}}_{BS}$")

  i = 6
  D = np.load(f"Models_output_correct_inoc/all_D{i}.npy")
  D = D[j]
  axes[j].plot(Q, D, alpha=0.8, linewidth=2, label="Model $\mathcal{{M}}_{IR}$")
  
  i = 5
  D = np.load(f"Models_output_correct_inoc/all_D{i}.npy")
  D = D[j]
  axes[j].plot(Q, D, alpha=0.8, linewidth=2, label="Model $\mathcal{{M}}_{IR}$")

  #SV Set experiment label
  axes[j].set_ylabel(f"{exp_name}", fontsize=12)
  axes[j].grid()
  axes[j].legend(fontsize=14)

axes[j].set_xlabel("q", fontsize=12)

plt.tight_layout()
plt.show()

```

## Extraction of the csv tables with theta, mu and r

```{python}
# STAMP Model This code builds a csv file with the data. The last column corresponds to theta

with open('Analysis_output/tab_tot.csv', newline='') as file:
    data = list(csv.reader(file, delimiter=','))

def read_file(data, configuration, tags_range, configuration_column, validation_column, time_of_sampling_column, bacterial_load_column, sample_weight_column, sample_index_column, Ntypes):
    frequencies = []
    times = []
    loads = []
    weights = []
    indexes = []

    for row in data:
        if row[configuration_column] == configuration[0]: # Check condition
            if row[validation_column] == '1': # Check sequencing validation
                counts = [int(row[j]) for j in range(tags_range, Ntypes + tags_range)]  # Adjust for weight column
                counts_sum = sum(counts)
                if counts_sum > 0:
                    frequencies.append([c / counts_sum for c in counts])
                    times.append(float(row[time_of_sampling_column]))
                    loads.append(float(row[bacterial_load_column]))
                    weights.append(float(row[sample_weight_column]))
                    indexes.append(row[sample_index_column])

    return np.array(frequencies), np.array(times), np.array(loads), np.array(weights), np.array(indexes)


def load_pickle(filename):
    with open(filename, "rb") as f:
        return pickle.load(f)

# Load theta values
theta_values = pickle.load(open('Models_output_correct_inoc/theta_values1.pkl', 'rb'))

# Load precomputed rates (r) per config
# Here we recompute the rates so we can align them


# Read CSV into list
with open('Analysis_output/tab_tot.csv', newline='') as file:
    data = list(csv.reader(file))

# Compute r per valid row to match theta
# Compute rates for all configs
rates = {}
all_indexes = {}
for configuration in lconfigurations:
    config_key = tuple(configuration)
    frequencies, times, loads, weights, indexes = read_file(data, configuration, tags_range, configuration_column, validation_column, time_of_sampling_column, bacterial_load_column, sample_weight_column, sample_index_column, Ntypes)
    
    if config_key in theta_values:
        theta_list = np.array(theta_values[config_key]['theta_list'])
        p1 = np.max(frequencies, axis=1)
        r_ = np.full_like(p1, np.nan)
        mask = (p1>0) & (loads>0) & (times!=0) & (theta_list>0)
        r_[mask] = (np.log(p1[mask]*loads[mask]*weights[mask])+(1.0/theta_list[mask]))/times[mask]
        rates[config_key] = r_
        all_indexes[config_key] = indexes


# Prepare output file with header
#input_file = "Models_output/Data_py.csv"
input_file = "Analysis_output/tab_tot.csv"
output_file = "Models_output_correct_inoc/data_and_estimators_model_STAMP.csv"
with open(input_file, newline='') as infile:
    reader = list(csv.reader(infile))

# Header with only theta
header = ['index'] + ['theta_STAMP']

# Track per-config index to select correct theta
idx_per_config = {tuple(c): 0 for c in lconfigurations}

with open(output_file, mode='w', newline='') as outfile:
    writer = csv.writer(outfile)
    writer.writerow(header)
    for row in reader[1:]:
        configuration = (row[configuration_column],)
        index_val = ""
        theta_val = ""  # default blank
        if configuration in theta_values and row[validation_column] == '1':
            i = idx_per_config[configuration]
            theta_list = np.array(theta_values[configuration]['theta_list'])
            index_list = all_indexes[configuration]
            if i < len(theta_list):
                index_val = index_list[i]
                theta_val = theta_list[i]
                idx_per_config[configuration] += 1
        writer.writerow([index_val] + [theta_val])

print(f"OK Saved file to {output_file}")

```

```{python}
# CR Model This code builds a csv file with the data. The last three columns correspond to theta, r, mu

with open('Analysis_output/tab_tot.csv', newline='') as file:
    data = list(csv.reader(file, delimiter=','))

def read_file(data, configuration, tags_range, configuration_column, validation_column, time_of_sampling_column, bacterial_load_column, sample_weight_column, sample_index_column, Ntypes):
    frequencies = []
    times = []
    loads = []
    weights = []
    indexes = []

    for row in data:
        if row[configuration_column] == configuration[0]: # Check condition
            if row[validation_column] == '1': # Check sequencing validation
                counts = [int(row[j]) for j in range(tags_range, Ntypes + tags_range)]  # Adjust for weight column
                counts_sum = sum(counts)
                if counts_sum > 0:
                    frequencies.append([c / counts_sum for c in counts])
                    times.append(float(row[time_of_sampling_column]))
                    loads.append(float(row[bacterial_load_column]))
                    weights.append(float(row[sample_weight_column]))
                    indexes.append(row[sample_index_column])

    return np.array(frequencies), np.array(times), np.array(loads), np.array(weights), np.array(indexes)


def load_pickle(filename):
    with open(filename, "rb") as f:
        return pickle.load(f)

# Load theta values
theta_values = pickle.load(open('Models_output_correct_inoc/theta_values3.pkl', 'rb'))

# Load precomputed rates (r) per config
# Here we recompute the rates so we can align them


# Read CSV into list
with open('Analysis_output/tab_tot.csv', newline='') as file:
    data = list(csv.reader(file))

# Compute r per valid row to match theta
# Compute rates for all configs
rates = {}
all_indexes = {}
for configuration in lconfigurations:
    config_key = tuple(configuration)
    frequencies, times, loads, weights, indexes = read_file(data, configuration, tags_range, configuration_column, validation_column, time_of_sampling_column, bacterial_load_column, sample_weight_column, sample_index_column, Ntypes)
    
    if config_key in theta_values:
        theta_list = np.array(theta_values[config_key]['theta_list'])
        p1 = np.max(frequencies, axis=1)
        r_ = np.full_like(p1, np.nan)
        mask = (p1>0) & (loads>0) & (times!=0) & (theta_list>0)
        r_[mask] = (np.log(p1[mask]*loads[mask]*weights[mask])+(1.0/theta_list[mask]))/times[mask]
        rates[config_key] = r_
        all_indexes[config_key] = indexes


# Prepare output file with header
#input_file = "Models_output/Data_py.csv"
input_file = "Analysis_output/tab_tot.csv"
output_file = "Models_output_correct_inoc/data_and_estimators_model_CR.csv"
with open(input_file, newline='') as infile:
    reader = list(csv.reader(infile))

# Header with only theta
header = ['index', 'theta_CR', 'r', 'mu']

# Track per-config index to select correct theta
idx_per_config = {tuple(c): 0 for c in lconfigurations}

with open(output_file, mode='w', newline='') as outfile:
    writer = csv.writer(outfile)
    writer.writerow(header)
    for row in reader[1:]:
        configuration = (row[configuration_column],)
        index_val = ""
        theta_val = ""
        r_val = ""
        mu_val = ""
        if configuration in rates and row[validation_column] == '1':
            i = idx_per_config[configuration]
            theta_list = np.array(theta_values[configuration]['theta_list'])
            r_list = rates[configuration]
            index_list = all_indexes[configuration]
            if i < len(theta_list):
                index_val = index_list[i]
                theta_val = theta_list[i]
                r_val = r_list[i]
                mu_val = theta_val * r_val
                idx_per_config[configuration] += 1
        writer.writerow([index_val, theta_val, r_val, mu_val])

print(f"OK Saved file to {output_file}")

```

```{python}
# IR Model This code builds a csv file with the data. The last three columns correspond to theta, r, mu

k = 0.78

with open('Analysis_output/tab_tot.csv', newline='') as file:
    data = list(csv.reader(file, delimiter=','))

def read_file(data, configuration, tags_range, configuration_column, validation_column, time_of_sampling_column, bacterial_load_column, sample_weight_column, sample_index_column, Ntypes):
    frequencies = []
    times = []
    loads = []
    weights = []
    indexes = []

    for row in data:
        if row[configuration_column] == configuration[0]: # Check condition
            if row[validation_column] == '1': # Check sequencing validation
                counts = [int(row[j]) for j in range(tags_range, Ntypes + tags_range)]  # Adjust for weight column
                counts_sum = sum(counts)
                if counts_sum > 0:
                    frequencies.append([c / counts_sum for c in counts])
                    times.append(float(row[time_of_sampling_column]))
                    loads.append(float(row[bacterial_load_column]))
                    weights.append(float(row[sample_weight_column]))
                    indexes.append(row[sample_index_column])

    return np.array(frequencies), np.array(times), np.array(loads), np.array(weights), np.array(indexes)


def load_pickle(filename):
    with open(filename, "rb") as f:
        return pickle.load(f)

# Load theta values
theta_values = pickle.load(open('Models_output_correct_inoc/theta_values6.pkl', 'rb'))

# Load precomputed rates (r) per config
# Here we recompute the rates so we can align them


# Read CSV into list
with open('Analysis_output/tab_tot.csv', newline='') as file:
    data = list(csv.reader(file))

# Compute r per valid row to match theta
# Compute rates for all configs
rates = {}
all_indexes = {}
for configuration in lconfigurations:
    config_key = tuple(configuration)
    frequencies, times, loads, weights, indexes = read_file(data, configuration, tags_range, configuration_column, validation_column, time_of_sampling_column, bacterial_load_column, sample_weight_column, sample_index_column, Ntypes)
    
    if config_key in theta_values:
        theta_list = np.array(theta_values[config_key]['theta_list'])
        p1 = np.max(frequencies, axis=1)
        r_ = np.full_like(p1, np.nan)
        mask = (p1>0) & (loads>0) & (times!=0) & (theta_list>0)
        r_[mask] = (np.log(p1[mask]*loads[mask]*weights[mask])+(1/k * np.log(1 + np.log(2) * k / theta_list[mask])))/times[mask]
        rates[config_key] = r_
        all_indexes[config_key] = indexes


# Prepare output file with header
#input_file = "Models_output/Data_py.csv"
input_file = "Analysis_output/tab_tot.csv"
output_file = "Models_output_correct_inoc/data_and_estimators_model_IR.csv"
with open(input_file, newline='') as infile:
    reader = list(csv.reader(infile))

# Header with only theta
header = ['index', 'theta_IR', 'r', 'mu']

# Track per-config index to select correct theta
idx_per_config = {tuple(c): 0 for c in lconfigurations}

with open(output_file, mode='w', newline='') as outfile:
    writer = csv.writer(outfile)
    writer.writerow(header)
    for row in reader[1:]:
        configuration = (row[configuration_column],)
        index_val = ""
        theta_val = ""
        r_val = ""
        mu_val = ""
        if configuration in rates and row[validation_column] == '1':
            i = idx_per_config[configuration]
            theta_list = np.array(theta_values[configuration]['theta_list'])
            r_list = rates[configuration]
            index_list = all_indexes[configuration]
            if i < len(theta_list):
                index_val = index_list[i]
                theta_val = theta_list[i]
                r_val = r_list[i]
                mu_val = theta_val * r_val
                idx_per_config[configuration] += 1
        writer.writerow([index_val, theta_val, r_val, mu_val])

print(f"OK Saved file to {output_file}")
```

```{python}
# BS Model This code builds a csv file with the data. The last column corresponds to theta

with open('Analysis_output/tab_tot.csv', newline='') as file:
    data = list(csv.reader(file, delimiter=','))

def read_file(data, configuration, tags_range, configuration_column, validation_column, time_of_sampling_column, bacterial_load_column, sample_weight_column, sample_index_column, Ntypes):
    frequencies = []
    times = []
    loads = []
    weights = []
    indexes = []

    for row in data:
        if row[configuration_column] == configuration[0]: # Check condition
            if row[validation_column] == '1': # Check sequencing validation
                counts = [int(row[j]) for j in range(tags_range, Ntypes + tags_range)]  # Adjust for weight column
                counts_sum = sum(counts)
                if counts_sum > 0:
                    frequencies.append([c / counts_sum for c in counts])
                    times.append(float(row[time_of_sampling_column]))
                    loads.append(float(row[bacterial_load_column]))
                    weights.append(float(row[sample_weight_column]))
                    indexes.append(row[sample_index_column])

    return np.array(frequencies), np.array(times), np.array(loads), np.array(weights), np.array(indexes)


def load_pickle(filename):
    with open(filename, "rb") as f:
        return pickle.load(f)

# Load theta values
theta_values = pickle.load(open('Models_output_correct_inoc/theta_values4.pkl', 'rb'))

# Load precomputed rates (r) per config
# Here we recompute the rates so we can align them


# Read CSV into list
with open('Analysis_output/tab_tot.csv', newline='') as file:
    data = list(csv.reader(file))

# Compute r per valid row to match theta
# Compute rates for all configs
rates = {}
all_indexes = {}
for configuration in lconfigurations:
    config_key = tuple(configuration)
    frequencies, times, loads, weights, indexes = read_file(data, configuration, tags_range, configuration_column, validation_column, time_of_sampling_column, bacterial_load_column, sample_weight_column, sample_index_column, Ntypes)
    
    if config_key in theta_values:
        theta_list = np.array(theta_values[config_key]['theta_list'])
        p1 = np.max(frequencies, axis=1)
        r_ = np.full_like(p1, np.nan)
        mask = (p1>0) & (loads>0) & (times!=0) & (theta_list>0)
        r_[mask] = (np.log(p1[mask]*loads[mask]*weights[mask])+(1.0/theta_list[mask]))/times[mask]
        rates[config_key] = r_
        all_indexes[config_key] = indexes


# Prepare output file with header
input_file = "Analysis_output/tab_tot.csv"
output_file = "Models_output_correct_inoc/data_and_estimators_model_BS.csv"
with open(input_file, newline='') as infile:
    reader = list(csv.reader(infile))

# Header with only theta
header = ['index'] + ['1 / theta_BS']

# Track per-config index to select correct theta
idx_per_config = {tuple(c): 0 for c in lconfigurations}

with open(output_file, mode='w', newline='') as outfile:
    writer = csv.writer(outfile)
    writer.writerow(header)
    for row in reader[1:]:
        configuration = (row[configuration_column],)
        index_val = ""
        theta_val = ""  # default blank
        if configuration in theta_values and row[validation_column] == '1':
            i = idx_per_config[configuration]
            theta_list = np.array(theta_values[configuration]['theta_list'])
            index_list = all_indexes[configuration]
            if i < len(theta_list):
                index_val = index_list[i]
                theta_val = theta_list[i]
                idx_per_config[configuration] += 1
        writer.writerow([index_val] + [theta_val])

print(f"OK Saved file to {output_file}")

```

## Visualisation of models outputs

Visualisation of the output parameters theta, mu and r

```{r Graph}

tab = tab_tot

tab_temp = read.csv("Models_output_correct_inoc/data_and_estimators_model_STAMP.csv", header= TRUE, check.names= FALSE)
tab$Theta_STAMP = tab_temp$theta_STAMP
tab_temp = read.csv("Models_output_correct_inoc/data_and_estimators_model_CR.csv", header= TRUE, check.names= FALSE)
tab$Theta_CR = tab_temp$theta_CR
tab$Mu_CR = tab_temp$mu
tab$r_CR = tab_temp$r
tab_temp = read.csv("Models_output_correct_inoc/data_and_estimators_model_BS.csv", header= TRUE, check.names= FALSE)
tab$Theta_BS = tab_temp$`1 / theta_BS`
tab$Theta_BS = 1 / tab$Theta_BS
tab_temp = read.csv("Models_output_correct_inoc/data_and_estimators_model_IR.csv", header= TRUE, check.names= FALSE)
tab$Theta_IR = tab_temp$theta
tab$Mu_IR = tab_temp$mu
tab$r_IR = tab_temp$r

tab = tab %>%
  filter(Zone == "Root") %>%
  filter(`PCR2 validation on gel` > 0)


####################
# Visualisation of the STAMP model (Nb)
####################

# Kruskal-wallis statistical test
test = Graph_kruskal(tab= tab, y= "Theta_STAMP", x= "Condition2")

# Display helper
values = tab %>% 
  group_by(Condition2, .drop= FALSE) %>% 
  summarise(mean = mean(Theta_STAMP), sd = sd(Theta_STAMP))


# Graph
ggplot(data= tab, aes(x= Condition2, y= Theta_STAMP)) +
  
  geom_boxplot(outlier.shape= NA, width=0.9, fill= "lightblue") +
  geom_point(aes(colour= Replicate),
                  position= position_jitterdodge(jitter.width= 0.3, dodge.width= 0.4), alpha= 1) +
  
  scale_colour_manual(values= c("#E41A1C", "#339900", "#FFD92F")) +
  geom_text(data = values, aes(label = test$groups, y = 1 , x = Condition2), vjust = -22, inherit.aes = FALSE) +
  scale_x_discrete(drop= FALSE) +
  scale_y_continuous(trans= "log10", limits= c(1,280)) +
  labs(title= "Instantaneous bottleneck STAMP model", y = "Bottleneck Nb (Number of bacteria)") +
  theme_gray()


####################
# Visualisation of the CR model (theta, mu and r)
####################

# Theta CR
# Kruskal-wallis statistical test
test = Graph_kruskal(tab= tab, y= "Theta_CR", x= "Condition2")

# Display helper
values = tab %>% 
  group_by(Condition2, .drop= FALSE) %>% 
  summarise(mean = mean(Theta_CR), sd = sd(Theta_CR))

# Graph theta CR
ggplot(data= tab, aes(x= Condition2, y= Theta_CR)) +
  
  geom_boxplot(outlier.shape= NA, width=0.9, fill= "pink") +
  geom_point(aes(colour= Replicate),
                  position= position_jitterdodge(jitter.width= 0.3, dodge.width= 0.4), alpha= 1) +
  
  scale_colour_manual(values= c("#E41A1C", "#339900", "#FFD92F")) +
  geom_text(data = values, aes(label = test$groups, y = 1 , x = Condition2), vjust = -21.7, inherit.aes = FALSE) +
  scale_x_discrete(drop= FALSE) +
  scale_y_continuous(limits= c(0,52)) +
  labs(title= "Constant entry rate CR model", y = "Bottleneck effect mu/r (no units)") +
  theme_gray()


# Mu CR
# Kruskal-wallis statistical test
test = Graph_kruskal(tab= tab, y= "Mu_CR", x= "Condition2")

# Display helper
values = tab %>% 
  group_by(Condition2, .drop= FALSE) %>% 
  summarise(mean = mean(Mu_CR), sd = sd(Mu_CR))

# Graph mu CR
ggplot(data= tab, aes(x= Condition2, y= Mu_CR), ) +
  
  geom_boxplot(outlier.shape= NA, width=0.9, fill= "pink") +
  geom_point(aes(colour= Replicate),
                  position= position_jitterdodge(jitter.width= 0.3, dodge.width= 0.4), alpha= 1) +
  
  scale_colour_manual(values= c("#E41A1C", "#339900", "#FFD92F")) +
  geom_text(data = values, aes(label = test$groups, y = 1 , x = Condition2), vjust = -21.7, inherit.aes = FALSE) +
  scale_x_discrete(drop= FALSE) +
  scale_y_continuous(limits= c(0,250)) +
  labs(title= "Constant entry rate CR model", y = "Permeability mu (CFU/d)") +
  theme_gray()


# r CR
# Kruskal-wallis statistical test
test = Graph_kruskal(tab= tab, y= "r_CR", x= "Condition2")

# Display helper
values = tab %>% 
  group_by(Condition2, .drop= FALSE) %>% 
  summarise(mean = mean(r_CR), sd = sd(r_CR))

# Graph r CR
ggplot(data= tab, aes(x= Condition2, y= r_CR), ) +
  
  geom_boxplot(outlier.shape= NA, width=0.9, fill= "pink") +
  geom_point(aes(colour= Replicate),
                  position= position_jitterdodge(jitter.width= 0.3, dodge.width= 0.4), alpha= 1) +
  
  scale_colour_manual(values= c("#E41A1C", "#339900", "#FFD92F")) +
  geom_text(data = values, aes(label = test$groups, y = 1 , x = Condition2), vjust = -20.7, inherit.aes = FALSE) +
  scale_x_discrete(drop= FALSE) +
  scale_y_continuous(limits= c(0,6.5)) +
  labs(title= "Constant entry rate CR model", y = "Growth rate r (d-1)") +
  theme_gray()

####################
# Visualisation of the BS model (theta)
####################

# Theta BS
# Kruskal-wallis statistical test
test = Graph_kruskal(tab= tab, y= "Theta_BS", x= "Condition2")

# Display helper
values = tab %>% 
  group_by(Condition2, .drop= FALSE) %>% 
  summarise(mean = mean(Theta_BS), sd = sd(Theta_BS))

# Graph theta BS
ggplot(data= tab, aes(x= Condition2, y= Theta_BS), ) +
  
  geom_boxplot(outlier.shape= NA, width=0.9, fill= "lightgreen") +
  geom_point(aes(colour= Replicate),
                  position= position_jitterdodge(jitter.width= 0.3, dodge.width= 0.4), alpha= 1) +
  
  scale_colour_manual(values= c("#E41A1C", "#339900", "#FFD92F")) +
  geom_text(data = values, aes(label = test$groups, y = 1 , x = Condition2), vjust = -22, inherit.aes = FALSE) +
  scale_x_discrete(drop= FALSE) +
  scale_y_continuous(trans= "log10", limits= c(1,23), breaks= c(1,3,5,10,20)) +
  labs(title= "Bolthausen-Sznitsman BS model", y = "Normalised entry speed v (no units)") +
  theme_gray()


# Graph theta BS v2
ggplot(data= tab, aes(x= Condition2, y= Theta_BS, fill= Genotype), ) +
  
  geom_boxplot(outlier.shape= NA, width=0.8) +
  geom_point(aes(colour= Replicate),
                  position= position_jitterdodge(jitter.width= 0.3, dodge.width= 0.4), alpha= 1) +
  
  scale_fill_manual(values= c("#6699FF", "#FF9900")) +
  scale_colour_manual(values= c("#E41A1C", "#339900", "#FFD92F")) +
  geom_text(data = values, aes(label = test$groups, y = 1 , x = Condition2), vjust = -23.5, inherit.aes = FALSE) +
  scale_x_discrete(drop= FALSE) +
  scale_y_continuous(trans= "log10", limits= c(1,23), breaks= c(1,3,5,10,20)) +
  labs(y = bquote("Bottleneck effect"~theta[BS]*" (no units)")) +
  theme_gray()


####################
# Visualisation of the IR model (theta, mu and r)
####################

# Theta IR
# Kruskal-wallis statistical test
test = Graph_kruskal(tab= tab, y= "Theta_IR", x= "Condition2")

# Display helper
values = tab %>% 
  group_by(Condition2, .drop= FALSE) %>% 
  summarise(mean = mean(Theta_IR), sd = sd(Theta_IR))

# Graph theta IR
ggplot(data= tab, aes(x= Condition2, y= Theta_IR), ) +
  
  geom_boxplot(outlier.shape= NA, width=0.9, fill= "tomato") +
  geom_point(aes(colour= Replicate),
                  position= position_jitterdodge(jitter.width= 0.3, dodge.width= 0.4), alpha= 1) +
  
  scale_colour_manual(values= c("#E41A1C", "#339900", "#FFD92F")) +
  geom_text(data = values, aes(label = test$groups, y = 1 , x = Condition2), vjust = -0.9, inherit.aes = FALSE) +
  scale_x_discrete(drop= FALSE) +
  scale_y_continuous(limits= c(0,1.05)) +
  labs(title= "Increasing entry rate IR model", y = "Bacterial capacity index mu/r (no units)") +
  theme_gray()


# Mu IR
# Kruskal-wallis statistical test
test = Graph_kruskal(tab= tab, y= "Mu_IR", x= "Condition2")

# Display helper
values = tab %>% 
  group_by(Condition2, .drop= FALSE) %>% 
  summarise(mean = mean(Mu_IR), sd = sd(Mu_IR))

# Graph mu IR
ggplot(data= tab, aes(x= Condition2, y= Mu_IR), ) +
  
  geom_boxplot(outlier.shape= NA, width=0.9, fill= "tomato") +
  geom_point(aes(colour= Replicate),
                  position= position_jitterdodge(jitter.width= 0.3, dodge.width= 0.4), alpha= 1) +
  
  scale_colour_manual(values= c("#E41A1C", "#339900", "#FFD92F")) +
  geom_text(data = values, aes(label = test$groups, y = 1 , x = Condition2), vjust = -17.8, inherit.aes = FALSE) +
  scale_x_discrete(drop= FALSE) +
  scale_y_continuous(limits= c(0,5.3)) +
  labs(title= "Increasing entry rate IR model", y = "Permeability mu (CFU/d)") +
  theme_gray()

# Graph mu IR v2
ggplot(data= tab, aes(x= Condition2, y= Mu_IR, fill= Genotype), ) +
  
  geom_boxplot(outlier.shape= NA, width=0.8) +
  geom_point(aes(colour= Replicate),
                  position= position_jitterdodge(jitter.width= 0.3, dodge.width= 0.4), alpha= 1) +
  
  scale_fill_manual(values= c("#6699FF", "#FF9900")) +
  scale_colour_manual(values= c("#E41A1C", "#339900", "#FFD92F")) +
  geom_text(data = values, aes(label = test$groups, y = 1 , x = Condition2), vjust = -18.9, inherit.aes = FALSE) +
  scale_x_discrete(drop= FALSE) +
  scale_y_continuous(limits= c(0,5.3)) +
  labs(y = bquote("Roots permeability multiplier"~mu[IR]*" (no units)")) +
  theme_gray()


# r IR
# Kruskal-wallis statistical test
test = Graph_kruskal(tab= tab, y= "r_IR", x= "Condition2")

# Display helper
values = tab %>% 
  group_by(Condition2, .drop= FALSE) %>% 
  summarise(mean = mean(r_IR), sd = sd(r_IR))

# Graph r IR
ggplot(data= tab, aes(x= Condition2, y= r_IR), ) +
  
  geom_boxplot(outlier.shape= NA, width=0.9, fill= "tomato") +
  geom_point(aes(colour= Replicate),
                  position= position_jitterdodge(jitter.width= 0.3, dodge.width= 0.4), alpha= 1) +
  
  scale_colour_manual(values= c("#E41A1C", "#339900", "#FFD92F")) +
  geom_text(data = values, aes(label = test$groups, y = 1 , x = Condition2), vjust = -18.5, inherit.aes = FALSE) +
  scale_x_discrete(drop= FALSE) +
  scale_y_continuous(limits= c(0,7)) +
  labs(title= "Increasing entry rate IR model", y = "Growth rate r (d-1)") +
  theme_gray()

# Graph r IR v2
ggplot(data= tab, aes(x= Condition2, y= r_IR, fill= Genotype), ) +
  
  geom_boxplot(outlier.shape= NA, width=0.9) +
  geom_point(aes(colour= Replicate),
                  position= position_jitterdodge(jitter.width= 0.3, dodge.width= 0.4), alpha= 1) +
  
  scale_fill_manual(values= c("#6699FF", "#FF9900")) +
  scale_colour_manual(values= c("#E41A1C", "#339900", "#FFD92F")) +
  geom_text(data = values, aes(label = test$groups, y = 1 , x = Condition2), vjust = -20, inherit.aes = FALSE) +
  scale_x_discrete(drop= FALSE) +
  scale_y_continuous(limits= c(0,7)) +
  labs(y = bquote("In planta growth rate"~r[IR]*" ("~d^-1*")")) +
  theme_gray()  

```

Statistical tests to see the contribution of theta, mu and r to the infection

```{r Calculation}

tab = tab_tot

tab_temp = read.csv("Models_output_correct_inoc/data_and_estimators_model_STAMP.csv", header= TRUE, check.names= FALSE)
tab$Theta_STAMP = tab_temp$theta_STAMP
tab_temp = read.csv("Models_output_correct_inoc/data_and_estimators_model_CR.csv", header= TRUE, check.names= FALSE)
tab$Theta_CR = tab_temp$theta_CR
tab$Mu_CR = tab_temp$mu
tab$r_CR = tab_temp$r
tab_temp = read.csv("Models_output_correct_inoc/data_and_estimators_model_BS.csv", header= TRUE, check.names= FALSE)
tab$Theta_BS = tab_temp$`1 / theta_BS`
tab$Theta_BS = 1 / tab$Theta_BS
tab_temp = read.csv("Models_output_correct_inoc/data_and_estimators_model_IR.csv", header= TRUE, check.names= FALSE)
tab$Theta_IR = tab_temp$theta
tab$Mu_IR = tab_temp$mu
tab$r_IR = tab_temp$r

tab = tab %>%
  filter(Zone == "Root") %>%
  filter(`PCR2 validation on gel` > 0)

tab_temp = select(tab, c("Replicate","Genotype", "Temperature (°C)", "Root condition", "Collected fresh weight (g)", "Theta_STAMP", "Theta_CR", "Theta_BS", "Theta_IR", "r_CR", "r_IR", "Mu_CR", "Mu_IR", "Shannon diversity", "Bacterial load (CFU/g)", "Time post-inoculation (d)"))

# Encode each condition to logical
# M = 0, H = 1
tab_temp$Genotype = as.integer(tab_temp$Genotype != "M")
# 27°C = 0, 32°C = 1
tab_temp$`Temperature (°C)` = as.integer(tab_temp$`Temperature (°C)` != "27°C")
# Intact = 0, Cut = 1
tab_temp$`Root condition` = as.integer(tab_temp$`Root condition` != "Intact")


####################
# Testing homoscedasticity between conditions
####################

a = levene_test(tab, Theta_BS ~ Condition4)
print(a)

a = levene_test(tab, Mu_IR ~ Condition4)
print(a)

a = levene_test(tab, r_IR ~ Condition4)
print(a)


####################
# Multiple linear regression
# Contribution of the different experimental conditions to the models outputs
####################

# Multiple linear regression test with principal effects only then principal effect + secondary interactions

# Bottleneck NB in the STAMP model
# Principal effects
a = lm(Theta_STAMP ~ (`Root condition` + Genotype + `Temperature (°C)`), data = tab_temp)
print(summary(a))

# Principal effects + secondary interactions
a = lm(Theta_STAMP ~ (`Root condition` + Genotype + `Temperature (°C)`)^2, data = tab_temp)
print(summary(a))


# Bottleneck theta in the BS model
# Principal effects
a = lm(Theta_BS ~ (`Root condition` + Genotype + `Temperature (°C)`), data = tab_temp)
print(summary(a))
# Principal effects + secondary interactions
a = lm(Theta_BS ~ (`Root condition` + Genotype + `Temperature (°C)`)^2, data = tab_temp)
print(summary(a))


# Growth rate in the IR model
# Principal effects
a = lm(r_IR ~ (`Root condition` + Genotype + `Temperature (°C)`), data = tab_temp)
print(summary(a))
# Principal effects + secondary interactions
a = lm(r_IR ~ (`Root condition` + Genotype + `Temperature (°C)`)^2, data = tab_temp)
print(summary(a))


# Permeability in the IR model
# Principal effects
a = lm(Mu_IR ~ (`Root condition` + Genotype + `Temperature (°C)`), data = tab_temp)
print(summary(a))
# Principal effects + secondary interactions
a = lm(Mu_IR ~ (`Root condition` + Genotype + `Temperature (°C)`)^2, data = tab_temp)
print(summary(a))


####################
# Non-parametric PERMANOVA test
# Contribution of the different experimental conditions to the models outputs
####################

# Bottleneck Nb in the STAMP model
#Principal effects
test = adonis2(formula = tab_temp$Theta_STAMP ~ (`Root condition` + Genotype + `Temperature (°C)`),
                       data = tab_temp, permutations= 1000, by= "terms")

print(test)


# Principal effects + secondary interactions
test = adonis2(formula = tab_temp$Theta_STAMP ~ (`Root condition` + Genotype + `Temperature (°C)`)^2,
                       data = tab_temp, permutations= 1000, by= "terms")
print(test)


# Bottleneck theta in the BS model
# Principal effects
test = adonis2(formula = tab_temp$Theta_BS ~ (`Root condition` + Genotype + `Temperature (°C)`),
                       data = tab_temp, permutations= 1000, by= "terms")

# Add omega squared effect size
test$parOmegaSq = adonis_OmegaSq(test)$parOmegaSq
test = test %>%
  relocate("F", .after = last_col()) %>%
  relocate("Pr(>F)", .after = last_col())

print(test)


# Principal effects + secondary interactions
test = adonis2(formula = tab_temp$Theta_BS ~ (`Root condition` + Genotype + `Temperature (°C)`)^2,
                       data = tab_temp, permutations= 1000, by= "terms")
print(test)


# Growth rate r in the IR model
# Principal effects
test = adonis2(formula = tab_temp$r_IR ~ (`Root condition` + Genotype + `Temperature (°C)`),
                       data = tab_temp, permutations= 1000, by= "terms")

# Add omega squared effect size
test$parOmegaSq = adonis_OmegaSq(test)$parOmegaSq
test = test %>%
  relocate("F", .after = last_col()) %>%
  relocate("Pr(>F)", .after = last_col())

print(test)


# Principal effects + secondary interactions
test = adonis2(formula = tab_temp$r_IR ~ (`Root condition` + Genotype + `Temperature (°C)`)^2,
                       data = tab_temp, permutations= 1000, by= "terms")
print(test)


# Permeability mu in the IR model
# Principal effects
test = adonis2(formula = tab_temp$Mu_IR ~ (`Root condition` + Genotype + `Temperature (°C)`),
                       data = tab_temp, permutations= 1000, by= "terms")

# Add omega squared effect size
test$parOmegaSq = adonis_OmegaSq(test)$parOmegaSq
test = test %>%
  relocate("F", .after = last_col()) %>%
  relocate("Pr(>F)", .after = last_col())

print(test)

# Principal effects + secondary interactions
test = adonis2(formula = tab_temp$Mu_IR ~ (`Root condition` + Genotype + `Temperature (°C)`)^2,
                       data = tab_temp, permutations= 1000, by= "terms")
print(test)


# Contribution of plant weight to mu and r in the IR model
# r
test = adonis2(formula = tab_temp$r_IR ~ (`Collected fresh weight (g)` +`Root condition` + Genotype + `Temperature (°C)`),
                       data = tab_temp, permutations= 1000, by= "terms")

# Add omega squared effect size
test$parOmegaSq = adonis_OmegaSq(test)$parOmegaSq
test = test %>%
  relocate("F", .after = last_col()) %>%
  relocate("Pr(>F)", .after = last_col())

print(test)


test = adonis2(formula = tab_temp$r_IR ~ (`Collected fresh weight (g)` +`Root condition` + Genotype + `Temperature (°C)`)^2,
                       data = tab_temp, permutations= 1000, by= "terms")
print(test)

# Mu
test = adonis2(formula = tab_temp$Mu_IR ~ (`Collected fresh weight (g)` +`Root condition` + Genotype + `Temperature (°C)`),
                       data = tab_temp, permutations= 1000, by= "terms")

# Add omega squared effect size
test$parOmegaSq = adonis_OmegaSq(test)$parOmegaSq
test = test %>%
  relocate("F", .after = last_col()) %>%
  relocate("Pr(>F)", .after = last_col())

print(test)

test = adonis2(formula = tab_temp$Mu_IR ~ (`Collected fresh weight (g)` +`Root condition` + Genotype + `Temperature (°C)`)^2,
                       data = tab_temp, permutations= 1000, by= "terms")
print(test)

```

## Global figure with Hill numbers for models STAMP, CR, BS, IR

```{python, fig.width=12, fig.height=18}

# Function to load pickle files
def load_pickle(filename):
    with open(filename, "rb") as f:
        return pickle.load(f)

# Load the saved data
RF = load_pickle("Models_output_correct_inoc/RF.pkl")
BF1 = load_pickle("Models_output_correct_inoc/BF1.pkl")
BF3 = load_pickle("Models_output_correct_inoc/BF3.pkl")
BF4 = load_pickle("Models_output_correct_inoc/BF4.pkl")
BF6 = load_pickle("Models_output_correct_inoc/BF6.pkl")
BF5 = load_pickle("Models_output_correct_inoc/BF5.pkl")

# Extract experiment names from RF
experiment_names = list(RF.keys())

# Define Q values (x-axis for Hill numbers)
Q = np.arange(0, 4.1, 0.1)
#Q = np.load("Q1.npy")
# Function to compute Hill numbers
def hill_number(V, q):
    """Compute Hill numbers for a given frequency vector V and order q."""
    if q == 1:
        return np.exp(-np.sum(V[V > 0] * np.log(V[V > 0])))
    elif q == 0:
        return np.sum(V > 0)
    else:
        return (np.sum(V ** q)) ** (1 / (1 - q))

# Function to compute Hill curves for multiple frequency distributions
def compute_hill_curves(frequency_data):
    """Compute Hill curves for all frequency distributions in a configuration."""
    hill_curves = []
    for freq_vector in frequency_data:
        curve = [hill_number(freq_vector, q) for q in Q]
        hill_curves.append(curve)
    return np.array(hill_curves)

# Number of configurations and models
num_configurations = 7
num_models = 5  # Models 1, 3, 4, 6


# Create figure with subplots (7 rows, 5 columns)
fig, axes = plt.subplots(num_configurations, num_models + 1, figsize=(12, 18), sharex=True, sharey=True)

# Iterate over configurations and plot Hill curves
for i, exp_name in enumerate(experiment_names):
    # Compute Hill curves for observed and models
    observed_hill = compute_hill_curves(RF[exp_name])
    bf1_hill = compute_hill_curves(BF1[exp_name])
    bf3_hill = compute_hill_curves(BF3[exp_name])
    bf4_hill = compute_hill_curves(BF4[exp_name])
    bf6_hill = compute_hill_curves(BF6[exp_name])
    bf5_hill = compute_hill_curves(BF5[exp_name])

    # Plot observed frequencies (first column)
    for curve in observed_hill:
        axes[i, 0].plot(Q, curve, alpha=0.8, linewidth=1)
    axes[i, 0].set_title("Observed", fontsize=12)

    # Plot Model 1 (second column)
    for curve in bf1_hill:
        axes[i, 1].plot(Q, curve, alpha=0.8, linewidth=1)
    axes[i, 1].set_title("Model $\mathcal{M}_{STAMP}$", fontsize=12)

    # Plot Model 3 (third column)
    for curve in bf3_hill:
        axes[i, 2].plot(Q, curve, alpha=0.8, linewidth=1)
    axes[i, 2].set_title("Model $\mathcal{M}_{CR}$", fontsize=12)

    # Plot Model 4 (fourth column)
    for curve in bf4_hill:
        axes[i, 3].plot(Q, curve, alpha=0.8, linewidth=1)
    axes[i, 3].set_title("Model $\mathcal{M}_{BS}$", fontsize=12)


    # Plot Model 6 (5th column)
    for curve in bf6_hill:
        axes[i, 4].plot(Q, curve, alpha=0.8, linewidth=1)
    axes[i, 4].set_title("Model $\mathcal{M}_{IR}$", fontsize=12)
    
    # Plot Model 5 (6th column)
    for curve in bf5_hill:
        axes[i, 5].plot(Q, curve, alpha=0.8, linewidth=1)
    axes[i, 5].set_title("Model $\mathcal{M}_{K}$", fontsize=12)

# Set labels for rows with experiment names
for i, exp_name in enumerate(experiment_names):
    axes[i, 0].set_ylabel(f"{exp_name}", fontsize=12)

# Set shared x-label
for ax in axes[-1, :]:
    ax.set_xlabel("q", fontsize=12)

plt.tight_layout()
#plt.savefig("all_models_hill.png", dpi=300, bbox_inches='tight')

plt.show()
```

## Barcodes: experiments vs models

```{python, fig.width=10, fig.height=6}

# =======================
# Load pickle files
# =======================

def load_pickle(filename):
    if not os.path.exists(filename):
        raise FileNotFoundError(f"'{filename}' not found in repository.")
    with open(filename, "rb") as f:
        return pickle.load(f)

# Load data
RF = load_pickle("Models_output_correct_inoc/RF.pkl")
BF1 = load_pickle("Models_output_correct_inoc/BF1.pkl")
BF3 = load_pickle("Models_output_correct_inoc/BF3.pkl")
BF4 = load_pickle("Models_output_correct_inoc/BF4.pkl")
BF6 = load_pickle("Models_output_correct_inoc/BF6.pkl")
theta_values = load_pickle("Models_output_correct_inoc/theta_values1.pkl")  # Adjust the file name if needed

# =======================
# Helper function
# =======================

def plot_stacked_bars(ax, frequencies_list, title=None):
    n_replicates = len(frequencies_list)
    x_positions = np.arange(n_replicates)

    for i, freqs in enumerate(frequencies_list):
        freqs_sorted = np.sort(freqs)[::-1]
        bottom = 0
        for freq in freqs_sorted:
            ax.bar(x_positions[i], freq, bottom=bottom, width=0.5, edgecolor='black')
            bottom += freq

    # Clean axes
    ax.set_xticks([])
    ax.set_yticks([])
    for spine in ax.spines.values():
        spine.set_visible(False)

    ax.set_ylim(0, 1)

    if title:
        ax.set_title(title, fontsize=10)

# =======================
# Row names (no K model)
# =======================

row_names = [
    "Observed",
    "Model $\mathcal{M}_{STAMP}$",
    "Model $\mathcal{M}_{CR}$",
    "Model $\mathcal{M}_{BS}$",
    "Model $\mathcal{M}_{IR}$",
]

# =======================
# Select experiments
# =======================

# Extract experiment names
experiment_names = list(RF.keys())

# Select experiments to process
# selected_experiments = [experiment_names[0], experiment_names[2]]
selected_experiments = [experiment_names[0]]  # <-- change if needed

# =======================
# Main loop over experiments
# =======================

for exp_name in selected_experiments:
    observed_list = RF[exp_name]
    bf1_list = BF1[exp_name]
    bf3_list = BF3[exp_name]
    bf4_list = BF4[exp_name]
    bf6_list = BF6[exp_name]

    n_replicates = len(observed_list)

    # === Sorting according to descending theta_list ===
    theta_list = theta_values[exp_name]['theta_list']
    sorted_indices = np.argsort(theta_list)[::-1]  # Indices of descending theta

    # Reorder all replicate lists based on sorted indices
    observed_list = [observed_list[i] for i in sorted_indices]
    bf1_list = [bf1_list[i] for i in sorted_indices]
    bf3_list = [bf3_list[i] for i in sorted_indices]
    bf4_list = [bf4_list[i] for i in sorted_indices]
    bf6_list = [bf6_list[i] for i in sorted_indices]

    # Create figure: 5 rows, 1 column
    fig, axes = plt.subplots(
        nrows=5, ncols=1,
        figsize=(max(6, n_replicates * 0.7), 10),
        sharex=True, sharey=True
    )

    fig.suptitle(f"Configuration: {exp_name}", fontsize=18)

    all_data = [
        observed_list,
        bf1_list,
        bf3_list,
        bf4_list,
        bf6_list,
    ]

    for row_idx, data_list in enumerate(all_data):
        ax = axes[row_idx] if n_replicates > 1 else axes
        plot_stacked_bars(ax, data_list)

        ax.set_ylabel(row_names[row_idx], fontsize=12, rotation=0, labelpad=40, ha='right')

        # Top axis: add replicate numbers
        if row_idx == 0:
            ax.set_xticks(np.arange(n_replicates))
            ax.set_xticklabels([f"Rep {i}" for i in sorted_indices], fontsize=8)
            ax.tick_params(axis='x', which='both', length=0)

    plt.tight_layout(rect=[0, 0, 1, 0.95])

    # Save the figure
    #filename = f"{exp_name}_results_compact_sorted.png"
    #fig.savefig(filename, dpi=300, bbox_inches='tight')

    # Show the figure
    plt.show()


# Extraction of the simulated barcodes frequencies from the selected experiment into .csv files
np.savetxt("Models_output_correct_inoc/Barcodes_best_prediction_MI27/observed_list.csv",
        observed_list,
        delimiter =";",
        fmt ='% s')

np.savetxt("Models_output_correct_inoc/Barcodes_best_prediction_MI27/STAMP_list.csv",
        bf1_list,
        delimiter =";",
        fmt ='% s')

np.savetxt("Models_output_correct_inoc/Barcodes_best_prediction_MI27/CR_list.csv",
        bf3_list,
        delimiter =";",
        fmt ='% s')

np.savetxt("Models_output_correct_inoc/Barcodes_best_prediction_MI27/BS_list.csv",
        bf4_list,
        delimiter =";",
        fmt ='% s')

np.savetxt("Models_output_correct_inoc/Barcodes_best_prediction_MI27/IR_list.csv",
        bf6_list,
        delimiter =";",
        fmt ='% s')

```

## Visualisation of the shape of diversity according to each model

```{r Graph}

# Retrieving table of simulated barcodes according to each model (for condition M_Intact_27)

tab_predict = read_xlsx("Models_output_correct_inoc/Barcodes_best_prediction_MI27/Manual_all_in_one_table.xlsx")

tab_predict$`Data type` = factor(tab_predict$`Data type`,
                                 levels= c("Observed", "STAMP", "CR", "BS", "IR"),
                                 labels = c("Observed", "Best fit STAMP", "Best fit CR", "Best fit BS", "Best fit IR"))

tab_predict$`Plant ID` = factor(tab_predict$`Plant ID`)

tab = tab_predict

tab_temp = tab

####################
# Visualising of the real data versus the best fitting simulations attempting to reproduce them according to each model
####################

# Ordering by frequency value
for (i in c(1:nrow(tab_temp))) {
  tab_temp[i,5:124] = tab_temp[i,5:124][,rev(order(tab_temp[i,5:124]))]
}

tab_temp = pivot_longer(tab_temp, cols = c(5:124))
tab = pivot_longer(tab, cols = c(5:124))

# Removing zero values for graph
tab_temp = tab_temp %>%
  filter(value > 0)

# Converting tag ID to number
tab_temp$name = as.numeric(substring(tab_temp$name, 4))

# retrieving the default pyplot colour palette
colour_tab10 = c("#1f77b4", "#ff7f0e", "#2ca02c", "#d62728", "#9467bd", "#8c564b", "#e377c2", "#7f7f7f", "#bcbd22", "#17becf", "#1f77b4", "#ff7f0e", "#2ca02c", "#d62728", "#9467bd", "#8c564b", "#e377c2")


# Graph
ggplot(data= tab_temp, aes(x= name, y= value, colour= `Plant ID`)) +
  
  geom_point(shape= 1, size= 1) +
  
  facet_grid(~ `Data type`) +
  scale_colour_discrete(type= colour_tab10) +
  scale_x_continuous(limits= c(1,120), breaks= c(1,40,80,120)) +
  scale_y_continuous(trans= "log10", limits= c(10^-5,1)) +
  labs(x= "Tags sorted by decreasing frequency", y= "Frequency") +
  theme_pubr() +
  theme(panel.spacing= unit(0.7, "lines"), legend.position = "none",
        title = element_text(size= 11), axis.text = element_text(size= 9), legend.text = element_text(size= 9))
  
```

## Correlations between estimated parameters in the models

Helper functions

```{python}

with open('Analysis_output/tab_tot.csv', newline='') as file:
    data = list(csv.reader(file, delimiter=','))

def read_file(data, configuration, tags_range, configuration_column, validation_column, time_of_sampling_column, bacterial_load_column, sample_weight_column, sample_index_column, Ntypes):
    frequencies = []
    times = []
    loads = []
    weights = []
    indexes = []

    for row in data:
        if row[configuration_column] == configuration[0]: # Check condition
            if row[validation_column] == '1': # Check sequencing validation
                counts = [int(row[j]) for j in range(tags_range, Ntypes + tags_range)]  # Adjust for weight column
                counts_sum = sum(counts)
                if counts_sum > 0:
                    frequencies.append([c / counts_sum for c in counts])
                    times.append(float(row[time_of_sampling_column]))
                    loads.append(float(row[bacterial_load_column]))
                    weights.append(float(row[sample_weight_column]))
                    indexes.append(row[sample_index_column])

    return np.array(frequencies), np.array(times), np.array(loads), np.array(weights), np.array(indexes)

def load_freq_load(data, lconfigurations):
    all_real_frequencies = {}  # Stores frequencies per config
    all_times = {}             # Stores times per config
    all_loads = {}             # Stores loads per config
    all_weights = {}
    all_indexes = {}

    for configuration in lconfigurations:
        real_frequencies, times, loads, weights, indexes = read_file(data, configuration, tags_range, configuration_column, validation_column, time_of_sampling_column, bacterial_load_column, sample_weight_column, sample_index_column, Ntypes)
        if len(real_frequencies) == 0:
            print(f"No data for configuration: {configuration}")
            continue

        config_key = tuple(configuration)

        all_real_frequencies[config_key] = real_frequencies
        all_times[config_key] = times
        all_loads[config_key] = loads
        all_weights[config_key] = weights
        all_indexes[config_key] = indexes

    return all_real_frequencies, all_times, all_loads, all_weights, all_indexes


def load_pickle(filename):
    with open(filename, "rb") as f:
        return pickle.load(f)

```

IR model

```{python}

k = 0.78

# Build the r values with theta-dependent expression
RF, all_times, all_loads, all_weights, all_indexes = load_freq_load(data, lconfigurations)

theta_values = load_pickle("Models_output_correct_inoc/theta_values6.pkl")

results = {}

for configuration in RF:
    frequencies = RF[configuration]     # Shape: (n_samples, 120)
    times = all_times[configuration]  # Shape: (n_samples,)
    loads = all_loads[configuration]  # Shape: (n_samples,)
    weights = all_weights[configuration]  # Shape: (n_samples,)
    indexes = all_indexes[configuration]
    theta_list = np.array(theta_values[configuration]['theta_list'])  # Shape: (n_samples,)

    # Compute p1: max frequency per sample (row-wise max)
    p1 = np.max(frequencies, axis=1)  # Shape: (n_samples,)

    # Initialize r values with NaNs
    r_ = np.full_like(p1, np.nan, dtype=float)

    # Apply mask to avoid division by zero or invalid log
    safe_mask = (p1 > 0) & (loads > 0) & (times != 0) & (theta_list > 0)

    # Compute updated r using the new formula
    f1_load_product = p1[safe_mask] * loads[safe_mask] * weights[safe_mask]
    r_[safe_mask] = (
        np.log(f1_load_product) + 1/k * np.log(1 + np.log(2) * k / theta_list[safe_mask])
    ) / times[safe_mask]

    # Store results
    results[configuration] = {
        'p1': p1,
        'r_': r_
    }

```

Correlation between $\mu$ and r

```{python, fig.width=10, fig.height=6}

# Load data
theta_values = load_pickle("theta_values6.pkl")

# Create the scatter plot
plt.figure(figsize=(10, 6))

# Font configuration
plt.rcParams.update({
    'font.size': 14,
    'axes.labelsize': 16,
    'axes.titlesize': 18,
    'legend.fontsize': 12,
    'xtick.labelsize': 13,
    'ytick.labelsize': 13
})

# Store global lists for Spearman
all_mu = []
all_r = []

# Color cycle
color_cycle = plt.rcParams['axes.prop_cycle'].by_key()['color']

for i, configuration in enumerate(RF):
    theta_list = theta_values[configuration]['theta_list']
    r_values = results[configuration]['r_']

    # Convert and align
    theta_array = np.array(theta_list)
    r_array = np.array(r_values)
    min_len = min(len(theta_array), len(r_array))
    theta_array = theta_array[:min_len]
    r_array = r_array[:min_len]

    # Filter invalid r
    valid_mask = np.isfinite(r_array)
    theta_array = theta_array[valid_mask]
    r_array = r_array[valid_mask]
    mu_array = theta_array * r_array

    # Append to global Spearman lists
    all_mu.extend(mu_array)
    all_r.extend(r_array)

    # Clean config label (remove 'Root')
    label = " - ".join([x for x in configuration if x.lower() != 'root'])

    # Plot
    color = color_cycle[i % len(color_cycle)]
    plt.scatter(mu_array, r_array, label=label, alpha=0.7, color=color)
    plt.plot(np.mean(mu_array), np.mean(r_array), marker='x', markersize=10, markeredgewidth=2, color=color)

# Labels and style
plt.xlabel(r"$\mu = \theta \times r$")
plt.ylabel("$r$")
plt.legend(bbox_to_anchor=(1.05, 1), loc='upper left', title="Configurations")
plt.grid(True)
plt.tight_layout()

# Save and show
#plt.savefig("mu_vs_r_IR.png", dpi=300)
plt.show()

# --- Spearman correlation analysis ---
all_mu = np.array(all_mu)
all_r = np.array(all_r)

spearman_corr, spearman_pval = spearmanr(all_mu, all_r)

print(f"Spearman rank correlation coefficient (μ vs r): {spearman_corr:.4f}")
print(f"P-value: {spearman_pval:.4e}")

```

Correlation between $\mu$ and weight

```{python, fig.width=10, fig.height=6}

# Load data
theta_values = load_pickle("theta_values6.pkl")

# Create the scatter plot
plt.figure(figsize=(10, 6))

# Font configuration
plt.rcParams.update({
    'font.size': 14,
    'axes.labelsize': 16,
    'axes.titlesize': 18,
    'legend.fontsize': 12,
    'xtick.labelsize': 13,
    'ytick.labelsize': 13
})

# Store global lists for Spearman
all_mu = []
all_weights2 = []

# Color cycle
color_cycle = plt.rcParams['axes.prop_cycle'].by_key()['color']

for i, configuration in enumerate(RF):
    theta_list = theta_values[configuration]['theta_list']
    r_values = results[configuration]['r_']
    weight_values = all_weights[configuration]

    # Convert and align
    theta_array = np.array(theta_list)
    r_array = np.array(r_values)
    weight_array = np.array(weight_values)
    min_len = min(len(theta_array), len(r_array), len(weight_array))
    theta_array = theta_array[:min_len]
    r_array = r_array[:min_len]
    weight_array = weight_array[:min_len]

    # Filter valid data
    valid_mask = np.isfinite(r_array) & np.isfinite(weight_array)
    theta_array = theta_array[valid_mask]
    r_array = r_array[valid_mask]
    weight_array = weight_array[valid_mask]
    mu_array = theta_array * r_array

    # Append to global lists
    all_mu.extend(mu_array)
    all_weights2.extend(weight_array)

    # Clean config label (remove 'Root')
    label = " - ".join([x for x in configuration if x.lower() != 'root'])

    # Plot
    color = color_cycle[i % len(color_cycle)]
    plt.scatter(mu_array, weight_array, label=label, alpha=0.7, color=color)
    plt.plot(np.mean(mu_array), np.mean(weight_array), marker='x', markersize=10, markeredgewidth=2, color=color)

# Labels and style
plt.xlabel(r"$\mu = \theta \times r$")
plt.ylabel("Weight")
plt.legend(bbox_to_anchor=(1.05, 1), loc='upper left', title="Configurations")
plt.grid(True)
plt.tight_layout()

# Save and show
#plt.savefig("mu_vs_weight.png", dpi=300)
plt.show()

# --- Spearman correlation analysis ---
all_mu = np.array(all_mu)
all_weights2 = np.array(all_weights2)

spearman_corr, spearman_pval = spearmanr(all_mu, all_weights2)

print(f"Spearman rank correlation coefficient (μ vs weight): {spearman_corr:.4f}")
print(f"P-value: {spearman_pval:.4e}")

```

```{python}
# --- Spearman correlation analysis: theta vs weight ---
all_theta = np.array([theta for configuration in RF for theta in theta_values[configuration]['theta_list']])
all_weights2 = np.concatenate([all_weights[configuration][:len(theta_values[configuration]['theta_list'])] for configuration in RF])

# Filter out NaNs or mismatched lengths
valid_mask = np.isfinite(all_theta) & np.isfinite(all_weights2)
theta_valid = all_theta[valid_mask]
weight_valid = all_weights2[valid_mask]

# Compute Spearman
spearman_corr_theta, spearman_pval_theta = spearmanr(theta_valid, weight_valid)

print(f"Spearman rank correlation coefficient (θ vs weight): {spearman_corr_theta:.4f}")
print(f"P-value: {spearman_pval_theta:.4e}")

```

# Section of the reanalysis of the kinetics of infection in Marmande tomato plants (tab_kinetic)

Launch this at the beginning of the session to load data

N.B.: don't forget to set up your Python environment in Rstudio.

```{python}

import numpy as np
import matplotlib.pyplot as plt
import csv
import os
import time
from matplotlib.animation import FuncAnimation
from matplotlib import pylab
import math
import pickle
import multiprocessing as mp
from scipy.stats import gamma
import sympy as sp
from scipy.stats import t
from scipy.optimize import minimize_scalar
import matplotlib.cm as cm
from scipy.stats import spearmanr


# Info retrieved from document "Analysis_output/tab_kinetic.csv" in the functions "read_file"
Ntypes = 120 # Number of different tags
tags_range = 55 # Starting column for tags
sample_index_column = 1 # Column with sample names
configuration_column = 17 # Column that contains all tested samples conditions
time_of_sampling_column = 17 # Column with DPI of sampling
bacterial_load_column = 33
sample_weight_column = 19
validation_column = 221 # Column to check that the sample is validated
q_values = np.linspace(0, 8, 100)  # Range of q-values for Hill numbers


# Configurations
lconfigurations = [
    ['3'],
    ['4'],
    ['5'],
    ['6']
    ]

```

# I. NULL

# II. Model testing on the experiment of kinetics of infection in tomato plants (do not relaunch unless necessary; outputs are already saved)

## Initialisation of the helper functions

```{python, fig.width=8, fig.height=6}
# @title Main functions common to all models. Launch before the estimation

def sample_theta(K, theta_range, distribution_model):
    theta = np.random.uniform(*theta_range)
    V = distribution_model(K, theta)
    return V / np.sum(V)

def quantile_distance(theta_range, frequencies, distribution_model, parameter_name="theta", Nsim=100, quantile=0.05):
    K = len(frequencies)
    Q = np.arange(0, 4.1, 0.1)

    def hill_number(V, q):
        if q == 1:
            return np.exp(-np.sum(V[V > 0] * np.log(V[V > 0])))
        elif q == 0:
            return np.sum(V > 0)
        else:
            return (np.sum(V ** q)) ** (1 / (1 - q))

    def calculate_distance(V1, V2):
        DS = 0
        for q in Q:
            Hq1 = hill_number(V1, q)
            Hq2 = hill_number(V2, q)
            DS += abs(Hq1 - Hq2) / (Hq1 + Hq2)
        return DS / len(Q)

    distances = []
    for _ in range(Nsim):
        V = sample_theta(K, theta_range, distribution_model)
        D = calculate_distance(V, frequencies)
        distances.append(D)

    return np.quantile(distances, quantile)

def find_best_theta(frequencies, theta_range, distribution_model, distance_function, parameter_name="theta", step=1):
    distances = []
    theta_values = np.arange(theta_range[0], theta_range[1] + step, step)

    for theta in theta_values:
        dist = distance_function((theta, theta), frequencies, distribution_model, parameter_name)
        distances.append(dist)

    best_theta = theta_values[np.argmin(distances)]
    min_distance = min(distances)
    return best_theta, min_distance
  

def read_file(data, configuration, tags_range, configuration_column, validation_column, sample_index_column, time_of_sampling_column, bacterial_load_column, Ntypes):
    frequencies = []
    conditions = []
    for row in data:
        if row[configuration_column] == configuration[0]: # Check condition
            if row[validation_column] == '1': # Check sequencing validation
                counts = [int(row[j]) for j in range(tags_range, Ntypes + tags_range)]
                counts_sum = sum(counts)
                if counts_sum > 0:
                    frequencies.append([c / counts_sum for c in counts])
                    time = float(row[17])
                    conditions.append((row[sample_index_column], int(row[time_of_sampling_column]), float(row[bacterial_load_column])))

    return np.array(frequencies), np.array(conditions)

def process_configurations(data, lconfigurations, distribution_model, theta_range, step, distance_function=quantile_distance, parameter_name="theta", model_name="Adaptable Frequency Model"):
    theta_values = {}  # Dictionary to store theta lists, mean values, and performance
    results = []

    for configuration in lconfigurations:
        frequencies, _ = read_file(data, configuration, tags_range, configuration_column, validation_column, sample_index_column, time_of_sampling_column, bacterial_load_column, Ntypes)
        if len(frequencies) == 0:
            print(f"No data for configuration: {configuration}")
            continue

        mle_theta = []
        dist_s = []
        for freq in frequencies:
            best_theta, dist_ = find_best_theta(freq, theta_range, distribution_model, distance_function, parameter_name, step)
            mle_theta.append(best_theta)
            dist_s.append(dist_)

        # Compute mean theta and mean distance across all replicates
        mean_mle_theta = np.mean(mle_theta)
        mean_distance = np.mean(dist_s)  # Performance metric

        # Store both all best theta values, mean theta, and mean performance
        theta_values[tuple(configuration)] = {
            "theta_list": mle_theta,  # List of all best theta values
            "mean_theta": mean_mle_theta,  # Mean of best theta values
            "mean_distance": mean_distance  # Mean distance (performance metric)
        }

        # Store and print results (only mean values)
        results.append((configuration, mean_mle_theta, mean_distance))
        print(f"Modèle: {model_name}, Configuration: {configuration}, Mean theta: {mean_mle_theta:.2f}, Performance (Mean Distance): {mean_distance:.4f}")

    return theta_values  # Return both detailed and summarized theta values
  

with open('Analysis_output/tab_kinetic.csv', newline='') as file:
    data = list(csv.reader(file, delimiter=','))

# Functions to plot the Hill curves
def hill_number(V, q):
    """Compute Hill numbers for a given frequency vector V and order q."""
    if q == 1:
        return np.exp(-np.sum(V[V > 0] * np.log(V[V > 0])))
    elif q == 0:
        return np.sum(V > 0)
    else:
        return (np.sum(V ** q)) ** (1 / (1 - q))

def calculate_distance_per_q(V1, V2, Q):
    """Compute the diversity-based distance for each q."""
    distances = []
    for q in Q:
        Hq1 = hill_number(V1, q)
        Hq2 = hill_number(V2, q)
        distance = abs(Hq1 - Hq2) / (Hq1 + Hq2) if (Hq1 + Hq2) != 0 else 0
        distances.append(distance)
    return distances

def generate_best_frequency(K, theta, distribution_model, real_frequency, num_sim=100):
    """Generate 100 frequency vectors and return the closest one to the real frequency."""
    simulated_vectors = [sample_theta(K, (theta, theta), distribution_model) for _ in range(num_sim)]
    distances = [np.mean(calculate_distance_per_q(V, real_frequency, np.arange(0, 4.1, 0.1))) for V in simulated_vectors]
    best_index = np.argmin(distances)  # Index of lowest distance
    return simulated_vectors[best_index]  # Return the best-matching frequency

def plot_hill_curves(real_frequencies, best_simulated_frequencies):
    """Plot Hill curves for real and best simulated frequencies in separate subplots."""
    Q = np.arange(0, 4.1, 0.1)
    num_freqs = len(real_frequencies)
    color_map = plt.colormaps["tab10"]

    fig, axes = plt.subplots(1, 2, figsize=(12, 6), sharey=True)

    # Plot each real frequency in the left subplot
    for i, freq in enumerate(real_frequencies):
        hill_curve = [hill_number(freq, q) for q in Q]
        color = color_map(i / max(1, num_freqs - 1))
        axes[0].plot(Q, hill_curve, color=color, alpha=0.9, linewidth=2)

    # Plot each best simulated frequency in the right subplot
    for i, best_freq in enumerate(best_simulated_frequencies):
        hill_curve = [hill_number(best_freq, q) for q in Q]
        color = color_map(i / max(1, num_freqs - 1))
        axes[1].plot(Q, hill_curve, color=color, alpha=0.9, linewidth=2)

    # Formatting subplots
    axes[0].set_title("Observed Frequencies")
    axes[1].set_title("Best Simulated Frequencies")
    for ax in axes:
        ax.set_xlabel("q")
        ax.set_ylabel("Hill Number")
        ax.set_ylim(0, 120)

    plt.tight_layout()
    plt.show()

def process_with_plots(data, lconfigurations, distribution_model, theta_values):
    """Process configurations, generate the best frequency vectors, and plot Hill curves and global mean distance."""
    Q = np.arange(0, 4.1, 0.1)
    global_mean_distances = np.zeros(len(Q))  # Accumulate mean distances over all configurations
    total_samples = 0  # Count total frequencies processed
    all_configurations_mean_distances = [] # Mean distances for all individual configuration
    configuration_mean_distances = np.zeros(len(Q))  # Accumulate mean distances within each configuration
    configuration_samples = 0 # Count frequencies processed within each configuration

    all_real_frequencies = {}  # Store real frequencies for all configurations
    all_best_simulated_frequencies = {}  # Store best simulated frequencies for all configurations

    for configuration in lconfigurations:
        real_frequencies, _ = read_file(data, configuration, tags_range, configuration_column, validation_column, sample_index_column, time_of_sampling_column, bacterial_load_column, Ntypes)
        if len(real_frequencies) == 0:
            print(f"No data for configuration: {configuration}")
            continue

        K = len(real_frequencies[0])  # Number of species/types
        config_key = tuple(configuration)
        theta_list = theta_values.get(config_key, {}).get("theta_list", None)

        if theta_list is None or len(theta_list) != len(real_frequencies):
            print(f"Missing or mismatched theta values for configuration: {configuration}")
            continue

        best_simulated_frequencies = []
        for real_freq, theta in zip(real_frequencies, theta_list):
            best_freq = generate_best_frequency(K, theta, distribution_model, real_freq)
            best_simulated_frequencies.append(best_freq)

            # Compute distance for this frequency
            distances = calculate_distance_per_q(real_freq, best_freq, Q)
            global_mean_distances += np.array(distances)
            total_samples += 1
            
            # Distance retriever for the configuration
            configuration_mean_distances += np.array(distances)
            configuration_samples += 1

        # Compute and append distance for the configuration
        all_configurations_mean_distances.append(configuration_mean_distances / configuration_samples)

        # Reset configuration distance and sample count
        configuration_mean_distances = np.zeros(len(Q))
        configuration_samples = 0
        
        # Store results for this configuration
        all_real_frequencies[config_key] = real_frequencies
        all_best_simulated_frequencies[config_key] = best_simulated_frequencies

        # Plot Hill curves for this configuration
        plot_hill_curves(real_frequencies, best_simulated_frequencies)

    # Compute final averaged distances
    if total_samples > 0:
        global_mean_distances /= total_samples

    # Plot global error figure
    plt.figure(figsize=(8, 6))
    plt.plot(Q, global_mean_distances, color="black", linewidth=2)
    plt.xlabel("q")
    plt.ylabel("Mean Distance")
    plt.title("Mean Distance Averaged Over All Configurations")
    plt.show()

    return Q, global_mean_distances, all_real_frequencies, all_best_simulated_frequencies, all_configurations_mean_distances

# functions to Save and load the results
def save_pickle(theta_values, filename="theta_values1.pkl"):
    """
    Save theta_values dictionary as a Pickle file.
    """
    with open(filename, "wb") as f:
        pickle.dump(theta_values, f)
    print(f"Saved to {filename}")

def load_pickle(filename="theta_values1.pkl"):
    """
    Load theta_values dictionary from a Pickle file.
    """
    with open(filename, "rb") as f:
        return pickle.load(f)

```

## Multinomial model

```{python}
# define the model
def distribution_modelM1(K, theta):
    Nb = theta
    return np.random.multinomial(Nb, [1 / K] * K)
```

```{python}

# define the range for parameter values
theta_range = (1, 200)
step = 1


print('Model performance')
# Run the processing function
theta_values1 = process_configurations(data, lconfigurations, distribution_modelM1, theta_range, step, quantile_distance, parameter_name="Nb", model_name = "STAMP Model")
save_pickle(theta_values1,"Models_output_kinetic/theta_values1.pkl")
```

```{python}
theta_values1 = load_pickle("Models_output_kinetic/theta_values1.pkl")  # Loads it back
Q1, D1, RF, BF1, all_D1 = process_with_plots(data, lconfigurations, distribution_modelM1, theta_values1)
np.save("Models_output_kinetic/Q1.npy", Q1)
np.save("Models_output_kinetic/D1.npy", D1)
np.save("Models_output_kinetic/all_D1.npy", all_D1)
save_pickle(RF, filename="Models_output_kinetic/RF.pkl")
save_pickle(BF1, filename="Models_output_kinetic/BF1.pkl")

```

## Constant rate model CR

```{python}
# define the model
def distribution_modelM3(K, theta):
    mu = theta
    Ntypes = K
    r = 1
    T = 10
    # on simule les temps d'entrée :
    tau = [ np.random.exponential( 1/mu ) ]
    while tau[-1] < T:
        tau += [ tau[-1] + np.random.exponential( 1/mu ) ]
    qt_prov = [ np.exp( r * (T-tau[i]) ) for i in range(len(tau)) ]
    types = [ np.random.randint(0, Ntypes) for i in range(len(tau))]
    qt_finale = []
    for i in range(len(tau)):
        if types[i] != -1:
            qt_cur = qt_prov[i]
            for j in range(i+1, len(tau)):
                if types[j] == types[i] and types[j] != -1:
                    qt_cur += qt_prov[j]
                    types[j] = -1
            types[i] = -1
            qt_finale += [ qt_cur ]
    if len(qt_finale) > Ntypes: # impossible, normalement
        print(len(qt_finale), "types différents sont passés")
        qt_finale = qt_finale[:Ntypes]
    else:
        qt_finale += (Ntypes - len(qt_finale)) * [0]
    res = np.array(qt_finale) #/ np.sum(np.array(qt_finale))

    return np.sort(res)[::-1]
```

```{python}
# define the range for parameter values
theta_range = (1, 50)
step = 1

print('Model performance')
# Run the processing function
theta_values3 = process_configurations(data, lconfigurations, distribution_modelM3, theta_range, step, quantile_distance, parameter_name="Nb", model_name = "CR1 Model")
save_pickle(theta_values3,"Models_output_kinetic/theta_values3.pkl")
```

```{python}
theta_values3 = load_pickle("Models_output_kinetic/theta_values3.pkl")
Q3, D3, RF, BF3, all_D3 = process_with_plots(data, lconfigurations, distribution_modelM3, theta_values3)
np.save("Models_output_kinetic/Q3.npy", Q3)
np.save("Models_output_kinetic/D3.npy", D3)
np.save("Models_output_kinetic/all_D3.npy", all_D3)
save_pickle(BF3, filename="Models_output_kinetic/BF3.pkl")
```

## Bolthausen-Sznitman coalescent model BS

```{python}
#**Step 1**: construct the initial random recursive tree.
def update_edges( l, k ):
    current = k
    while current != 0:
        current = l[current][0]
        l[current][2] += [k]


def init_bs( n ):
    l_edges = [ [0,[0],[]] ]
    new_edge = [0,[0],[]]
    for k in range(1,n):
        parent_new = np.random.randint(k)
        new_edge = [ parent_new, [k], []]
        l_edges += [ new_edge ]
        update_edges( l_edges, k )
    return l_edges
#**Step 2**: each edge is "lifted" at rate $1$ (in the terminology of *Berestycki 2009*).
def step_bs( tree, k ): # lift edge k of tree
    loss = 0
    parent = tree[k][0]
    tree[parent][1] += [k] + tree[k][2] # the edge is lifted to the parent
    for i in [k] + tree[k][2]:
        if tree[i][1][0] != -1:
            tree[i][1] = [-1] # unactivated
            loss += 1
    return tree, loss # tree : resulting tree; loss: number of blocks lost
#**Step 3:** the following function runs a Bolthausen-Sznitman coalescent for a time [t], starting from an initial tree [tree] with specified number of blocks [nblocks]).

def run_bs( nblocks, t, tree ):

    t_cur = np.random.exponential(1/nblocks)

    while t_cur < t and nblocks > 1:


        i = np.random.randint( 1, nblocks ) # choose the edge to be lifted
        edge = 0
        while i > -1:
            if tree[ edge ][1][0] != -1:
                edge += 1
                i -= 1
            else:
                edge += 1
        tree, loss = step_bs( tree, edge - 1 )
        nblocks -= loss

        t_cur += np.random.exponential(1/nblocks)

    return tree, nblocks


#run_bs(10000,0.5)
def display( tree ):
    for k in range( len(tree) ):
        if tree[k][1][0] != -1:
            print(tree[k][1], end='  ')
    print("")
def get_prop_bs( N, t ):
    tree,_ = run_bs( N, t, init_bs( N ) )
    prop = [] # get proportions
    for e in tree:
        if e[1][0] != -1:
            prop += [ len(e[1]) ]
    prop = prop / np.sum( prop )
    return prop
def sample_tags(prop, Ntags):
    prop_tags = np.zeros( Ntags )
    for p in prop:
        tag = np.random.randint( Ntags )
        prop_tags[ tag ] += p
    return prop_tags
```

```{python}
# define the model
def distribution_modelM4(K, theta):
    t = theta
    return sample_tags( get_prop_bs(2000, t), K )
```

```{python}
# define the range for parameter values
theta_range = (0.05, 1)
step = 0.05

print('Model performance')
# Run the processing function
theta_values4 = process_configurations(data, lconfigurations, distribution_modelM4, theta_range, step, quantile_distance, parameter_name="t", model_name = "BS Model")
save_pickle(theta_values4,"Models_output_kinetic/theta_values4.pkl")
```

```{python}
theta_values4 = load_pickle("Models_output_kinetic/theta_values4.pkl")
Q4, D4, RF, BF4, all_D4 = process_with_plots(data, lconfigurations, distribution_modelM4, theta_values4)
np.save("Models_output_kinetic/Q4.npy", Q4)
np.save("Models_output_kinetic/D4.npy", D4)
np.save("Models_output_kinetic/all_D4.npy", all_D4)
save_pickle(BF4, filename="Models_output_kinetic/BF4.pkl")
```

## Kingman coalescent

```{python}
def run_km( t, prop = [] ):

    nblocks = len(prop)
    t_cur = 0

    if nblocks > 1:
        t_cur = np.random.exponential(2/(nblocks * (nblocks-1)))

    while t_cur < t and nblocks > 1:
        t_cur += np.random.exponential(2/(nblocks * (nblocks-1)))

        i,j=[0,0]
        while i == j:
            i,j = np.random.randint( 0, nblocks, 2)
        prop[i] += prop[j] # j gets to i
        prop.pop(j) # remove j
        nblocks -= 1
    return prop

def sample_tags(prop, Ntags):
    prop_tags = np.zeros( Ntags )
    for p in prop:
        tag = np.random.randint( Ntags )
        prop_tags[ tag ] += p
    return prop_tags
```

```{python}
# define the model
def distribution_modelM5(K, theta):
    t = theta
    return sample_tags( run_km(t, [1/1000] * 1000), K )
```

```{python}
# define the range for parameter values
theta_range = (0.05, 1)
step = 0.05
print('Model performance')
# Run the processing function
theta_values5 = process_configurations(data, lconfigurations, distribution_modelM5, theta_range, step, quantile_distance, parameter_name="t", model_name = "K Model")
save_pickle(theta_values5,"Models_output_kinetic/theta_values5.pkl")
```

```{python}
theta_values5 = load_pickle("Models_output_kinetic/theta_values5.pkl")
Q5, D5, RF, BF5, all_D5 = process_with_plots(data, lconfigurations, distribution_modelM5, theta_values5)
np.save("Models_output_kinetic/Q5.npy", Q5)
np.save("Models_output_kinetic/D5.npy", D5)
np.save("Models_output_kinetic/all_D5.npy", all_D5)
save_pickle(BF5, filename="Models_output_kinetic/BF5.pkl")
```

## Increasing entry rate model IR

```{python}

# NHPP inverse cumulative intensity function for λ(t) = mu * exp(k * t)
def nhpp_exponential_inverse(s, mu, k):
    return (1.0 / k) * np.log(k * s / mu + 1)

# define the model
def distribution_modelM6(K, theta):
    mu = theta
    k = 0.78
    Ntypes = K
    T = 10

    # Step 1: Simulate NHPP arrival times
    tau = []
    s = 0.0
    while True:
        s += np.random.exponential(1.0)  # Homogeneous Poisson interarrival
        t_candidate = nhpp_exponential_inverse(s, mu, k)
        if t_candidate > T:
            break
        tau.append(t_candidate)

    # Step 2: Compute provisional quantity
    qt_prov = [np.exp((T - t)) for t in tau]

    # Step 3: Assign and merge types
    types = [np.random.randint(0, Ntypes) for _ in tau]
    qt_finale = []
    for i in range(len(tau)):
        if types[i] != -1:
            qt_cur = qt_prov[i]
            for j in range(i + 1, len(tau)):
                if types[j] == types[i] and types[j] != -1:
                    qt_cur += qt_prov[j]
                    types[j] = -1
            types[i] = -1
            qt_finale.append(qt_cur)

    # Step 4: Padding or trimming
    if len(qt_finale) > Ntypes:
        print(len(qt_finale), "types différents sont passés")
        qt_finale = qt_finale[:Ntypes]
    else:
        qt_finale += (Ntypes - len(qt_finale)) * [0]

    # Return sorted descending quantities
    return np.sort(np.array(qt_finale))[::-1]

```

```{python}
# define the range for parameter values
theta_range = (0.05, 1)
step = 0.05

print('Model performance')
# Run the processing function
theta_values6 = process_configurations(data, lconfigurations, distribution_modelM6, theta_range, step, quantile_distance, parameter_name="mu", model_name = "VR Model")
save_pickle(theta_values6,"Models_output_kinetic/theta_values6.pkl")
```

```{python}
theta_values6 = load_pickle("Models_output_kinetic/theta_values6.pkl")
Q6, D6, RF, BF6, all_D6 = process_with_plots(data, lconfigurations, distribution_modelM6, theta_values6)
np.save("Models_output_kinetic/Q6.npy", Q6)
np.save("Models_output_kinetic/D6.npy", D6)
np.save("Models_output_kinetic/all_D6.npy", all_D6)
save_pickle(BF6, filename="Models_output_kinetic/BF6.pkl")
```

# III. Some graphs

## Helper function

```{python, fig.width=10, fig.height=6}

# =======================
# Load pickle files
# =======================

def load_pickle(filename):
    if not os.path.exists(filename):
        raise FileNotFoundError(f"'{filename}' not found in repository.")
    with open(filename, "rb") as f:
        return pickle.load(f)

# Load data
RF = load_pickle("Models_output_kinetic/RF.pkl")
BF1 = load_pickle("Models_output_kinetic/BF1.pkl")
BF3 = load_pickle("Models_output_kinetic/BF3.pkl")
BF4 = load_pickle("Models_output_kinetic/BF4.pkl")
BF6 = load_pickle("Models_output_kinetic/BF6.pkl")
theta_values = load_pickle("Models_output_kinetic/theta_values1.pkl")  # Adjust the file name if needed

```

## Error curves

```{python, fig.width=8, fig.height=6}

# Initialize plot
plt.figure(figsize=(8, 6))

i = 1
Q = np.load(f"Models_output_kinetic/Q{i}.npy")  # Load Qi
D = np.load(f"Models_output_kinetic/D{i}.npy")  # Load Di
plt.plot(Q, D, linewidth=2, label="Model $\mathcal{{M}}_{STAMP}$")  # Plot with label
i = 3
Q = np.load(f"Models_output_kinetic/Q{i}.npy")  # Load Qi
D = np.load(f"Models_output_kinetic/D{i}.npy")  # Load Di
plt.plot(Q, D, linewidth=2, label="Model $\mathcal{{M}}_{CR}$")  # Plot with label
i = 4
Q = np.load(f"Models_output_kinetic/Q{i}.npy")  # Load Qi
D = np.load(f"Models_output_kinetic/D{i}.npy")  # Load Di
plt.plot(Q, D, linewidth=2, label="Model $\mathcal{{M}}_{BS}$")  # Plot with label
i = 6
Q = np.load(f"Models_output_kinetic/Q{i}.npy")  # Load Qi
D = np.load(f"Models_output_kinetic/D{i}.npy")  # Load Di
plt.plot(Q, D, linewidth=2, label="Model $\mathcal{{M}}_{IR}$")  # Plot with label


# Formatting the plot
plt.xlabel("$q$", fontsize=16)
plt.ylabel("Mean relative error", fontsize=16)
plt.legend(fontsize=14)
plt.xticks(fontsize=16)
plt.yticks(fontsize=16)
plt.grid()
plt.tight_layout()
#Save the figure as a PNG file
#plt.savefig("mean_error.png", dpi=300, bbox_inches='tight')

plt.show()
```

```{python, fig.width=6, fig.height=32}

# Plot mean error for each condition
# Load the saved data
RF = load_pickle("Models_output_kinetic/RF.pkl")

# Extract experiment names from RF
experiment_names = list(RF.keys())

Q = np.arange(0, 4.1, 0.1)

# Create figure with subplots (7 rows, 1 columns)
fig, axes = plt.subplots(9, 1, figsize=(6, 32), sharex=True, sharey=True)

# Iterate over configurations and plot Hill curves
for j, exp_name in enumerate(experiment_names):

  i = 1
  D = np.load(f"Models_output_kinetic/all_D{i}.npy")
  D = D[j]
  axes[j].plot(Q, D, alpha=0.8, linewidth=2, label="Model $\mathcal{{M}}_{STAMP}$")

  i = 3
  D = np.load(f"Models_output_kinetic/all_D{i}.npy")
  D = D[j]
  axes[j].plot(Q, D, alpha=0.8, linewidth=2, label="Model $\mathcal{{M}}_{CR}$")

  i = 4
  D = np.load(f"Models_output_kinetic/all_D{i}.npy")
  D = D[j]
  axes[j].plot(Q, D, alpha=0.8, linewidth=2, label="Model $\mathcal{{M}}_{BS}$")

  i = 6
  D = np.load(f"Models_output_kinetic/all_D{i}.npy")
  D = D[j]
  axes[j].plot(Q, D, alpha=0.8, linewidth=2, label="Model $\mathcal{{M}}_{IR}$")

  #SV Set experiment label
  axes[j].set_ylabel(f"{exp_name}", fontsize=12)
  axes[j].grid()
  axes[j].legend(fontsize=14)

axes[j].set_xlabel("q", fontsize=12)

plt.tight_layout()
plt.show()

```

## Extraction of the csv tables with theta, mu and r

```{python}
# STAMP Model This code builds a csv file with the data. The last column corresponds to theta

with open('Analysis_output/tab_kinetic.csv', newline='') as file:
    data = list(csv.reader(file, delimiter=','))

def read_file(data, configuration, tags_range, configuration_column, validation_column, time_of_sampling_column, bacterial_load_column, sample_weight_column, sample_index_column, Ntypes):
    frequencies = []
    times = []
    loads = []
    weights = []
    indexes = []

    for row in data:
        if row[configuration_column] == configuration[0]: # Check condition
            if row[validation_column] == '1': # Check sequencing validation
                counts = [int(row[j]) for j in range(tags_range, Ntypes + tags_range)]  # Adjust for weight column
                counts_sum = sum(counts)
                if counts_sum > 0:
                    frequencies.append([c / counts_sum for c in counts])
                    times.append(float(row[time_of_sampling_column]))
                    loads.append(float(row[bacterial_load_column]))
                    weights.append(float(row[sample_weight_column]))
                    indexes.append(row[sample_index_column])

    return np.array(frequencies), np.array(times), np.array(loads), np.array(weights), np.array(indexes)


def load_pickle(filename):
    with open(filename, "rb") as f:
        return pickle.load(f)

# Load theta values
theta_values = pickle.load(open('Models_output_kinetic/theta_values1.pkl', 'rb'))

# Load precomputed rates (r) per config
# Here we recompute the rates so we can align them


# Read CSV into list
with open('Analysis_output/tab_kinetic.csv', newline='') as file:
    data = list(csv.reader(file))

# Compute r per valid row to match theta
# Compute rates for all configs
rates = {}
all_indexes = {}
for configuration in lconfigurations:
    config_key = tuple(configuration)
    frequencies, times, loads, weights, indexes = read_file(data, configuration, tags_range, configuration_column, validation_column, time_of_sampling_column, bacterial_load_column, sample_weight_column, sample_index_column, Ntypes)
    
    if config_key in theta_values:
        theta_list = np.array(theta_values[config_key]['theta_list'])
        p1 = np.max(frequencies, axis=1)
        r_ = np.full_like(p1, np.nan)
        mask = (p1>0) & (loads>0) & (times!=0) & (theta_list>0)
        r_[mask] = (np.log(p1[mask]*loads[mask]*weights[mask])+(1.0/theta_list[mask]))/times[mask]
        rates[config_key] = r_
        all_indexes[config_key] = indexes


# Prepare output file with header
#input_file = "Models_output/Data_py.csv"
input_file = "Analysis_output/tab_kinetic.csv"
output_file = "Models_output_kinetic/data_and_estimators_model_STAMP.csv"
with open(input_file, newline='') as infile:
    reader = list(csv.reader(infile))

# Header with only theta
header = ['index'] + ['theta_STAMP']

# Track per-config index to select correct theta
idx_per_config = {tuple(c): 0 for c in lconfigurations}

with open(output_file, mode='w', newline='') as outfile:
    writer = csv.writer(outfile)
    writer.writerow(header)
    for row in reader[1:]:
        configuration = (row[configuration_column],)
        index_val = ""
        theta_val = ""  # default blank
        if configuration in theta_values and row[validation_column] == '1':
            i = idx_per_config[configuration]
            theta_list = np.array(theta_values[configuration]['theta_list'])
            index_list = all_indexes[configuration]
            if i < len(theta_list):
                index_val = index_list[i]
                theta_val = theta_list[i]
                idx_per_config[configuration] += 1
        writer.writerow([index_val] + [theta_val])

print(f"OK Saved file to {output_file}")

```

```{python}
# CR Model This code builds a csv file with the data. The last three columns correspond to theta, r, mu

with open('Analysis_output/tab_kinetic.csv', newline='') as file:
    data = list(csv.reader(file, delimiter=','))

def read_file(data, configuration, tags_range, configuration_column, validation_column, time_of_sampling_column, bacterial_load_column, sample_weight_column, sample_index_column, Ntypes):
    frequencies = []
    times = []
    loads = []
    weights = []
    indexes = []

    for row in data:
        if row[configuration_column] == configuration[0]: # Check condition
            if row[validation_column] == '1': # Check sequencing validation
                counts = [int(row[j]) for j in range(tags_range, Ntypes + tags_range)]  # Adjust for weight column
                counts_sum = sum(counts)
                if counts_sum > 0:
                    frequencies.append([c / counts_sum for c in counts])
                    times.append(float(row[time_of_sampling_column]))
                    loads.append(float(row[bacterial_load_column]))
                    weights.append(float(row[sample_weight_column]))
                    indexes.append(row[sample_index_column])

    return np.array(frequencies), np.array(times), np.array(loads), np.array(weights), np.array(indexes)


def load_pickle(filename):
    with open(filename, "rb") as f:
        return pickle.load(f)

# Load theta values
theta_values = pickle.load(open('Models_output_kinetic/theta_values3.pkl', 'rb'))

# Load precomputed rates (r) per config
# Here we recompute the rates so we can align them


# Read CSV into list
with open('Analysis_output/tab_kinetic.csv', newline='') as file:
    data = list(csv.reader(file))

# Compute r per valid row to match theta
# Compute rates for all configs
rates = {}
all_indexes = {}
for configuration in lconfigurations:
    config_key = tuple(configuration)
    frequencies, times, loads, weights, indexes = read_file(data, configuration, tags_range, configuration_column, validation_column, time_of_sampling_column, bacterial_load_column, sample_weight_column, sample_index_column, Ntypes)
    
    if config_key in theta_values:
        theta_list = np.array(theta_values[config_key]['theta_list'])
        p1 = np.max(frequencies, axis=1)
        r_ = np.full_like(p1, np.nan)
        mask = (p1>0) & (loads>0) & (times!=0) & (theta_list>0)
        r_[mask] = (np.log(p1[mask]*loads[mask]*weights[mask])+(1.0/theta_list[mask]))/times[mask]
        rates[config_key] = r_
        all_indexes[config_key] = indexes


# Prepare output file with header
#input_file = "Models_output/Data_py.csv"
input_file = "Analysis_output/tab_kinetic.csv"
output_file = "Models_output_kinetic/data_and_estimators_model_CR.csv"
with open(input_file, newline='') as infile:
    reader = list(csv.reader(infile))

# Header with only theta
header = ['index', 'theta_CR', 'r', 'mu']

# Track per-config index to select correct theta
idx_per_config = {tuple(c): 0 for c in lconfigurations}

with open(output_file, mode='w', newline='') as outfile:
    writer = csv.writer(outfile)
    writer.writerow(header)
    for row in reader[1:]:
        configuration = (row[configuration_column],)
        index_val = ""
        theta_val = ""
        r_val = ""
        mu_val = ""
        if configuration in rates and row[validation_column] == '1':
            i = idx_per_config[configuration]
            theta_list = np.array(theta_values[configuration]['theta_list'])
            r_list = rates[configuration]
            index_list = all_indexes[configuration]
            if i < len(theta_list):
                index_val = index_list[i]
                theta_val = theta_list[i]
                r_val = r_list[i]
                mu_val = theta_val * r_val
                idx_per_config[configuration] += 1
        writer.writerow([index_val, theta_val, r_val, mu_val])

print(f"OK Saved file to {output_file}")

```

```{python}
# IR Model This code builds a csv file with the data. The last three columns correspond to theta, r, mu

k = 0.78

with open('Analysis_output/tab_kinetic.csv', newline='') as file:
    data = list(csv.reader(file, delimiter=','))

def read_file(data, configuration, tags_range, configuration_column, validation_column, time_of_sampling_column, bacterial_load_column, sample_weight_column, sample_index_column, Ntypes):
    frequencies = []
    times = []
    loads = []
    weights = []
    indexes = []

    for row in data:
        if row[configuration_column] == configuration[0]: # Check condition
            if row[validation_column] == '1': # Check sequencing validation
                counts = [int(row[j]) for j in range(tags_range, Ntypes + tags_range)]  # Adjust for weight column
                counts_sum = sum(counts)
                if counts_sum > 0:
                    frequencies.append([c / counts_sum for c in counts])
                    times.append(float(row[time_of_sampling_column]))
                    loads.append(float(row[bacterial_load_column]))
                    weights.append(float(row[sample_weight_column]))
                    indexes.append(row[sample_index_column])

    return np.array(frequencies), np.array(times), np.array(loads), np.array(weights), np.array(indexes)


def load_pickle(filename):
    with open(filename, "rb") as f:
        return pickle.load(f)

# Load theta values
theta_values = pickle.load(open('Models_output_kinetic/theta_values6.pkl', 'rb'))

# Load precomputed rates (r) per config
# Here we recompute the rates so we can align them


# Read CSV into list
with open('Analysis_output/tab_kinetic.csv', newline='') as file:
    data = list(csv.reader(file))

# Compute r per valid row to match theta
# Compute rates for all configs
rates = {}
all_indexes = {}
for configuration in lconfigurations:
    config_key = tuple(configuration)
    frequencies, times, loads, weights, indexes = read_file(data, configuration, tags_range, configuration_column, validation_column, time_of_sampling_column, bacterial_load_column, sample_weight_column, sample_index_column, Ntypes)
    
    if config_key in theta_values:
        theta_list = np.array(theta_values[config_key]['theta_list'])
        p1 = np.max(frequencies, axis=1)
        r_ = np.full_like(p1, np.nan)
        mask = (p1>0) & (loads>0) & (times!=0) & (theta_list>0)
        r_[mask] = (np.log(p1[mask]*loads[mask]*weights[mask])+(1/k * np.log(1 + np.log(2) * k / theta_list[mask])))/times[mask]
        rates[config_key] = r_
        all_indexes[config_key] = indexes


# Prepare output file with header
#input_file = "Models_output/Data_py.csv"
input_file = "Analysis_output/tab_kinetic.csv"
output_file = "Models_output_kinetic/data_and_estimators_model_IR.csv"
with open(input_file, newline='') as infile:
    reader = list(csv.reader(infile))

# Header with only theta
header = ['index', 'theta_IR', 'r', 'mu']

# Track per-config index to select correct theta
idx_per_config = {tuple(c): 0 for c in lconfigurations}

with open(output_file, mode='w', newline='') as outfile:
    writer = csv.writer(outfile)
    writer.writerow(header)
    for row in reader[1:]:
        configuration = (row[configuration_column],)
        index_val = ""
        theta_val = ""
        r_val = ""
        mu_val = ""
        if configuration in rates and row[validation_column] == '1':
            i = idx_per_config[configuration]
            theta_list = np.array(theta_values[configuration]['theta_list'])
            r_list = rates[configuration]
            index_list = all_indexes[configuration]
            if i < len(theta_list):
                index_val = index_list[i]
                theta_val = theta_list[i]
                r_val = r_list[i]
                mu_val = theta_val * r_val
                idx_per_config[configuration] += 1
        writer.writerow([index_val, theta_val, r_val, mu_val])

print(f"OK Saved file to {output_file}")
```

```{python}
# BS Model This code builds a csv file with the data. The last column corresponds to theta

with open('Analysis_output/tab_kinetic.csv', newline='') as file:
    data = list(csv.reader(file, delimiter=','))

def read_file(data, configuration, tags_range, configuration_column, validation_column, time_of_sampling_column, bacterial_load_column, sample_weight_column, sample_index_column, Ntypes):
    frequencies = []
    times = []
    loads = []
    weights = []
    indexes = []

    for row in data:
        if row[configuration_column] == configuration[0]: # Check condition
            if row[validation_column] == '1': # Check sequencing validation
                counts = [int(row[j]) for j in range(tags_range, Ntypes + tags_range)]  # Adjust for weight column
                counts_sum = sum(counts)
                if counts_sum > 0:
                    frequencies.append([c / counts_sum for c in counts])
                    times.append(float(row[time_of_sampling_column]))
                    loads.append(float(row[bacterial_load_column]))
                    weights.append(float(row[sample_weight_column]))
                    indexes.append(row[sample_index_column])

    return np.array(frequencies), np.array(times), np.array(loads), np.array(weights), np.array(indexes)


def load_pickle(filename):
    with open(filename, "rb") as f:
        return pickle.load(f)

# Load theta values
theta_values = pickle.load(open('Models_output_kinetic/theta_values4.pkl', 'rb'))

# Load precomputed rates (r) per config
# Here we recompute the rates so we can align them


# Read CSV into list
with open('Analysis_output/tab_kinetic.csv', newline='') as file:
    data = list(csv.reader(file))

# Compute r per valid row to match theta
# Compute rates for all configs
rates = {}
all_indexes = {}
for configuration in lconfigurations:
    config_key = tuple(configuration)
    frequencies, times, loads, weights, indexes = read_file(data, configuration, tags_range, configuration_column, validation_column, time_of_sampling_column, bacterial_load_column, sample_weight_column, sample_index_column, Ntypes)
    
    if config_key in theta_values:
        theta_list = np.array(theta_values[config_key]['theta_list'])
        p1 = np.max(frequencies, axis=1)
        r_ = np.full_like(p1, np.nan)
        mask = (p1>0) & (loads>0) & (times!=0) & (theta_list>0)
        r_[mask] = (np.log(p1[mask]*loads[mask]*weights[mask])+(1.0/theta_list[mask]))/times[mask]
        rates[config_key] = r_
        all_indexes[config_key] = indexes


# Prepare output file with header
input_file = "Analysis_output/tab_kinetic.csv"
output_file = "Models_output_kinetic/data_and_estimators_model_BS.csv"
with open(input_file, newline='') as infile:
    reader = list(csv.reader(infile))

# Header with only theta
header = ['index'] + ['1 / theta_BS']

# Track per-config index to select correct theta
idx_per_config = {tuple(c): 0 for c in lconfigurations}

with open(output_file, mode='w', newline='') as outfile:
    writer = csv.writer(outfile)
    writer.writerow(header)
    for row in reader[1:]:
        configuration = (row[configuration_column],)
        index_val = ""
        theta_val = ""  # default blank
        if configuration in theta_values and row[validation_column] == '1':
            i = idx_per_config[configuration]
            theta_list = np.array(theta_values[configuration]['theta_list'])
            index_list = all_indexes[configuration]
            if i < len(theta_list):
                index_val = index_list[i]
                theta_val = theta_list[i]
                idx_per_config[configuration] += 1
        writer.writerow([index_val] + [theta_val])

print(f"OK Saved file to {output_file}")

```

## Visualisation of models outputs

Visualisation of the output parameters theta, mu and r

```{r Graph}

tab = tab_kinetic %>%
  filter(Zone == "Root") %>%
  filter(Condition2 == "M_Intact_27") %>%
  filter(`PCR2 validation on gel` > 0)

tab_temp = read.csv("Models_output_kinetic/data_and_estimators_model_STAMP.csv", header= TRUE, check.names= FALSE)
tab$Theta_STAMP = tab_temp$theta_STAMP
tab_temp = read.csv("Models_output_kinetic/data_and_estimators_model_CR.csv", header= TRUE, check.names= FALSE)
tab$Theta_CR = tab_temp$theta_CR
tab$Mu_CR = tab_temp$mu
tab$r_CR = tab_temp$r
tab_temp = read.csv("Models_output_kinetic/data_and_estimators_model_BS.csv", header= TRUE, check.names= FALSE)
tab$Theta_BS = tab_temp$`1 / theta_BS`
tab$Theta_BS = 1 / tab$Theta_BS
tab_temp = read.csv("Models_output_kinetic/data_and_estimators_model_IR.csv", header= TRUE, check.names= FALSE)
tab$Theta_IR = tab_temp$theta
tab$Mu_IR = tab_temp$mu
tab$r_IR = tab_temp$r

tab$`Time post-inoculation (d)` = factor(tab$`Time post-inoculation (d)`, levels= unique(tab$`Time post-inoculation (d)`))


####################
# Visualisation of the STAMP model (Nb)
####################

# Kruskal-wallis statistical test
test = Graph_kruskal(tab= tab, y= "Theta_STAMP", x= "Time post-inoculation (d)", skip_h = TRUE)

# Display helper
values = tab %>% 
  group_by(`Time post-inoculation (d)`, .drop= FALSE) %>% 
  summarise(mean = mean(Theta_STAMP), sd = sd(Theta_STAMP))


# Graph
ggplot(data= tab, aes(x= `Time post-inoculation (d)`, y= Theta_STAMP)) +
  
  geom_boxplot(outlier.shape= NA, width=0.9, fill= "lightblue") +
  geom_point(position= position_jitter(width= 0.3), alpha= 1) +
  
  geom_text(data = values, aes(label = test$groups, y = 1 , x = `Time post-inoculation (d)`), vjust = -22, inherit.aes = FALSE) +
  scale_x_discrete(drop= FALSE) +
  scale_y_continuous(trans= "log10", limits= c(NA,NA)) +
  labs(title= "Instantaneous bottleneck STAMP model", y = "Bottleneck Nb (Number of bacteria)") +
  theme_gray()


####################
# Visualisation of the CR model (theta, mu and r)
####################

# Theta CR
# Kruskal-wallis statistical test
test = Graph_kruskal(tab= tab, y= "Theta_CR", x= "Time post-inoculation (d)", skip_h = TRUE)

# Display helper
values = tab %>% 
  group_by(`Time post-inoculation (d)`, .drop= FALSE) %>% 
  summarise(mean = mean(Theta_CR), sd = sd(Theta_CR))

# Graph theta CR
ggplot(data= tab, aes(x= `Time post-inoculation (d)`, y= Theta_CR)) +
  
  geom_boxplot(outlier.shape= NA, width=0.9, fill= "pink") +
  geom_point(position= position_jitter(width= 0.3), alpha= 1) +
  
  geom_text(data = values, aes(label = test$groups, y = 1 , x = `Time post-inoculation (d)`), vjust = -21.7, inherit.aes = FALSE) +
  scale_x_discrete(drop= FALSE) +
  scale_y_continuous(limits= c(NA,NA)) +
  labs(title= "Constant entry rate CR model", y = "Bottleneck effect mu/r (no units)") +
  theme_gray()


# Mu CR
# Kruskal-wallis statistical test
test = Graph_kruskal(tab= tab, y= "Mu_CR", x= "Time post-inoculation (d)", skip_h = TRUE)

# Display helper
values = tab %>% 
  group_by(`Time post-inoculation (d)`, .drop= FALSE) %>% 
  summarise(mean = mean(Mu_CR), sd = sd(Mu_CR))

# Graph mu CR
ggplot(data= tab, aes(x= `Time post-inoculation (d)`, y= Mu_CR), ) +
  
  geom_boxplot(outlier.shape= NA, width=0.9, fill= "pink") +
  geom_point(position= position_jitter(width= 0.3), alpha= 1) +
  
  geom_text(data = values, aes(label = test$groups, y = 1 , x = `Time post-inoculation (d)`), vjust = -21.7, inherit.aes = FALSE) +
  scale_x_discrete(drop= FALSE) +
  scale_y_continuous(limits= c(NA,NA)) +
  labs(title= "Constant entry rate CR model", y = "Permeability mu (CFU/d)") +
  theme_gray()


# r CR
# Kruskal-wallis statistical test
test = Graph_kruskal(tab= tab, y= "r_CR", x= "Time post-inoculation (d)", skip_h = TRUE)

# Display helper
values = tab %>% 
  group_by(`Time post-inoculation (d)`, .drop= FALSE) %>% 
  summarise(mean = mean(r_CR), sd = sd(r_CR))

# Graph r CR
ggplot(data= tab, aes(x= `Time post-inoculation (d)`, y= r_CR), ) +
  
  geom_boxplot(outlier.shape= NA, width=0.9, fill= "pink") +
  geom_point(position= position_jitter(width= 0.3), alpha= 1) +
  
  geom_text(data = values, aes(label = test$groups, y = 1 , x = `Time post-inoculation (d)`), vjust = -0.9, inherit.aes = FALSE) +
  scale_x_discrete(drop= FALSE) +
  scale_y_continuous(limits= c(NA,NA)) +
  labs(title= "Constant entry rate CR model", y = "Growth rate r (d-1)") +
  theme_gray()

####################
# Visualisation of the BS model (theta)
####################

# Theta BS
# Kruskal-wallis statistical test
test = Graph_kruskal(tab= tab, y= "Theta_BS", x= "Time post-inoculation (d)", skip_h = TRUE)

# Display helper
values = tab %>% 
  group_by(`Time post-inoculation (d)`, .drop= FALSE) %>% 
  summarise(mean = mean(Theta_BS), sd = sd(Theta_BS))

# Graph theta BS
ggplot(data= tab, aes(x= `Time post-inoculation (d)`, y= Theta_BS), ) +
  
  geom_boxplot(outlier.shape= NA, width=0.9, fill= "lightgreen") +
  geom_point(position= position_jitter(width= 0.3), alpha= 1) +
  
  geom_text(data = values, aes(label = test$groups, y = 1 , x = `Time post-inoculation (d)`), vjust = -22, inherit.aes = FALSE) +
  scale_x_discrete(drop= FALSE) +
  scale_y_continuous(trans= "log10", limits= c(NA,NA), breaks= c(1,3,5,10,20)) +
  labs(title= "Bolthausen-Sznitsman BS model", y = "Normalised entry speed v (no units)") +
  theme_gray()


####################
# Visualisation of the IR model (theta, mu and r)
####################

# Theta IR
# Kruskal-wallis statistical test
test = Graph_kruskal(tab= tab, y= "Theta_IR", x= "Time post-inoculation (d)", skip_h = TRUE)

# Display helper
values = tab %>% 
  group_by(`Time post-inoculation (d)`, .drop= FALSE) %>% 
  summarise(mean = mean(Theta_IR), sd = sd(Theta_IR))

# Graph theta IR
ggplot(data= tab, aes(x= `Time post-inoculation (d)`, y= Theta_IR), ) +
  
  geom_boxplot(outlier.shape= NA, width=0.9, fill= "tomato") +
  geom_point(position= position_jitter(width= 0.3), alpha= 1) +
  
  geom_text(data = values, aes(label = test$groups, y = 1 , x = `Time post-inoculation (d)`), vjust = -0.9, inherit.aes = FALSE) +
  scale_x_discrete(drop= FALSE) +
  scale_y_continuous(limits= c(NA,NA)) +
  labs(title= "Increasing entry rate IR model", y = "Bacterial capacity index mu/r (no units)") +
  theme_gray()


# Graph theta IR v2
ggplot(data= tab, aes(x= `Time post-inoculation (d)`, y= Theta_IR), ) +
  
  geom_boxplot(outlier.shape= NA, width=0.7) +
  geom_point(position= position_jitter(width= 0.3), alpha= 1) +
  
  geom_text(data = values, aes(label = test$groups, y = 1 , x = `Time post-inoculation (d)`), vjust = -0.9, inherit.aes = FALSE) +
  scale_x_discrete(drop= FALSE) +
  scale_y_continuous(limits= c(0,1.05)) +
  labs(x= "Organ", y = bquote("Bottleneck effect"~theta[IR]*" (no units)")) +
  theme_gray()


# Mu IR
# Kruskal-wallis statistical test
test = Graph_kruskal(tab= tab, y= "Mu_IR", x= "Time post-inoculation (d)", skip_h = TRUE)

# Display helper
values = tab %>% 
  group_by(`Time post-inoculation (d)`, .drop= FALSE) %>% 
  summarise(mean = mean(Mu_IR), sd = sd(Mu_IR))

# Graph mu IR
ggplot(data= tab, aes(x= `Time post-inoculation (d)`, y= Mu_IR), ) +
  
  geom_boxplot(outlier.shape= NA, width=0.9, fill= "tomato") +
  geom_point(position= position_jitter(width= 0.3), alpha= 1) +
  
  geom_text(data = values, aes(label = test$groups, y = 0.51 , x = `Time post-inoculation (d)`), vjust = 0, inherit.aes = FALSE) +
  scale_x_discrete(drop= FALSE) +
  scale_y_continuous(limits= c(NA,NA)) +
  labs(title= "Increasing entry rate IR model", y = "Permeability mu (CFU/d)") +
  theme_gray()

# Graph mu IR v2
ggplot(data= tab, aes(x= `Time post-inoculation (d)`, y= Mu_IR), ) +
  
  geom_boxplot(outlier.shape= NA, width=0.8) +
  geom_point(position= position_jitter(width= 0.3), alpha= 1) +
  
  geom_text(data = values, aes(label = test$groups, y = 0.51 , x = `Time post-inoculation (d)`), vjust = 0, inherit.aes = FALSE) +
  scale_x_discrete(drop= FALSE) +
  scale_y_continuous(limits= c(NA,NA)) +
  labs(x= "Organ", y = bquote("Apparent entry rate"~mu[IR]*" (no units)")) +
  theme_gray()


# r IR
# Kruskal-wallis statistical test
test = Graph_kruskal(tab= tab, y= "r_IR", x= "Time post-inoculation (d)", skip_h = TRUE)

# Display helper
values = tab %>% 
  group_by(`Time post-inoculation (d)`, .drop= FALSE) %>% 
  summarise(mean = mean(r_IR), sd = sd(r_IR))

# Graph r IR
ggplot(data= tab, aes(x= `Time post-inoculation (d)`, y= r_IR), ) +
  
  geom_boxplot(outlier.shape= NA, width=0.9, fill= "tomato") +
  geom_point(position= position_jitter(width= 0.3), alpha= 1) +
  
  geom_text(data = values, aes(label = test$groups, y = 0.81 , x = `Time post-inoculation (d)`), vjust = 0, inherit.aes = FALSE) +
  scale_x_discrete(drop= FALSE) +
  scale_y_continuous(limits= c(NA,NA)) +
  labs(title= "Increasing entry rate IR model", y = "Growth rate r (h-1)") +
  theme_gray()

# Graph r IR v2
ggplot(data= tab, aes(x= `Time post-inoculation (d)`, y= r_IR), ) +
  
  geom_boxplot(outlier.shape= NA, width=0.9) +
  geom_point(position= position_jitter(width= 0.3), alpha= 1) +
  
  geom_text(data = values, aes(label = test$groups, y = 0.81 , x = `Time post-inoculation (d)`), vjust = 0, inherit.aes = FALSE) +
  scale_x_discrete(drop= FALSE) +
  scale_y_continuous(limits= c(NA,NA)) +
  labs(x= "Organ", y = bquote("Apparent In vivo growth rate"~r[IR]*" ("~h^-1*")")) +
  theme_gray()  

```

## Classical growth rate with an exponential linear regression versus estimated models' parameters

```{r Graph}

# Loading table and adding models' output parameters
tab = tab_kinetic %>%
  filter(Zone == "Root") %>%
  filter(Condition2 == "M_Intact_27")

tab_temp = read.csv("Models_output_kinetic/data_and_estimators_model_STAMP.csv", header= TRUE, check.names= FALSE)
colnames(tab_temp)[colnames(tab_temp) == "theta_STAMP"] = "Theta_STAMP"

tab_temp = merge(tab_temp, read.csv("Models_output_kinetic/data_and_estimators_model_CR.csv", header= TRUE, check.names= FALSE))
colnames(tab_temp)[colnames(tab_temp) == "theta_CR"] = "Theta_CR"
colnames(tab_temp)[colnames(tab_temp) == "r"] = "r_CR"
colnames(tab_temp)[colnames(tab_temp) == "mu"] = "Mu_CR"

tab_temp = merge(tab_temp, read.csv("Models_output_kinetic/data_and_estimators_model_BS.csv", header= TRUE, check.names= FALSE))
colnames(tab_temp)[colnames(tab_temp) == "1 / theta_BS"] = "Theta_BS"

tab_temp = merge(tab_temp, read.csv("Models_output_kinetic/data_and_estimators_model_IR.csv", header= TRUE, check.names= FALSE))
colnames(tab_temp)[colnames(tab_temp) == "theta_IR"] = "Theta_IR"
colnames(tab_temp)[colnames(tab_temp) == "r"] = "r_IR"
colnames(tab_temp)[colnames(tab_temp) == "mu"] = "Mu_IR"

colnames(tab_temp)[colnames(tab_temp) == "index"] = "Sample ID"

tab = merge(tab, tab_temp, all.x= TRUE)


# Separating data availability
tab$`Data availability` = tab$`PCR2 validation on gel`
tab$`Data availability`[tab$`Data availability` > 0] = "CFU + sequences"
tab$`Data availability`[tab$`Data availability` == 0] = "CFU only"
tab$`Data availability`[is.na(tab$`Data availability`)] = "CFU only"


####################
# Bacterial load over time
####################

# Formating data for linear regression
tab_temp = data.frame(Zone= factor(), N0= numeric(), std_error_N0= numeric(),
                      Rate= numeric(), std_error_Rate= numeric(), Adj.rr= numeric())

a = summary(lm(data= filter(tab, Zone == "Root"), log(`Bacterial load (CFU/g)`) ~ as.integer(`Time post-inoculation (d)`)))
tab_temp = rbind(tab_temp, c("Root", a$coefficients[1], a$coefficients[3],
                             a$coefficients[2], a$coefficients[4], a$adj.r.squared))

names(tab_temp) = c("Zone", "N0", "std_error_N0", "Rate", "std_error_Rate", "Adj.rr")

tab_temp[,2:6] = lapply(tab_temp[,2:6], as.numeric)

tab_temp$Equation = paste(tab_temp$Zone,
                          "   y= exp(",round(tab_temp$Rate, 2), "t",
                          ifelse(tab_temp$N0 >= 0, "+",""),
                          round(tab_temp$N0, 2) , ")", sep= "")


# Graph v1 
ggplot(data= tab, aes(x= as.numeric(as.character(`Time post-inoculation (d)`)), y= `Bacterial load (CFU/g)`, colour= "blue")) +
  
  geom_point(aes(shape= `Data availability`)) +
  geom_smooth(method= "lm", formula= y~x, se= FALSE) +
  #stat_regline_equation(aes(label= paste(after_stat(eq.label), after_stat(rr.label), sep= "~~~~")), formula = I(log(10^y))~x) +
  annotate("text", colour= "blue", x=3, y= c(10e7),
           label= rev(paste(tab_temp$Equation, "   R²= ", round(as.numeric(tab_temp$Adj.rr), 2), sep= ""))) +
  
  scale_colour_manual(values= "blue") +
  scale_shape_manual(values= c(19,1)) +
  scale_y_continuous(trans= "log10",
                     breaks = trans_breaks("log10", function(x) 10^x),
                     labels = trans_format("log10", math_format(10^.x))) +
  labs(x= "Time post-inoculation (d)") +
  theme_gray() +
  theme(legend.position="bottom")


####################
# Usin the linear regression to calculate a bacterial load normalised over time and comparaison with growth rate from the IR model
####################

# Calculating a normalised load at t0 based on the parameter r=2.23 found by fitting an exponential growth on data
tab$`Normalised load t0 (CFU/g)` = tab$`Bacterial load (CFU/g)` / (exp(2.23 * as.numeric(tab$`Time post-inoculation (d)`)))


# Graph v2 
ggplot(data= tab, aes(x= "", y= r_IR)) +
  
  geom_boxplot() +
  geom_point() +
  
  stat_mean(colour= "red", size= 15, shape= 45) +
  
  labs(x= "All samples with CFU + sequence", y= "Estimated growth rate r_IR in the IR model") +
  theme_gray() +
  theme(legend.position="bottom")


# Multiple linear correlation
a = lm(log(`Normalised load t0 (CFU/g)`) ~ (Mu_IR + r_IR), data = tab)
print(summary(a))


```

# Section reanalysis of the experiment of kinetics of infection of tomato with correction with the starting inoculum frequencies

Launch this at the beginning of the session to load data

N.B.: don't forget to set up your Python environment in Rstudio.

```{python}

import numpy as np
import matplotlib.pyplot as plt
import csv
import os
import time
from matplotlib.animation import FuncAnimation
from matplotlib import pylab
import math
import pickle
import multiprocessing as mp
from scipy.stats import gamma
import sympy as sp
from scipy.stats import t
from scipy.optimize import minimize_scalar
import matplotlib.cm as cm
from scipy.stats import spearmanr

# Info retrieved from document "Analysis_output/tab_kinetic.csv" in the functions "read_file"
Ntypes = 120 # Number of different tags
tags_range = 55 # Starting column for tags
sample_index_column = 1 # Column with sample names
configuration_column = 17 # Column that contains all tested samples conditions
time_of_sampling_column = 17 # Column with DPI of sampling
bacterial_load_column = 33
sample_weight_column = 19
validation_column = 221 # Column to check that the sample is validated
inoculum_index_column = 214 # Index of the inoculum that was used for the infection
q_values = np.linspace(0, 8, 100)  # Range of q-values for Hill numbers


# Configurations
lconfigurations = [
    ['3'],
    ['4'],
    ['5'],
    ['6']
    ]

# Info regarding the inoculum table document "Analysis_output/tab_inoc.csv"
Ntypes = Ntypes # Samples and inoculum should have identical number of tags
corresponding_inoculum_index_column = 144 # Index of the inoculum that was used for the infection
corresponding_inoculum_tags_range = 146 # Starting column for tags in inoculum
inoculum_validation_column = 145 # Column to check that the inoculum is validated

```

# I. NULL

# II. Reanalysis of the kinetics with inoculum correction (do not relaunch unless necessary; outputs are already saved)

## Initialisation of the helper functions

```{python, fig.width=8, fig.height=6}
# @title Main functions common to all models. Launch before the estimation

def sample_theta(K, theta_range, starting_frequencies, distribution_model):
    theta = np.random.uniform(*theta_range)
    V = distribution_model(K, theta, starting_frequencies)
    return V / np.sum(V)

def quantile_distance(theta_range, frequencies, starting_frequencies, distribution_model, parameter_name="theta", Nsim=100, quantile=0.05):
    K = len(frequencies)
    Q = np.arange(0, 4.1, 0.1)

    def hill_number(V, q):
        if q == 1:
            return np.exp(-np.sum(V[V > 0] * np.log(V[V > 0])))
        elif q == 0:
            return np.sum(V > 0)
        else:
            return (np.sum(V ** q)) ** (1 / (1 - q))

    def calculate_distance(V1, V2):
        DS = 0
        for q in Q:
            Hq1 = hill_number(V1, q)
            Hq2 = hill_number(V2, q)
            DS += abs(Hq1 - Hq2) / (Hq1 + Hq2)
        return DS / len(Q)

    distances = []
    for _ in range(Nsim):
        V = sample_theta(K, theta_range, starting_frequencies, distribution_model)
        D = calculate_distance(V, frequencies)
        distances.append(D)

    return np.quantile(distances, quantile)

def find_best_theta(frequencies, starting_frequencies, theta_range, distribution_model, distance_function, parameter_name="theta", step=1):
    distances = []
    theta_values = np.arange(theta_range[0], theta_range[1] + step, step)

    for theta in theta_values:
        dist = distance_function((theta, theta), frequencies, starting_frequencies, distribution_model, parameter_name)
        distances.append(dist)

    best_theta = theta_values[np.argmin(distances)]
    min_distance = min(distances)
    return best_theta, min_distance


def read_file(data, configuration, tags_range, configuration_column, validation_column, sample_index_column, time_of_sampling_column, bacterial_load_column, inoculum_index_column, Ntypes):
    frequencies = []
    conditions = []
    for row in data:
        if row[configuration_column] == configuration[0]: # Check condition
            if row[validation_column] == '1': # Check sequencing validation
                counts = [int(row[j]) for j in range(tags_range, Ntypes + tags_range)]
                counts_sum = sum(counts)
                if counts_sum > 0:
                    frequencies.append([c / counts_sum for c in counts])
                    time = float(row[17])
                    conditions.append((row[sample_index_column], int(row[time_of_sampling_column]),float(row[bacterial_load_column]), row[inoculum_index_column]))

    return np.array(frequencies), np.array(conditions)


def read_starting_freq(inoc, corresponding_inoculum_tags_range, corresponding_inoculum_index_column, inoculum_validation_column, Ntypes):
    frequencies = []
    conditions = []
    for row in inoc:
      if row[inoculum_validation_column] == '1': # Check sequencing validation
          counts = [int(row[j]) for j in range(corresponding_inoculum_tags_range, Ntypes + corresponding_inoculum_tags_range)]
          counts_sum = sum(counts)
          if counts_sum > 0:
              frequencies.append([c / counts_sum for c in counts])
              time = float(row[17])
              conditions.append((row[corresponding_inoculum_index_column]))
            
    return np.array(frequencies), np.array(conditions)

def process_configurations(data, inoc, lconfigurations, distribution_model, theta_range, step, distance_function=quantile_distance, parameter_name="theta", model_name="Adaptable Frequency Model"):
    theta_values = {}  # Dictionary to store theta lists, mean values, and performance
    results = []
    
    starting_frequencies, inoculum_index = read_starting_freq(inoc, corresponding_inoculum_tags_range, corresponding_inoculum_index_column, inoculum_validation_column, Ntypes)

    for configuration in lconfigurations:
        frequencies, metadata = read_file(data, configuration, tags_range, configuration_column, validation_column, sample_index_column, time_of_sampling_column, bacterial_load_column, inoculum_index_column, Ntypes)
        
        if len(frequencies) == 0:
            print(f"No data for configuration: {configuration}")
            continue
        
        mle_theta = []
        dist_s = []
        
        if len(frequencies) > 0:
            for i, freq in enumerate(frequencies):
            
                # get the starting frequencies corresponding to the inoculum of the sample
                meta = metadata[i]
                start_freq = starting_frequencies[np.searchsorted(inoculum_index, meta[3])]
            
                best_theta, dist_ = find_best_theta(freq, start_freq, theta_range, distribution_model, distance_function, parameter_name, step)
                mle_theta.append(best_theta)
                dist_s.append(dist_)

        # Compute mean theta and mean distance across all replicates
        mean_mle_theta = np.mean(mle_theta)
        mean_distance = np.mean(dist_s)  # Performance metric

        # Store both all best theta values, mean theta, and mean performance
        theta_values[tuple(configuration)] = {
            "theta_list": mle_theta,  # List of all best theta values
            "mean_theta": mean_mle_theta,  # Mean of best theta values
            "mean_distance": mean_distance  # Mean distance (performance metric)
        }

        # Store and print results (only mean values)
        results.append((configuration, mean_mle_theta, mean_distance))
        print(f"Modèle: {model_name}, Configuration: {configuration}, Mean theta: {mean_mle_theta:.2f}, Performance (Mean Distance): {mean_distance:.4f}")

    return theta_values  # Return both detailed and summarized theta values
  

with open('Analysis_output/tab_kinetic.csv', newline='') as file:
    data = list(csv.reader(file, delimiter=','))

with open('Analysis_output/tab_inoc.csv', newline='') as file:
    inoc = list(csv.reader(file, delimiter=','))

# Functions to plot the Hill curves
def hill_number(V, q):
    """Compute Hill numbers for a given frequency vector V and order q."""
    if q == 1:
        return np.exp(-np.sum(V[V > 0] * np.log(V[V > 0])))
    elif q == 0:
        return np.sum(V > 0)
    else:
        return (np.sum(V ** q)) ** (1 / (1 - q))

def calculate_distance_per_q(V1, V2, Q):
    """Compute the diversity-based distance for each q."""
    distances = []
    for q in Q:
        Hq1 = hill_number(V1, q)
        Hq2 = hill_number(V2, q)
        distance = abs(Hq1 - Hq2) / (Hq1 + Hq2) if (Hq1 + Hq2) != 0 else 0
        distances.append(distance)
    return distances

def generate_best_frequency(K, theta, starting_frequencies, distribution_model, real_frequency, num_sim=100):
    """Generate 100 frequency vectors and return the closest one to the real frequency."""
    simulated_vectors = [sample_theta(K, (theta, theta), starting_frequencies, distribution_model) for _ in range(num_sim)]
    distances = [np.mean(calculate_distance_per_q(V, real_frequency, np.arange(0, 4.1, 0.1))) for V in simulated_vectors]
    best_index = np.argmin(distances)  # Index of lowest distance
    return simulated_vectors[best_index]  # Return the best-matching frequency

def plot_hill_curves(real_frequencies, best_simulated_frequencies):
    """Plot Hill curves for real and best simulated frequencies in separate subplots."""
    Q = np.arange(0, 4.1, 0.1)
    num_freqs = len(real_frequencies)
    color_map = plt.colormaps["tab10"]

    fig, axes = plt.subplots(1, 2, figsize=(12, 6), sharey=True)

    # Plot each real frequency in the left subplot
    for i, freq in enumerate(real_frequencies):
        hill_curve = [hill_number(freq, q) for q in Q]
        color = color_map(i / max(1, num_freqs - 1))
        axes[0].plot(Q, hill_curve, color=color, alpha=0.9, linewidth=2)

    # Plot each best simulated frequency in the right subplot
    for i, best_freq in enumerate(best_simulated_frequencies):
        hill_curve = [hill_number(best_freq, q) for q in Q]
        color = color_map(i / max(1, num_freqs - 1))
        axes[1].plot(Q, hill_curve, color=color, alpha=0.9, linewidth=2)

    # Formatting subplots
    axes[0].set_title("Observed Frequencies")
    axes[1].set_title("Best Simulated Frequencies")
    for ax in axes:
        ax.set_xlabel("q")
        ax.set_ylabel("Hill Number")
        ax.set_ylim(0, 120)

    plt.tight_layout()
    plt.show()

def process_with_plots(data, inoc, lconfigurations, distribution_model, theta_values):
    """Process configurations, generate the best frequency vectors, and plot Hill curves and global mean distance."""
    Q = np.arange(0, 4.1, 0.1)
    global_mean_distances = np.zeros(len(Q))  # Accumulate mean distances over all configurations
    total_samples = 0  # Count total frequencies processed
    all_configurations_mean_distances = [] # Mean distances for all individual configuration
    configuration_mean_distances = np.zeros(len(Q))  # Accumulate mean distances within each configuration
    configuration_samples = 0 # Count frequencies processed within each configuration

    all_real_frequencies = {}  # Store real frequencies for all configurations
    all_best_simulated_frequencies = {}  # Store best simulated frequencies for all configurations
    
    starting_frequencies, inoculum_index = read_starting_freq(inoc, corresponding_inoculum_tags_range, corresponding_inoculum_index_column, inoculum_validation_column, Ntypes)
    
    
    for configuration in lconfigurations:
        real_frequencies, metadata = read_file(data, configuration, tags_range, configuration_column, validation_column, sample_index_column, time_of_sampling_column, bacterial_load_column, inoculum_index_column, Ntypes)
        
        if len(real_frequencies) == 0:
            print(f"No data for configuration: {configuration}")
            continue

        K = len(real_frequencies[0])  # Number of species/types
        config_key = tuple(configuration)
        theta_list = theta_values.get(config_key, {}).get("theta_list", None)

        if theta_list is None or len(theta_list) != len(real_frequencies):
            print(f"Missing or mismatched theta values for configuration: {configuration}")
            continue

        best_simulated_frequencies = []
        i = 0
        
        if len(real_frequencies) > 0:
            for real_freq, theta in zip(real_frequencies, theta_list):
            
                # get the starting frequencies corresponding to the inoculum of the sample
                meta = metadata[i]
                start_freq = starting_frequencies[np.searchsorted(inoculum_index, meta[3])]
                i = i + 1
                
                best_freq = generate_best_frequency(K, theta, start_freq, distribution_model, real_freq)
                best_simulated_frequencies.append(best_freq)

                # Compute distance for this frequency
                distances = calculate_distance_per_q(real_freq, best_freq, Q)
                global_mean_distances += np.array(distances)
                total_samples += 1
            
                # Distance retriever for the configuration
                configuration_mean_distances += np.array(distances)
                configuration_samples += 1

        # Compute and append distance for the configuration
        all_configurations_mean_distances.append(configuration_mean_distances / configuration_samples)

        # Reset configuration distance and sample count
        configuration_mean_distances = np.zeros(len(Q))
        configuration_samples = 0
        
        # Store results for this configuration
        all_real_frequencies[config_key] = real_frequencies
        all_best_simulated_frequencies[config_key] = best_simulated_frequencies

        # Plot Hill curves for this configuration
        plot_hill_curves(real_frequencies, best_simulated_frequencies)

    # Compute final averaged distances
    if total_samples > 0:
        global_mean_distances /= total_samples

    # Plot global error figure
    plt.figure(figsize=(8, 6))
    plt.plot(Q, global_mean_distances, color="black", linewidth=2)
    plt.xlabel("q")
    plt.ylabel("Mean Distance")
    plt.title("Mean Distance Averaged Over All Configurations")
    plt.show()

    return Q, global_mean_distances, all_real_frequencies, all_best_simulated_frequencies, all_configurations_mean_distances

# functions to Save and load the results
def save_pickle(theta_values, filename="theta_values1.pkl"):
    """
    Save theta_values dictionary as a Pickle file.
    """
    with open(filename, "wb") as f:
        pickle.dump(theta_values, f)
    print(f"Saved to {filename}")

def load_pickle(filename="theta_values1.pkl"):
    """
    Load theta_values dictionary from a Pickle file.
    """
    with open(filename, "rb") as f:
        return pickle.load(f)

```

## Multinomial model

```{python}
# define the model
def distribution_modelM1(K, theta, starting_frequencies):
    Nb = theta
    return np.random.multinomial(Nb, starting_frequencies)
```

```{python}

# define the range for parameter values
theta_range = (1, 200)
step = 1

print('Model performance')
# Run the processing function
theta_values1 = process_configurations(data, inoc, lconfigurations, distribution_modelM1, theta_range, step, quantile_distance, parameter_name="Nb", model_name = "STAMP Model")
save_pickle(theta_values1,"Models_output_kinetic_correct_inoc/theta_values1.pkl")
```

```{python}
theta_values1 = load_pickle("Models_output_kinetic_correct_inoc/theta_values1.pkl")  # Loads it back
Q1, D1, RF, BF1, all_D1 = process_with_plots(data, inoc, lconfigurations, distribution_modelM1, theta_values1)
np.save("Models_output_kinetic_correct_inoc/Q1.npy", Q1)
np.save("Models_output_kinetic_correct_inoc/D1.npy", D1)
np.save("Models_output_kinetic_correct_inoc/all_D1.npy", all_D1)
save_pickle(RF, filename="Models_output_kinetic_correct_inoc/RF.pkl")
save_pickle(BF1, filename="Models_output_kinetic_correct_inoc/BF1.pkl")

```

## Constant rate model CR

```{python}
# define the model
def distribution_modelM3(K, theta, starting_frequencies):
    mu = theta
    r = 1
    T = 10
    # simulate entry times :
    tau = [ np.random.exponential( 1/mu ) ]
    while tau[-1] < T:
        tau += [ tau[-1] + np.random.exponential( 1/mu ) ]
    qt_prov = [ np.exp( r * (T-tau[i]) ) for i in range(len(tau)) ]
    res = sample_tags( qt_prov, K, starting_frequencies )
    return np.sort(res)[::-1]

def sample_tags(prop, Ntags, starting_frequencies):
    prop_tags = np.zeros( Ntags )
    for pp in prop:
        tag = np.random.choice( range(Ntags), p = starting_frequencies) 
        prop_tags[ tag ] += pp
    return prop_tags

```

```{python}
# define the range for parameter values
theta_range = (1, 50)
step = 1

print('Model performance')
# Run the processing function
theta_values3 = process_configurations(data, inoc, lconfigurations, distribution_modelM3, theta_range, step, quantile_distance, parameter_name="Nb", model_name = "CR1 Model")

save_pickle(theta_values3,"Models_output_kinetic_correct_inoc/theta_values3.pkl")
```

```{python}
theta_values3 = load_pickle("Models_output_kinetic_correct_inoc/theta_values3.pkl")
Q3, D3, RF, BF3, all_D3 = process_with_plots(data, inoc, lconfigurations, distribution_modelM3, theta_values3)
np.save("Models_output_kinetic_correct_inoc/Q3.npy", Q3)
np.save("Models_output_kinetic_correct_inoc/D3.npy", D3)
np.save("Models_output_kinetic_correct_inoc/all_D3.npy", all_D3)
save_pickle(BF3, filename="Models_output_kinetic_correct_inoc/BF3.pkl")
```

## Bolthausen-Sznitman coalescent model BS

```{python}
#**Step 1**: construct the initial random recursive tree.
def update_edges( l, k ):
    current = k
    while current != 0:
        current = l[current][0]
        l[current][2] += [k]


def init_bs( n ):
    l_edges = [ [0,[0],[]] ]
    new_edge = [0,[0],[]]
    for k in range(1,n):
        parent_new = np.random.randint(k)
        new_edge = [ parent_new, [k], []]
        l_edges += [ new_edge ]
        update_edges( l_edges, k )
    return l_edges
#**Step 2**: each edge is "lifted" at rate $1$ (in the terminology of *Berestycki 2009*).
def step_bs( tree, k ): # lift edge k of tree
    loss = 0
    parent = tree[k][0]
    tree[parent][1] += [k] + tree[k][2] # the edge is lifted to the parent
    for i in [k] + tree[k][2]:
        if tree[i][1][0] != -1:
            tree[i][1] = [-1] # unactivated
            loss += 1
    return tree, loss # tree : resulting tree; loss: number of blocks lost
#**Step 3:** the following function runs a Bolthausen-Sznitman coalescent for a time [t], starting from an initial tree [tree] with specified number of blocks [nblocks]).

def run_bs( nblocks, t, tree ):

    t_cur = np.random.exponential(1/nblocks)

    while t_cur < t and nblocks > 1:


        i = np.random.randint( 1, nblocks ) # choose the edge to be lifted
        edge = 0
        while i > -1:
            if tree[ edge ][1][0] != -1:
                edge += 1
                i -= 1
            else:
                edge += 1
        tree, loss = step_bs( tree, edge - 1 )
        nblocks -= loss

        t_cur += np.random.exponential(1/nblocks)

    return tree, nblocks


#run_bs(10000,0.5)
def display( tree ):
    for k in range( len(tree) ):
        if tree[k][1][0] != -1:
            print(tree[k][1], end='  ')
    print("")
def get_prop_bs( N, t ):
    tree,_ = run_bs( N, t, init_bs( N ) )
    prop = [] # get proportions
    for e in tree:
        if e[1][0] != -1:
            prop += [ len(e[1]) ]
    prop = prop / np.sum( prop )
    return prop
def sample_tags(prop, Ntags, starting_frequencies):
    prop_tags = np.zeros( Ntags )
    for pp in prop:
        tag = np.random.choice( range(Ntags), p = starting_frequencies) 
        prop_tags[ tag ] += pp
    return prop_tags
```

```{python}
# define the model
def distribution_modelM4(K, theta, starting_frequencies):
    t = theta
    return sample_tags( get_prop_bs(2000, t), K, starting_frequencies)
```

```{python}
# define the range for parameter values
theta_range = (0.05, 1)
step = 0.05

print('Model performance')
# Run the processing function
theta_values4 = process_configurations(data, inoc, lconfigurations, distribution_modelM4, theta_range, step, quantile_distance, parameter_name="t", model_name = "BS Model")
save_pickle(theta_values4,"Models_output_kinetic_correct_inoc/theta_values4.pkl")
```

```{python}
theta_values4 = load_pickle("Models_output_kinetic_correct_inoc/theta_values4.pkl")
Q4, D4, RF, BF4, all_D4 = process_with_plots(data, inoc, lconfigurations, distribution_modelM4, theta_values4)
np.save("Models_output_kinetic_correct_inoc/Q4.npy", Q4)
np.save("Models_output_kinetic_correct_inoc/D4.npy", D4)
np.save("Models_output_kinetic_correct_inoc/all_D4.npy", all_D4)
save_pickle(BF4, filename="Models_output_kinetic_correct_inoc/BF4.pkl")
```

## Kingman coalescent

```{python}
def run_km( t, prop = [] ):

    nblocks = len(prop)
    t_cur = 0

    if nblocks > 1:
        t_cur = np.random.exponential(2/(nblocks * (nblocks-1)))

    while t_cur < t and nblocks > 1:
        t_cur += np.random.exponential(2/(nblocks * (nblocks-1)))

        i,j=[0,0]
        while i == j:
            i,j = np.random.randint( 0, nblocks, 2)
        prop[i] += prop[j] # j gets to i
        prop.pop(j) # remove j
        nblocks -= 1
    return prop
def sample_tags(prop, Ntags, starting_frequencies):
    prop_tags = np.zeros( Ntags )
    for pp in prop:
        tag = np.random.choice( range(Ntags), p = starting_frequencies) 
        prop_tags[ tag ] += pp
    return prop_tags

```

```{python}
# define the model
def distribution_modelM5(K, theta, starting_frequencies):
    t = theta
    return sample_tags( run_km(t, [1/1000] * 1000), K, starting_frequencies)
```

```{python}
# define the range for parameter values
theta_range = (0.05, 1)
step = 0.05
print('Model performance')
# Run the processing function
theta_values5 = process_configurations(data, inoc, lconfigurations, distribution_modelM5, theta_range, step, quantile_distance, parameter_name="t", model_name = "K Model")
save_pickle(theta_values5,"Models_output_kinetic_correct_inoc/theta_values5.pkl")
```

```{python}
theta_values5 = load_pickle("Models_output_kinetic_correct_inoc/theta_values5.pkl")
Q5, D5, RF, BF5, all_D5 = process_with_plots(data, inoc, lconfigurations, distribution_modelM5, theta_values5)
np.save("Models_output_kinetic_correct_inoc/Q5.npy", Q5)
np.save("Models_output_kinetic_correct_inoc/D5.npy", D5)
np.save("Models_output_kinetic_correct_inoc/all_D5.npy", all_D5)
save_pickle(BF5, filename="Models_output_kinetic_correct_inoc/BF5.pkl")
```

## Increasing entry rate model IR

```{python}

# NHPP inverse cumulative intensity function for λ(t) = mu * exp(k * t)
def nhpp_exponential_inverse(s, mu, k):
    return (1.0 / k) * np.log(k * s / mu + 1)

def distribution_modelM6(K, theta, starting_frequencies):
    mu = theta
    k = 0.78
    Ntypes = K
    T = 10

    tau = []
    s = 0.0
    while True:
        s += np.random.exponential(1.0)  # Homogeneous Poisson interarrival
        t_candidate = nhpp_exponential_inverse(s, mu, k)
        if t_candidate > T:
            break
        tau.append(t_candidate)

    qt_prov = [np.exp((T - t)) for t in tau]

    res = sample_tags( qt_prov, K, starting_frequencies )

    return np.sort(res)[::-1]

def sample_tags(prop, Ntags, starting_frequencies):
    prop_tags = np.zeros( Ntags )
    for pp in prop:
        tag = np.random.choice( range(Ntags), p = starting_frequencies) 
        prop_tags[ tag ] += pp
    return prop_tags

```

```{python}
# define the range for parameter values
theta_range = (0.05, 1)
step = 0.05

print('Model performance')
# Run the processing function
theta_values6 = process_configurations(data, inoc, lconfigurations, distribution_modelM6, theta_range, step, quantile_distance, parameter_name="mu", model_name = "VR Model")
save_pickle(theta_values6,"Models_output_kinetic_correct_inoc/theta_values6.pkl")
```

```{python}
theta_values6 = load_pickle("Models_output_kinetic_correct_inoc/theta_values6.pkl")
Q6, D6, RF, BF6, all_D6 = process_with_plots(data, inoc, lconfigurations, distribution_modelM6, theta_values6)
np.save("Models_output_kinetic_correct_inoc/Q6.npy", Q6)
np.save("Models_output_kinetic_correct_inoc/D6.npy", D6)
np.save("Models_output_kinetic_correct_inoc/all_D6.npy", all_D6)
save_pickle(BF6, filename="Models_output_kinetic_correct_inoc/BF6.pkl")
```

# III. Some graphs

## Helper function

```{python, fig.width=10, fig.height=6}

# =======================
# Load pickle files
# =======================

def load_pickle(filename):
    if not os.path.exists(filename):
        raise FileNotFoundError(f"'{filename}' not found in repository.")
    with open(filename, "rb") as f:
        return pickle.load(f)

# Load data
RF = load_pickle("Models_output_kinetic_correct_inoc/RF.pkl")
BF1 = load_pickle("Models_output_kinetic_correct_inoc/BF1.pkl")
BF3 = load_pickle("Models_output_kinetic_correct_inoc/BF3.pkl")
BF4 = load_pickle("Models_output_kinetic_correct_inoc/BF4.pkl")
BF6 = load_pickle("Models_output_kinetic_correct_inoc/BF6.pkl")
theta_values = load_pickle("Models_output_kinetic_correct_inoc/theta_values1.pkl")  # Adjust the file name if needed

```

## Error curves

```{python, fig.width=8, fig.height=6}

# Initialize plot
plt.figure(figsize=(8, 6))

i = 1
Q = np.load(f"Models_output_kinetic_correct_inoc/Q{i}.npy")  # Load Qi
D = np.load(f"Models_output_kinetic_correct_inoc/D{i}.npy")  # Load Di
plt.plot(Q, D, linewidth=2, label="Model $\mathcal{{M}}_{STAMP}$")  # Plot with label
i = 3
Q = np.load(f"Models_output_kinetic_correct_inoc/Q{i}.npy")  # Load Qi
D = np.load(f"Models_output_kinetic_correct_inoc/D{i}.npy")  # Load Di
plt.plot(Q, D, linewidth=2, label="Model $\mathcal{{M}}_{CR}$")  # Plot with label
i = 4
Q = np.load(f"Models_output_kinetic_correct_inoc/Q{i}.npy")  # Load Qi
D = np.load(f"Models_output_kinetic_correct_inoc/D{i}.npy")  # Load Di
plt.plot(Q, D, linewidth=2, label="Model $\mathcal{{M}}_{BS}$")  # Plot with label
i = 6
Q = np.load(f"Models_output_kinetic_correct_inoc/Q{i}.npy")  # Load Qi
D = np.load(f"Models_output_kinetic_correct_inoc/D{i}.npy")  # Load Di
plt.plot(Q, D, linewidth=2, label="Model $\mathcal{{M}}_{IR}$")  # Plot with label


# Formatting the plot
plt.xlabel("$q$", fontsize=16)
plt.ylabel("Mean relative error", fontsize=16)
plt.legend(fontsize=14)
plt.xticks(fontsize=16)
plt.yticks(fontsize=16)
plt.grid()
plt.tight_layout()
#Save the figure as a PNG file
#plt.savefig("mean_error.png", dpi=300, bbox_inches='tight')

plt.show()
```

```{python, fig.width=6, fig.height=32}

# Plot mean error for each condition
# Load the saved data
RF = load_pickle("Models_output_kinetic_correct_inoc/RF.pkl")

# Extract experiment names from RF
experiment_names = list(RF.keys())

Q = np.arange(0, 4.1, 0.1)

# Create figure with subplots (7 rows, 1 columns)
fig, axes = plt.subplots(9, 1, figsize=(6, 32), sharex=True, sharey=True)

# Iterate over configurations and plot Hill curves
for j, exp_name in enumerate(experiment_names):

  i = 1
  D = np.load(f"Models_output_kinetic_correct_inoc/all_D{i}.npy")
  D = D[j]
  axes[j].plot(Q, D, alpha=0.8, linewidth=2, label="Model $\mathcal{{M}}_{STAMP}$")

  i = 3
  D = np.load(f"Models_output_kinetic_correct_inoc/all_D{i}.npy")
  D = D[j]
  axes[j].plot(Q, D, alpha=0.8, linewidth=2, label="Model $\mathcal{{M}}_{CR}$")

  i = 4
  D = np.load(f"Models_output_kinetic_correct_inoc/all_D{i}.npy")
  D = D[j]
  axes[j].plot(Q, D, alpha=0.8, linewidth=2, label="Model $\mathcal{{M}}_{BS}$")

  i = 6
  D = np.load(f"Models_output_kinetic_correct_inoc/all_D{i}.npy")
  D = D[j]
  axes[j].plot(Q, D, alpha=0.8, linewidth=2, label="Model $\mathcal{{M}}_{IR}$")

  #SV Set experiment label
  axes[j].set_ylabel(f"{exp_name}", fontsize=12)
  axes[j].grid()
  axes[j].legend(fontsize=14)

axes[j].set_xlabel("q", fontsize=12)

plt.tight_layout()
plt.show()

```

## Extraction of the csv tables with theta, mu and r

```{python}
# STAMP Model This code builds a csv file with the data. The last column corresponds to theta

with open('Analysis_output/tab_kinetic.csv', newline='') as file:
    data = list(csv.reader(file, delimiter=','))

def read_file(data, configuration, tags_range, configuration_column, validation_column, time_of_sampling_column, bacterial_load_column, sample_weight_column, sample_index_column, Ntypes):
    frequencies = []
    times = []
    loads = []
    weights = []
    indexes = []

    for row in data:
        if row[configuration_column] == configuration[0]: # Check condition
            if row[validation_column] == '1': # Check sequencing validation
                counts = [int(row[j]) for j in range(tags_range, Ntypes + tags_range)]  # Adjust for weight column
                counts_sum = sum(counts)
                if counts_sum > 0:
                    frequencies.append([c / counts_sum for c in counts])
                    times.append(float(row[time_of_sampling_column]))
                    loads.append(float(row[bacterial_load_column]))
                    weights.append(float(row[sample_weight_column]))
                    indexes.append(row[sample_index_column])

    return np.array(frequencies), np.array(times), np.array(loads), np.array(weights), np.array(indexes)


def load_pickle(filename):
    with open(filename, "rb") as f:
        return pickle.load(f)

# Load theta values
theta_values = pickle.load(open('Models_output_kinetic_correct_inoc/theta_values1.pkl', 'rb'))

# Load precomputed rates (r) per config
# Here we recompute the rates so we can align them


# Read CSV into list
with open('Analysis_output/tab_kinetic.csv', newline='') as file:
    data = list(csv.reader(file))

# Compute r per valid row to match theta
# Compute rates for all configs
rates = {}
all_indexes = {}
for configuration in lconfigurations:
    config_key = tuple(configuration)
    frequencies, times, loads, weights, indexes = read_file(data, configuration, tags_range, configuration_column, validation_column, time_of_sampling_column, bacterial_load_column, sample_weight_column, sample_index_column, Ntypes)
    
    if config_key in theta_values:
        theta_list = np.array(theta_values[config_key]['theta_list'])
        p1 = np.max(frequencies, axis=1)
        r_ = np.full_like(p1, np.nan)
        mask = (p1>0) & (loads>0) & (times!=0) & (theta_list>0)
        r_[mask] = (np.log(p1[mask]*loads[mask]*weights[mask])+(1.0/theta_list[mask]))/times[mask]
        rates[config_key] = r_
        all_indexes[config_key] = indexes


# Prepare output file with header
#input_file = "Models_output/Data_py.csv"
input_file = "Analysis_output/tab_kinetic.csv"
output_file = "Models_output_kinetic_correct_inoc/data_and_estimators_model_STAMP.csv"
with open(input_file, newline='') as infile:
    reader = list(csv.reader(infile))

# Header with only theta
header = ['index'] + ['theta_STAMP']

# Track per-config index to select correct theta
idx_per_config = {tuple(c): 0 for c in lconfigurations}

with open(output_file, mode='w', newline='') as outfile:
    writer = csv.writer(outfile)
    writer.writerow(header)
    for row in reader[1:]:
        configuration = (row[configuration_column],)
        index_val = ""
        theta_val = ""  # default blank
        if configuration in theta_values and row[validation_column] == '1':
            i = idx_per_config[configuration]
            theta_list = np.array(theta_values[configuration]['theta_list'])
            index_list = all_indexes[configuration]
            if i < len(theta_list):
                index_val = index_list[i]
                theta_val = theta_list[i]
                idx_per_config[configuration] += 1
        writer.writerow([index_val] + [theta_val])

print(f"OK Saved file to {output_file}")

```

```{python}
# CR Model This code builds a csv file with the data. The last three columns correspond to theta, r, mu

with open('Analysis_output/tab_kinetic.csv', newline='') as file:
    data = list(csv.reader(file, delimiter=','))

def read_file(data, configuration, tags_range, configuration_column, validation_column, time_of_sampling_column, bacterial_load_column, sample_weight_column, sample_index_column, Ntypes):
    frequencies = []
    times = []
    loads = []
    weights = []
    indexes = []

    for row in data:
        if row[configuration_column] == configuration[0]: # Check condition
            if row[validation_column] == '1': # Check sequencing validation
                counts = [int(row[j]) for j in range(tags_range, Ntypes + tags_range)]  # Adjust for weight column
                counts_sum = sum(counts)
                if counts_sum > 0:
                    frequencies.append([c / counts_sum for c in counts])
                    times.append(float(row[time_of_sampling_column]))
                    loads.append(float(row[bacterial_load_column]))
                    weights.append(float(row[sample_weight_column]))
                    indexes.append(row[sample_index_column])

    return np.array(frequencies), np.array(times), np.array(loads), np.array(weights), np.array(indexes)


def load_pickle(filename):
    with open(filename, "rb") as f:
        return pickle.load(f)

# Load theta values
theta_values = pickle.load(open('Models_output_kinetic_correct_inoc/theta_values3.pkl', 'rb'))

# Load precomputed rates (r) per config
# Here we recompute the rates so we can align them


# Read CSV into list
with open('Analysis_output/tab_kinetic.csv', newline='') as file:
    data = list(csv.reader(file))

# Compute r per valid row to match theta
# Compute rates for all configs
rates = {}
all_indexes = {}
for configuration in lconfigurations:
    config_key = tuple(configuration)
    frequencies, times, loads, weights, indexes = read_file(data, configuration, tags_range, configuration_column, validation_column, time_of_sampling_column, bacterial_load_column, sample_weight_column, sample_index_column, Ntypes)
    
    if config_key in theta_values:
        theta_list = np.array(theta_values[config_key]['theta_list'])
        p1 = np.max(frequencies, axis=1)
        r_ = np.full_like(p1, np.nan)
        mask = (p1>0) & (loads>0) & (times!=0) & (theta_list>0)
        r_[mask] = (np.log(p1[mask]*loads[mask]*weights[mask])+(1.0/theta_list[mask]))/times[mask]
        rates[config_key] = r_
        all_indexes[config_key] = indexes


# Prepare output file with header
#input_file = "Models_output/Data_py.csv"
input_file = "Analysis_output/tab_kinetic.csv"
output_file = "Models_output_kinetic_correct_inoc/data_and_estimators_model_CR.csv"
with open(input_file, newline='') as infile:
    reader = list(csv.reader(infile))

# Header with only theta
header = ['index', 'theta_CR', 'r', 'mu']

# Track per-config index to select correct theta
idx_per_config = {tuple(c): 0 for c in lconfigurations}

with open(output_file, mode='w', newline='') as outfile:
    writer = csv.writer(outfile)
    writer.writerow(header)
    for row in reader[1:]:
        configuration = (row[configuration_column],)
        index_val = ""
        theta_val = ""
        r_val = ""
        mu_val = ""
        if configuration in rates and row[validation_column] == '1':
            i = idx_per_config[configuration]
            theta_list = np.array(theta_values[configuration]['theta_list'])
            r_list = rates[configuration]
            index_list = all_indexes[configuration]
            if i < len(theta_list):
                index_val = index_list[i]
                theta_val = theta_list[i]
                r_val = r_list[i]
                mu_val = theta_val * r_val
                idx_per_config[configuration] += 1
        writer.writerow([index_val, theta_val, r_val, mu_val])

print(f"OK Saved file to {output_file}")

```

```{python}
# IR Model This code builds a csv file with the data. The last three columns correspond to theta, r, mu

k = 0.78

with open('Analysis_output/tab_kinetic.csv', newline='') as file:
````
