## Supplementary material for "Time-resolved bottlenecks reveal concurrent pathogen entry and growth during bacterial infection dynamics": Code 2: SV_Dorado_analysis.sh

```
#! /usr/bin/bash

# Pipeline for a metagenomic based analysis of STAMP obtained by Nanopore sequencing
# Author: Sylvain Vicente, LIPME, Toulouse, France, 06/2024

# Linux and Conda (Python) environment advised


#dorado download --model dna_r10.4.1_e8.2_400bps_sup@v5.0.0


# Get directory of script

SCRIPT_PATH="$(realpath "${BASH_SOURCE[0]}")"
SCRIPT_DIR="$(dirname "$SCRIPT_PATH")"

read -p "Directory of folder containing the pod5 folders" EXP_DIR

read -p "Run number" RUN_NUM

###########################################################################
### Basecalling
###########################################################################

mkdir -p "${SCRIPT_DIR}/Run$RUN_NUM"

dorado basecaller \
  --min-qscore 10 \
  --recursive \
  --emit-sam \
  --kit-name EXP-PBC096 \
  --no-trim \
  "/mnt/GRP_RAP/GRP_RAP/svicente/dna_r10.4.1_e8.2_400bps_sup@v5.0.0/" \
  $EXP_DIR > "${SCRIPT_DIR}/Run$RUN_NUM/00_Total_reads.sam"

dorado demux \
  --kit-name EXP-PBC096 \
  --emit-fastq \
  --emit-summary \
  --output-dir "${SCRIPT_DIR}/Run$RUN_NUM/02_Uncompressed_fastq" \
  "${SCRIPT_DIR}/Run$RUN_NUM/00_Total_reads.sam"
```
