## Supplementary material for "Time-resolved bottlenecks reveal concurrent pathogen entry and growth during bacterial infection dynamics": Code 2: SV_STAMP_nanopore.sh

```
#! /usr/bin/bash

# Pipeline for a metagenomic based analysis of STAMP obtained by Nanopore sequencing
# Author: Sylvain Vicente, LIPME, Toulouse, France, 06/2024

# Linux and Conda (Python) environment advised
# /!\ You should activate your Qiime2 Conda environment prior to the use of this script


###########################################################################
### Initial conda installation
###########################################################################
#
#conda install wget
#
#wget https://raw.githubusercontent.com/qiime2/distributions/dev/latest/passed/qiime2-amplicon-ubuntu-latest-conda.yml
#
#conda env create -n SV_STAMP_nanopore --file qiime2-amplicon-ubuntu-latest-conda.yml
#
#rm qiime2-amplicon-ubuntu-latest-conda.yml
#
#conda activate SV_STAMP_nanopore
#
#cd
#htop


###########################################################################

# Get directory of script

SCRIPT_PATH="$(realpath "${BASH_SOURCE[0]}")"
SCRIPT_DIR="$(dirname "$SCRIPT_PATH")"

read -p "Run number (write as #)" RUN_NUM

READS_DIR="$SCRIPT_DIR/Run$RUN_NUM/02_Uncompressed_fastq"
OUTPUT_DIR=$READS_DIR

read -p "Sequences basecalled using SV_Dorado_analysis.sh? If yes, code will skip barcode fastq retrieval and uncompressing (y/n)" SKIP1

if [ "$SKIP1" == "n" ]
then
read -p "Target fastq pass directory" READS_DIR
fi

read -p "Skip sequence filtering steps? If yes, skip filtering steps and start directly from 05_Dereplicated_sequences.qza (y/n)" SKIP2

read -p "Internal barcode used (write as #)" BARCODE_NUM

IBARCODE="Internal_Barcode_$BARCODE_NUM"

read -p "Vsearch alignement quality on internal barcode to identify run identity?" QTHRESHOLD1

read -p "Minimum coverage on reference to accept a read" COVERAGE1

read -p "Vsearch alignement quality on STAMP tag clustering?" QTHRESHOLD2


###########################################################################
### Concatenatation of the obtained fastq.gz into a single file for each Nanopore barcode (sample)
###########################################################################

###########################################################################
# Loop to find Nanopore identifier in the specified and concatenate them in a created folder "01_Concatenated"

if [ "$SKIP1" == "n" ]
then

mkdir -p "${SCRIPT_DIR}/Run$RUN_NUM/01_Concatenated"

OUTPUT_DIR="${SCRIPT_DIR}/Run$RUN_NUM/01_Concatenated"

for ((i=1; i<=96; i++)); do
    identifier=$(printf "barcode%02d" $i)
    find $READS_DIR -type f -name "*$identifier*" -exec cat {} + > $OUTPUT_DIR/concatenated_output_$identifier.fastq.gz
done


# Unzip fastq.gz to fastq in a folder "02_Uncompressed_fastq"

mkdir -p "${SCRIPT_DIR}/Run$RUN_NUM/02_Uncompressed_fastq"

READS_DIR=$OUTPUT_DIR
OUTPUT_DIR="${SCRIPT_DIR}/Run$RUN_NUM/02_Uncompressed_fastq"


for ((i=1; i<=96; i++)); do
    identifier=$(printf "barcode%02d" $i)
    find $READS_DIR -type f -name "*$identifier*" -exec gzip -d -c {} + > $OUTPUT_DIR/$identifier.fastq
done
fi

###########################################################################
# Filter sequences to keep only sequences between 160pb and 200pb in a folder "03_Filtered_fastq"
# This filter corresponds to the selection of the expected final length after trimming barcodes
# Reads under 160 are expected to be too small to contain all information

if [ "$SKIP2" == "n" ]
then

if [ "$SKIP1" == "y" ]
then
for ((i=1; i<=96; i++)); do
    identifier=$(printf "barcode%02d" $i)
    
    find "$READS_DIR" -type f -name "*$identifier*" | while read -r file; do
        
        mv "$file" "$OUTPUT_DIR/${identifier}.fastq"
    done
done
fi

mkdir -p "${SCRIPT_DIR}/Run$RUN_NUM/03_Filtered_fastq"

READS_DIR=$OUTPUT_DIR
OUTPUT_DIR="${SCRIPT_DIR}/Run$RUN_NUM/03_Filtered_fastq"

for ((i=1; i<=96; i++)); do
    identifier=$(printf "barcode%02d" $i)
    find $READS_DIR -type f -name "*$identifier*" -exec awk -v max_length=200 -v min_length=160 -v identifier="$identifier" '
    BEGIN {
        line_number = 1
        header_line = ""
        quality_line = ""
        plus_line = ""
        dna_line = ""
    }
    {
        if (line_number % 4 == 1) {
            header_line = $0
        }
        else if (line_number % 4 == 2) {
            dna_line = $0
        }
        else if (line_number % 4 == 3) {
            plus_line = $0
        }
        else if (line_number % 4 == 0) {
            quality_line = $0
            if (length(dna_line) <= max_length && length(dna_line) >= min_length) {
                print header_line
                print dna_line
                print plus_line
                print quality_line
            }
        }
        line_number++
    }
    ' {} + > $OUTPUT_DIR/Filtered_$identifier.fastq
done


###########################################################################
### Metagenomic pipeline using Qiime2 suite of Python packages
###########################################################################

###########################################################################
# Import sequences to Qiime2 in a file "04_Imported_fastq_artifact.qza" using a manually made manifest

mkdir -p "${SCRIPT_DIR}/Run$RUN_NUM/Quality_reports"

qiime tools import \
  --type 'SampleData[SequencesWithQuality]' \
  --input-path "$SCRIPT_DIR/Manifest1_Run$RUN_NUM.tsv" \
  --output-path $SCRIPT_DIR/Run$RUN_NUM/04_Filtered_demux.qza \
  --input-format SingleEndFastqManifestPhred33V2

qiime tools import \
  --type 'SampleData[SequencesWithQuality]' \
  --input-path "$SCRIPT_DIR/Manifest2_Run$RUN_NUM.tsv" \
  --output-path $SCRIPT_DIR/Run$RUN_NUM/04_Non_filtered_demux.qza \
  --input-format SingleEndFastqManifestPhred33V2

# Quality check step 04: size filtered reads vs non filtered reads

qiime demux summarize \
  --i-data $SCRIPT_DIR/Run$RUN_NUM/04_Filtered_demux.qza \
  --p-n 10000 \
  --o-visualization  ${SCRIPT_DIR}/Run$RUN_NUM/Quality_reports/04_Quality_summary_filtered.qzv

qiime demux summarize \
  --i-data $SCRIPT_DIR/Run$RUN_NUM/04_Non_filtered_demux.qza \
  --p-n 10000 \
  --o-visualization  ${SCRIPT_DIR}/Run$RUN_NUM/Quality_reports/04_Quality_summary_non_filtered.qzv


###########################################################################
# Dereplicatation of the sequences in a file "05_Dereplicated_sequences.qza" with its corresponding table of occurence "05_Table.qza"

qiime vsearch dereplicate-sequences \
  --i-sequences $SCRIPT_DIR/Run$RUN_NUM/04_Filtered_demux.qza \
  --o-dereplicated-table $SCRIPT_DIR/Run$RUN_NUM/05_Table.qza \
  --o-dereplicated-sequences $SCRIPT_DIR/Run$RUN_NUM/05_Dereplicated_sequences.qza

fi

mkdir -p "${SCRIPT_DIR}/Run$RUN_NUM/Quality_reports"

###########################################################################
# Import reference sequences containing internal barcodes, these barcodes are used to descriminate reads from different runs performed on a same sequencing chip
# Classify internal barcode consensus for each read and filtering out all wrong or ambiguous internal barcodes in "06_Filtered_sequences.qza"

qiime tools import \
  --input-path $SCRIPT_DIR/Internal_barcode_reads.fasta \
  --output-path $SCRIPT_DIR/Run$RUN_NUM/06_Internal_barcodes.qza \
  --type 'FeatureData[Sequence]'

qiime tools import \
  --type 'FeatureData[Taxonomy]' \
  --input-format HeaderlessTSVTaxonomyFormat \
  --input-path $SCRIPT_DIR/Internal_barcode_taxonomy.tsv \
  --output-path $SCRIPT_DIR/Run$RUN_NUM/06_Run_taxonomy.qza

qiime feature-classifier classify-consensus-vsearch \
  --i-query $SCRIPT_DIR/Run$RUN_NUM/05_Dereplicated_sequences.qza \
  --i-reference-reads $SCRIPT_DIR/Run$RUN_NUM/06_Internal_barcodes.qza \
  --i-reference-taxonomy $SCRIPT_DIR/Run$RUN_NUM/06_Run_taxonomy.qza \
  --p-maxaccepts 1 \
  --p-perc-identity $QTHRESHOLD1 \
  --p-query-cov $COVERAGE1 \
  --p-strand 'both' \
  --p-top-hits-only TRUE \
  --p-maxhits 1 \
  --o-classification $SCRIPT_DIR/Run$RUN_NUM/06_Seq_classification.qza \
  --o-search-results $SCRIPT_DIR/Run$RUN_NUM/06_Taxonomy_search_result.qza

qiime taxa filter-seqs \
  --i-sequences $SCRIPT_DIR/Run$RUN_NUM/05_Dereplicated_sequences.qza \
  --i-taxonomy $SCRIPT_DIR/Run$RUN_NUM/06_Seq_classification.qza \
  --p-include $IBARCODE \
  --o-filtered-sequences $SCRIPT_DIR/Run$RUN_NUM/06_Filtered_sequences.qza

qiime taxa filter-table \
  --i-table $SCRIPT_DIR/Run$RUN_NUM/05_Table.qza \
  --i-taxonomy $SCRIPT_DIR/Run$RUN_NUM/06_Seq_classification.qza \
  --p-include $IBARCODE \
  --o-filtered-table $SCRIPT_DIR/Run$RUN_NUM/06_Filtered_table.qza

# Quality check step 06: Proportion of eliminated reads at run recognition

qiime taxa barplot \
  --i-table $SCRIPT_DIR/Run$RUN_NUM/05_Table.qza \
  --i-taxonomy $SCRIPT_DIR/Run$RUN_NUM/06_Seq_classification.qza \
  --o-visualization $SCRIPT_DIR/Run$RUN_NUM/Quality_reports/06_Recognised_reads_barplot.qzv

qiime taxa collapse \
  --i-table $SCRIPT_DIR/Run$RUN_NUM/05_Table.qza \
  --i-taxonomy $SCRIPT_DIR/Run$RUN_NUM/06_Seq_classification.qza \
  --p-level 1 \
  --o-collapsed-table $SCRIPT_DIR/Run$RUN_NUM/06_Classification_table.qza

qiime feature-table summarize \
  --i-table $SCRIPT_DIR/Run$RUN_NUM/06_Classification_table.qza \
  --o-visualization $SCRIPT_DIR/Run$RUN_NUM/Quality_reports/06_Quality_summary_run.qzv

qiime metadata tabulate \
  --m-input-file $SCRIPT_DIR/Run$RUN_NUM/06_Classification_table.qza \
  --o-visualization $SCRIPT_DIR/Run$RUN_NUM/Quality_reports/06_Validated_reads_count.qzv


###########################################################################
# Import reference reads of the 120 STAMP tags
# Classify STAMP tag consensus for each read

qiime tools import \
  --input-path $SCRIPT_DIR/STAMP123_tags.fasta \
  --output-path $SCRIPT_DIR/Run$RUN_NUM/07_STAMP_tags.qza \
  --type 'FeatureData[Sequence]'

qiime tools import \
  --type 'FeatureData[Taxonomy]' \
  --input-format HeaderlessTSVTaxonomyFormat \
  --input-path $SCRIPT_DIR/STAMP123_tags_taxonomy.tsv \
  --output-path $SCRIPT_DIR/Run$RUN_NUM/07_Tags_taxonomy.qza

#qiime feature-classifier classify-consensus-vsearch \
#  --i-query $SCRIPT_DIR/06_Filtered_sequences.qza \
#  --i-reference-reads $SCRIPT_DIR/07_STAMP_tags.qza \
#  --i-reference-taxonomy $SCRIPT_DIR/07_Tags_taxonomy.qza \
#  --p-maxaccepts 1 \
#  --p-perc-identity 0.85 \
#  --p-query-cov 0.60 \
#  --p-strand 'both' \
#  --p-top-hits-only TRUE \
#  --p-maxhits 1 \
#  --o-classification $SCRIPT_DIR/07_Seq_classification.qza \
#  --o-search-results $SCRIPT_DIR/07_Taxonomy_search_result.qza

qiime vsearch cluster-features-closed-reference \
  --i-sequences $SCRIPT_DIR/Run$RUN_NUM/06_Filtered_sequences.qza \
  --i-table $SCRIPT_DIR/Run$RUN_NUM/06_Filtered_table.qza \
  --i-reference-sequences $SCRIPT_DIR/Run$RUN_NUM/07_STAMP_tags.qza \
  --p-perc-identity $QTHRESHOLD2 \
  --p-strand 'both' \
  --o-clustered-table $SCRIPT_DIR/Run$RUN_NUM/07_Table.qza \
  --o-clustered-sequences $SCRIPT_DIR/Run$RUN_NUM/07_Matched_reads.qza \
  --o-unmatched-sequences $SCRIPT_DIR/Run$RUN_NUM/07_Unmatched_reads.qza

# Quality check step 07: Tag attribution 

qiime taxa barplot \
  --i-table $SCRIPT_DIR/Run$RUN_NUM/07_Table.qza \
  --o-visualization $SCRIPT_DIR/Run$RUN_NUM/Quality_reports/07_Tag_clustering_barplot.qzv

qiime feature-table summarize \
  --i-table $SCRIPT_DIR/Run$RUN_NUM/07_Table.qza \
  --o-visualization $SCRIPT_DIR/Run$RUN_NUM/Quality_reports/07_STAMP_Tags_summary.qzv
  
qiime metadata tabulate \
  --m-input-file $SCRIPT_DIR/Run$RUN_NUM/07_Table.qza \
  --o-visualization $SCRIPT_DIR/Run$RUN_NUM/Quality_reports/07_STAMP_Tags_count.qzv


###########################################################################
# Summarising STAMP tags count
# File "08_Tag_clustering_barplot.qzv" for visualisation of STAMP tags abundances

qiime feature-table summarize-plus \
  --i-table $SCRIPT_DIR/Run$RUN_NUM/07_Table.qza \
  --o-feature-frequencies $SCRIPT_DIR/Run$RUN_NUM/08_Final_freq_table.qza \
  --o-sample-frequencies $SCRIPT_DIR/Run$RUN_NUM/08_Final_sample_freq_table.qza \
  --o-summary $SCRIPT_DIR/Run$RUN_NUM/Quality_reports/08_Final_summary.qzv
```
