## Supplementary material for "Time-resolved bottlenecks reveal concurrent pathogen entry and growth during bacterial infection dynamics": Code 2: ReadMe.txt

06/2024

# Pipeline for the analysis of a neutral barcode collection in R. pseudosolanacearum sequenced in Nanopore. Linux and Conda (Python) environment advised.

This code is distributed under a Creative Commons CC BY 4.0 license (https://creativecommons.org/licenses/by/4.0/). You are free to share and adapt the code for any uses. If this code substantially participated your work, please consider citing the corresponding article.

This code was written by Sylvain Vicente.

# How to use

The Shell scripts "SV_Dorado_analysis.sh" and "SV_STAMP_nanopore.sh" control the code for the analysis of the raw .pod5 data obtained in Nanopore through a Qiime2 pipeline for metabarcoding analysis. The scripts also contain additional annotations regarding how to set up the Python environnent for the pipeline.

The first script "SV_Dorado_analysis.sh" launches the Nanopore Dorado basecaller on your raw .pod5. The default settings of the basecaller is the super accurate model "dna_r10.4.1_e8.2_400bps_sup@v5.0.0" with a minimum Q score of 10 to accept reads. A prompt will ask you for the location of the .pod5 folder and will ask for the number index of the run you want to analyse. The script will then create a new folder "Run#" according to the run index and will demultiplex the samples in a subfolder called "02_Uncompressed_fastq".

The second script "SV_STAMP_nanopore.sh" launches the metabarcoding analysis on the demultiplexed reads. The script will ask you the number index of the run you want to analyse. First, you need to specify the path of your filtered reads by modifying the files "Manifest1_RunX.tsv" and "Manifest2_RunX.tsv". When this is done, you can launch the script and it will then ask the following prompts:
- "Sequences basecalled using SV_Dorado_analysis.sh? If yes, code will skip barcode fastq retrieval and uncompressing (y/n)"
If you choose "y", the script will automatically start the analysis using the subfolder "02_Uncompressed_fastq", if no then you will be prompted for the location of the fastq files.

- "Skip sequence filtering steps? If yes, skip filtering steps and start directly from 05_Dereplicated_sequences.qza (y/n)"
Use "y" to restart an analysis without having to repeat the first filtering steps, pick "n" if it is your first time analysing the data.

- "Internal barcode used (write as #)"
The number of the internal barcode used for the run (the PCR1 barcode). All reads are aligned against a closed reference that contain all PCR1 barcodes used in the study. This steps allows to filter out potential cross-contaminations between runs performed on a same flowcell.

"Vsearch alignement quality on internal barcode to identify run identity?"
Minimum quality to keep a read when aligned on the PCR1 internal barcode. Used value: 0.85

"Minimum coverage on reference to accept a read"
Minimum coverage of to accept a read when mapped against the 123-tags collection. Whatever the selected value, ambiguous reads are by default filtered out. Used value: 0.3

"Vsearch alignement quality on STAMP tag clustering?"
Minimum quality to keep a read when aligned on the the 123-tags collection. Used value: 0.85

# After the analysis

Quality reports will be generated at different steps of the pipeline. In order to retrieve the reads table and the quality reports. Open https://view.qiime2.org/ to use the Qiime2 visualisation tool.

- Retrieve "04_Quality_summary_non_filtered.qzv" as a .tsv for the initial reads counts per samples.
- Retrieve "04_Quality_summary_filtered.qzv" as a .tsv for the length-validated reads counts.
- Retrieve "06_Recognised_reads_barplot.qzv" as a .csv for the validated reads counts based on the internal barcode (PCR1 barcode).
- Retrieve "07_Tag_clustering_barplot.qzv" as a .csv for the final 123-tags counts.

# N.B.

Manifest files and quality reports for Run2 have been left in the folder as examples.
