## Supplementary material for "Time-resolved bottlenecks reveal concurrent pathogen entry and growth during bacterial infection dynamics": code_tikz_figure.tex

\begin{figure}[h]
  \begin{center}
    \begin{tikzpicture}
      \node (0) at (0,0) {1};
      \node (2) at (-1,-1) {2};
      \node (3) at (-1,-2) {3};
      \node (4) at (-1,-3) {4};
      \node (5) at (1,-1) {5};
      \node (6) at (1,-2) {6};
      \node (7) at (1,-3) {7};
      \draw[-] (0) -- (2) ;
      \draw[-] (0) -- (5) ;
      \draw[very thick] (2) -- (3) ;
      \draw[-] (3) -- (4) ;
      \draw[-] (5) -- (6) ;
      \draw[-] (6) -- (7) ;
    \end{tikzpicture}
    \qquad
    \begin{tikzpicture}
      \node (0) at (0,0) {1};
      \node (2) at (-1,-1) {2,3,4};
      \node (5) at (1,-1) {5};
      \node (6) at (1,-2) {6};
      \node (7) at (1,-3) {7};
      \draw[-] (0) -- (2) ;
      \draw[-] (0) -- (5) ;
      \draw[-] (5) -- (6) ;
      \draw[very thick] (6) -- (7) ;
    \end{tikzpicture}
    \qquad
    \begin{tikzpicture}
      \node (0) at (0,0) {1};
      \node (2) at (-1,-1) {2,3,4};
      \node (5) at (1,-1) {5};
      \node (6) at (1,-2) {6,7};
      \draw[-] (0) -- (2) ;
      \draw[-] (0) -- (5) ;
      \draw[very thick] (5) -- (6) ;
      \node (align) at (0,-3) {};
    \end{tikzpicture}    
    \qquad
    \begin{tikzpicture}
      \node (0) at (0,0) {1};
      \node (2) at (-1,-1) {2,3,4};
      \node (5) at (1,-1) {5,6,7};
      \draw[-] (0) -- (2) ;
      \draw[-] (0) -- (5) ;
      \node (align) at (0,-3) {};
    \end{tikzpicture}
  \end{center}
\end{figure}
